# Non-exonic read content reorganises lncRNA co-expression networks in renal and other cancers

**DOI:** 10.64898/2026.10.09.757827

**Authors:** Gehad Youssef, Namshik Han

## Abstract

Co-expression networks are used to find prognostic lncRNA programmes, but lncRNA quantification depends on unprocessed RNA in the library. In renal cancer (1,062 patients), the leading lncRNA axis tracked the non-exonic (non-feature) read fraction, a routine alignment metric, at Spearman rho 0.90. Without adjustment, most of the lncRNA network merged into one module. Adjustment preserved agreement with single-cell tumour-normal differences. Under rules fixed with the renal results known, 9 of 25 TCGA cancer types had a lncRNA-specific axis and 4 had network collapse (8 and 2 of 23 non-renal types). Intronic lncRNAs tracked the metric more than intergenic ones in all 25, and the metric was not consistently prognostic. All five renal modules that survived adjustment kept their direction in an external cohort, two nominally significant and none at a 5% false discovery rate. Three localised to malignant or epithelial cells. Together they added no discrimination over TNM and grade.

## Introduction

Prognostication in renal cell carcinoma rests on pathological stage, nuclear grade and composite scores built from them. The Mayo Clinic SSIGN score combines stage, size, grade and coagulative necrosis to predict cancer-specific survival after radical nephrectomy^1^. The Leibovich score predicts metastatic progression from similar variables^2^, and the UCLA Integrated Staging System combines stage, grade and performance status^3^. These routinely recorded variables are the comparator a molecular marker has to beat.

Many transcriptomic signatures, increasingly built from long non-coding RNAs (lncRNAs), have been reported to improve on these scores. Six lncRNA signatures illustrate the pattern, all derived from the same TCGA-KIRC expression matrix under different biological labels: nine lncRNAs with a survival nomogram^4^, eight autophagy-related^5^, five m6A-related^6^ and four cuproptosis-related lncRNAs^7^, a pyroptosis-related model^8^ and six disulfidptosis-related lncRNAs^9^. They share the discovery cohort and modelling strategy, and none states RNA integrity, mapping rate or per-sample sequencing quality control in its Methods.

Such signatures have been questioned on statistical grounds. When 28 published clear-cell renal cell carcinoma biomarkers were evaluated systematically in 350 TCGA patients, only tumour stage and the ccB expression signature remained independent predictors after multivariable adjustment^10^. In breast cancer, most randomly drawn gene signatures are associated with outcome once they correlate with a dominant biological axis such as proliferation^11^.

A separate problem is technical. The fraction of reads outside annotated exons varies systematically between RNA-seq libraries, largely with extraction method and library selection^12^, which is why the exonic rate is a routine per-sample quality metric^13^. RNA degradation widely distorts expression estimates, and covariate adjustment removes much of this distortion where normalisation does not^14,15^. In TCGA RNA-seq, library size, tumour purity and processing plate affect gene co-expression and expression-survival associations^16^. Technical confounders inflate co-expression network edges, and removing leading principal components before network inference reduces false connections^17^. Such correction can also remove biology, a risk not yet tested against single-cell or cell-type ground truth. How far a single routine alignment metric carries the co-expression structure and apparent prognostic signal of a whole gene class has not been quantified.

Non-exonic reads should matter most for lncRNAs, which are low in abundance, often retained in the nucleus and often inside or opposite protein-coding loci, where unspliced and intronic reads accumulate^18^. Among the lncRNAs analysed, 58.3% overlap or sit opposite a protein-coding gene (Supplementary Table 1d), and intron retention is widespread in cancer transcriptomes^19^. Apparent lncRNA expression should therefore depend on a library’s unprocessed RNA content, and a co-expression network built on it should inherit that dependence before any survival model is fitted.

To measure this dependence and its consequences, we assembled four renal cancer cohorts, processed through one Genomic Data Commons STAR-Counts pipeline, and four single-cell cohorts without outcome data. One renal cohort used a different library protocol. We related the leading lncRNA axis to the alignment metrics and network structure of the same libraries, then residualised expression on those metrics before defining modules. We tested modules and published signatures for prognostic value with and without adjustment, and asked whether surviving modules hold in an external cohort, transfer across protocols, localise to cells and add to TNM with grade. Finally, we tested the correction against single-cell ground truth and measured the axis and network reorganisation across 25 TCGA cancer types under decision rules fixed before any non-renal cancer type was analysed.

## Results

### Four cohorts differ in library protocol and in non-exonic read content

To relate lncRNA structure to library content, we first defined four renal cohorts and compared their library protocols and alignment metrics. From 614 TCGA-KIRC STAR-Counts files we excluded 73 non-primary-tumour samples, 5 failed libraries, 4 duplicate aliquots and 4 with non-positive or missing survival time, leaving 528 patients with 173 deaths and a reverse Kaplan-Meier median follow-up of 1,657 days (Fig. 1a, Table 1). Nodal status was explicitly assessed in 259 of 528 patients (17 node-positive) and M category in 494 (79 M1). Two stage IV patients without T4 disease or an explicit M category were reconciled to M1, giving 81 M1 patients of 528. Models requiring the full clinical covariate set used 520 patients with 172 deaths. The CPTAC-3 validation cohort kept 200 patients with 32 deaths from 353 primary-tumour files after removal of duplicate aliquots, non-clear-cell cases and missing survival times. The subtype cohorts were 269 TCGA-KIRP patients and 65 TCGA-KICH patients with 9 deaths, and the 263 TCGA-KIRP patients with a recorded T category had 40 deaths (Fig. 1b, Table 1, Supplementary Data 1).

**Fig. 1.**
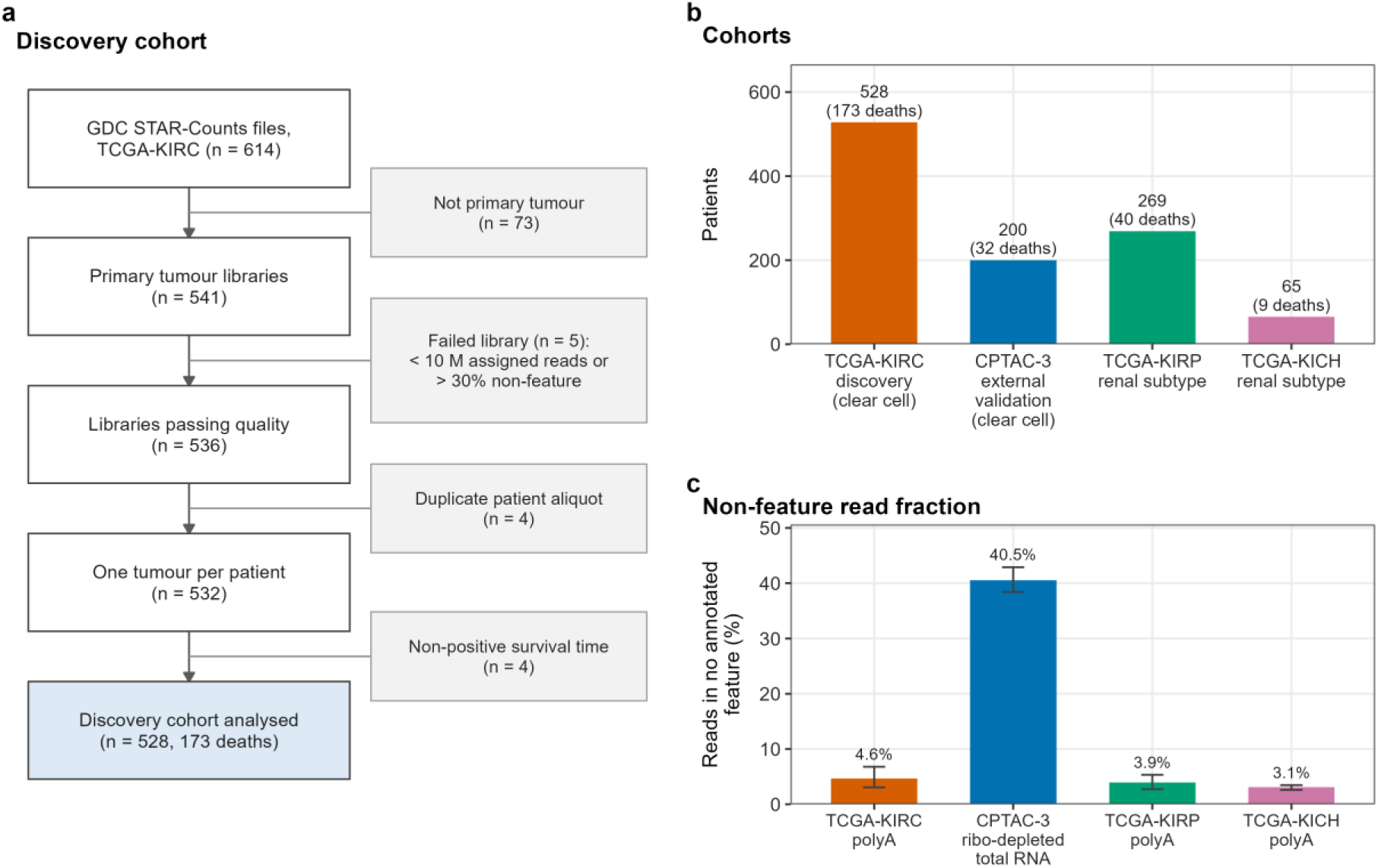
Study design, cohorts and library protocol. **a** Assembly of the discovery cohort. Of 614 TCGA-KIRC STAR-Counts files downloaded from the Genomic Data Commons, 73 were not from primary tumours, leaving 541 primary-tumour libraries. Five failed the library rule (fewer than 10 million assigned reads or more than 30% non-feature reads), leaving 536. Four were duplicate aliquots, leaving 532 tumours from distinct patients, and 4 patients had non-positive or missing survival time. The shaded box is the analysed discovery cohort of 528 patients with 173 deaths. **b** Patients per cohort, with deaths in parentheses: TCGA-KIRC 528 (173), CPTAC-3 200 (32), TCGA-KIRP 269 (40) and TCGA-KICH 65 (9), totalling 1,062 patients and 254 deaths. The TCGA-KIRP bar counts the 40 deaths among the 263 patients with a recorded T category, who enter the adjusted models. Of all 269 TCGA-KIRP patients with module scores, 42 died. Colours identify cohorts, as in **c**. **c** Median non-feature read fraction per cohort, the percentage of reads in the STAR gene-counts summary assigned to no annotated feature. The denominator sums the unmapped, multimapping, non-feature, ambiguous and gene-assigned counts. Bars are medians and whiskers interquartile ranges: TCGA-KIRC 4.6% (3.1 to 6.8), CPTAC-3 40.5% (38.4 to 42.9), TCGA-KIRP 3.9% (2.7 to 5.3) and TCGA-KICH 3.1% (2.6 to 3.5), on 528, 200, 269 and 65 libraries. Library protocols are given under the cohort labels. The ribo-depleted total-RNA cohort is about 9 to 13 times higher than the poly(A) cohorts (8.7-fold against TCGA-KIRC and 13.0-fold against TCGA-KICH). Each cohort used one protocol, so protocol and cohort cannot be separated.

**Table 1.** Characteristics of the four renal cancer cohorts. All four cohorts were quantified with the same Genomic Data Commons STAR-Counts pipeline. TCGA libraries are poly(A)-selected and CPTAC-3 libraries are ribo-depleted total RNA. Median follow-up is the reverse Kaplan-Meier median. Percentages are of patients with the variable recorded. T category is recorded for 528 of 528 TCGA-KIRC, 199 of 200 CPTAC-3, 263 of 269 TCGA-KIRP and 65 of 65 TCGA-KICH patients. Grade is recorded for 520 of 528 TCGA-KIRC and 200 of 200 CPTAC-3 patients. The M1 column gives the percentage of all patients who are M1. Patients recorded MX are treated as M0, and patients with AJCC stage IV disease and a T category below T4 are recoded as M1 (2 TCGA-KIRC, 12 CPTAC-3, 5 TCGA-KIRP and 0 TCGA-KICH patients). The figure in parentheses is the percentage M1 among patients with an explicitly assessed M category (494 of 528, 70 of 200, 99 of 269 and 36 of 65). Nodal status was explicitly assessed in 259 of 528 TCGA-KIRC patients, 17 (6.6%) of them node-positive, and in 56 of 200 CPTAC-3 patients, 8 (14.3%) of them node-positive. It is not tabulated because the number of patients explicitly assessed is not recorded for TCGA-KIRP or TCGA-KICH. Grade is missing for every TCGA-KIRP and TCGA-KICH patient at this data release and is not a standard measure for chromophobe tumours, so it did not enter the subtype covariate set. For TCGA-KIRP and TCGA-KICH, deaths are counted among the patients entering the adjusted models, which are the 263 of 269 TCGA-KIRP patients with a recorded T category and all 65 TCGA-KICH patients. Of all 269 TCGA-KIRP patients with module scores, 42 died. The non-feature read fraction is defined in the Methods and the Fig. 1c legend. IQR, interquartile range.

| Cohort | Patients | Deaths | Median follow-up (days) | Median age (years) | Male (%) | T3 or T4 (%) | M1 (%) | Grade 3 or 4 (%) | Non-feature reads (%), median (IQR) |
| --- | --- | --- | --- | --- | --- | --- | --- | --- | --- |
| TCGA-KIRC (discovery, clear cell) | 528 | 173 | 1,657 | 61.1 | 65.0 | 36.0 | 15.3 (16.0) | 53.8 | 4.6 (3.1 to 6.8) |
| CPTAC-3 (external validation, clear cell) | 200 | 32 | 1,414 | 61.5 | 71.5 | 36.2 | 8.0 (5.7) | 40.0 | 40.5 (38.4 to 42.9) |
| TCGA-KIRP (papillary) | 269 | 40 | 805 | 62.2 | 72.9 | 23.2 | 5.2 (9.1) | not recorded | 3.9 (2.7 to 5.3) |
| TCGA-KICH (chromophobe) | 65 | 9 | 2,752 | 50.7 | 58.5 | 30.8 | 3.1 (5.6) | not recorded | 3.1 (2.6 to 3.5) |

The TCGA libraries are poly(A)-selected and the CPTAC-3 libraries ribo-depleted total RNA. The non-feature read fraction is the percentage of reads that STAR assigns to no annotated feature in its gene-counts summary (Methods). It measures a library’s non-exonic, largely pre-mRNA read content and, with the multimapping fraction and depth, is one of three alignment metrics used throughout. Its median was 4.6% in TCGA-KIRC (interquartile range 3.1 to 6.8), 40.5% in CPTAC-3 (38.4 to 42.9), 3.9% in TCGA-KIRP (2.7 to 5.3) and 3.1% in TCGA-KICH (2.6 to 3.5) (Fig. 1c), an 8.7-fold difference between the validation and discovery cohorts. The metric therefore varies within every cohort and differs 8.7- to 13-fold between protocols, so every adjustment for it is fitted within a cohort and never carried from one cohort to another.

### The leading lncRNA expression axis tracks non-exonic read content

We next asked whether the main source of lncRNA variation follows the non-feature fraction, as dependence on unprocessed RNA content would predict. The leading principal component of the observed lncRNA matrix explained 20.0% of total variance, 2.5 times the second (8.1%). It correlated with the non-feature read fraction at Spearman rho 0.90 (p 1.4e-186, n = 511, Fig. 2a) and only weakly with depth (-0.13) and the ESTIMATE stromal and immune scores (-0.16 and 0.04). The leading protein-coding component (13.7% of variance) does not track the metric (rho 0.05, p 0.30, n = 506, Fig. 2b). It tracks the multimapping fraction instead (-0.51), and its fourth and fifth components track the metric at 0.61 and 0.66.

**Fig. 2.**
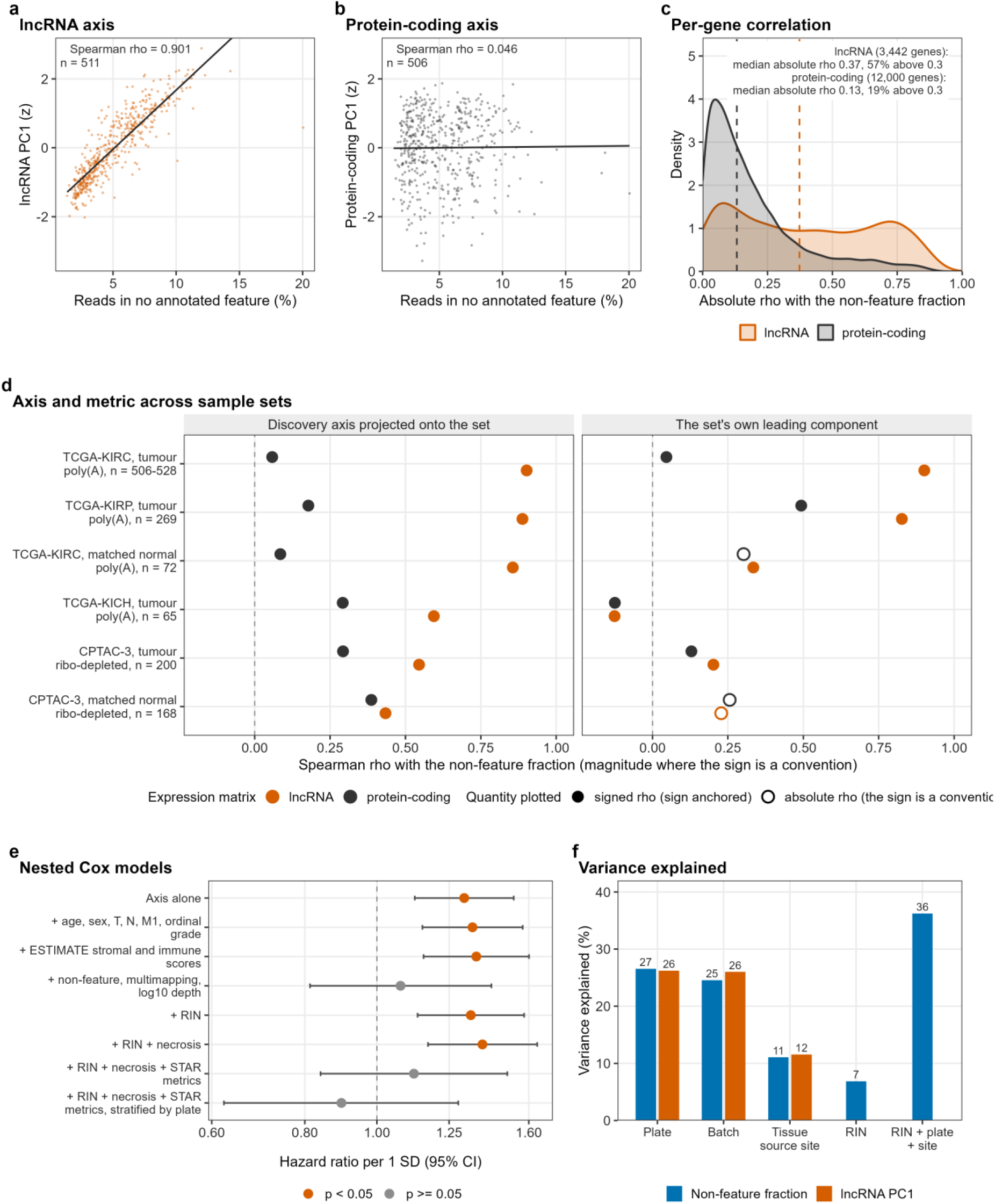
The leading lncRNA expression axis tracks non-exonic read content. **a** First principal component of the observed lncRNA matrix, log2(FPKM + 1), against the non-feature read fraction per library, with a least-squares fit (Spearman rho 0.90, n = 511). **b** As **a** for the protein-coding matrix, on identical axes (rho 0.05, n = 506). The observed protein-coding red eigengene (473 genes, RNA splicing) does track the metric (rho 0.84, Supplementary Fig. 5a). **c** Densities of per-gene absolute Spearman correlation with the metric. Dashed medians are 0.37 for 3,442 lncRNAs (56.8% above 0.3) and 0.13 for 12,000 protein-coding genes (18.8%). **d** Spearman correlation of the axis with the metric in six sample sets, faceted by the projected discovery axis and each set’s own leading component. Open symbols, plotted as magnitudes, mark the 3 of 12 own-component rows whose sign cannot be fixed because its correlations with mean expression and the projected axis are weak or disagree. The projected lncRNA axis gives rho 0.90, 0.89, 0.86, 0.59, 0.55 and 0.43 in TCGA-KIRC tumours (n = 528), TCGA-KIRP (269), TCGA-KIRC normal kidney (72), TCGA-KICH (65), CPTAC-3 tumours (200) and CPTAC-3 normal tissue (168). Within each set the projected lncRNA correlation exceeds the projected protein-coding correlation by 0.30 to 0.84 in the four poly(A) sets and by 0.05 to 0.25 in the two ribo-depleted sets. **e** Hazard ratio per standard deviation of the lncRNA axis under eight nested Cox models (503 patients, 163 events), with 95% confidence intervals on a logarithmic axis. Red marks p below 0.05. The hazard ratio is 1.31 to 1.39 before the three STAR metrics enter and 1.08 after. It is 1.12 with RNA integrity and slide necrosis added to the STAR metrics, and 0.90 with plate stratification. **f** Variance in the metric and the axis explained by plate (26.6% and 26.2%), batch (24.6% and 26.0%) and tissue source site (11.1% and 11.6%), as one-way analysis of variance R-squared on 511 to 528 libraries. Two further bars, for the metric only, give regression R-squared on the 527 patients with an integrity number, for RNA integrity alone (6.9%, 1 degree of freedom) and for RNA integrity with plate and site (36.2%, 25 degrees of freedom). The joint bar comes from a single model, so it need not equal the sum of the other bars.

At gene level the effect is concentrated in lncRNAs. The median absolute correlation with the metric was 0.37 across the 3,442 lncRNAs, with 56.8% of genes above 0.3, against 0.13 and 18.8% across the 12,000 protein-coding genes (Fig. 2c). The exception is the red module (473 genes) of the protein-coding network described below. Its observed eigengene correlates with the metric at rho 0.84, with a median per-gene correlation of 0.68 and 94.5% of genes above 0.3, against 0.22 in the next strongest module. It is enriched for the Gene Ontology terms RNA splicing (adjusted p 4.5e-15) and mRNA processing (8.9e-14), as a shift towards unspliced and intronic read content would predict.

Normalisation could create such an axis, and Genomic Data Commons FPKM is one route into it, because it divides by reads assigned to protein-coding genes while intronic and overlapping reads accumulate at lncRNA loci. The lncRNA read share tracks the metric at rho 0.87 (Supplementary Table 2b). Yet the axis persists under counts per million, trimmed mean of M-values, a variance-stabilising transform and quantile normalisation of the same 511 libraries, with the leading component tracking the metric at absolute rho 0.93, 0.91, 0.92 and 0.88 (Supplementary Fig. 1). On the trimmed-mean scale, the hazard ratios of the 16 lncRNA modules defined below shift downward relative to FPKM for 7 without adjustment and for none after residualisation (Supplementary Table 2).

The signal is carried by genes whose exons lie inside or opposite the transcribed span of a protein-coding gene. With lncRNAs classified by position against GENCODE v36, intronic and exonic-sense lncRNAs had median absolute correlations with the metric of 0.53 and 0.52. The intergenic and divergent classes, which overlap no protein-coding gene, both had 0.25. The class difference (Kruskal-Wallis chi-square 263.31 on 5 degrees of freedom, p 7.7e-55) survives adjustment for mean expression and log10 gene length (F 54.80, p 6.2e-55), although other gene properties could still contribute (Supplementary Table 1).

To test whether the axis requires tumour tissue, we projected the fixed discovery direction onto the 72 matched TCGA-KIRC normal-kidney libraries (72 patients, 71 in the discovery set). It correlates with the metric at rho 0.86 (p 8.9e-22), and the projected protein-coding direction does not (0.09, Supplementary Fig. 2). The unpaired tumour-normal difference in the metric (1.73 percentage points) largely reflects plate confounding, falling to 0.15 points on shared plates and within patient (Supplementary Table 3).

Across cohorts, the projected discovery direction correlates with the non-feature fraction at rho 0.90 in TCGA-KIRC tumours, 0.89 in TCGA-KIRP, 0.86 in TCGA-KIRC normal kidney and 0.59 in TCGA-KICH, and at 0.55 and 0.43 in CPTAC-3 tumours and normal kidney (Fig. 2d, Supplementary Table 4). The correlation is graded by library protocol, and the axis transfers to ribo-depleted libraries at reduced strength. Its lncRNA specificity is a property of poly(A)-selected libraries. Against the projected protein-coding direction the contrast is 0.90 to 0.06 in TCGA-KIRC tumours but 0.55 to 0.29 in CPTAC-3, because retaining unprocessed RNA raises intronic content for every gene.

### The axis reorganises the lncRNA co-expression network

An axis this strong could set the structure of any network built on these matrices, so we built the discovery networks on expression residualised on the three alignment metrics and compared the lncRNA network with one built without adjustment. From the 12,000 protein-coding and 3,442 lncRNA genes that passed filtering, the adjusted protein-coding network (506 samples, soft power 18) resolved 13 modules, with 4,069 genes (33.9%) unassigned. The adjusted lncRNA network (511 samples, soft power 10, fixed by an outcome-blind sweep) resolved 16 modules, with 966 genes (28.1%) unassigned (Supplementary Fig. 3). Each of the 29 module eigengenes is a rotation fitted once on the residualised discovery matrix and reproduces the WGCNA eigengene at r = 1.000 (Supplementary Data 2).

Rebuilt on the same 511 libraries with identical parameters but without adjustment, the lncRNA network resolves into 3 modules instead of 16. Its largest module holds 1,982 of 3,442 genes (57.6%), against 438 (12.7%) in the adjusted network, and its eigengene correlates at r 0.98 with the observed leading component and 0.91 with the metric. Mean intramodular connectivity over assigned genes, each computed on its own matrix, is 70.5 against 3.3 after adjustment. The two partitions share almost no structure (adjusted Rand index 0.062 over all genes, 0.047 over the 1,722 genes assigned in both, Supplementary Table 5).

The largest unadjusted module, hereafter the dominant module, absorbs the positional classes that carry the metric’s signal. It holds 71.5% of intronic and 69.8% of exonic-sense lncRNAs, against 43.7% of intergenic lncRNAs and 57.6% overall (false discovery rate 8.7e-17 for intronic, 1.8e-29 for intergenic). The adjusted lncRNA black, green and turquoise modules lie almost wholly inside it (99.5% to 100% of their genes), whereas the unadjusted network left 86.1% of greenyellow and 63.4% of blue unassigned (Supplementary Table 5). Adjustment therefore splits the dominant module and resolves modules that the unadjusted network could not, so any downstream analysis depends on whether expression was adjusted.

### Across TCGA the mechanism is general and varies in strength

To test whether the renal finding is specific to one dataset, we repeated the measurements in every TCGA project under two decision rules. We fixed them before analysing any non-renal project but after seeing the renal results (Methods), so the counts below are also given for the 23 non-renal types alone. Of 11,505 STAR-Counts files from 33 projects, 10,722 libraries were retained, and 25 projects had at least 100 retained primary tumours (Supplementary Table 6). Through the pan-cancer pipeline, TCGA-KIRC reproduced the discovery axis (rho 0.91) and the collapsed network (3 modules, 57.5% of genes in the largest, adjusted Rand index 0.06).

The leading lncRNA component tracked the non-feature fraction at absolute rho of at least 0.5 in 19 of the 25 cancer types (median 0.72, against 0.28 for the protein-coding component, Fig. 3a), 17 of them non-renal. The locked rule for a lncRNA-specific axis (Rule A), which adds a margin of 0.3 over the protein-coding component, was met in 9: KIRC, LUAD, LUSC, OV, STAD, READ, PAAD, COAD and LGG, 8 of them among the 23 non-renal types. Of the other ten, five (GBM, SKCM, KIRP, SARC and BLCA) had a protein-coding component that also reached absolute rho of at least 0.5, and the rest missed the margin, THCA and ESCA narrowly. The following comparisons are descriptive and outside the locked rules. At gene level, lncRNAs tracked the metric more than protein-coding genes in 22 of 25 types, and intronic more than intergenic lncRNAs in all 25 (median 0.35 against 0.19, six-class Kruskal-Wallis p at most 1.3e-15 in every type, Fig. 3b). In 13 normal-tissue sets with at least 20 libraries, the lncRNA component reached absolute rho of at least 0.5 in 8 and met the full rule in 2. In TCGA-KIRC normal kidney the projected discovery direction tracks the metric at 0.86 but the samples’ own leading component only at 0.33, so the axis is present without dominating the variance. Network collapse is the extreme form of the effect. The locked rule for a reorganised network (Rule C) requires a largest observed module holding at least 40% of genes, with an eigengene tracking the metric at absolute r of at least 0.7. It also requires an adjusted Rand index below 0.2 between observed and adjusted partitions. The rule was met in 4 of 25 types (GBM, KIRC, SKCM and KIRP), 2 of them among the 23 non-renal types, with LUAD just outside (44%, 0.21) (Fig. 3c). Short of collapse, the largest observed module tracked the metric at absolute r of at least 0.7 in 17 of 25 types, and the partitions agreed only moderately (median adjusted Rand index 0.31).

**Fig. 3.**
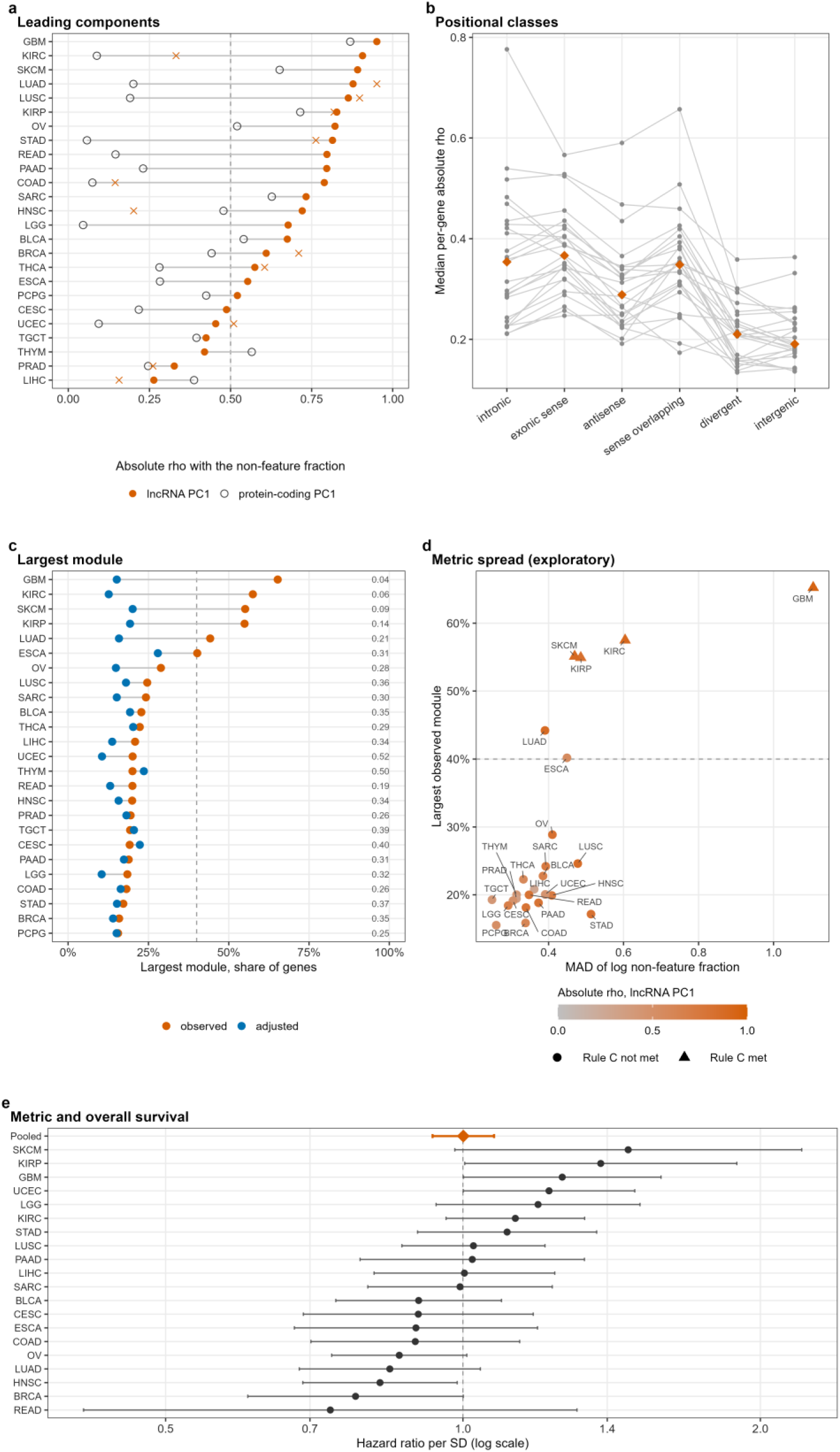
The lncRNA axis and network reorganisation across TCGA. Twenty-five cancer types with at least 100 retained primary tumours. Dashed lines are the decision rules, fixed before any non-renal project was analysed (Methods). **a** Absolute Spearman correlation with the non-feature read fraction of the leading principal component of the lncRNA (filled) and protein-coding (open) matrices, and of the lncRNA component in solid-tissue normals with at least 20 libraries (crosses). Rule A, for a lncRNA-specific axis (absolute rho at least 0.5 and at least 0.3 above protein-coding), is met in 9 of 25, 8 of them non-renal. **b** Median per-gene absolute correlation with the metric by positional lncRNA class, one line per cancer type, with diamonds for the median over types. Intronic exceed intergenic lncRNAs in all 25. **c** Share of lncRNAs in the largest module of the observed (orange) and adjusted (blue) networks, with the adjusted Rand index between the two partitions at right. Rule C, for a reorganised network, is met in GBM, KIRC, SKCM and KIRP. **d** Exploratory panel, specified after the locked rules were applied. Within-cohort spread of the metric (median absolute deviation of its logarithm) against the share of genes in the largest observed module, coloured by the absolute rho of the lncRNA component. Triangles meet Rule C. Spread correlates with the largest-module share at Spearman rho 0.69 (false discovery rate 0.0016) and with the absolute rho of the lncRNA component at 0.73 (6.1e-04). **e** Hazard ratio for overall survival per standard deviation of the non-feature fraction in the 20 cancer types with at least 20 deaths, adjusted for age, sex and stage where recorded and stratified by sequencing plate. Error bars are 95% confidence intervals, and the diamond is the random-effects pooled estimate, 1.00 (0.93 to 1.08, I-squared 50%). Cancer types are TCGA project codes. Source data are in Supplementary Table 6.

To see why the effect varies in strength, we asked whether it follows the within-cohort variability of the metric, in an exploratory analysis specified after the locked rules were applied. Spread was measured as the median absolute deviation of the log non-feature fraction. Across the 25 cancer types, spread correlated with lncRNA axis strength (Spearman rho 0.73, false discovery rate 6.1e-04) and with the share of genes in the largest observed module (0.69, 0.0016) (Fig. 3d, Supplementary Table 6d). The first association held for all four spread measures (0.63 to 0.73), and the second weakened with the standard deviation (0.38, 0.081). No spread measure predicted the adjusted Rand index, and GBM, with by far the most variable libraries, contributes much of the pattern.

### The metric is prognostic in renal cancer but not consistently across cancer types

Studies that treat lncRNA scores as biology would read the axis as a prognostic marker, so we tested it, and the metric itself, in survival models. Hazard ratios are per standard deviation of the exposure or eigengene throughout. In nested Cox models (503 patients, 163 deaths), the axis gave 1.31 alone (95% confidence interval 1.12 to 1.53) and 1.34 to 1.39 after adding clinical terms, ESTIMATE scores, RNA integrity and slide necrosis, so it appears prognostic independently of the recorded clinical variables. The three STAR metrics reduced it to 1.08 (0.81 to 1.43, p 0.61) with a variance inflation factor of 4.5, so collinearity contributes to the collapse. Stratifying the fullest model by sequencing plate gave 0.90 (0.62 to 1.29) (Fig. 2e, Supplementary Table 7a).

Tested directly, the non-feature fraction is itself prognostic. In 520 patients (172 deaths) it gave 1.28 alone (1.13 to 1.46, p 1.2e-04) and 1.36 after clinical adjustment (1.20 to 1.55, p 3.0e-06), or 1.44 on the log10 scale (1.23 to 1.68) (Supplementary Table 8c). Adding ESTIMATE scores and stratifying by tissue source site gave 1.45 (1.25 to 1.69, p 1.6e-06). In the 503 patients with RNA integrity recorded, it remained prognostic within sequencing plate at 1.23 (1.02 to 1.50, p 0.035) after adjustment for RNA integrity, slide necrosis, the multimapping fraction and depth, which removes the axis (Supplementary Table 7b). In CPTAC-3 it was not prognostic, with a reversed point estimate of 0.76 (0.45 to 1.27, p 0.29, 199 patients, 32 events). Two explanations fit, and these data cannot separate them: 32 events give little power, and under that cohort’s protocol the metric has a different distribution and possibly a different meaning.

If the renal association reflected a general effect of library content on outcome, it should recur in other cancer types, so we tested the 20 eligible types with at least 20 deaths. Plate-stratified Cox models adjusted for age, sex and, where recorded, stage gave a pooled hazard ratio per standard deviation of the metric of 1.00 (0.93 to 1.08, I-squared 50%, heterogeneity p 0.0047, prediction interval 0.79 to 1.26). Estimates exceeded one in 10 types, and 4 were nominally significant, three above one (GBM, KIRP and UCEC) and one below (HNSC) (Fig. 3e, Supplementary Table 6). Without plate stratification the pooled estimate was 1.01 (0.94 to 1.08). For TCGA-KIRC, the pan-cancer pipeline (Clinical Data Resource endpoint, stage in place of T, N, M and grade) gives 1.27 (1.12 to 1.43) without and 1.13 (0.96 to 1.33) with plate stratification. In renal cancer, the axis therefore loses its association with survival once the alignment metrics or sequencing plate enter the model, whereas the metric itself remains associated with survival in the discovery models (1.23 within plate). Across cancer types the metric is not a consistent marker of outcome.

### Processing batch and RNA integrity explain a third of the metric’s variance

To see what the metric reflects, we partitioned its variance. It is structured by batch: sequencing plate explains 26.6% of its variance (p 6.5e-27), Biospecimen Core Resource batch 24.6% and tissue source site 11.1%, and the same three explain 26.2%, 26.0% and 11.6% of variance in the axis (Fig. 2f, Supplementary Table 7d). Plate medians range from 2.6% to 7.1% across the 14 plates with at least ten patients. RNA integrity, recorded for 527 of the 528 patients, correlates with the metric at -0.25 (p 9.4e-09) and explains 6.9% of its variance alone and 36.2% jointly with plate and site (Supplementary Table 7c, e). Slide percent necrosis is uninformative (median zero, correlation -0.05, p 0.28), and no clinical variable is associated with the metric (smallest p 0.21, Supplementary Fig. 4, Supplementary Table 9f).

Biological correlates point the same way. In 501 patients, a block of annotated tumour biology (hypoxia metagene, proliferation score, ESTIMATE scores and slide tumour-nuclei percentage) explains 6.1% of the metric’s variance, against 36.8% for RNA integrity, plate and site. Blocks built from observed expression explain more but cannot be identified as biology, because random gene sets of the same sizes reach 61.0%. Singly, proliferation is uncorrelated with the metric (rho -0.01) and hypoxia correlates in the unexpected direction (-0.14). Stratifying the clinically adjusted model by plate moves the hazard ratio from 1.37 to 1.24 (1.04 to 1.48) for the metric and from 1.36 to 1.13 (0.94 to 1.36) for the axis. No biological covariate moves the metric’s log hazard ratio by more than 5% (Supplementary Fig. 5, Supplementary Table 9).

Driver genotype was not associated with the metric or the axis (5% false discovery rate) in the 368 patients with open-access mutation calls, a subset with a lower metric and half the death rate of the rest. Adjusting for genotype left the metric’s hazard ratio at 1.34 (1.10 to 1.63, Supplementary Table 1). The recorded batch and integrity variables therefore explain about a third of the variance in the metric, annotated tumour biology explains little of it, and most of the variance remains unexplained (Discussion).

### Comparison with single-cell ground truth finds no loss of tumour biology after correction

Residualisation could remove tumour biology along with artefact, so we compared its effect with single-cell data, which do not depend on bulk library preparation. In 72 TCGA-KIRC patients with tumour and matched normal libraries, the bulk tumour-minus-normal difference of 507 lncRNAs agreed with their single-cell fold change between malignant and normal epithelial cells at Spearman rho 0.775, and 92.5% of the 213 single-cell-significant lncRNAs agreed in sign (Supplementary Fig. 6, Supplementary Table 10). After correction, paired agreement fell slightly but detectably to 0.771 (-0.004, 95% interval -0.008 to -0.001). This paired contrast is insensitive, because a patient’s tumour and normal libraries differ little in the correction covariates (0.13 standard deviations in the non-feature fraction). The unpaired contrast of all 532 tumours against all 72 normals (Methods) is more sensitive, because the gap there is 0.86 standard deviations. Agreement was 0.773 before and 0.774 after correction (change 0.001, - 0.013 to 0.014), and sign agreement 93.9% and 91.5% (change -2.4 points, -5.6 to 0.9). The correction therefore neither improved nor detectably degraded agreement with single-cell biology.

Within tumours, residualisation removes by construction the correlation of any lncRNA score with the metric. The informative quantity is therefore how far a score’s coherence with a protein-coding malignant-marker score exceeds that of lncRNA sets that single-cell data classify as non-specific. A score from the 38 single-cell tumour-specific lncRNAs, fixed before any bulk step, correlated with the marker score at 0.55 (0.48 to 0.62) before and 0.48 (0.41 to 0.55) after correction. Abundance-matched non-specific sets gave 0.37 and 0.28, an excess of 0.19 before and 0.20 after. All sets lose some coherence because the marker score itself correlates with the metric at 0.22. Three of the outcomes we recorded in advance were not met (Methods). Paired agreement fell slightly, and the non-specific sets were not neutral. The 15 candidates set aside in bulk as technically sensitive lost their correlation with the metric (0.50 to 0.84 before, -0.08 to 0.11 after) but had no room to show recovery, because four of the five with a normal-epithelium estimate already agreed with single-cell direction before correction. Residualisation can therefore precede network construction without detectable loss of the tumour biology that single cells report, within the lncRNAs that single cells can measure.

### Residualisation and RUV-III both remove the axis, prevent collapse and preserve the survival associations, but disagree on module membership

To test whether these results depend on the correction method, we repeated the analyses on the same 528 tumours with RUV-III using pseudo-replicates of pseudo-samples^16^ (Supplementary Table 11). Pseudo-replicates were built within ccA/ccB and stage groups, across plates and across extremes of the metric and depth. RUV-III removed the link between the leading lncRNA component and the metric (rho 0.05, against 0.06 after residualisation and 0.91 observed) and prevented network collapse (largest module 14% of genes, against 14% and 58%). The five modules that remain prognostic after full adjustment (described below) stayed associated with survival at hazard ratios similar to those after residualisation. The two corrections agreed only moderately on module membership (adjusted Rand index 0.23). RUV-III also removed more single-cell-confirmed tumour signal. The tumour-specific lncRNA score correlated with the malignant-marker score at 0.28 after RUV-III and 0.48 after residualisation, and its excess over abundance-matched non-specific sets drawn for this comparison was 0.11 and 0.22. RUV-III’s controls may capture some tumour-content variation that residualisation keeps by design. Removing the axis, preventing collapse and preserving the survival associations therefore do not depend on the correction method, but module membership does, so individual module identities should be interpreted with caution.

### Adjusting for the alignment metrics changes which modules appear prognostic

A technical axis matters only if it changes what a study concludes, so we next asked whether adjustment, with module membership held fixed, changes which modules appear prognostic. Across the 29 modules, with the clinical model and Benjamini-Hochberg control within each network, 15 were significant with eigengenes from observed expression (11 of them lncRNA modules), ten with expression residualised on the three STAR metrics (Fig. 4a) and 3 with observed eigengenes plus the three metrics as covariates (Supplementary Table 12a). Nine modules were lost on adjustment, four gained, six significant in both arms and ten in neither. The largest change is in the protein-coding red (splicing) module, from 1.45 per standard deviation (1.24 to 1.69, FDR 2.5e-05) to 1.17 (0.99 to 1.40, FDR 0.16). With RNA integrity and slide necrosis added to the clinical model, 12 of 29 are significant without adjustment and 4 with it (Supplementary Table 12c).

**Fig. 4.**
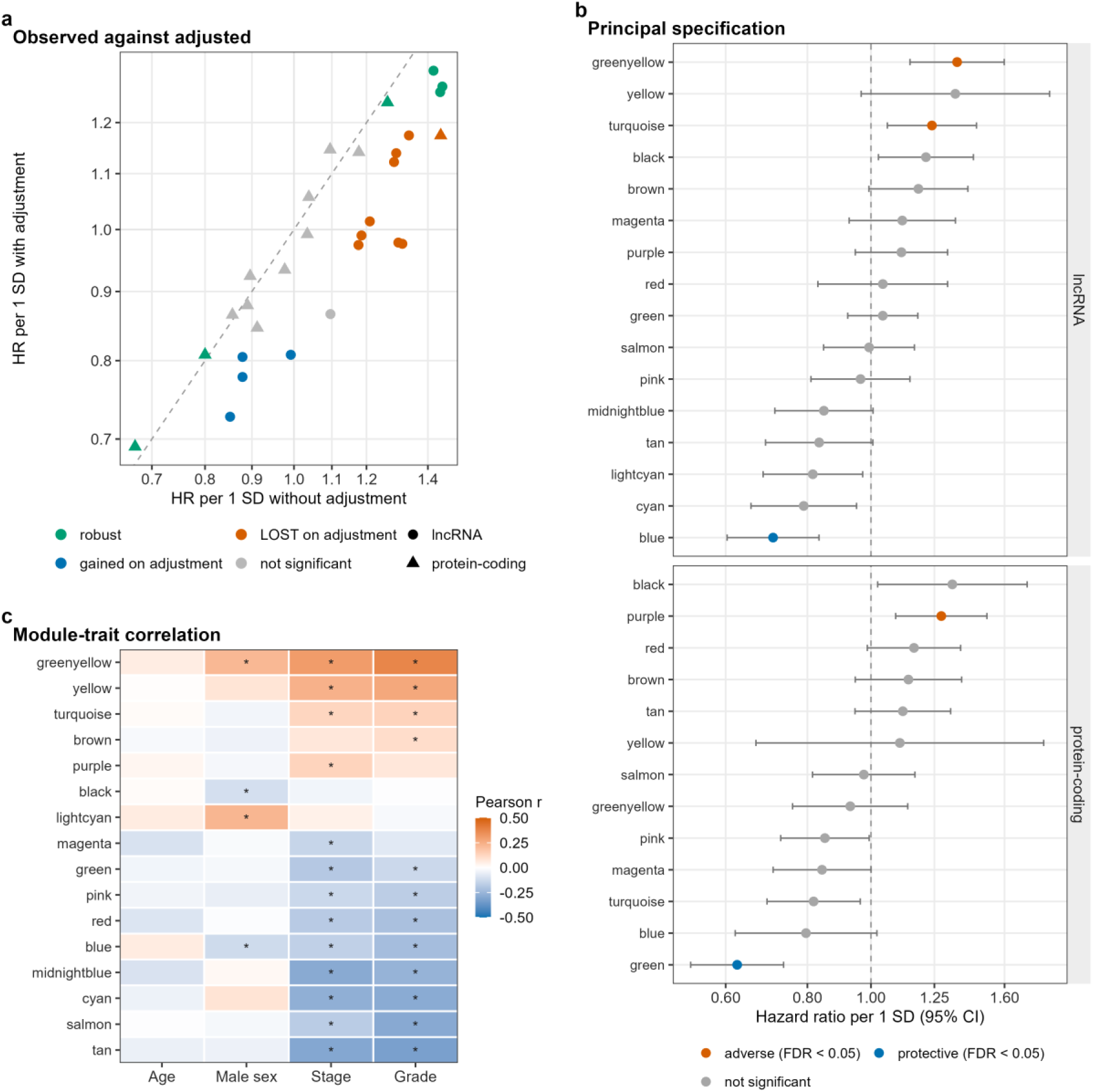
Adjusting for the alignment metrics changes which co-expression modules are prognostic. **a** Hazard ratio per standard deviation of each of the 29 module eigengenes, computed on the observed matrix (horizontal axis) and on the matrix residualised on the three STAR alignment metrics (vertical axis). Module membership is the same on both axes, so only the adjustment differs. Axes are logarithmic and the dashed line marks equality. Colour shows the outcome of adjustment and shape the network. Of 29 modules, 15 were significant at a 5% false discovery rate without adjustment and 10 with it: 9 lost significance, 4 gained it, 6 were significant in both arms and 10 in neither. Models adjust for age, sex, T category, nodal status, M1 and ordinal grade and are fitted on 504 patients with 163 events (lncRNA network) and 502 with 165 (protein-coding network). False discovery rates are controlled within network and arm. **b** Hazard ratio per standard deviation of each module eigengene under the principal specification, by network, with 95% confidence intervals on a logarithmic axis. The principal specification adds the eigengene to age, sex, T category, nodal status, M1, ordinal grade, the ESTIMATE stromal and immune scores and the three STAR metrics, in 504 patients with 163 events (lncRNA network) and 502 with 165 (protein-coding network). Colour marks the direction of modules with a within-network false discovery rate below 0.05, and grey marks non-significant modules. Five of 29 modules meet that threshold: protein-coding green and purple, and lncRNA blue, greenyellow and turquoise. **c** Pearson correlation of the 16 lncRNA module eigengenes with age, male sex, ordinal AJCC stage and ordinal grade in 511 patients. Asterisks mark a Benjamini-Hochberg false discovery rate below 0.05 across the panel’s 64 tests (29 of 64).

Applied to published work, the same test gives a more qualified answer. Six published TCGA-KIRC lncRNA prognostic signatures were re-scored on the 528 discovery tumours, with 34 of their 38 genes present and scoreable in the current annotation. Every signature correlates with the metric, five positively and the Yu signature negatively. Spearman correlations were 0.17 and 0.43 for the two with published coefficients, Hong 2022^7^ and Jiang 2019^4^, and 0.11 to 0.43 for the three whose weights were signed by the univariate Cox coefficient in the same patients, Liu 2023^8^, Gui 2021^5^ and Feng 2024^9^. Yu 2021^6^ mapped only 2 of its 5 lncRNAs and lost the gene with the largest published coefficient, so its truncated score runs opposite to the published model (-0.24) and is reported as a footnote. After residualisation every correlation lies within 0.04 of zero (all p above 0.41).

No signature is abolished by adjustment for the alignment metrics. With clinical adjustment (520 patients, 172 events), the two with published coefficients gave 1.67 (1.45 to 1.93) for Jiang and 1.75 (1.51 to 2.03) for Hong, and the three weighted in sample 1.50 to 1.63 (Supplementary Table 13). Adding the three STAR metrics changed the log hazard ratio by -2.6% and -4.9% for the first two and by -19.0% to -11.1% for the other three, and every adjusted estimate kept p below 0.001. Nor are the signatures less correlated with the metric than chance: against a sign-randomised null, three of the six sit at or above the null median (Supplementary Fig. 7). All six were derived in TCGA-KIRC, so their increments over a clinical model in this cohort (0.021 and 0.024 for the two with published coefficients) are apparent figures.

### Five modules remain associated with survival after full adjustment

To find modules that stay prognostic once the alignment metrics and recorded confounders are accounted for, we used a principal specification that adds the module eigengene to age, sex, T category, nodal status, M1, ordinal grade, the ESTIMATE stromal and immune scores and the three STAR metrics. Under it, five modules remained significant at a within-network false discovery rate below 0.05, and exactly these five survive Benjamini-Hochberg control across all 29 modules of both networks (Table 2, Fig. 4b, Supplementary Data 3). Two are protein-coding (502 patients, 165 events): green (670 genes) at 0.62 (0.53 to 0.74, FDR 2.2e-07) and purple (216 genes) at 1.28 (1.09 to 1.50, FDR 0.016). Three are lncRNA modules (504 patients, 163 events): blue (246 genes) at 0.71 (0.60 to 0.83, FDR 4.8e-04), greenyellow (108 genes) at 1.35 (1.15 to 1.60, FDR 0.0027) and turquoise (438 genes) at 1.24 (1.06 to 1.45, FDR 0.039). Two of the three lncRNA modules, greenyellow and blue, were largely unassigned in the unadjusted network, and the third, turquoise, lay almost wholly inside its dominant module. The lncRNA eigengenes also track stage and grade (Fig. 4c), greenyellow at r 0.31 with ordinal AJCC stage and 0.38 with ordinal grade, and blue at -0.16 and -0.22.

**Table 2.** Module eigengenes associated with overall survival in TCGA-KIRC and their external estimates in CPTAC-3. Hazard ratios (HR) are per 1 standard deviation of the module eigengene, computed on the expression matrix residualised on the alignment metrics. The principal specification is the module eigengene with age, sex, T category, nodal status, M1, ordinal grade, the ESTIMATE stromal and immune scores, and the three STAR alignment metrics (non-feature read fraction, multimapping fraction and log10 assigned reads). The clinical model is age, sex, T category, nodal status, M1 and ordinal grade. False discovery rates (FDR) are Benjamini-Hochberg, controlled within network across the 13 protein-coding or 16 lncRNA modules and, in the column so labelled, jointly across all 29 modules. The five modules listed meet a 5% within-network false discovery rate under the principal specification, and the same five meet the joint threshold. The two discovery sample sizes differ because the networks exclude different connectivity outliers and models are fitted on complete cases. In CPTAC-3, module membership and eigengene loadings were fixed in discovery, only the module coefficient was re-estimated, and eigengenes were standardised within CPTAC-3. Panel (b) therefore reports an association re-estimated in an external cohort, and Table 3 reports the performance of the fixed prediction rule. The 12-parameter model uses the principal covariate set and carries 2.7 events per parameter, below the threshold of 5 applied elsewhere. The 6-parameter model (module eigengene, age, sex, T category, M1 and ordinal grade) carries 5.3 and meets that threshold. Its false discovery rates are controlled across the five modules tested. Replication is defined in the Methods, and no module meets it. Standardising CPTAC-3 with the discovery means and standard deviations in place of within-cohort standardisation changes no hazard ratio in panel (b) by more than 0.01 (Supplementary Table 21g).

| Network | Module | Genes | n | Events | HR, principal specification (95% CI) | FDR, within network | FDR, all 29 modules | HR, clinical model (95% CI) | FDR, clinical model |
| --- | --- | --- | --- | --- | --- | --- | --- | --- | --- |
| lncRNA | blue | 246 | 504 | 163 | 0.71 (0.60 to 0.83) | 4.8e-04 | 4.4e-04 | 0.73 (0.62 to 0.85) | 0.0015 |
| lncRNA | greenyellow | 108 | 504 | 163 | 1.35 (1.15 to 1.60) | 0.0027 | 0.0033 | 1.31 (1.11 to 1.54) | 0.0092 |
| lncRNA | turquoise | 438 | 504 | 163 | 1.24 (1.06 to 1.45) | 0.039 | 0.043 | 1.26 (1.08 to 1.49) | 0.024 |
| protein-coding | green | 670 | 502 | 165 | 0.62 (0.53 to 0.74) | 2.2e-07 | 4.9e-07 | 0.69 (0.59 to 0.81) | 3.8e-05 |
| protein-coding | purple | 216 | 502 | 165 | 1.28 (1.09 to 1.50) | 0.016 | 0.018 | 1.24 (1.06 to 1.45) | 0.036 |

*(b) External estimates, CPTAC-3 (199 patients, 32 events)*

| Network | Module | HR, 12-parameter model (95% CI) | p | FDR | HR, 6-parameter model (95% CI) | p | FDR | Direction as in discovery |
| --- | --- | --- | --- | --- | --- | --- | --- | --- |
| lncRNA | blue | 0.78 (0.55 to 1.11) | 0.17 | 0.21 | 0.79 (0.57 to 1.10) | 0.16 | 0.20 | yes |
| lncRNA | greenyellow | 1.59 (1.12 to 2.25) | 0.0097 | 0.042 | 1.42 (1.07 to 1.90) | 0.016 | 0.068 | yes |
| lncRNA | turquoise | 1.11 (0.81 to 1.53) | 0.51 | 0.51 | 1.10 (0.85 to 1.43) | 0.46 | 0.46 | yes |
| protein-coding | green | 0.73 (0.51 to 1.04) | 0.082 | 0.14 | 0.74 (0.54 to 1.03) | 0.071 | 0.12 | yes |
| protein-coding | purple | 1.52 (1.08 to 2.14) | 0.017 | 0.042 | 1.41 (1.04 to 1.92) | 0.027 | 0.068 | yes |

The five associations do not depend on the coding of unassessed nodal and metastasis status. Alternative codings moved every hazard ratio by at most 0.026, and restriction to patients with both assessed left all five in the same direction (Supplementary Table 14). The eigengene term meets the proportional-hazards assumption in every model (p 0.45 to 0.95 for the five modules), and global-test violations lie in the clinical covariates.

Because module eigengenes differ by sex (8 of 29 at a 5% false discovery rate), we also fitted sex-specific models. Greenyellow is nominally stronger in women in each cohort where a sex-specific fit converges, at 1.97 (1.40 to 2.78) against 1.20 (0.98 to 1.47) in TCGA-KIRC, but no interaction survives multiplicity correction, so these estimates are descriptive (Supplementary Table 15). With membership and loadings fixed in discovery, these five modules are the set carried into the external cohort.

### All five modules keep direction in CPTAC-3 and the networks transfer

Associations selected in one cohort need confirmation in an independent one, so we projected the five modules onto CPTAC-3 with membership and loadings fixed in discovery and re-estimated only the module coefficient. All five kept their discovery direction (Table 2, Fig. 5a, one-sided sign test p 0.031). Evidence for individual modules is weaker. Replication of a module was defined as a false discovery rate below 0.05 under the six-parameter set, the one meeting the events-per-variable threshold of 5 (Methods), and by that rule none of the five replicated. In 199 patients with 32 deaths, lncRNA greenyellow gave 1.42 (1.07 to 1.90, p 0.016) and protein-coding purple 1.41 (1.04 to 1.92, p 0.027) under that set, nominally significant but at a false discovery rate of 0.068. Under the twelve-parameter principal set (2.7 events per variable) they gave 1.59 (1.12 to 2.25) and 1.52 (1.08 to 2.14), both at a false discovery rate of 0.042, but that set falls below the events-per-variable threshold and is not used to claim replication. Green, blue and turquoise kept their direction of association without nominal significance.

**Fig. 5.**
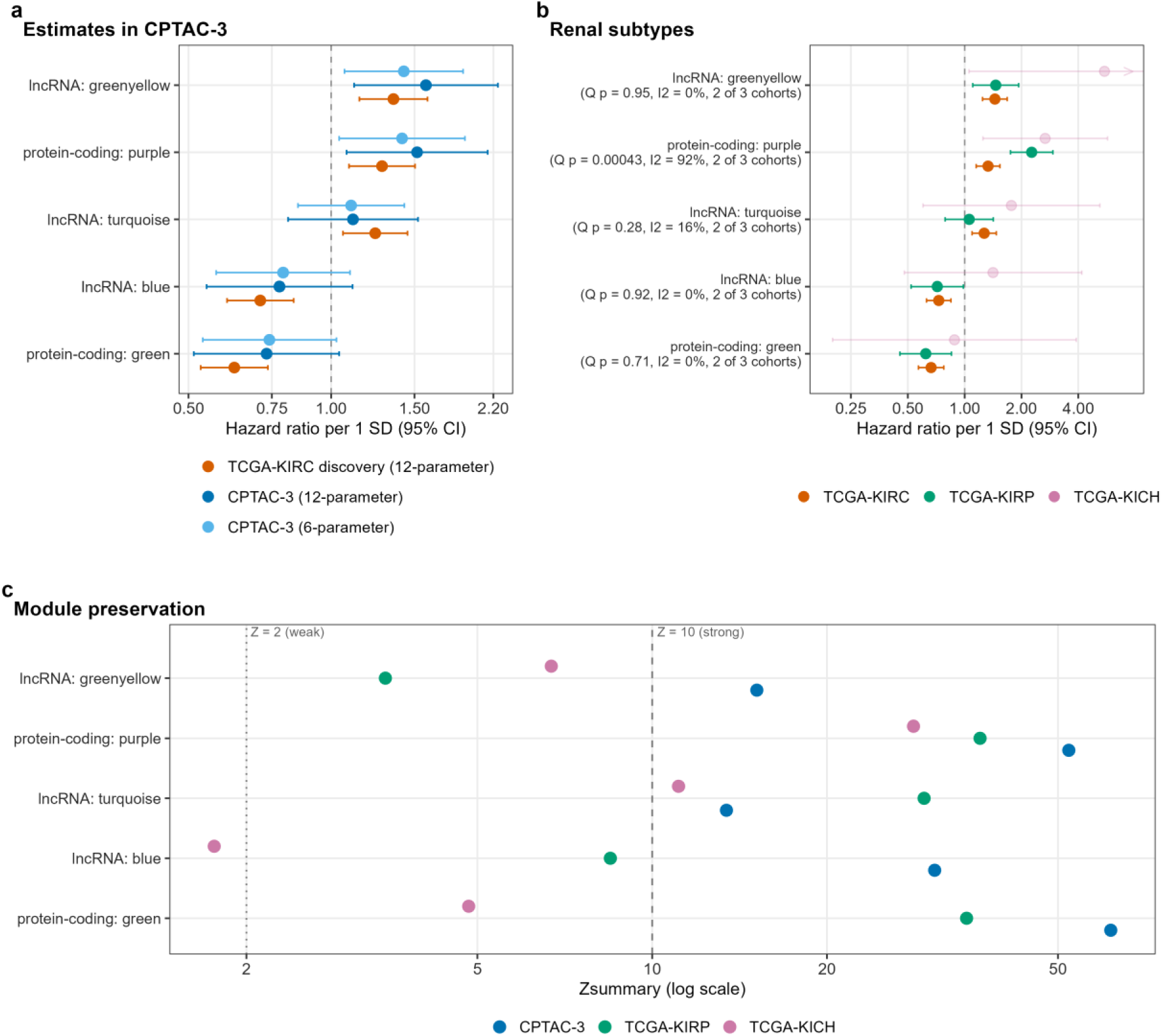
External association estimates, subtype estimates and module preservation. **a** Three estimates for each of the five modules significant in discovery. These are TCGA-KIRC under the principal specification (504 patients and 163 events for lncRNA modules, 502 and 165 for protein-coding modules), CPTAC-3 with the same twelve-parameter covariate set, and CPTAC-3 with a six-parameter set of module eigengene, age, sex, T category, M1 and ordinal grade. Both CPTAC-3 arms use 199 patients with 32 events, with membership and loadings fixed in discovery and only the module coefficient re-estimated. Points are hazard ratios per standard deviation with 95% confidence intervals on a logarithmic axis. All five keep the discovery direction in both arms (one-sided sign test p 0.031), and greenyellow and purple are nominally significant (p below 0.05) in both. Replication was defined as a false discovery rate below 0.05 in the six-parameter arm (5.3 events per parameter), and no module meets it. The twelve-parameter arm, in which greenyellow and purple reach a false discovery rate of 0.042, has 2.7 events per parameter, below the threshold of 5 used elsewhere, and is not used to claim replication. **b** The same five modules projected with the fixed discovery loadings onto TCGA-KIRC (497 patients, 160 events), TCGA-KIRP (263, 40) and TCGA-KICH (65, 9), adjusted for one parsimonious set of age, sex, T category, M1 and the non-feature fraction. Error bars are 95% confidence intervals on a logarithmic axis. Row labels give Cochran Q p values and I-squared for TCGA-KIRC and TCGA-KIRP, the two cohorts meeting the events-per-parameter rule. TCGA-KICH, at 1.5 events per parameter, is faded and descriptive only. Its lncRNA greenyellow interval extends beyond the axis limit and is marked. **c** Preservation of the five modules in CPTAC-3 (n = 200), TCGA-KIRP (269) and TCGA-KICH (65), as WGCNA Zsummary on a logarithmic axis, from 100 permutations for the lncRNA network and 30 for the protein-coding network. Reference lines mark the conventional weak (Z = 2) and strong (Z = 10) thresholds, and 10 of the 15 module-cohort pairs lie above 10.

We then projected the modules onto papillary renal cell carcinoma with a six-parameter model omitting grade, and four of the five kept direction and significance (263 patients, 40 events). Protein-coding green gave 0.62 (0.45 to 0.85, p 0.0030), lncRNA blue 0.72 (0.52 to 0.98, p 0.040), lncRNA greenyellow 1.46 (1.10 to 1.93, p 0.0080) and protein-coding purple 2.27 (1.75 to 2.93, p 5.8e-10), and lncRNA turquoise was null at 1.06 (0.79 to 1.42, p 0.71) (Fig. 5b). Only purple differs between TCGA-KIRC and TCGA-KIRP (Cochran Q 12.38, p 4.3e-04, I-squared 92.0%), the other four giving I-squared of 0.0% to 16.0%. Chromophobe renal cell carcinoma provided 9 events for six parameters and is not interpreted (Supplementary Table 16). To test whether the network structure transfers as well as the coefficients, we computed preservation statistics. In CPTAC-3, 13 of 16 lncRNA and 13 of 13 protein-coding modules are strongly preserved by the WGCNA permutation summary statistic^20^ (Zsummary above 10, 100 permutations for the lncRNA and 30 for the protein-coding network). Zsummary increases with module and test-set size, so values are read only against the conventional bands. Every prognostic module is strongly preserved there: lncRNA blue 30.7, greenyellow 15.1 and turquoise 13.4, protein-coding green 61.6 and purple 52.2. In TCGA-KIRP the protein-coding modules and lncRNA turquoise are strongly preserved (34.8, 36.7 and 29.4), while lncRNA blue (8.5) and greenyellow (3.5) fall in the weak-to-moderate band. In TCGA-KICH lncRNA blue is not preserved (1.8) (Fig. 5c, Supplementary Fig. 8, Supplementary Table 17).

### Surviving modules are differentiation, proliferation and malignant-cell programmes

To interpret the surviving modules, we examined their gene content, cellular origin and relations to one another. The protective protein-coding green module is a catabolic programme. Its three strongest Gene Ontology terms are small molecule catabolic process (adjusted p 4.1e-38, 85 of 613 mapped genes), organic acid catabolic process (6.2e-37) and carboxylic acid catabolic process (6.2e-37) (Fig. 6a). The adverse protein-coding purple module is a proliferation programme, with chromosome segregation at 1.5e-44 (Supplementary Data 4). Because these gene sets were selected by correlation, their enrichment p values serve as descriptive rankings. The green module is better described as a proximal-tubule differentiation programme than as catabolism alone. Seventeen of its top 20 hubs have a Human Protein Atlas single-cell specificity string naming proximal tubule^21^, although only LRP2 is exclusive to it. A 21-gene proximal-tubule marker score^21^, including 14 green-module genes, correlates with the green eigengene at r 0.84 (n = 506). In a common set of 502 patients with 165 events, clinically adjusted hazard ratios are 0.69 (0.59 to 0.81) for the eigengene and 0.70 (0.59 to 0.82) for the marker score. The two cannot be separated statistically (Wald p 0.48), so the module’s value beyond the marker score is unresolved (Supplementary Table 18c).

**Fig. 6.**
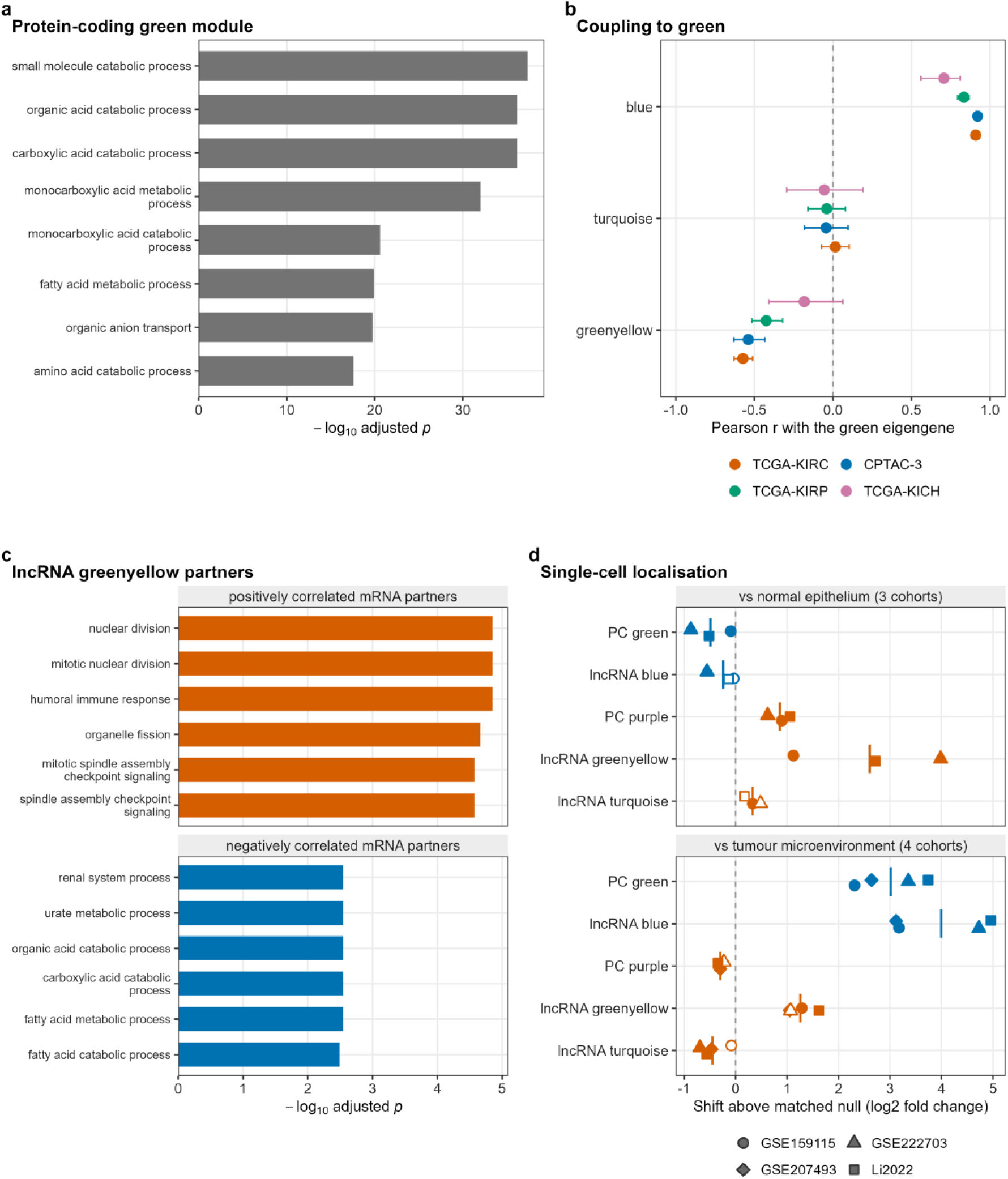
What the prognostic modules are coupled to and where they are expressed. **a** The eight Gene Ontology biological-process terms with the smallest adjusted p values for the protein-coding green module, against the 12,000 protein-coding genes entering the network. Bars are - log10 Benjamini-Hochberg adjusted p. Under the principal specification the module’s hazard ratio is 0.62 per standard deviation (95% confidence interval 0.53 to 0.74, false discovery rate 2.2e-07). **b** Pearson correlation, with 95% confidence intervals, of each prognostic lncRNA module eigengene with the protein-coding green eigengene in 497 TCGA-KIRC, 199 CPTAC-3, 269 TCGA-KIRP and 65 TCGA-KICH patients. Two of three intervals exclude zero in TCGA-KIRC, CPTAC-3 and TCGA-KIRP, and one of three in TCGA-KICH. The turquoise interval spans zero in every cohort. **c** Gene Ontology terms of the protein-coding genes whose expression tracks the lncRNA greenyellow eigengene. The 300 most strongly correlated partners of each sign are tested separately, and the six terms with the smallest adjusted p are shown per sign. The module’s hazard ratio is 1.35 (1.15 to 1.60, false discovery rate 0.0027). Positively correlated partners are enriched for a proliferation and humoral-immune programme, and negatively correlated partners for the catabolic and renal-tubular programme of **a**. **d** Single-cell localisation of the five prognostic modules. For each module and cohort, the median log2 fold change of members in malignant cells against normal kidney epithelium (GSE159115, GSE222703 and Li 2022) and against pooled tumour immune, endothelial and stromal cells (the same three and GSE207493), minus the mean of 2,000 abundance-matched random gene sets of the same gene class. Shape marks the cohort, colour the hazard direction, filled symbols an empirical p below 0.05, and vertical bars the mean over cohorts. Members tested per cohort range from 14 to 18 for greenyellow against normal epithelium and are listed with all values in Supplementary Table 19.

The module is not stromal or immune contamination: 4 of its 670 genes overlap the ESTIMATE stromal signature against 6.4 expected, whereas protein-coding black carries 28 of 323 (p 1.5e-19, Supplementary Table 18e). Nor, in discovery, is it normal-kidney admixture, as no normal library exceeds the tumour maximum on the green eigengene. In CPTAC-3, however, 6 of 168 normals do, so this argument does not extend there (Supplementary Table 18f). Consistent with ccA biology^22,23^, the adverse hazard of ccB class after clinical adjustment (1.62, 1.16 to 2.27) disappears when the green eigengene is added (1.03, 0.67 to 1.58), a partly definitional contrast (Supplementary Table 18d).

Two of the three prognostic lncRNA modules are coupled to this programme, with signs matching the direction of risk. The protective blue eigengene correlates with green at r 0.91 in TCGA-KIRC (n = 497), 0.92 in CPTAC-3 (199), 0.83 in TCGA-KIRP (269) and 0.71 in TCGA-KICH (65), and the adverse greenyellow at -0.57, -0.54, -0.42 and -0.18 (p 0.14 in TCGA-KICH). The adverse turquoise eigengene is uncorrelated with green in every cohort (0.02, -0.04, -0.04 and -0.06) (Fig. 6b). Blue and green are too close to separate: entered together in 493 patients with 159 events, blue carries no independent hazard (1.18, 0.79 to 1.77, p 0.41) against 0.61 for green (0.40 to 0.91, p 0.015), with variance inflation 7.5. By likelihood-ratio test, turquoise adds to green (p 6.9e-04) and greenyellow does not (p 0.19).

Gene Ontology terms of protein-coding partners, split by correlation sign, reproduce the coupling without using module membership. Blue’s positive partners are enriched for small molecule catabolic process and its negative partners for extracellular matrix organization (Supplementary Fig. 9a). Greenyellow’s negative partners are enriched for renal system and organic acid catabolic processes, and its positive partners for a mitotic programme (Fig. 6c). Turquoise’s positive partners are enriched for RNA splicing and its negative partners for proteasomal catabolism, with no green-module overlap (Supplementary Fig. 9b).

Matched normal kidney distinguishes the three modules. Projected onto the 72 normal libraries, only greenyellow keeps its tumour-versus-normal direction after adjustment, lying 0.76 tumour standard deviations below the tumour mean (p 3.2e-11, Supplementary Table 18f). The apparent separation of blue and turquoise in normal tissue is explained by tumour-normal differences in the alignment metrics and collapses on residualisation.

Most members of the three modules are unnamed clone-based accessions (140 of blue’s 246, 68 of greenyellow’s 108 and 321 of turquoise’s 438) (Supplementary Fig. 9c). Three greenyellow members have prior functional studies in this disease: LUCAT1^24^ and LNCSRLR^25^, reported as adverse like the module, and FILNC1^26^, reported as a tumour suppressor, contrary to the module. To locate these programmes in cells, we turned to four independent single-cell cohorts^27,28,29,30^. Donor-level pseudobulk contrasts of malignant cells were available against normal kidney epithelium in three cohorts and against the rest of the tumour microenvironment in four (Methods). Each module’s median member log2 fold change was compared with 2,000 random gene sets of the same gene class matched on single-cell abundance (Fig. 6d, Supplementary Table 19, Supplementary Fig. 10). Green was higher in malignant cells than in the microenvironment in all four cohorts (median log2 fold change 2.3 to 3.6, null near zero) and lower than in normal epithelium in all three (-0.09 to -0.98), the pattern of an epithelial differentiation programme retained by tumour cells but reduced relative to normal tubule. Purple, the proliferation programme, was higher in malignant cells than in normal epithelium in all three cohorts (0.69 to 1.26) but lower than in the microenvironment in all four (-0.31 to -0.52), as expected of a programme shared with cycling immune cells. The lncRNA blue module followed green, higher than the microenvironment in all four cohorts (3.6 to 5.3, with 78% to 89% of tested members individually significant) and below the null against normal epithelium in all three, significantly in one.

The adverse greenyellow module was malignant-specific. Against normal epithelium its members shifted upwards in all three cohorts (median log2 fold changes 1.04, 4.38 and 2.86, empirical p 0.0025, 0.0005 and 0.0005). Averaged across cohorts this is the largest shift of the 16 lncRNA modules, and none of its tested members was significantly lower in malignant cells. It was also higher than the microenvironment in all four cohorts (1.5 to 1.9). In the pooled single-cell meta-analysis, members significantly raised in malignant cells include LUCAT1 and CYTOR against normal epithelium and LNCSRLR and FILNC1 against the microenvironment. Turquoise, the surviving module most exposed to the metric, showed only weak shifts and no malignant-specific enrichment. Its members sat slightly above the null against normal epithelium in all three cohorts (median log2 fold changes 0.24 to 0.82, significant in one) and below it against the microenvironment in all four (significant in three). Coverage is partial because lncRNAs are sparsely detected in single cells: 25% of greenyellow, 44% of blue and 18% of turquoise members were testable against normal epithelium in at least one cohort.

### Modules add no demonstrable value over the clinical model for overall survival

A molecular marker is useful only if it improves on routinely recorded variables, so we measured incremental value over the clinical model by offering all 29 module eigengenes to a penalised Cox model, in 493 discovery patients (159 events) and 199 CPTAC-3 patients (32 events) (Fig. 7a, Table 3, Supplementary Table 20). The apparent concordance-index increment, scoring the locked model on the patients it was fitted to, was +0.021 (0.002 to 0.040, p 0.026). Under ten repeats of ten-fold cross-validation with module definitions fixed on the full cohort, a paired patient bootstrap of the repeat-averaged out-of-fold predictor gave +0.009 (-0.007 to 0.026, p 0.27). Rebuilding both networks and all loadings inside each training fold (three folds, five random fold assignments) gave +0.006 (standard deviation 0.011). Against the augmented model, which adds the ESTIMATE scores and three STAR metrics, the bootstrap increment is +0.010 (-0.007 to 0.029, p 0.22) and the fold-wise rebuild +0.012 (Table 3b).

**Fig. 7.**
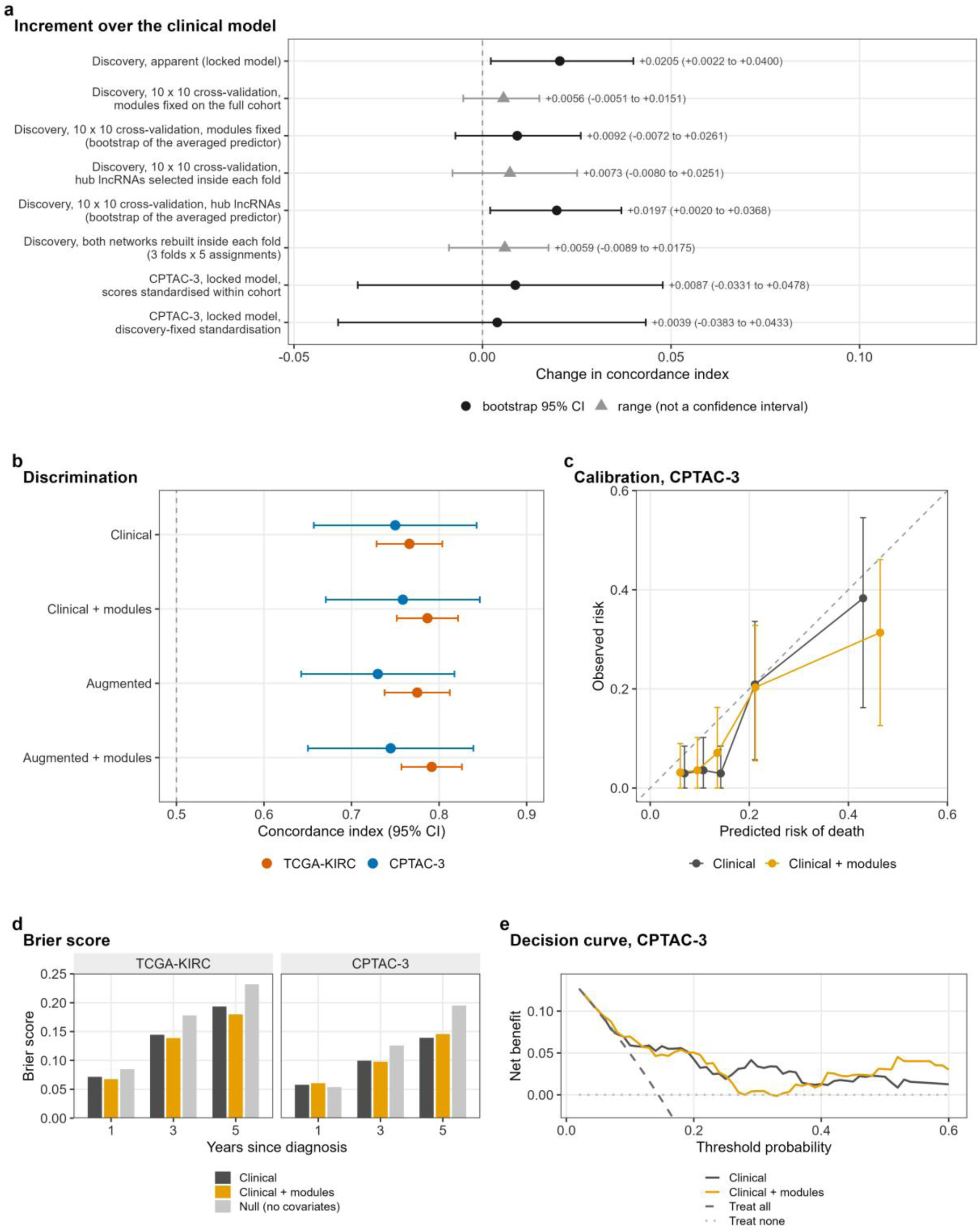
Module eigengenes add no demonstrable discrimination over the clinical model. All panels are for overall survival. The clinical model is age, sex, T category, nodal status, M1 and ordinal grade. The augmented model adds the ESTIMATE stromal and immune scores and the three STAR metrics. Module eigengenes enter a penalised Cox model with the comparator terms unpenalised (Table 3). **a** Increment in concordance index over the clinical model under six estimation schemes (eight rows), in 493 discovery patients with 159 events and 199 CPTAC-3 patients with 32. Black circles carry paired patient bootstrap 95% confidence intervals (2,000 resamples), and grey triangles carry ranges across repeats, which are not confidence intervals. Row one is the locked model scored in sample (+0.0205). Rows two to five are ten repeats of ten-fold cross-validation with module eigengenes (+0.0092) and with hub lncRNAs selected inside each fold (+0.0197), each range row preceding its bootstrap row. Row six rebuilds both networks inside each training fold, over three folds and five assignments (+0.0059). The last two rows are the locked model in CPTAC-3 under cohort (+0.0087) and discovery-fixed (+0.0039) standardisation. **b** Concordance index with 95% confidence intervals for four locked models in both cohorts (493 patients with 159 events in TCGA-KIRC, 199 with 32 in CPTAC-3), eigengenes standardised within cohort. The dashed line marks 0.5, and TCGA-KIRC values are apparent (in-sample). **c** Three-year calibration in CPTAC-3 (199 patients, 32 events) by quintile of predicted risk, with the baseline hazard fixed in discovery. The dashed line is perfect calibration, and error bars are Kaplan-Meier 95% confidence intervals. The calibration slope is 0.97 (0.60 to 1.34) for the clinical model and 0.89 (0.58 to 1.20) with modules, and observed over expected risk is 0.75 and 0.74. **d** Brier scores at one, three and five years, weighted by the inverse probability of censoring (lower is better), for the clinical model, the model with modules and a null model assigning every patient the Kaplan-Meier risk. Modules improve the score at 4 of 6 cohort-by-horizon combinations, three of them apparent TCGA-KIRC scores, and at 1 of 3 horizons in CPTAC-3. **e** Three-year decision curve in CPTAC-3 with the same fixed baseline, thresholds 0.02 to 0.60. The median change in net benefit over thresholds 0.10 to 0.40 is -0.004, and the change is positive at 9 of 31 thresholds. The treat-all curve lies below the plotted range.

**Table 3.**
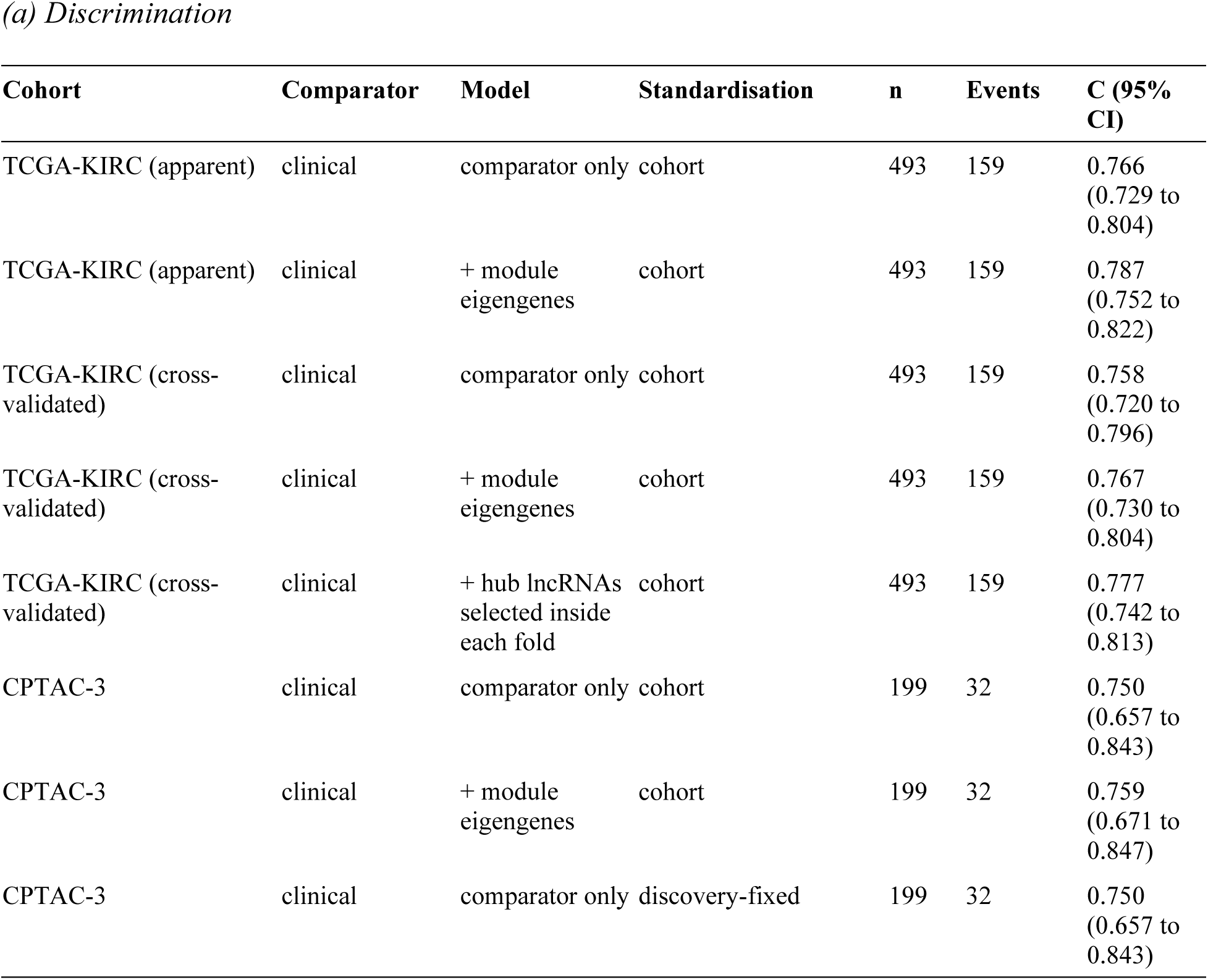
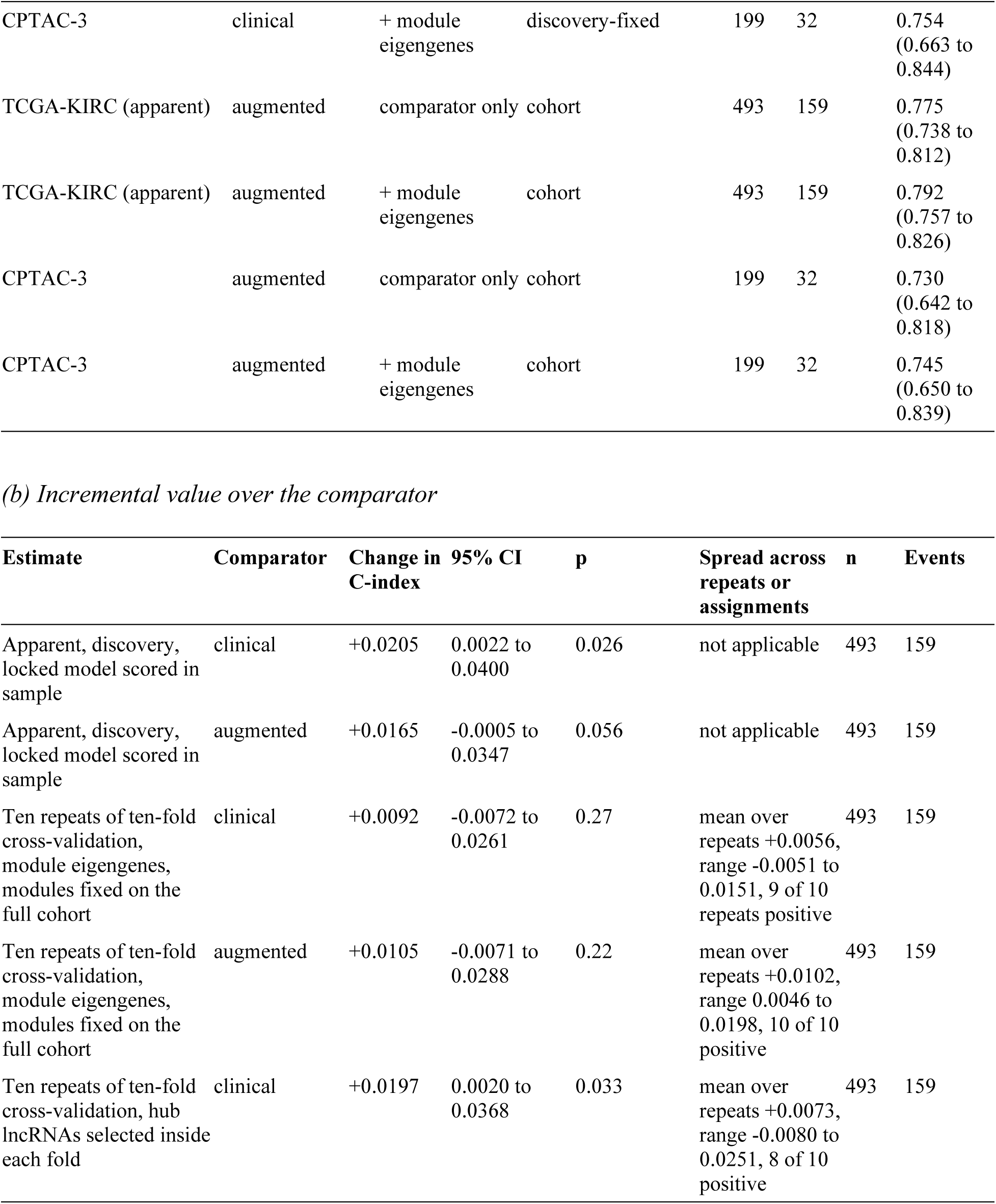
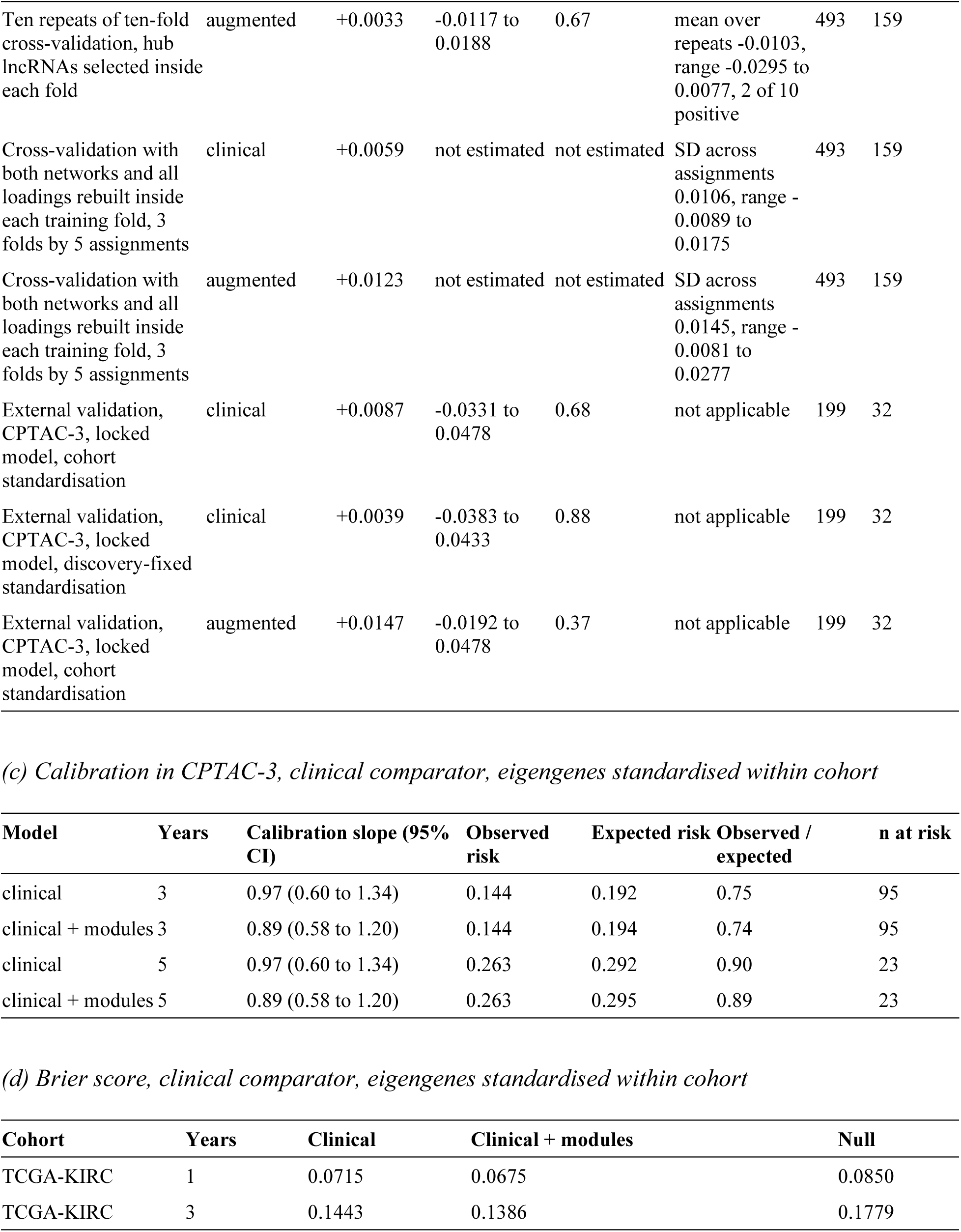
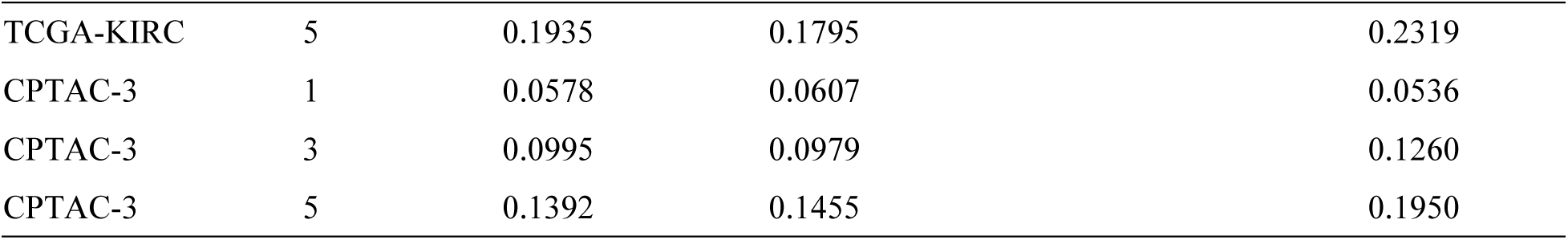
Performance of the locked prediction model for overall survival. The clinical comparator is age, sex, T category, nodal status, M1 and ordinal grade. The augmented comparator adds the ESTIMATE stromal and immune scores and the three STAR alignment metrics. Module eigengenes are the 29 discovery module eigengenes (13 protein-coding and 16 lncRNA), entered into a penalised Cox model with the comparator terms unpenalised. Rows marked apparent in panel (a), and the first two rows of panel (b), are scored on the patients the locked model was fitted to. Rows marked cross-validated are the repeat-averaged out-of-fold predictors of ten repeats of ten-fold cross-validation, so the optimism of the clinical comparator is the difference between 0.766 and 0.758. CPTAC-3 rows apply the discovery coefficients unchanged. Cohort standardisation centres and scales the module eigengenes within the cohort being scored. Discovery-fixed standardisation applies the discovery constants, a fixed rule that does not use the validation population. In panel (b), the change in C-index for a cross-validated row is the increment of the out-of-fold predictor averaged over repeats, the quantity to which the confidence interval and p value belong. The mean over individual repeats is a different quantity, given with its range in the spread column. That range is not a confidence interval. Confidence intervals and p values are from a paired patient bootstrap over 2,000 resamples. The fold-wise rows report the mean, standard deviation and range over five random fold assignments and carry no bootstrap interval. In panel (c), absolute risks use the baseline hazard fixed in discovery. The calibration slope is the coefficient of the linear predictor in a Cox model and does not depend on the horizon, so the same slope appears at three and five years. The n at risk is the number of patients still at risk at that horizon, not the analysis sample size. Under discovery-fixed standardisation the combined model over-predicts risk roughly twofold, with an observed over expected ratio of 0.55 at three years (Supplementary Table 21b, Supplementary Fig. 12b). In panel (d), lower is better, the null model assigns every patient the Kaplan-Meier risk, and Brier scores are weighted by the inverse probability of censoring. TCGA-KIRC rows are apparent (in-sample) scores. No interval is computed for a Brier score, so differences between models should not be read as a ranking. The largest difference in CPTAC-3 is 0.0063, at five years. In CPTAC-3 the combined model is worse than the clinical model at one and five years, and at one year both models are worse than the null.

Individual lncRNAs might carry signal that module eigengenes average out, so we also built a signature of hub lncRNAs, with the candidate pool selected inside each training fold. Over the clinical model it adds +0.007 on average across the ten repeats (bootstrap +0.020, 0.002 to 0.037, p 0.033), and over the augmented model -0.010 (bootstrap +0.003, -0.012 to 0.019, p 0.67) (Supplementary Fig. 11). The lncRNA signal such selection finds largely restates the alignment metrics already in the augmented model.

In external validation the locked model gave a concordance index of 0.759 (0.671 to 0.847) against 0.750 (0.657 to 0.843) for the clinical model, an increment of +0.009 (-0.033 to 0.048, p 0.68), or +0.004 (-0.038 to 0.043, p 0.88) with discovery-fixed standardisation (Fig. 7b, Supplementary Table 21). Dropping each retained module in turn changed the increment by less than 0.02, and dropping all lncRNA modules left it at +0.010, so any external increment comes from the protein-coding modules.

Absolute risk and clinical utility do not favour the combined model either. In CPTAC-3 its calibration slope was 0.89 (0.58 to 1.20) against 0.97 (0.60 to 1.34) for the clinical model, and the three-year observed-to-expected ratio 0.74 against 0.75, so over-prediction in this lower-risk cohort comes from the transported baseline, not the modules (Table 3c, Fig. 7c). Brier scores favour the clinical model at one and five years and the combined model at three (Table 3d, Fig. 7d). At three years, decision-curve analysis gave a median change in net benefit of -0.004 (-0.022 to 0.006) over thresholds 0.10 to 0.40, with medians near zero in the 0.05 to 0.30 window and at five years. The curves show neither consistent benefit nor consistent harm (Fig. 7e, Supplementary Fig. 12, Supplementary Table 21).

This conclusion is specific to overall survival. For Clinical Data Resource endpoints the cross-validated increment was +0.023 for disease-specific survival (0.004 to 0.045, p 0.017, 482 patients, 99 events) and +0.014 for the progression-free interval (0.001 to 0.027, p 0.036, 491 patients, 146 events), against +0.008 for overall survival. Neither survives Benjamini-Hochberg adjustment (0.051 and 0.054 over three distinct endpoints, 0.068 and 0.072 if overall survival from the pipeline and the Clinical Data Resource counts twice). Under three-fold cross-validation the disease-specific increment falls to +0.013 whether modules are fixed or rebuilt in every fold. CPTAC-3 has neither endpoint (Supplementary Fig. 13, Supplementary Table 8). After clinical adjustment the metric itself predicts overall survival (1.36, 1.20 to 1.55) and disease-specific survival (1.31, 1.10 to 1.55) but not the progression-free interval (1.11, 0.95 to 1.30), and the axis behaves the same way (Supplementary Table 8).

## Discussion

The leading axis of lncRNA expression in TCGA-KIRC tracks a routine alignment metric, the non-feature read fraction (Spearman rho 0.90), and sets the structure of the lncRNA co-expression network. Without adjustment the lncRNA network collapses into one module holding 58% of genes. Adjusting for the three alignment metrics changes which modules exist and uncovers modules that the unadjusted network had left largely unassigned. With membership held fixed, it also cuts the significant modules from 15 of 29 to 10. The surviving modules keep the direction of their survival association in an external cohort and are preserved across library protocols, and in single cells three of them localise to malignant or epithelial cells. For overall survival they add no demonstrable discrimination over recorded TNM and grade. Across TCGA, under rules fixed before any non-renal type was analysed, a lncRNA-specific axis occurs in 9 of 25 cancer types and the network collapses in 4, renal clear-cell among them. Among the 23 non-renal types, the counts are 8 and 2.

Whether adjustment is appropriate depends on what the metric measures, and two readings lead to opposite conclusions. If it is pre-analytical, it indexes how the specimen was collected, stored and made into a library. If it is tissue-intrinsic, because necrotic tumour or tumour with impaired RNA processing yields more unprocessed RNA and coagulative necrosis is part of the SSIGN score^1^, residualising on it would remove biology. The evidence favours a large pre-analytical component. RNA integrity, plate and tissue source site explain 36.2% to 36.8% of its variance, against 6.1% for annotated tumour biology, and the axis is present in tumour-free kidney. Two further results bear on this. First, single-cell data, which do not depend on bulk library preparation, show no loss of tumour biology after correction. In the unpaired tumour-normal contrast, which the correction can alter, agreement with malignant-cell specificity was 0.773 before and 0.774 after, and a tumour-specific lncRNA score kept its excess coherence with malignant markers. This is consistent with the correction removing pre-analytical variation and sparing biology, though it does not show that everything removed was artefact. Second, the metric has no consistent association with death across cancer types. In renal cancer it is prognostic within sequencing plate in discovery, at 1.23 (1.02 to 1.50), and weaker through the pan-cancer pipeline, which uses a different endpoint source and stage in place of TNM and grade, at 1.13 (0.96 to 1.33). Across 20 cancer types the pooled hazard ratio is 1.00 (0.93 to 1.08), with heterogeneity and nominal associations in both directions. This argues against a general marker of tumour aggressiveness while leaving room for a tissue-specific effect. A third reading would explain why the metric predicts death in renal cancer: it is pre-analytical but confounded with outcome through specimen handling. A longer time from devascularisation to freezing, plausible in larger and more complex nephrectomies, would both raise the metric and mark higher-risk tumours. This reading fits the within-plate association, which survives adjustment for RNA integrity, and the finding that the recorded variables leave most of the metric’s variance unexplained. These data test it only partly. T category is the only size proxy recorded, and its T1 and T2 categories are defined by size (T1a at most 4 cm, T1b 4 to 7 cm, T2 above 7 cm), yet it shows no association with the metric (rho 0.06, p 0.21, and no clinical variable reaches a smaller p). The handling interval is not recorded for TCGA-KIRC. CPTAC-3 records the time from excision to freezing, at most 30 minutes under its protocol. A post hoc analysis in CPTAC-3 found that handling time was longer in higher-grade tumours (Spearman rho 0.32) but did not track the metric (-0.02, 175 tumours) or the axis in tumour or normal tissue (Supplementary Table 22). The protocol cap, the different library protocol and the metric’s lack of prognostic value in that cohort limit what the CPTAC-3 analysis can rule out.

The network result follows from how a signed, soft-thresholded co-expression network treats a shared axis. A variable that moves thousands of genes together raises every pairwise correlation among them, the soft threshold amplifies those correlations, and the most exposed genes, the lncRNAs lying inside or opposite a protein-coding gene, merge into one module. The positional gradient appears in every cancer type examined, so the mechanism is not specific to renal cancer. The within-cohort spread of the metric, which tracked the strength of the axis in an exploratory analysis, can be measured before any network is built. Adjusting for such covariates before building networks that include lncRNAs is established practice in other fields^31^, and the choice of adjustment is known to reshape network architecture^32^. Principal-component correction of TCGA correlations has so far excluded lncRNAs^33^, and the TCGA lncRNA literature has not adopted covariate adjustment. Two consequences follow. First, because the dominant module holds most of the unadjusted network, the modules, hubs and guilt-by-association annotations of an unadjusted lncRNA network largely describe the metric. Second, reproducibility does not separate technical from biological structure. Turquoise lay almost wholly inside the dominant module before adjustment, and its residualised eigengene still correlates with the metric (Spearman 0.20), yet it is strongly preserved in CPTAC-3 and TCGA-KIRP (Zsummary 13.4 and 29.4). Preservation alone does not show biology, so a check independent of library preparation is needed. The same applies to analyses that read the TCGA intronic signal as transcriptional biology^34^.

Cell-type localisation provides that check and shows what the surviving modules represent. Protein-coding green is enriched for small-molecule and organic-acid catabolism, the metabolic reprogramming described in this disease^35,36^. Its hubs are proximal-tubule genes^21^, and a 21-gene proximal-tubule marker score reproduces its hazard in the same patients. In single cells from every cohort, malignant cells express green far above the microenvironment and below normal epithelium, the signature of a differentiation programme retained to varying degrees by tumour cells. This explains its protective association and why it absorbs the ccA and ccB contrast^22^. Purple is a proliferation programme raised in malignant cells over normal epithelium. The lncRNA blue module follows green in bulk (r 0.91) and in single cells and carries no hazard of its own once green is in the model. Greenyellow is the one lncRNA programme that is both malignant-specific and prognostic. It is the lncRNA module most enriched in malignant cells over normal epithelium, is anti-correlated with the proximal-tubule programme and is coupled to proliferation, and its members raised in malignant cells include LUCAT1^24^ and CYTOR. In a joint model, however, it adds nothing over green (p 0.19), so its prognostic content is shared with the differentiation axis. Turquoise, the only lncRNA module that adds prognostic information to green (p 6.9e-04), shows no malignant-specific enrichment and was not nominally significant in CPTAC-3. None of the three lncRNA modules therefore carries prognostic information that is both independent of the protein-coding programmes and supported externally.

Applied to published TCGA-KIRC lncRNA signatures, the same test gives a mixed result. All six re-scored signatures correlate with the metric, and adjusting for it abolishes none. The clinically adjusted log hazard ratio moves by 2.6% and 4.9% for the two signatures that publish coefficients, and by 11.1% to 19.0% for the three whose hazard direction was set in sample, a figure inflated by that in-sample weighting. All six signatures were derived in TCGA-KIRC, and three had their hazard direction set in the same patients, so their increments over a clinical model in this cohort, 0.021 and 0.024 for the two with published coefficients, are apparent figures and are likely to be optimistic. We did not cross-validate these signatures, so the size of that optimism is not measured. Our own modules suggest its order of magnitude, with an increment of 0.021 apparent and 0.009 under cross-validation.

Against the strongest clinical comparator available, the modules add nothing demonstrable for overall survival, with increments of 0.009 (-0.007 to 0.026) in cross-validation and 0.009 (-0.033 to 0.048) in CPTAC-3, and calibration, Brier scores and net benefit do not favour the combined model. Hub lncRNAs selected inside each training fold add 0.020 over the clinical model but 0.003 over one that already contains the alignment metrics, so they largely restate those metrics. The disease-specific and progression-free increments, 0.023 and 0.014, do not survive Benjamini-Hochberg correction across endpoints and rest on one cohort. The comparator and cohort sizes set the scope of this result. Tumour size and performance status are not recorded, so the comparator is weaker than the composite scores used clinically and the reported increment is an upper bound on what a fully staged comparator would leave. CPTAC-3 contributed 32 deaths against the 100 to 200 recommended for external validation of a survival model^37^, and in discovery the smallest increment cross-validation could separate from zero was about 0.024. The result therefore bounds the gain in these cohorts without showing it to be zero. Overall survival counts deaths from any cause, including competing mortality, and no competing-risks model was fitted. No clinical test is proposed here.

The remaining limits concern the adjustment, the biology of the metric, the single-cell evidence and the pan-cancer design. Adjustment was per-gene linear residualisation on three STAR metrics. Module identities depend on the correction method (adjusted Rand index 0.23 against RUV-III) and should be interpreted with caution. The residual correlation of turquoise with the metric shows that a linear adjustment can leave some dependence on it. Biospecimen and batch variables exist for TCGA-KIRC only. In clear-cell renal cancer SETD2 loss causes intron retention and transcriptional read-through^38,39^, which would raise the metric biologically. SETD2-mutant tumours had a median metric 0.47 points above wild-type (46 of 368, p 0.54), small beside the plate differences, and the positional gradient appears in cancer types where SETD2 loss is rare. A tumour-intrinsic component in renal cancer remains possible. The metric indexes non-exonic read content and does not measure RNA degradation directly. Severe degradation lowers the intronic share of poly(A) libraries, because degraded pre-mRNA loses the poly(A)-captured intronic signal^40^. A moderate delay before freezing, with the RNA still largely intact, would instead plausibly add incompletely processed and read-through transcripts and raise the intronic share. The metric’s negative correlation with RNA integrity (-0.25) is compatible with the second process and not with the first, although it is also compatible with the tissue-intrinsic reading. Gene-level union counting, as in the GDC pipeline, under-quantifies overlapping and antisense lncRNAs^41^, and the controlled-access reads needed to requantify them were not used. Single-cell localisation covers 18% to 44% of lncRNA module members against normal epithelium, rests on donor-level inference in three or four cohorts, one with malignant labels transferred from another, and shows where a programme is expressed, not whether it drives outcome. All four single-cell cohorts are droplet data from dissociated whole cells, counted without intronic reads (Methods). The lncRNAs detectable in them are therefore biased towards abundant, polyadenylated, independently transcribed transcripts, whereas the classes carrying most of the metric’s signal are those overlapping protein-coding loci. The single-cell test of the correction thus covers the lncRNAs least exposed to the metric, and it was run in one bulk cohort. Function was inferred by guilt-by-association, and because the two networks were built separately one programme can appear in both, as blue and green do. The pan-cancer analysis applied the discovery network parameters to every cancer type without tuning and used one overall-survival endpoint, with stage where recorded, and its dose-response analysis was specified after the locked rules had been applied. Current bulk transcriptomic foundation models cover few or no lncRNAs^42,43^ and cannot yet stand in for these modules. Finally, the sex-disaggregated estimates are descriptive, both cohorts are convenience samples of consented research participants, race and ethnicity were incompletely recorded and not modelled, and the study was not registered.

Four recommendations follow for network and signature studies of bulk lncRNA expression. First, report per-sample alignment metrics and their within-cohort spread, and test whether the leading axes of the expression matrix and the eigengene of its largest module track them, preferably with a transcript-integrity measure computed from the sequencing files^44^. Second, where they do, build the network on adjusted expression and report module-level associations with and without adjustment. Third, do not treat module preservation as evidence that a module is biological, and check cellular localisation or another readout independent of library preparation. Fourth, benchmark prognostic claims against full TNM with grade, cross-validate feature definition, and validate across library protocols, stating the protocol difference. A routine alignment metric carries the leading axis of lncRNA expression in these renal cancer cohorts and, to a varying degree, across TCGA. It sets the structure of the lncRNA co-expression network and is itself structured by processing batch, and correcting for it does not detectably remove the tumour biology that single cells report. Adjustment changes the network and which modules look prognostic, and it shifts the hazard ratios of published signatures without abolishing any. The surviving programmes are proximal-tubule differentiation, proliferation and a malignant-cell lncRNA programme coupled to both. They keep the direction of their survival association in an external cohort, transfer across protocols and localise to specific cell types. None of this makes lncRNA expression uninformative in renal cancer. It means that a network or prognostic claim made from it can be assessed only when the alignment metrics are reported alongside it.

## Methods

### Cohorts and data sources

Four renal cancer cohorts were quantified by one Genomic Data Commons STAR-Counts pipeline^45,46^, giving comparable gene identifiers and FPKM definitions without cross-platform rescaling. TCGA-KIRC was the discovery cohort^35^, CPTAC-3 the external validation cohort^47,48^, and TCGA-KIRP and TCGA-KICH the subtype comparators. TCGA libraries are poly(A)-selected, whereas CPTAC-3 libraries are ribo-depleted stranded total RNA. All libraries came from primary tumour resection specimens. Systemic and adjuvant therapy is not recorded in the harmonised fields and was not modelled, so an unrecorded fraction of each cohort will have received systemic therapy after nephrectomy. TCGA-KIRC tumours came from 20 tissue source sites, and CPTAC-3 contributing institutions are not identified. Grade is Fuhrman nuclear grade in both cohorts. Grade and pathological stage were assigned by the contributing institutions’ pathologists and not re-reviewed.

Expression files, manifests, sample types and harmonised clinical fields came from Genomic Data Commons Data Release 46.0 (10 August 2026). TCGA-KIRC grade came from the Biospecimen Core Resource clinical XML supplements, somatic calls from the open-access masked MAF files of the aliquot-ensemble whole-exome workflow, and gene annotation from GENCODE v36^49^. STAR-Counts are gene-level, so every analysed feature is a gene. Four TCGA-KIRC barcodes carry two files, so expression, library size and STAR metrics were selected by file identifier.

### Sample inclusion and clinical variables

Only primary tumours were analysed. A library failed if assigned reads were below 10 million or the non-feature read fraction exceeded 30%. The non-feature threshold is an outlier rule within one library protocol and was applied to the poly(A) cohorts only, because under ribo-depleted total RNA the whole distribution of the metric lies above 30%. No CPTAC-3 library failed on read depth, and five TCGA-KIRC libraries failed. The rule was applied per file, before selecting one sample per patient, so a patient with a second usable aliquot kept it. Where a patient still had several aliquots, the one with the most assigned reads was retained.

Overall survival was days to death for deceased patients. For censored patients it was the longest follow-up across the Genomic Data Commons diagnosis and follow-up records and the Biospecimen Co re Resource follow-up records, because the harmonised field alone is missing for most living TCGA-KIRC patients. A death recorded in either source counted as an event. Follow-up was administratively censored at ten years, reached by 13 discovery patients. Median follow-up used reverse Kaplan-Meier. Patients with non-positive or missing survival time were excluded.

Stage was parsed from the roman numeral of the pathological stage field, ignoring the sub-stage letter, and T category from the pathological T field, never imputed from stage. NX and MX were coded N0 and M0 and flagged, following the SSIGN and Leibovich convention of assigning pNX to pN0, because lymphadenectomy is not routine in this disease. Patients with AJCC stage IV, a T category below T4 and no explicit M category were reconciled to M1 (2 TCGA-KIRC and 12 CPTAC-3 patients), and an explicit M0 was never overridden. Grades GX and not reported were treated as missing. Sex came from the harmonised clinical record, with female as reference. No primary analysis was disaggregated by sex, and the sex analyses (Additional analyses) are sensitivity analyses. Race and ethnicity, incompletely recorded in both programmes, were not modelled, and no fairness assessment was performed.

CPTAC-3 cases were retained with a primary diagnosis of clear cell or renal cell carcinoma, not otherwise specified, because the programme codes its clear-cell cohort under the latter term. The harmonised fields carry no accrual dates, so no accrual window is stated, and the Genomic Data Commons snapshot of 10 August 2026 is the effective end of follow-up.

### Ethics

The TCGA and CPTAC programmes obtained written informed consent from every participant under the approvals of their contributing institutions. This secondary analysis of de-identified, publicly available data required no further ethical approval. No patients or members of the public were involved in the design, conduct or reporting of the study.

### Expression processing and adjustment for the alignment metrics

Analyses used log2(FPKM + 1) of the Genomic Data Commons unstranded FPKM values, whose denominator is the number of reads mapped to protein-coding genes. More unprocessed RNA in a library raises both the non-feature fraction and the intronic reads counted to overlapping lncRNA loci, inflating lncRNA FPKM relative to protein-coding FPKM. The unstranded column was used in every cohort so that one quantity is compared throughout. For the stranded CPTAC-3 libraries it sums both strands, raising apparent antisense lncRNA expression relative to a strand-aware count. Protein-coding genes were retained with FPKM of at least 1.0 in at least 25% of samples, and lncRNAs with FPKM of at least 0.30 in at least 20%. The most variable genes by median absolute deviation were then kept, up to 12,000 protein-coding genes and 6,000 lncRNAs. The protein-coding cap was reached (12,000 of 19,962 genes), and 3,442 of the 16,889 unique GENCODE v36 lncRNA genes passed the abundance filter. Duplicate gene symbols kept the most variable Ensembl entry.

The four STAR summary rows (unmapped, multimapping, non-feature and ambiguous reads) were expressed for every library as percentages of the sum of those four counts and the reads assigned to annotated genes. The non-feature read fraction, the principal metric, is therefore the percentage of all reads in the STAR gene-counts summary assigned to no annotated feature. With the multimapping fraction and log10 assigned reads, it forms the three alignment metrics. Adjustment for the alignment metrics was applied to expression before network construction. Each gene was regressed by ordinary least squares on the three standardised metrics, and its residuals were kept with the gene mean added back. The stored centring, scaling and coefficients allow held-out samples to be adjusted with training-fold coefficients. The adjustment was fitted within each cohort and never transported between cohorts, because the CPTAC-3 non-feature fraction lies far outside the TCGA range. All analyses downstream of network construction use the adjusted matrix unless the observed matrix is named.

Tumour composition was not removed from expression, because stromal and immune content are prognostic in this disease. ESTIMATE stromal and immune scores from the full protein-coding matrix were instead entered as covariates^50^. The illumina platform setting does not return the Affymetrix-calibrated purity value, so the ESTIMATE score is an inverse-purity proxy.

### Biospecimen, plate and batch variables

For TCGA-KIRC, RNA integrity number, A260/A280 ratio, slide percent necrosis and slide percentages of tumour nuclei, stromal cells and normal cells were taken from the Genomic Data Commons biospecimen records, averaging the integrity number over the RNA analytes of the primary-tumour sample. One integrity number recorded as zero is a coded missing value, excluded from Supplementary Fig. 4a, b and retained elsewhere as one of the 527 recorded values. Sequencing plate and tissue source site were fields 6 and 2 of the 28-character RNA aliquot barcode, and Biospecimen Core Resource batch was the administrative batch number in the clinical XML. TCGA centralised extraction, library preparation and sequencing, so tissue source site is a proxy for collection, ischaemia and freezing practice. Levels with fewer than ten patients were pooled into an “other” stratum, recomputed on each analysis’s complete cases. The other three cohorts lack equivalent metadata, so these analyses are restricted to TCGA-KIRC.

### Network construction and module definition

Signed weighted gene co-expression networks with signed topological overlap were built separately for protein-coding genes and lncRNAs with WGCNA^51^, in a single block of up to 25,000 genes. Samples with standardised connectivity below -2.5 were removed as outliers, leaving 506 samples for the protein-coding network and 511 for the lncRNA network. Outlier removal preceded adjustment for the alignment metrics, followed by power selection and module detection, so module definitions do not inherit the axis. The two gene classes differ in abundance and variance scale and need separate soft thresholds, so they were kept apart. One programme can therefore appear as a module in each network, as lncRNA blue and protein-coding green do. For the protein-coding network the soft-thresholding power was the lowest reaching a signed scale-free fit R-squared of at least 0.85 over a grid of 1 to 10 and 12 to 30 in steps of 2, giving power 18 (R-squared 0.894). For the lncRNA network a parameter sweep fixed the power at 10 (R-squared 0.811), where the automatic rule would have given 12.

The sweep scored 96 configurations (three expression variants, four powers, two deep-split values, two merge heights and two minimum module-membership thresholds) with the network-construction call of the final network, without consulting survival, stage or grade. A configuration was admissible if its largest module held at most 35% of genes, at most 30% of genes were unassigned and 5 to 40 modules were produced, and 33 qualified. The eight configurations with the smallest largest module were ranked by mean adjusted Rand index against assignments rebuilt on ten random 80% patient subsamples. The selected configuration (power 10, deep split 2, merge height 0.15, minimum module membership 0.3) had a mean adjusted Rand index of 0.605, which counts unassigned genes as one class and so inflates agreement.

Modules were detected with a minimum size of 30, with the study seed set immediately before the call. Excluding the grey module throughout left 29 module eigengenes. Module-trait correlations used only age, sex, stage and grade, never observation time.

### Module eigengenes and projection

A module eigengene is the first principal component of the module’s gene-standardised expression, sign-aligned to correlate positively with its genes’ mean standardised expression and scaled to unit variance. This definition was used everywhere. The rotation was fitted once on the adjusted discovery matrix, and the stored gene means and standard deviations, loading vector and score constants were applied unchanged elsewhere. Genes and eigengenes are standardised within the cohort being scored, and every validation quantity was also computed with the discovery constants.

The leading lncRNA axis is the first principal component of the column-centred, unscaled observed lncRNA matrix of the 511 network samples, sign-aligned to per-sample mean expression. The same computation on the observed protein-coding matrix of 506 samples is the negative control. In comparisons across sample sets, the other set was centred on the discovery gene means, projected onto the discovery loadings and scaled by the discovery standard deviation, so one fixed quantity is compared. Each set’s own leading component is a different quantity and is labelled as such.

### Survival analysis

Three specifications were fitted for every module, without imputation. The unadjusted specification contains the module eigengene alone. The clinical model adds age, sex, T category, nodal status, M1 and ordinal grade, and the principal specification further adds the ESTIMATE stromal and immune scores and the three alignment metrics. Residualisation removes each gene’s linear dependence on the metrics by construction, so a near-zero correlation after adjustment does not show that their influence is gone. The metrics therefore also enter the principal specification as covariates, where they can absorb non-linear and score-level dependence. T category and grade were coded 1 to 4, and nodal status and M1 as binary. ESTIMATE and alignment-metric terms were standardised over all network samples. Kaplan-Meier curves use a median split and a log-rank test.

The false discovery rate was controlled by the Benjamini-Hochberg procedure^52^ at 5% within each network (13 protein-coding or 16 lncRNA modules) and is also reported over all 29 modules. Log-rank tests were adjusted in the same family, and the false discovery rate of the CPTAC-3 estimates was controlled across the five modules carried forward. Subtype estimates are nominal. Proportional hazards were tested by scaled Schoenfeld residuals for the eigengene term, globally for each model and for the assembled risk scores.

The leading lncRNA axis was tested in a nested sequence on one complete-case set: the axis alone, then successively adding the clinical terms, the ESTIMATE scores and the three alignment metrics, with the variance inflation factor at each step. The sequence was repeated adding the integrity number, adding the integrity number and slide necrosis, and stratifying by sequencing plate. A parallel nested sequence took the non-feature read fraction as the exposure and ended with stratification by tissue source site. Being right-skewed, the metric was fitted on both the linear and the base-ten logarithmic scale.

Because nodal status was unassessed in half of the discovery cohort, three further specifications were fitted for every prognostic module. They coded unassessed nodal and metastatic status as separate categories, restricted analysis to patients with both assessed, and took M1 as coded without reconciliation from stage IV. SSIGN, Leibovich and UISS scores could not be reconstructed, because tumour size is absent from every TCGA-KIRC clinical XML supplement and performance status is recorded for a minority of patients. The recorded slide percent necrosis has a median of 0 and is not the whole-tumour coagulative necrosis those scores require. TNM as recorded, with ordinal grade, is therefore the strongest clinical comparator available.

### Prediction models and internal validation

Penalised Cox models used an elastic-net mixing parameter^53^ of 0.5, because a pure lasso did not converge on near-collinear hub lncRNAs. The penalty was chosen by cross-validated partial likelihood inside each training fold over a 50-value path, and comparator covariates were never penalised. The analysis set was the 493 discovery patients present in both networks with complete clinical inputs and survival (159 events).

Two feature representations were compared. The eigengene representation offers all 29 module eigengenes from the adjusted matrices, without pre-selection. The hub representation offers individual lncRNAs on observed expression, closer to what a deployable assay would measure. Its candidates were chosen inside each training fold as the 200 genes with the highest module membership in lncRNA modules reaching a 5% false discovery rate in per-module Cox models on the training rows.

Discrimination is Harrell’s concordance index^54^ on out-of-fold linear predictors from 10 repeats of 10-fold cross-validation. Three interval types are reported and are not interchangeable. A paired patient bootstrap of 2,000 resamples on the repeat-averaged out-of-fold predictor gives a confidence interval and p value. The 2.5th and 97.5th percentiles across the ten repeats measure cross-validation noise, and the minimum and maximum across fold assignments give a range.

### Apparent concordance is labelled in-sample

Module definition was also cross-validated. In three-fold cross-validation over five random fold assignments, each training fold was residualised on its own alignment metrics, both networks were rebuilt and loadings fitted on it, and held-out patients were scored with that fold’s centring, scaling and rotation. The comparator was cross-validated on the same assignments, and a matched fixed-module arm used identical folds and inner splits.

No formal sample-size calculation was performed, and all eligible patients were used. The bootstrap interval for the cross-validated increment has a half-width of 0.017, implying a standard error of about 0.008, so the smallest concordance increment the design could separate from zero at 80% power is about 0.024. CPTAC-3 carries 32 events, well below the 100 to 200 recommended for external validation of a survival model^37^.

### External association estimates and predictive validation

Two validation analyses were run in CPTAC-3 and reported separately, because they hold different quantities fixed. The external association analysis fixed module membership and eigengene loadings in discovery, projected CPTAC-3 expression onto those loadings without recomputing components, and re-estimated only the Cox coefficient. Two covariate sets were fitted and both are reported. The twelve-parameter principal specification from discovery gives 2.7 events per variable on 32 events, below the threshold of five applied to the subtype cohorts. A parsimonious six-parameter set (eigengene, age, sex, T category, M1 and ordinal grade) gives 5.3 and meets that threshold, so replication of a module was defined as a false discovery rate below 0.05 in this arm, controlled across the five modules. Directional agreement across the five modules was summarised by a one-sided sign test. This analysis tests an association, and the predictive validation tests a fixed prediction rule.

The predictive validation also fixed the penalised coefficients in discovery and applied the linear predictor unchanged to CPTAC-3. Absolute risks use a Breslow baseline hazard estimated in discovery with the locked predictor as an offset, so validation risks are out-of-sample predictions. Calibration is summarised by the calibration slope, refitted in the cohort being scored, and by the Kaplan-Meier ratio of observed to expected risk. Brier scores use inverse-probability-of-censoring weighting^55^ and carry no interval. Decision-curve analysis^56^ on the discovery baseline assessed clinical utility at thresholds of 0.02 to 0.60, with net benefit summarised over the windows 0.10 to 0.40 and 0.05 to 0.30 by a paired bootstrap of 500 resamples of the median gain. No threshold is proposed as a decision rule. Three years is the primary horizon, because few CPTAC-3 patients remain at risk at five.

The alignment metrics are not comparable across library protocols. The CPTAC-3 non-feature fraction has a median of 40.5% against 4.6% in TCGA-KIRC, so a discovery-standardised value would lie many standard deviations from the discovery mean and dominate any linear predictor containing it. The expression-level adjustment and the augmented model are therefore refitted within each cohort, and both standardisation arms are reported for every validation quantity. A drop-one-module test set each retained coefficient to zero individually, then all retained lncRNA module coefficients together, and re-bootstrapped the change in concordance.

### Subtype specificity and module preservation

TCGA-KIRP and TCGA-KICH primary tumours were scored through the same path as every other cohort: log2(FPKM + 1) on the network gene sets, within-cohort adjustment for the alignment metrics, then the fixed discovery loadings with cohort standardisation. The same parsimonious adjustment set (age, sex, T category, M1 and the non-feature read fraction, six parameters per model) was used in TCGA-KIRC, TCGA-KIRP and TCGA-KICH, and only cohorts with at least five events per variable contributed to inference. TCGA-KIRC (26.7) and TCGA-KIRP (6.7) qualify and TCGA-KICH (1.5) does not, so TCGA-KICH estimates are descriptive and a null there is uninformative. Between-subtype heterogeneity was tested by Cochran’s Q with I-squared across qualifying cohorts only.

Module preservation was assessed with the WGCNA module preservation statistics^20^, with the adjusted discovery matrix as reference and test networks built as in the external and subtype analyses. Permutations numbered 100 for the lncRNA network and 30 for the protein-coding network, and test module size was capped at 1,000. A Zsummary above 10 was read as strong preservation, 2 to 10 as weak to moderate, and below 2 as none.

### Functional characterisation

Gene Ontology biological-process over-representation was tested with clusterProfiler^57^ against a universe of the 12,000 network protein-coding genes, with Benjamini-Hochberg adjustment and cut-offs of 0.05 on the adjusted p value and 0.10 on the q value. Because partner sets are selected by correlation with an eigengene, these p values are uncalibrated and serve only as descriptive rankings.

Because lncRNAs carry little direct annotation, prognostic lncRNA module function was inferred by guilt-by-association. Protein-coding genes were ranked by absolute correlation with the module eigengene, and over-representation was tested in the top 100, 300 and 500 partners on both the observed and the adjusted matrix. Partners were also split by correlation sign, and each side’s top 300 was tested by a separately implemented hypergeometric test with the same annotation and universe. This sign-split analysis correlated residualised lncRNA eigengenes with the observed protein-coding matrix, so its partner sets can carry variance from the alignment metrics. Coupling with the catabolic module was tested directly by correlating each prognostic lncRNA eigengene with the catabolic protein-coding eigengene in each cohort. The catabolic module is identified by its Gene Ontology enrichment, because colour labels change whenever a network is rebuilt. Named module members were screened by gene symbol against the renal cell carcinoma literature, without a systematic search. LNCSRLR is NCBI Gene 109729161 (HGNC:52753), retrieved 23 August 2026.

A marker score tested whether the catabolic module reflects retained proximal-tubule differentiation. It is the mean per-gene z score of documented proximal-tubule markers on the observed matrix^21^, computed for all 21 markers, the 14 inside the analysed network, those whose Human Protein Atlas specificity names proximal tubule, and the 7 outside the network. Tumours were placed on the ccA and ccB axis^22^ by correlating each standardised 34-gene profile with a template of +1 for ccA genes and -1 for ccB genes^23^. The published centroids are not distributed, so this score reproduces the ccA and ccB axis but not the classifier. Every module eigengene was evaluated both as adjusted and as refitted on the observed matrix, because only the latter can carry variance from the metric or from admixture.

### Additional analyses

Correlates of the metric. Candidate correlates of the non-feature read fraction were tested in TCGA-KIRC: a 51-gene hypoxia metagene^58^, a proliferation score, the ESTIMATE scores, the slide tumour-nuclei percentage, and the eigengenes of both networks on the observed and residualised matrices. The proliferation score is the mean per-gene z score of log2(FPKM + 1) over MKI67, TOP2A, PCNA, MCM2, BUB1, CCNB1, CDK1, TYMS, RRM2 and AURKA, a set defined ad hoc without an external source. Variance in the metric was partitioned between a block of annotated biology and a technical block of RNA integrity, sequencing plate and tissue source site. Any block built from observed expression recovers the metric from the matrix it was measured on, so such rows are reported only beside negative controls (eigengenes of size-matched random gene sets and the leading principal components of the same matrix). Attenuation was assessed by adding each candidate to the clinically adjusted exposure model, and rows whose added covariate is largely the exposure itself are labelled collinearity diagnostics (Supplementary Table 9).

Matched normal libraries. The 72 TCGA-KIRC and 168 CPTAC-3 Solid Tissue Normal files were processed as the tumours were, except that only the read-depth criterion of the library-failure rule applies to CPTAC-3. The discovery axis was projected onto each set, and each set’s own leading component was computed and compared with a size-matched baseline of 200 discovery subsamples. Tumour-versus-normal differences are reported unpaired, within patient, and restricted to plates occupied by both sides, because the unpaired contrast is confounded by plate (Supplementary Table 3).

Published signatures. Six published TCGA-KIRC lncRNA signatures were re-scored on the discovery tumours^4,5,6,7,8,9^. They were chosen purposively, without systematic search, one per successive biological label applied to TCGA-KIRC lncRNA models between 2019 and 2024, and are illustrative, not representative, of that literature. Each was scored as the weighted sum of log2(FPKM + 1) over its mapped genes, with the published weights and with coefficients re-estimated in TCGA-KIRC. Two signatures publish coefficients, three publish neither coefficients nor a hazard direction and were signed in sample, and one mapped too few genes to report beyond a footnote. Weight provenance is given on every table row, and ranges are never pooled across provenance classes. Each score was run alone, with the clinical terms, with the clinical terms and the three metrics, and on the adjusted matrix with the clinical terms. Two nulls of 1,000 random sets per signature size were built from one set of draws. Outcome-signing orients every random set along the cohort’s dominant prognostic axis, so the outcome-signed null^11^ is the reference for hazard ratios and a sign-randomised null that for correlation with the metric (Supplementary Table 13).

Normalisation. Raw STAR counts were re-read by file identifier and checked against recorded library sizes. Five lncRNA normalisations were compared: log2(FPKM + 1) as used, log2 counts per million by library size, log2 counts per million with trimmed-mean-of-M-values factors^59^, a blind variance-stabilising transformation^60^, and quantile-normalised log2(FPKM + 1)^61^. With module membership fixed, eigengenes were recomputed on the trimmed-mean scale, and the between-scale difference in log hazard ratio was bootstrapped over 2,000 paired resamples (Supplementary Table 2).

Alternative endpoints. Disease-specific survival and the progression-free interval came from the TCGA Clinical Data Resource, censored at ten years as for overall survival^62^. Overall survival from that source matched the pipeline endpoint on the event indicator for all 528 patients, with survival times agreeing at Spearman 0.998. The module increment was re-estimated for each endpoint by the same repeated cross-validation, with the inner fold split specified explicitly. The false discovery rate was controlled across the four endpoint analyses reported, counting overall survival twice (pipeline and Clinical Data Resource), and across the three distinct endpoints. The disease-specific analysis was repeated with both networks rebuilt inside every training fold, against a matched fixed-module arm. CPTAC-3 carries neither endpoint, so these are discovery-only sensitivity analyses (Supplementary Table 8).

Driver mutations and positional classes. Non-synonymous variants in VHL, PBRM1, BAP1, SETD2, KDM5C, MTOR and TP53, and the total non-synonymous count, were tested against the non-feature read fraction and the lncRNA axis, controlling the false discovery rate within each outcome over eight tests. Masked variant files were available for 368 of the 528 discovery patients, and availability itself is associated with the exposure, so this analysis is conditional on a biased subset. The 3,442 network lncRNAs were classified against GENCODE v36 protein-coding genes as exonic sense, antisense, intronic, sense-overlapping, divergent or intergenic. Per-gene correlation with the metric was compared across classes by Kruskal-Wallis and by sequential analysis of variance entering class after mean log2 expression and log10 gene length (Supplementary Table 1).

Specimen handling time. For each CPTAC-3 library, the times from excision and from clamping to freezing were taken from the Genomic Data Commons record of the sequenced sample and correlated with the alignment metrics, the projected lncRNA axis, T category, stage and grade by Spearman correlation, with 2,000-resample bootstrap intervals and Benjamini-Hochberg control within each sample set. Overall survival was modelled per standard deviation of the excision-to-freezing time, alone and with age, sex, T category, M1 and ordinal grade. This analysis is post hoc (Supplementary Table 22).

Sex. In TCGA-KIRC, CPTAC-3 and TCGA-KIRP, the five prognostic modules were refitted with a module-by-sex interaction and separately within women and within men. The sex-specific fits let every covariate and the baseline hazard differ by sex, so the two models estimate different quantities and are never presented in one row. Every module eigengene, the three alignment metrics and the lncRNA axis were tested for association with sex in all four cohorts. X- and Y-encoded module members were counted against the GENCODE background, and recorded sex was checked against XIST and the chromosome Y lncRNAs (Supplementary Table 15).

### Network reorganisation

The lncRNA network was rebuilt on the observed matrix of the same 511 libraries with every main-network parameter unchanged, including power, deep split, merge height and minimum module membership. Intramodular connectivity was computed on each network’s own matrix. The partitions were compared by the adjusted Rand index over all 3,442 genes, with unassigned genes as one class, and over genes assigned in both, and each adjusted module was traced into the unadjusted partition. Positional classes in the largest unadjusted module were compared with the network background by Fisher’s exact test, with Benjamini-Hochberg control across the six classes.

### Single-cell localisation of modules

Four single-cell RNA-seq cohorts of untreated primary clear-cell renal cell carcinoma, none with outcome data, were used: GSE159115^27^, the multi-region dataset of Li and colleagues^28^, GSE222703^29^ and GSE207493^30^. All four are single-cell RNA-seq of whole cells from enzymatically dissociated tissue on the 10x Genomics Chromium platform, quantified by the authors with Cell Ranger against a GRCh38 reference: GSE159115 with 3’ v2 chemistry and Cell Ranger 2.1.1, the Li dataset with 5’ chemistry and Cell Ranger 2.1.1, GSE222703 with 3’ v3.1 chemistry and Cell Ranger 4.0.0, and GSE207493 with 3’ v3 chemistry and Cell Ranger 3.1.0. Cell Ranger counts intronic reads by default from version 7 and on request from version 5, and earlier versions count them only with a pre-mRNA reference. None of the four reports using a pre-mRNA reference, so the deposited matrices should count reads in annotated exons only. Cells were assigned to malignant, normal epithelial and microenvironment compartments from the authors’ annotations. GSE207493 lacks deposited annotations and was labelled by reference transfer from GSE159115 using only protein-coding genes, so no lncRNA influenced the malignant call. Cells in its in-tumour epithelial cluster were individually reassigned as malignant if at least three of CA9, NDUFA4L2, ANGPTL4, VEGFA and EGLN3 were detected and neither PTPRC nor PECAM1 was. Counts were summed to donor-level pseudobulk^63^. Malignant cells were contrasted with normal epithelium, and with pooled tumour immune, endothelial and fibroblast or stromal cells, by edgeR quasi-likelihood^64^, paired by donor where at least three donors contributed both groups. A cohort entered a contrast only with enough donors on both sides for inference, giving three cohorts for the normal-epithelium contrast and four for the microenvironment contrast. These contrasts come from a separate, outcome-blind pipeline, run beforehand and not altered. That pipeline also supplies the restricted-maximum-likelihood random-effects meta-analysis of lncRNA fold changes across cohorts that gives the pooled per-gene estimates in the Results.

For each of the 29 modules and each cohort, the statistic was the median log2 fold change of the members tested in that cohort. A two-sided empirical p value came from 2,000 random sets of the same size, drawn from the same gene class among network genes tested in that cohort and matched on single-cell abundance by deciles of log10 mean counts per million. Cohorts were combined by an unweighted Stouffer sum of per-cohort z scores, with Benjamini-Hochberg control across the 29 modules within each contrast. The null treats genes as exchangeable although genes within a module are correlated, so combined z scores overstate precision and serve only to rank modules. Inference rests on per-cohort empirical p values and cross-cohort agreement.

### Pan-cancer analysis

Open-access STAR-Counts files for every TCGA project were retrieved from Genomic Data Commons Data Release 46.0 and verified against their md5 checksums. Primary tumours (peripheral-blood cancer for TCGA-LAML) and solid-tissue normals were processed with the discovery rules: the library-failure rule, one library per patient per group, the discovery gene filters applied within each project and group, and the discovery definition of the leading component. Each project therefore uses its own expressed genes, so its own-component values differ from those computed on the discovery genes (Supplementary Table 4). TCGA-KIRC run through this pipeline is the like-for-like check. The discovery classifier assigned a positional class to every GENCODE v36 lncRNA and reproduced the discovery class of all 3,442 network lncRNAs. The lncRNA network was built twice per eligible project, on observed and residualised expression, with the discovery lncRNA parameters and sample-outlier rule. Overall survival came from the TCGA Clinical Data Resource, censored at ten years. Stage was the AJCC pathologic stage where at least 70% of analysed patients had it, otherwise the clinical stage, and was otherwise omitted. Project estimates were combined by restricted-maximum-likelihood random-effects meta-analysis^65^.

Decision rules were written into the pipeline configuration, committed to version control and hashed before any non-renal pan-cancer project was analysed. We had analysed the renal projects during discovery, so we set the thresholds with those results known. Pipeline stages 39 to 41 verify the rules before running, and the rule choosing the source of tumour stage for survival models is fixed in the analysis script. A project was eligible with at least 100 retained primary tumours, and TCGA-LAML, a blood cancer, was excluded from inference. Rule A, for a lncRNA-specific axis, required the lncRNA component to track the non-feature fraction at an absolute Spearman rho of at least 0.5 and to exceed the protein-coding component by at least 0.3. Rule C, for a reorganised network, required a largest observed module holding at least 40% of genes, an eigengene correlating with the metric at an absolute Pearson r of at least 0.7, and an adjusted Rand index below 0.2. The letters correspond to the three measurements of the analysis plan. Measurement B, the gene-level and positional-class comparison, had no decision threshold and is reported descriptively. Counts are reported over all eligible projects and over the non-renal projects alone. Normal-tissue sets with at least 20 libraries were analysed descriptively, and projects with at least 20 deaths entered the meta-analysis. The dose-response analysis, specified after these rules were applied, correlated four measures of the metric’s within-cohort spread with the rule quantities and the plate share of the metric’s variance across projects, with Benjamini-Hochberg control across its 20 tests.

### Single-cell test of the correction

The test used the TCGA-KIRC primary tumours and solid-tissue normals of the pan-cancer build (532 tumours), 4,088 lncRNAs expressed in either group, and the single-cell results of the outcome-blind pipeline, whose candidate lncRNAs were locked before any bulk step. The pan-cancer build retains the 4 patients excluded from discovery for survival time. The correction was the one used throughout, fitted in the tumours and applied unchanged to the normals. A sensitivity arm fitted it on both groups with a tumour indicator and removed only the metric terms. Agreement was the Spearman correlation across lncRNAs between the random-effects single-cell fold change of malignant against normal epithelial cells and the mean paired bulk tumour-minus-normal difference, together with sign agreement among lncRNAs at a single-cell false discovery rate below 0.05. The change in agreement was assessed by 2,000 gene bootstraps. Agreement was computed for the paired contrast and for the unpaired contrast of all tumours against all normals, which was added post hoc because the paired contrast proved insensitive. Scores were means of per-gene z scores. The malignant-marker score used CA9, NDUFA4L2, ANGPTL4, VEGFA and EGLN3, the markers the single-cell pipeline used to check its malignant labels. The null was 1,000 sets of lncRNAs the single-cell analysis calls non-specific (absolute log2 fold change below 0.25, false discovery rate above 0.5), matched to the tumour-specific set on bulk abundance decile.

### Comparison with RUV-III

RUV-III^16^ was applied with the ruv package to log2(FPKM + 1) of the 12,000 discovery protein-coding genes and the expressed lncRNAs of the 528 TCGA-KIRC tumours with a ccA/ccB class, following the pseudo-replicate design of RUVprps^66^. Biology groups were ccA/ccB class crossed with stage I-II or III-IV. Pseudo-samples were means of at least three samples of one group on one plate, and of the ten samples with the lowest and the ten with the highest non-feature fraction or depth within a group, giving 12 replicate sets of 66 pseudo-samples. Negative controls were the 1,000 genes most associated with the metric and depth among the half least associated with the biology groups. Three unwanted factors were removed, with one and five as sensitivity analyses. The single-cell coherence test was repeated on these 528 tumours with 500 null sets, so its excess values differ slightly from those of the single-cell test of the correction. This comparison is a post hoc sensitivity analysis.

### Statistics and reproducibility

All tests were two-sided with alpha 0.05. The false discovery rate was controlled by the Benjamini-Hochberg procedure at 5% within the stated families, and every reported false discovery rate names its family. Hazard ratios are from Cox proportional-hazards models, per standard deviation of the eigengene or metric, with 95% confidence intervals. Correlations are Spearman unless stated otherwise. Confidence intervals on a concordance index are the estimate plus or minus 1.96 standard errors. The index depends on the cohort’s censoring distribution, so values are not comparable between cohorts and the within-cohort increment is the quantity interpreted. Bootstraps used 2,000 patient-level paired resamples (500 for the decision-curve medians), with the p value as twice the smaller tail proportion. Every model uses the complete cases of its own covariates, and its sample size and event count are given where it first appears. Randomisation and blinding do not apply to an analysis of existing cohorts. The global seed was 20260816, reset at the start of every stochastic stage, and the five fold assignments of the rebuild analyses used seeds 20260817 to 20260821.

Analyses used R 4.4.2 on Windows with the numeric locale set to C. The principal packages were WGCNA 1.73, survival 3.7-0, glmnet 4.1-10, data.table 1.16.4, clusterProfiler 4.12.6, estimate 1.0.13, edgeR 4.2.2, DESeq2 1.44.0, limma 3.60.6, GenomicRanges 1.56.2, metafor 4.8.0 and ruv 0.9.7.2. The full list with versions is in the repository. The study was not registered and had no separate protocol, so analyses are described as principal or sensitivity analyses, not as pre-specified. The network parameters, the seed and every analysis setting are fixed in the pipeline configuration file. For the single-cell test of the correction and the handling-time analysis, the expected direction of every test was recorded in the analysis script before its first run. Reporting follows the TRIPOD+AI statement^67^.

## Data availability

No new data were generated for this study. All primary data analysed are publicly available. Expression data are the augmented_star_gene_counts.tsv STAR-Counts files of the National Cancer Institute Genomic Data Commons (https://portal.gdc.cancer.gov), Data Release 46.0 of 10 August 2026, for projects TCGA-KIRC, TCGA-KIRP, TCGA-KICH and CPTAC-3. Harmonised clinical fields, sample types and file manifests were retrieved from the Genomic Data Commons application programming interface at the same release. Tumour grade for TCGA-KIRC was taken from the Biospecimen Core Resource clinical XML supplements. The biospecimen measurements, which the Genomic Data Commons biospecimen records carry for TCGA-KIRC only, were the RNA integrity number, the A260/A280 ratio, the slide percentages of tumour nuclei, stroma, normal tissue and necrosis, the sequencing plate and the Biospecimen Core Resource batch. Somatic variant calls are the open-access masked MAF files of the same release. Disease-specific survival and the progression-free interval are from the TCGA Clinical Data Resource^62^. Gene annotation is GENCODE v36^49^, the reference of the Genomic Data Commons STAR-Counts workflow. Single-cell data are the Gene Expression Omnibus series GSE159115, GSE222703 and GSE207493 and the dataset of Li and colleagues at Mendeley Data (g67bkbnhhg), all open access. The pan-cancer analysis used the STAR-Counts files of all 33 TCGA projects at the same release. Supplementary Data 5 lists every file with its identifier, md5 checksum and retention status.

The expression, clinical and mutation files are open access and require no authorisation. The parent studies are registered in dbGaP under accessions phs000178 (The Cancer Genome Atlas) and phs001287 (Clinical Proteomic Tumor Analysis Consortium). These accessions govern the controlled-access data of the same programmes, which were not used.

Supplementary Data 1 lists all 1,549 STAR-Counts expression files of the four projects, one row per file, with the Genomic Data Commons file and case identifiers, workflow version, data release, md5 checksum, sample type, and whether the file was retained or why it was excluded. Local copies of 1,491 files matched the Genomic Data Commons md5 checksums. The remaining 58 files, 32 TCGA-KIRP and 25 TCGA-KICH normal-tissue libraries and one TCGA-KIRP additional-new-primary sample, were never downloaded or analysed. Supplementary Data 2 assigns the 10,407 genes of the two networks to all 29 modules and gives per-gene module membership, eigengene loading, and the centring and scaling constants that define each module eigengene. Supplementary Data 3 gives the survival results for all 29 modules under the principal specification, and Supplementary Data 4 the Gene Ontology enrichment of the protein-coding modules. Source data for every figure panel are the result tables in the code repository. The repository README lists them by figure and table, and each supplementary figure legend names the table behind its panels.

## Code availability

The complete analysis pipeline is available at https://github.com/GehadYoussef/rcc-lncrna-networks under an MIT licence. The version used for this article (release v1.1.1, commit c8abeb5) is archived on Zenodo at https://doi.org/10.5281/zenodo.23255737.

The pipeline is written in R. The bulk pipeline in src/R comprises 49 files, namely the configuration file 00_config.R, 44 numbered stage scripts, the enrichment helper ora_hypergeometric.R, the pan-cancer helper pan_helpers.R, the download script 00_download_gdc.R and the driver run_all.R. The configuration file holds every analysis constant, threshold and random seed and the locked pan-cancer decision rules. The driver runs the stages in order and caches each stage’s output. The single-cell pipeline that produces the localisation contrasts is in src/singlecell, with its settings in src/config/singlecell.yml. The repository README lists the stage scripts and names the result tables behind every figure and table. The main figures are built by 13_figures.R, the supplementary figures by 32_supplementary_figures.R and the manuscript tables by 14_tables.R, all directly from the result tables, so that no table or figure can drift from the number in the text.

Every result table cited in the paper is in the results folder of the repository. Package versions are listed in the Methods (Statistics and reproducibility) and in results/31_package_versions.tsv, and each stage writes its session information when run.

## Acknowledgements

The results published here are in whole or part based upon data generated by the TCGA Research Network (https://www.cancer.gov/tcga) and by the National Cancer Institute Clinical Proteomic Tumor Analysis Consortium (CPTAC). The authors thank the patients and investigators who contributed to those programmes. G.Y. was funded by the Milner Therapeutics Institute and Cancer Research UK. N.H. was supported by the National Research Foundation of Korea grant funded by the Ministry of Science and ICT (RS-2025-18362970) and by the Brain Pool Plus Fellowship Program funded by the Ministry of Science and ICT (RS-2025-25427881). The funders played no role in study design, data collection, analysis and interpretation of data, or the writing of this manuscript.

## Author contributions

G.Y. conceived the study, designed the analysis, wrote the pipeline, assembled the cohorts, performed the statistical analysis, produced the figures and tables, and wrote the first draft. N.H. supervised the work, obtained the funding, administered the project, and reviewed and edited the manuscript. Both authors interpreted the results and read and approved the submitted manuscript.

