## Supplementary Figures for "Non-exonic read content reorganises lncRNA co-expression networks in renal and other cancers"

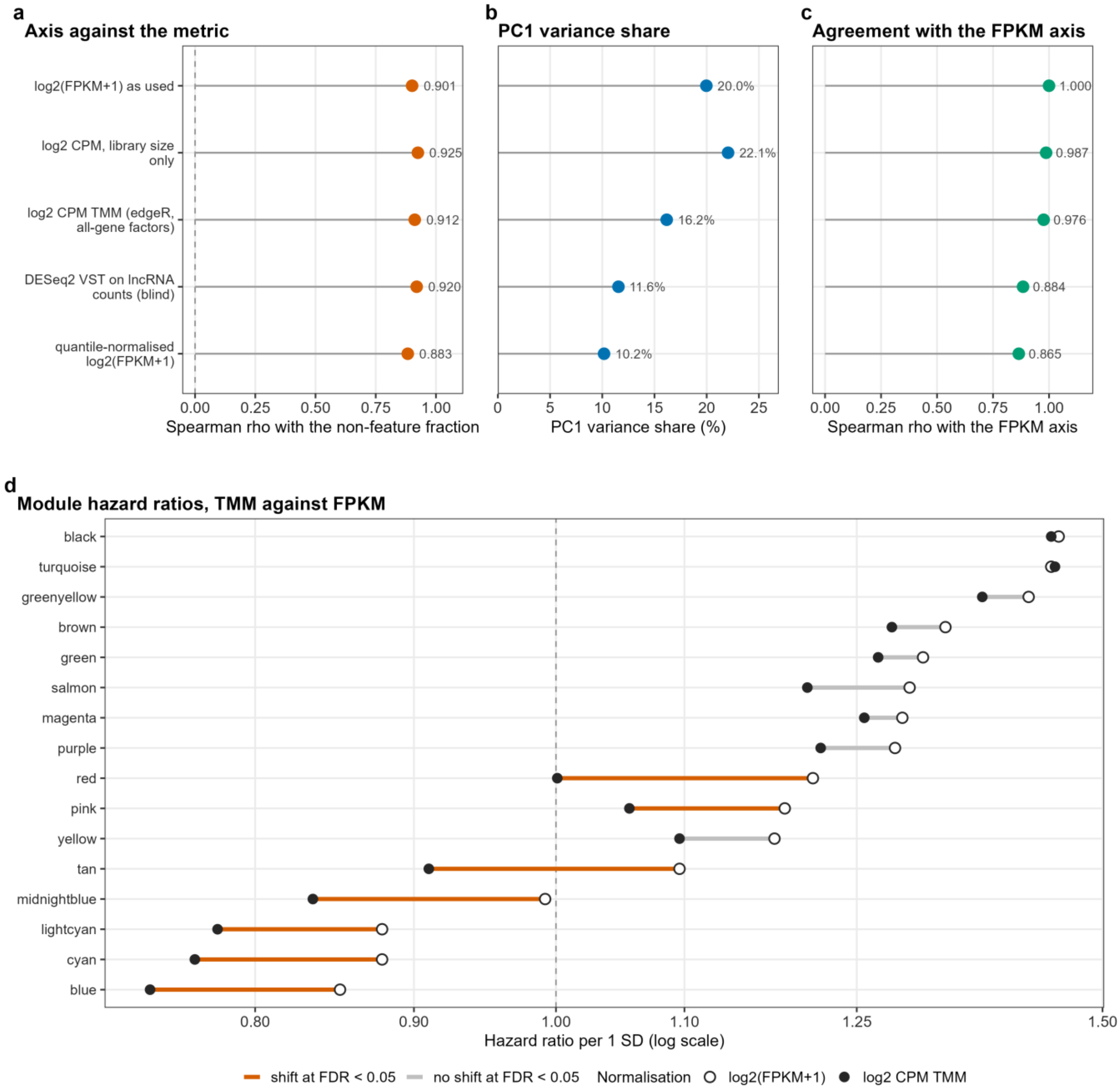


**Supplementary Fig. 1. The lncRNA axis under five normalisations.** **a** Spearman correlation of the leading lncRNA principal component with the non-feature fraction under five normalisations of the same 3,442 genes and 511 libraries. The five arms are log2(FPKM + 1) as used throughout, log2 counts per million with library size only or with edgeR TMM factors, a blind DESeq2 variance-stabilising transform, and quantile-normalised log2(FPKM + 1). **b** PC1 variance share. Each PC1 is oriented against the log2(FPKM + 1) axis. **c** Spearman correlation of each arm's PC1 with the FPKM axis. Quantile normalisation removes the per-sample mean, leaving that arm's sign anchor indeterminate, so it is quoted as a magnitude. **d** Paired patient bootstrap difference in log hazard ratio between the TMM and FPKM scales for the 16 lncRNA modules (2,000 resamples, 504 patients, 163 deaths), with the same modules, patients and covariates on both scales. With Benjamini-Hochberg control within each arm, 7 of 16 differences reach a false discovery rate below 0.05 on the observed matrix and 0 of 16 on the residualised matrix. Before multiplicity control, 9 and 2 intervals exclude zero. Source tables: `29_normalisation_pc1.tsv`, `29_normalisation_module_contrast.tsv`.

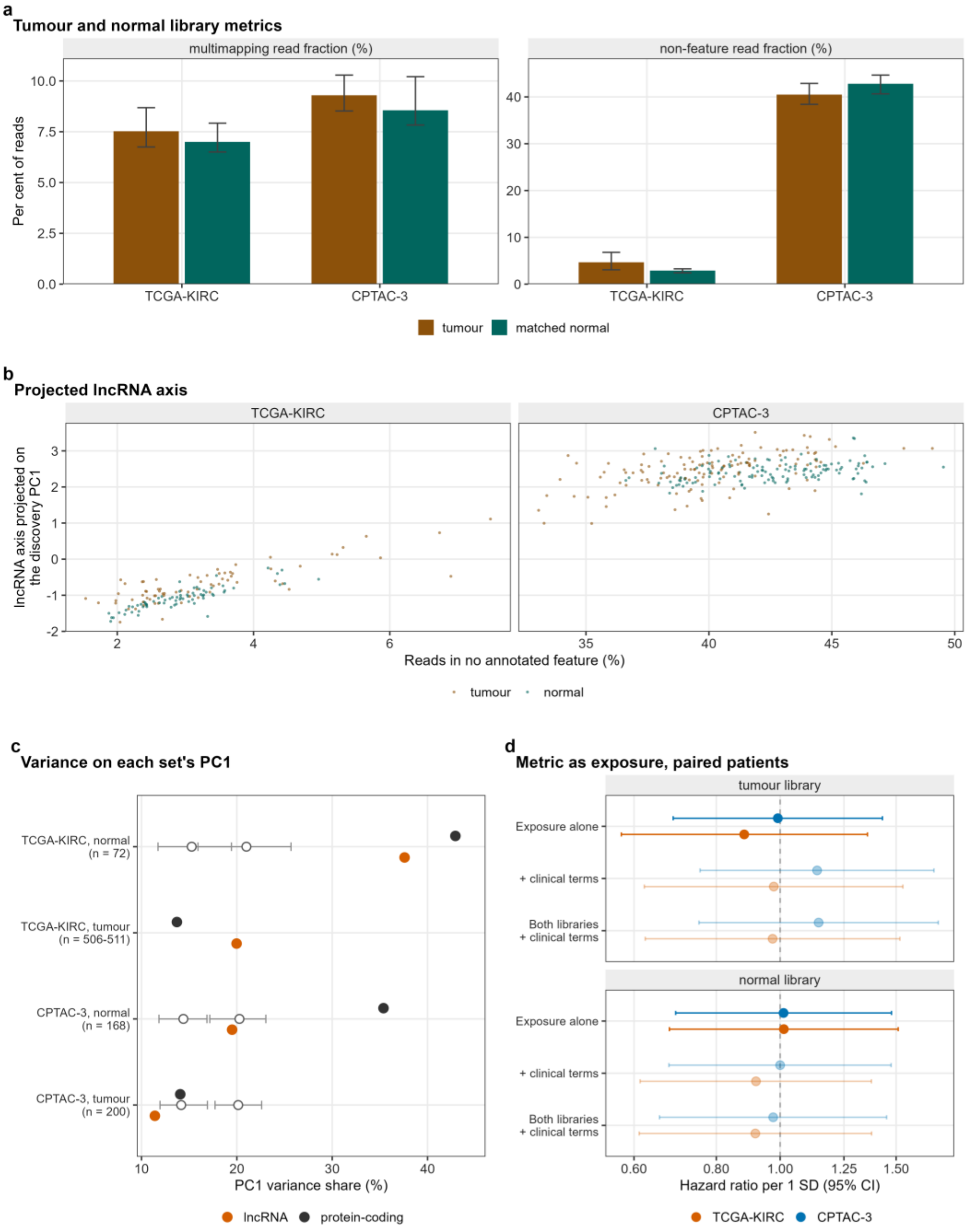


**Supplementary Fig. 2. Matched normal-kidney libraries. a** Median and interquartile range of the multimapping and non-feature fractions in tumour and matched normal libraries of TCGA-KIRC (528 tumour, 72 normal) and CPTAC-3 (200 tumour, 168 normal). All four unpaired rank-sum tests reach a false discovery rate below 0.05, but the unpaired contrast mixes tissue with plate. In TCGA-KIRC the non-feature difference falls from 1.73 percentage points unpaired to 0.16 on the 68 plate-matched normal libraries (p 0.049), to 0.15 for the normals of the analysed tumours on shared plates (144 tumour and 67 normal libraries, p 0.057), and to 0.15 within patient (71 pairs, signed-rank p 0.10). **b** Projected lncRNA axis (the discovery lncRNA PC1 direction applied to each sample set) against the non-feature fraction in the paired libraries, faceted by cohort and coloured by tissue type: rho 0.77 in TCGA-KIRC tumours and 0.86 in their normals (71 pairs), and 0.54 and 0.35 in CPTAC-3 (129 pairs). **c** PC1 variance share of each sample set's own lncRNA and protein-coding matrix against a size-matched discovery baseline from 200 resampling draws (open grey points and bars, the median and range of the draws, available for the six non-discovery rows). The baseline range is not a confidence interval. Three of the eight observed shares lie above it. A share carries no sign, and the sign of three of the eight components cannot be fixed. **d** Forest plot of the non-feature fraction (linear scale, per standard deviation), measured on the tumour or the normal library, as exposure for overall survival in the paired patients under three models. Faded rows have an events-per-variable ratio below 5, as does every adjusted model in the panel (71 patients and 26 events in TCGA-KIRC, 129 and 28 in CPTAC-3). Source tables: `25_tumour_vs_normal_distribution.tsv`, `25_paired_scores_per_sample.tsv`, `25_normal_axis_projection.tsv`, `25_normal_metric_survival.tsv`.

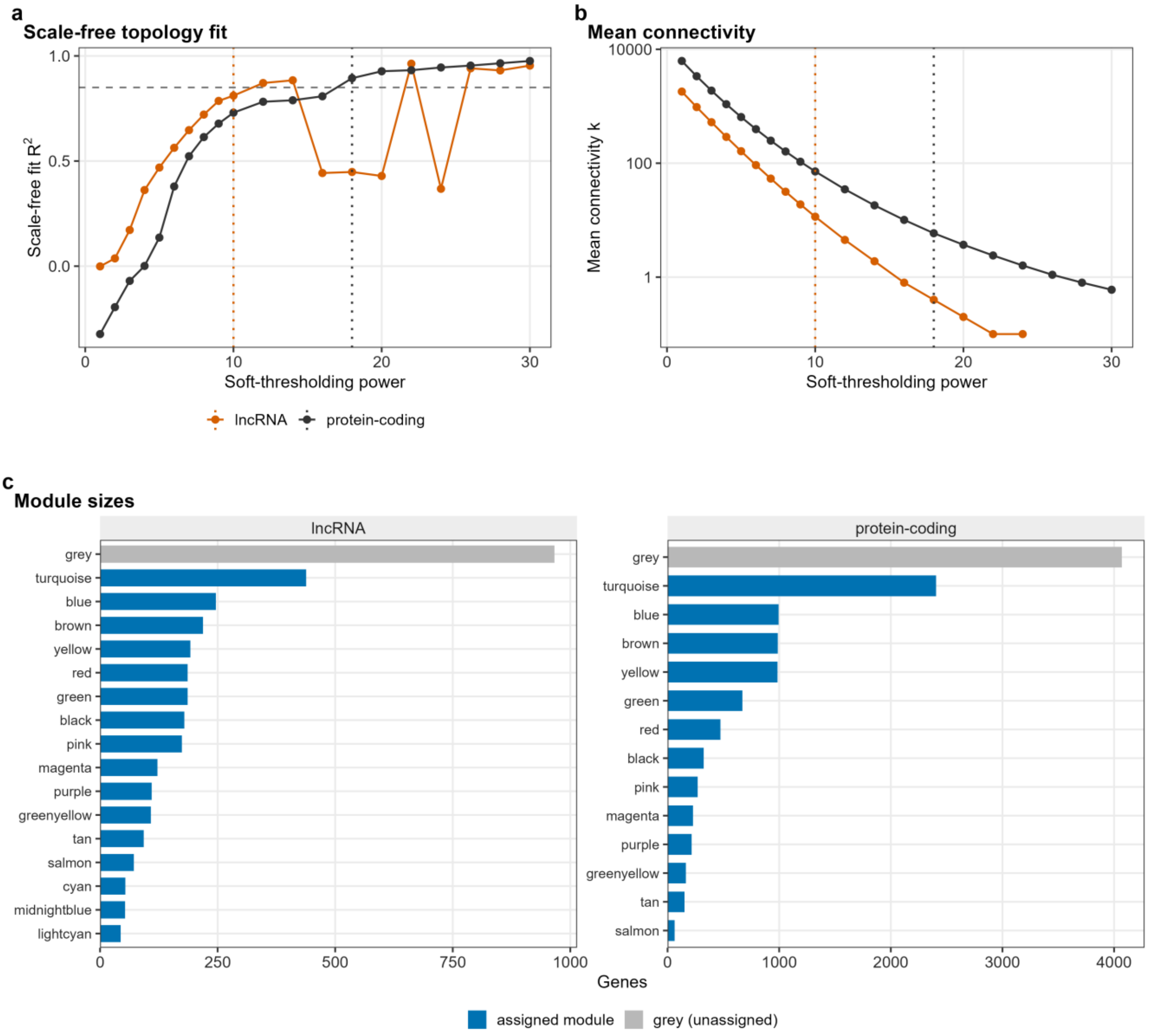


**Supplementary Fig. 3. Network construction diagnostics. a** Signed scale-free topology fit R-squared against soft-thresholding power for the protein-coding (12,000 genes, 506 samples) and lncRNA (3,442 genes, 511 samples) networks. The horizontal line marks the target of 0.85 and the vertical lines the chosen powers, 18 and 10, at which the signed fit is 0.894 and 0.811. The protein-coding power is the lowest reaching the target. The outcome-blind parameter sweep (Methods) fixed the lncRNA power at 10, where the automatic rule would have given 12. **b** Mean connectivity against power on a logarithmic axis, with the same chosen-power lines. **c** Module sizes, with unassigned (grey) genes shown separately. The protein-coding network resolves 13 non-grey modules, leaving 4,069 of 12,000 genes unassigned, and the lncRNA network 16, leaving 966 of 3,442. Source tables: `02_lncRNA_soft_threshold.tsv`, `02_mRNA_soft_threshold.tsv`, `02_network_summary.tsv`, `02_lncRNA_module_sizes.tsv`, `02_mRNA_module_sizes.tsv`.

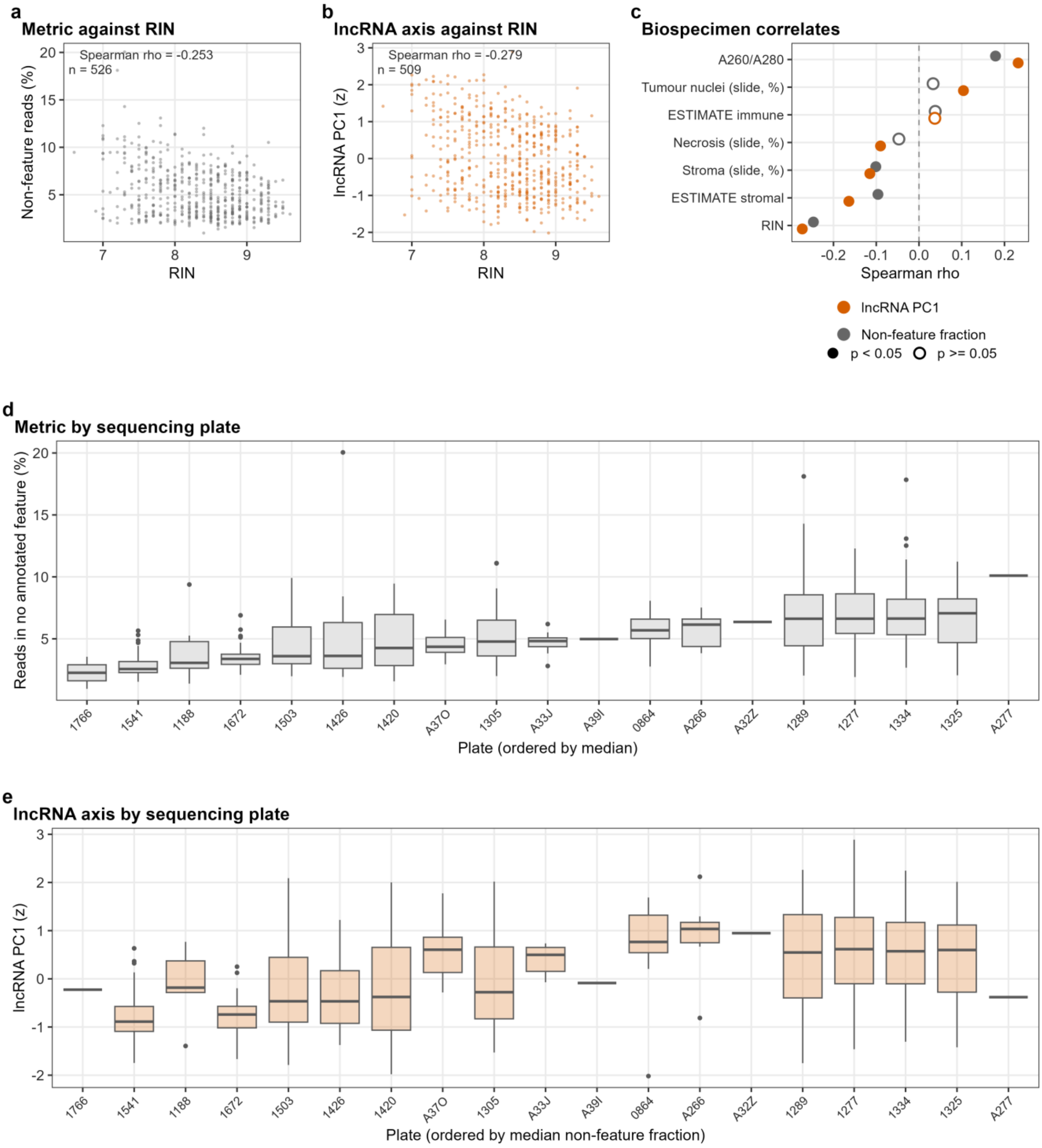


**Supplementary Fig. 4. RNA integrity number and sequencing plate. a** Non-feature read fraction against RNA integrity number in TCGA-KIRC, with the integrity number from the Biospecimen Core Resource biospecimen record. One library with an RNA integrity number recorded as 0, 6.6 units below the next lowest, is treated as a coded missing value and excluded from panels **a** and **b**, so their sample size is one below the correlation table's. **b** Observed lncRNA PC1, the axis of Fig. 2a, against RNA integrity number, with the same exclusion. **c** Spearman correlation of the non-feature fraction and the lncRNA axis with seven biospecimen measures: RNA integrity number, A260/A280 ratio, slide percent necrosis, tumour nuclei and stromal cells, and the ESTIMATE stromal and immune scores (507 to 528 patients). Filled symbols mark the 10 of 14 tests with unadjusted p below 0.05. **d** Non-feature fraction by sequencing plate, ordered by median, for 528 libraries on 19 plates (1 to 59 per plate). **e** lncRNA axis by plate, in the same order, for the 511 libraries with an axis score. Biospecimen and batch metadata exist only for TCGA-KIRC. Source tables: `23_biospecimen_kirc.tsv`, `23_axis_vs_biospecimen.tsv`, `27_axis_scores_per_sample.tsv`.

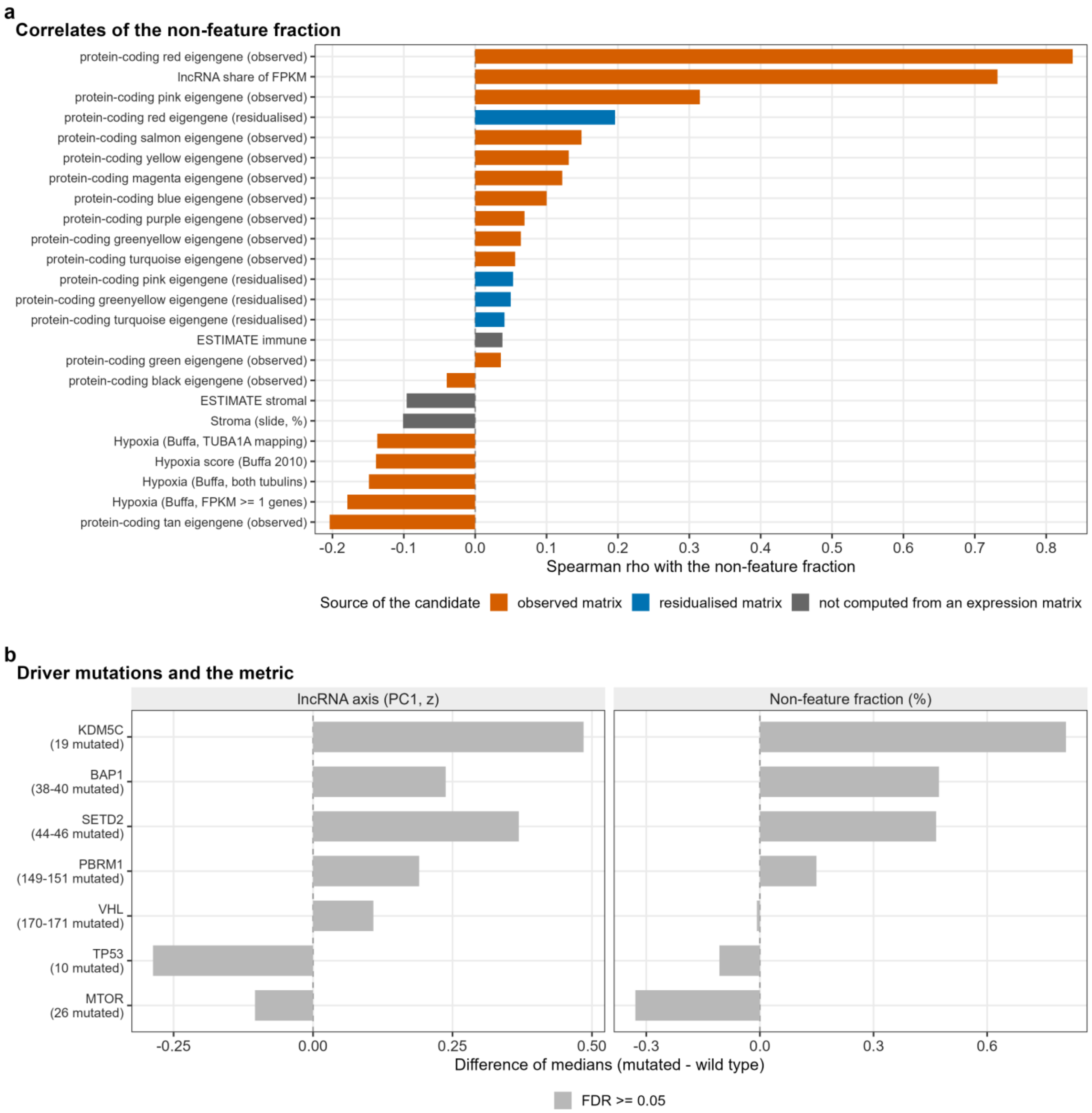


**Supplementary Fig. 5. What the non-feature read fraction correlates with. a** The 24 strongest candidate correlates of the non-feature fraction in TCGA-KIRC (506 to 528 patients), coloured as eigengenes of the observed or the residualised expression matrix, or quantities computed outside it. Observed eigengenes cannot be read as biology, because the metric was measured on the same matrix. The annotated biological candidates are the hypoxia metagene, the proliferation score, the ESTIMATE stromal and immune scores and the slide tumour-nuclei percentage. **b** Difference of medians in the non-feature fraction and the lncRNA axis between mutated and wild-type patients for seven recurrent clear-cell drivers. None of the 14 tests (7 genes by 2 outcomes) reaches a 5% false discovery rate, on 358 to 368 patients per test and as few as 10 mutated patients. Testing is conditional on the 368 of 528 patients with an open-access masked mutation file, whose median non-feature fraction is lower than that of the others. Source tables: `27_metric_biology_correlations.tsv`, `28_noFeature_vs_mutations.tsv`.

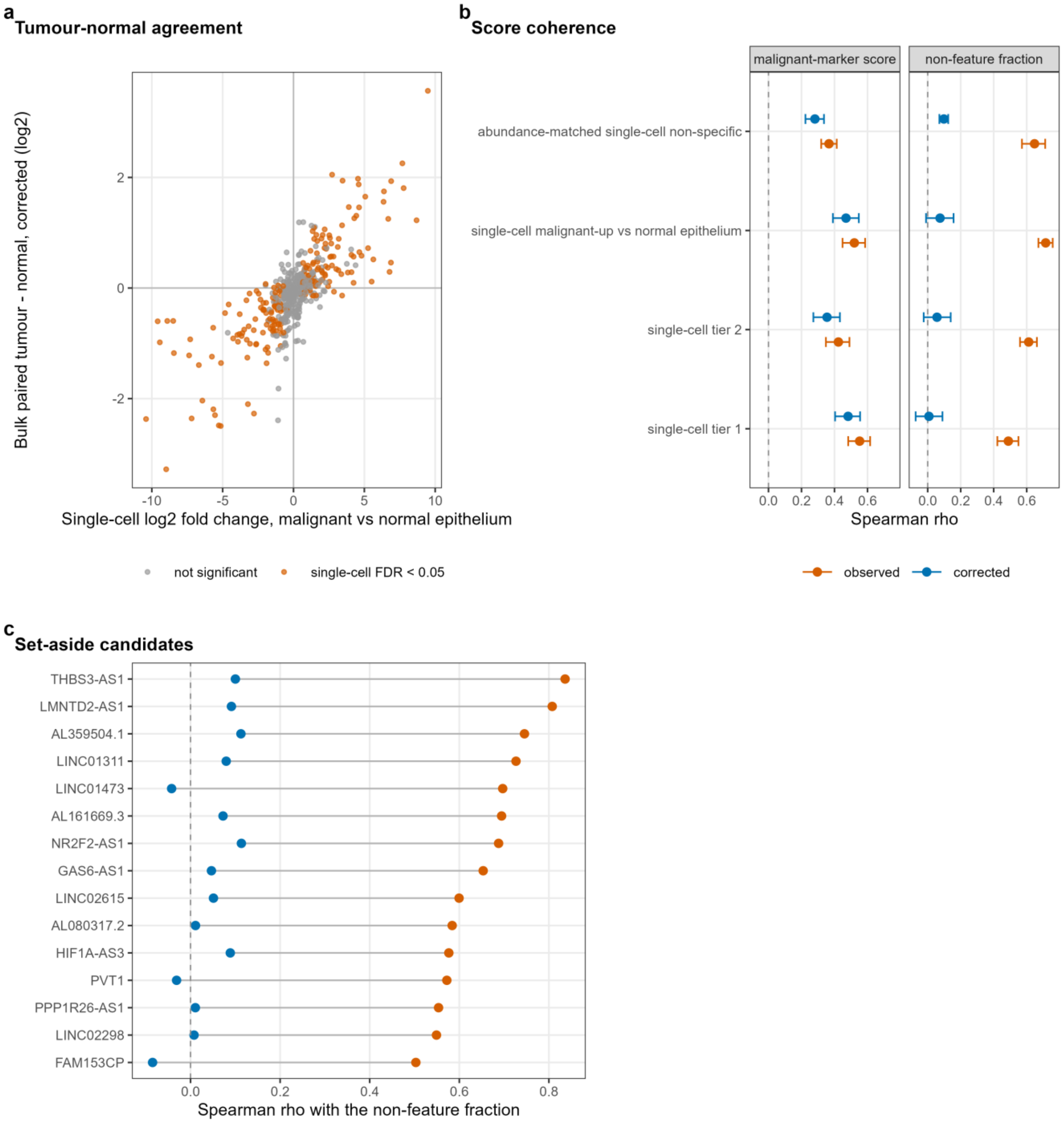


**Supplementary Fig. 6. Single-cell ground truth for the correction. a** Corrected bulk paired tumour-minus-normal difference against the single-cell fold change of malignant against normal epithelial cells, for 507 lncRNAs in 72 TCGA-KIRC patients. Orange, single-cell false discovery rate below 0.05. Agreement is Spearman rho 0.775 observed and 0.771 corrected, with sign agreement of 92.5% and 92.0% among single-cell-significant lncRNAs. **b** Spearman correlation of lncRNA scores in 532 tumours with the malignant-marker score and the non-feature fraction, observed (orange) and corrected (blue), with 95% bootstrap intervals. The marker score itself correlates with the metric at 0.22. The null row is 1,000 abundance-matched sets of single-cell non-specific lncRNAs. Correlation with the metric falls by construction, and the informative quantity is marker coherence relative to the null. **c** Correlation with the non-feature fraction, before and after correction, of the 15 single-cell candidates set aside in bulk as technically sensitive. Source tables: `42_sc_truth_concordance.tsv`, `42_sc_truth_score_coherence.tsv`, `42_sc_truth_technical_candidates.tsv`.

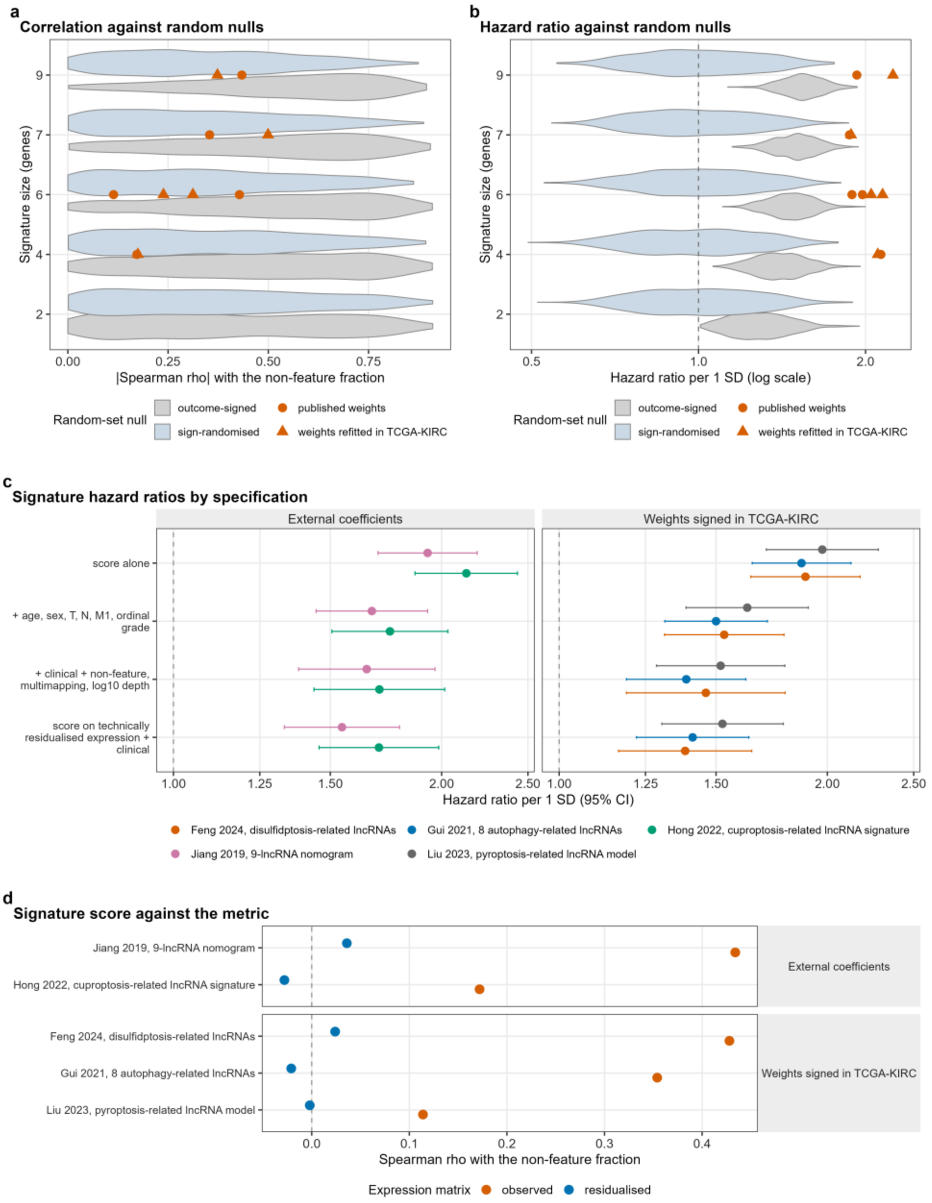


**Supplementary Fig. 7. Published lncRNA signatures against two random-signature nulls. a** Absolute Spearman correlation with the non-feature fraction (violins) for 1,000 random gene sets at each of five set sizes, under two weightings of the same draws: outcome-signed after Venet and colleagues^11^, and sign-randomised. The two nulls are the same 5,000 gene sets signed two ways. Points are the five primary-tier signatures, each with its published weights and with weights refitted in TCGA-KIRC. The Yu signature is held back for incomplete mapping. Against the sign-randomised null the signatures sit at percentiles 25.4 to 65.3 for the two with external coefficients and 19.1 to 64.9 for the three signed in sample, and against the outcome-signed null at 15.9 to 30.9 and 8.6 to 36.7. **b** Unadjusted hazard ratio per standard deviation (violins) for the same draws and weightings in 528 patients with 173 deaths, with the signatures overlaid. The sign-randomised null has a median hazard ratio of 1.00, whereas 95% of outcome-signed sets reach p below 0.05 by construction. The signatures sit at percentiles 99.8 to 100.0 (external coefficients) and 99.7 to 99.9 (signed in sample) of the outcome-signed null. **c** Hazard ratio per standard deviation (95% confidence interval) for the five primary-tier signatures under four specifications, grouped by weight provenance (520 to 528 patients, 172 to 173 deaths). All 20 intervals exclude 1. Adding the three STAR metrics changes the clinically adjusted log hazard ratio by -4.9% to -2.6% with external coefficients and -19.0% to -11.1% with weights signed in sample. Residualising the expression instead changes it by -15.0% to -5.1% and -23.6% to -13.2%. **d** Spearman correlation of each signature score with the non-feature fraction (528 patients), observed against residualised expression, by weight provenance: 0.172 to 0.434 observed with external coefficients and 0.114 to 0.428 signed in sample, and below 0.04 in absolute value for every signature after residualisation. Outcome-signing itself induces correlation with the metric in the null, so outcome-signed percentiles cannot show a signature to be less correlated with the metric than chance. Source tables: `26_random_signature_null.tsv`, `26_published_signatures_models.tsv`, `26_published_signatures_correlations.tsv`, `26_headline_ranges_by_provenance.tsv`.

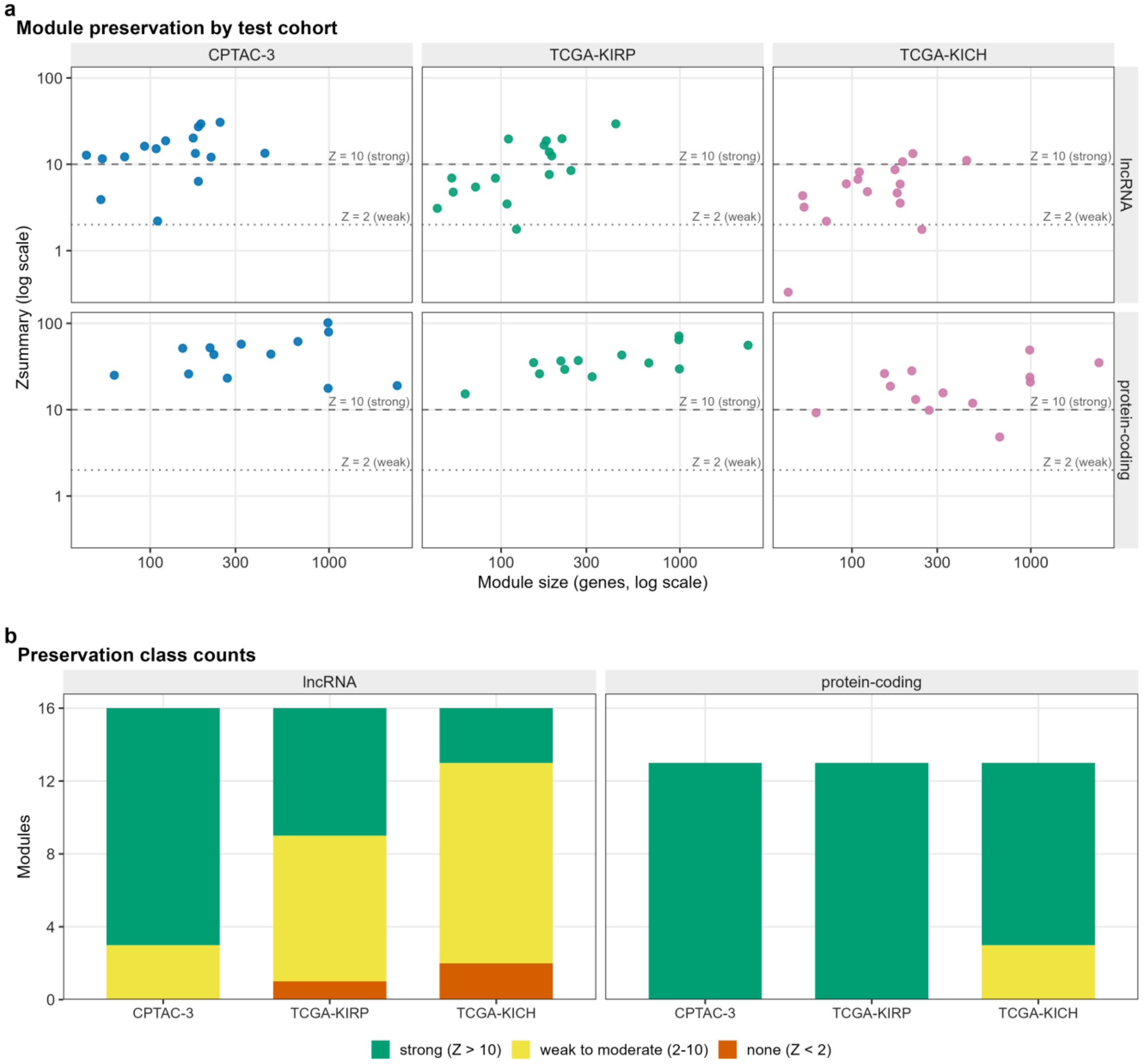


**Supplementary Fig. 8. Module preservation of the discovery networks in the three external cohorts. a** WGCNA Zsummary against discovery module size by network and test cohort, for 29 modules and 87 module-cohort pairs, with lines at the conventional bands of Zsummary 2 and 10. Module size is the true discovery size, because the preservation function caps reported size at 1,000, which affects one module. The reference is the residualised TCGA-KIRC matrix used to define the modules, and each test network is residualised on its own cohort's STAR metrics. The lncRNA network used 100 permutations, the protein-coding network 30. **b** Stacked counts of modules strongly, weakly to moderately or not preserved, per cohort and network. Strongly preserved lncRNA modules number 13 of 16 in CPTAC-3, 7 of 16 in TCGA-KIRP and 3 of 16 in TCGA-KICH, and protein-coding modules 13 of 13, 13 of 13 and 10 of 13. The gold and grey rows of the underlying tables are WGCNA internal reference sets, not modules. Source tables: `24_module_preservation_lncRNA.tsv`, `24_module_preservation_mRNA.tsv`, `24_module_preservation_summary.tsv`, `02_lncRNA_module_sizes.tsv`, `02_mRNA_module_sizes.tsv`.

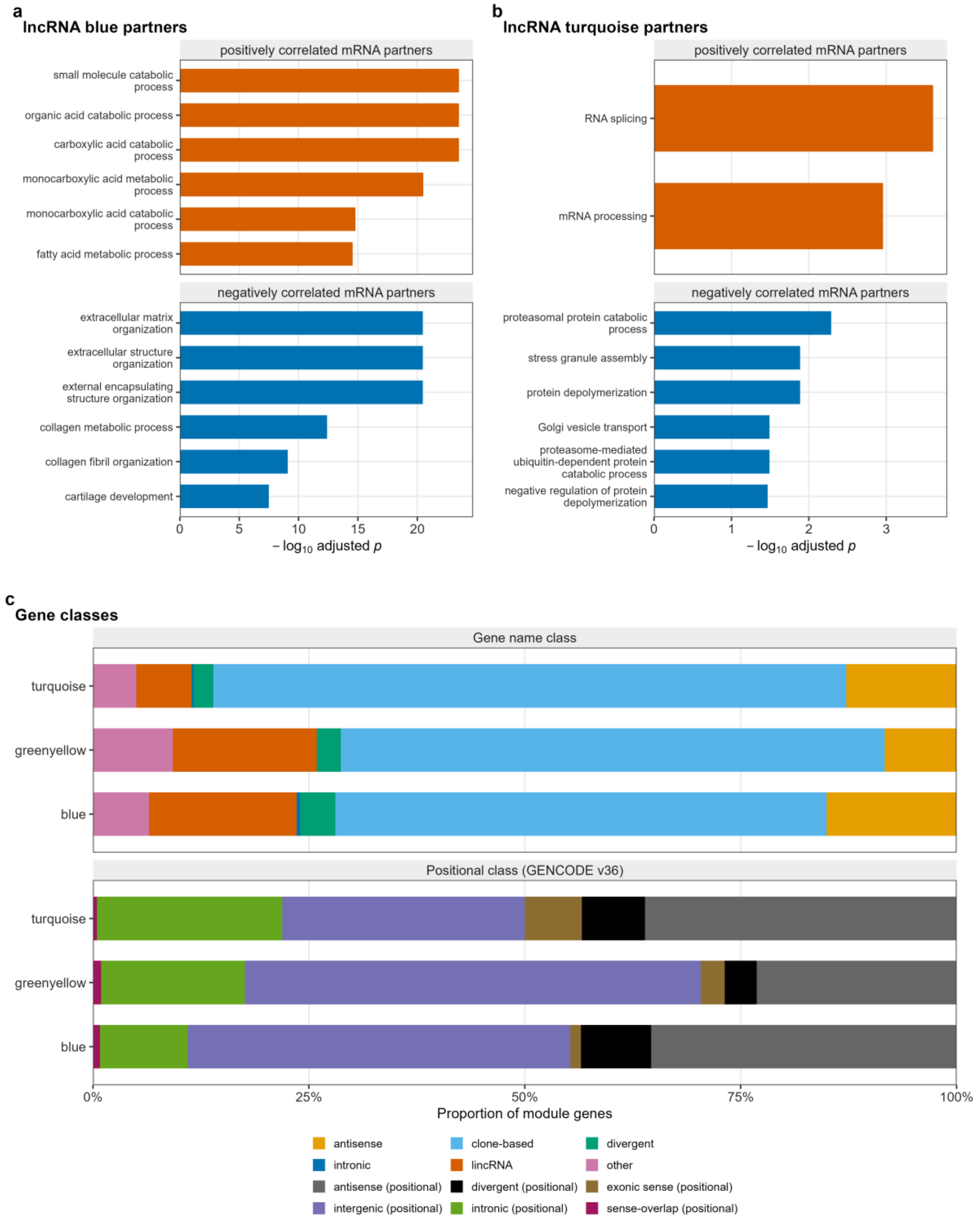


**Supplementary Fig. 9. Gene Ontology terms of correlated protein-coding partners and gene classes of the prognostic lncRNA modules. a**, **b** Gene Ontology biological-process terms of the protein-coding partners of the lncRNA blue (**a**) and turquoise (**b**) modules, drawn as in Fig. 6c for greenyellow. Partners are split by the sign of their correlation with the module eigengene, the 300 most strongly correlated of each sign are tested separately against the 12,000 network protein-coding genes, and up to six terms with the smallest Benjamini-Hochberg adjusted p value are shown per sign. Turquoise has two enriched terms among its positive partners. Because partners are selected by correlation, the adjusted p values rank terms and are not calibrated tests. Under the principal specification the hazard ratio per standard deviation is 0.71 (0.60 to 0.83, false discovery rate 4.8e-04) for blue and 1.24 (1.06 to 1.45, 0.039) for turquoise. **c** Gene-class composition, as stacked proportions, of the three prognostic lncRNA modules, which hold 246 genes (blue), 108 (greenyellow) and 438 (turquoise), under two annotation schemes. The name-based classes are antisense, clone-based unnamed, divergent, intronic, named long intergenic and other. The positional classes, relative to GENCODE v36 protein-coding genes, are exonic sense, antisense, intronic, sense-overlapping, divergent and intergenic. Source tables: `10_lncRNA_guilt_by_association_GO_signed.tsv` (**a**, **b**), `10_lncRNA_module_composition.tsv` and `28_module_composition_by_class.tsv` (**c**).

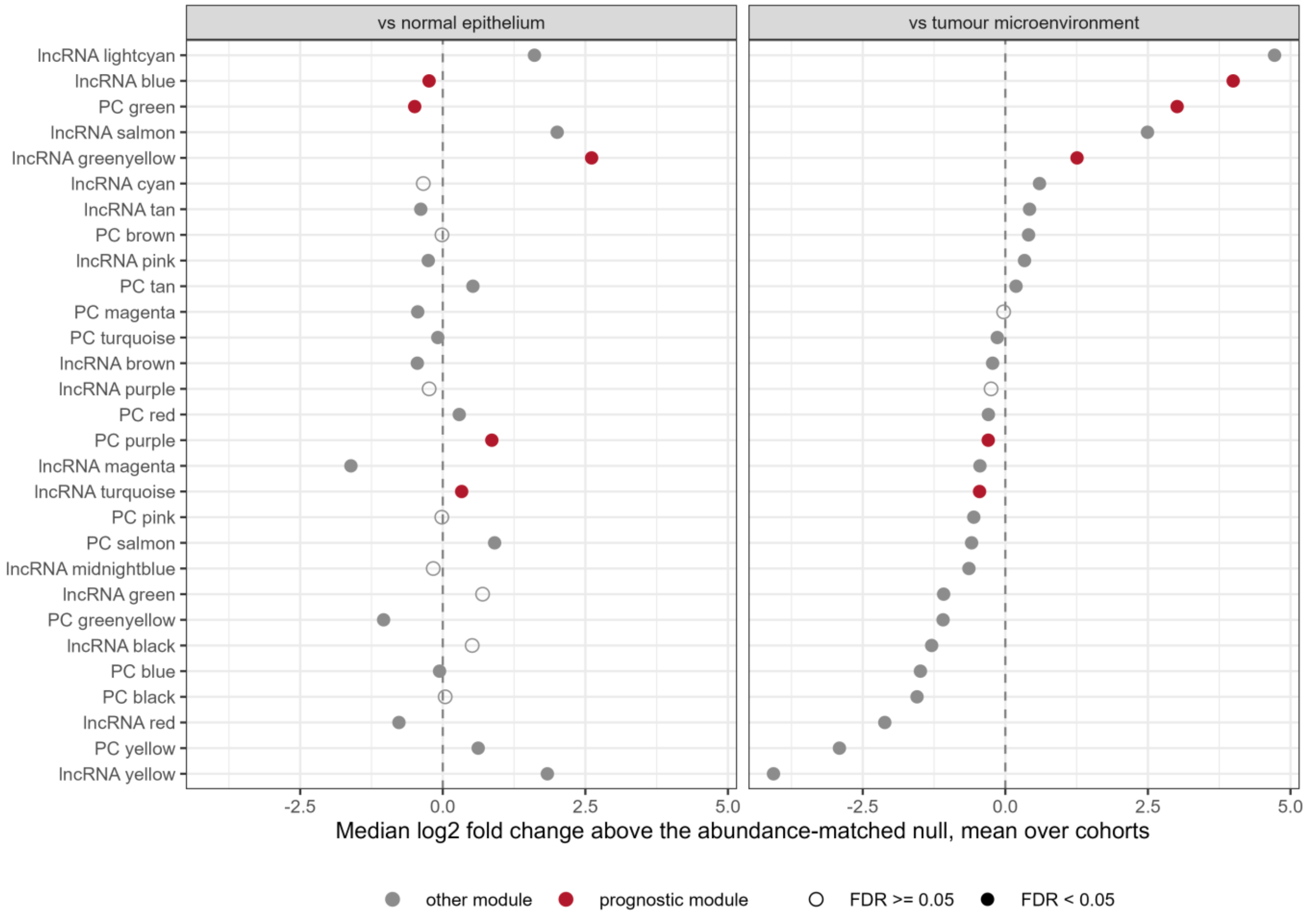


**Supplementary Fig. 10. Single-cell localisation of the 29 discovery modules.** For each module, the median log2 fold change of its members minus the mean of 2,000 abundance-matched random gene sets, averaged over cohorts, for malignant cells against normal kidney epithelium (three cohorts, left) and against the rest of the tumour microenvironment (four cohorts, right). Positive values mean higher in malignant cells than a matched random set. Red points, the five prognostic modules. Filled symbols, a within-contrast false discovery rate below 0.05 by Stouffer combination of per-cohort z scores, which overstates precision because genes within a module are correlated. Per-cohort values are in Supplementary Table 19. Source table: `36_sc_module_localisation_combined.tsv`.

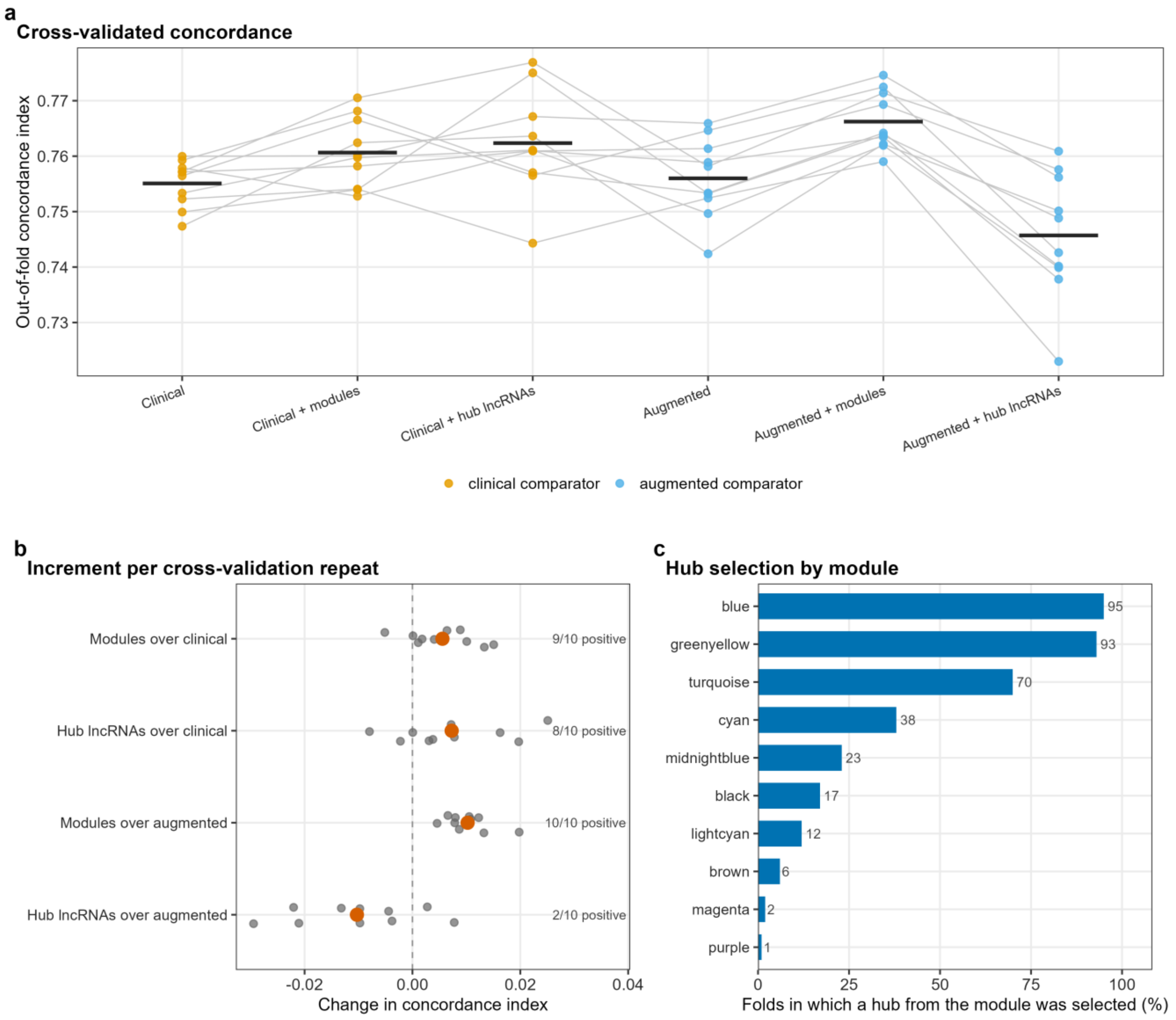


**Supplementary Fig. 11. Cross-validation of hub lncRNAs selected inside each fold. a** Out-of-fold concordance index of six models over ten repeats of ten-fold cross-validation in 493 discovery patients with 159 events. The arms are the clinical and augmented models, each alone, with the 29 module eigengenes or with up to 200 hub lncRNAs added under an elastic-net penalty. Lines join the same repeat, and bars are the mean over repeats. **b** Per-repeat increment for four comparisons, with the number of repeats in which it is positive. Grey points are single repeats and red points the mean over the ten repeats. **c** Percentage of the 100 training folds (10 repeats by 10 folds, bar labels giving fold counts) in which a hub lncRNA from each module entered the penalised model, 95 of 100 for lncRNA blue, 93 for greenyellow and 70 for turquoise. Because the candidate pool is chosen inside each training fold, the panel shows the stability of feature selection, not a fixed signature. Source tables: `05_cv_cindex_per_repeat.tsv`, `05_hub_module_selection_frequency.tsv`.

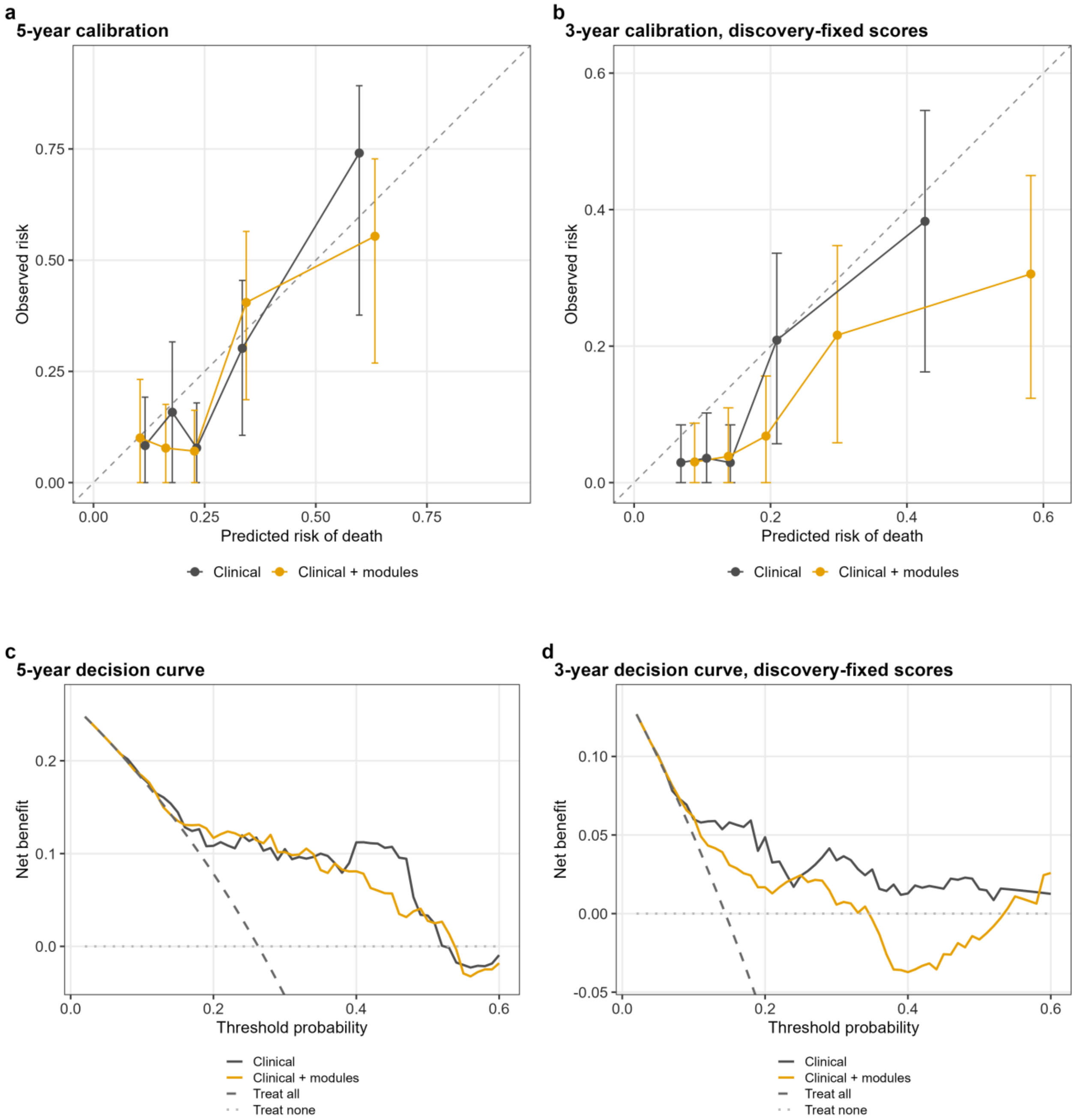


**Supplementary Fig. 12. Sensitivity analyses of calibration and clinical utility.** All panels are CPTAC-3, and error bars in **a** and **b** are Kaplan-Meier 95% confidence intervals. **a** Calibration at five years under cohort standardisation, by quintile of predicted risk, with the identity line. With twenty-three patients at risk at five years, this is the least stable estimate reported. The calibration slope is 0.97 (0.60 to 1.34) for the clinical model and 0.89 (0.58 to 1.20) with modules, and observed over expected risk 0.90 and 0.89. **b** Calibration at three years, the primary horizon, under discovery-fixed standardisation, with 95 patients at risk. Under discovery-fixed standardisation the combined model over-predicts risk about two-fold: the slope is 0.97 (0.61 to 1.34) for the clinical model and 0.89 (0.58 to 1.20) with modules, and observed over expected risk 0.76 and 0.55. **c** Decision curve at five years, cohort standardisation. The median change in net benefit over thresholds 0.10 to 0.40 is +0.003, positive at 19 of 31 thresholds. **d** Decision curve at three years, discovery-fixed standardisation, with a median change of -0.027, positive at 3 of 31 thresholds. In **c** and **d** the treat-all curve lies below the plotted range. All decision curves use the baseline hazard fixed in discovery, and none re-estimates it in CPTAC-3. Source tables: `12_calibration_bins.tsv`, `12_calibration_summary.tsv`, `12_validation_decision_curve.tsv`.

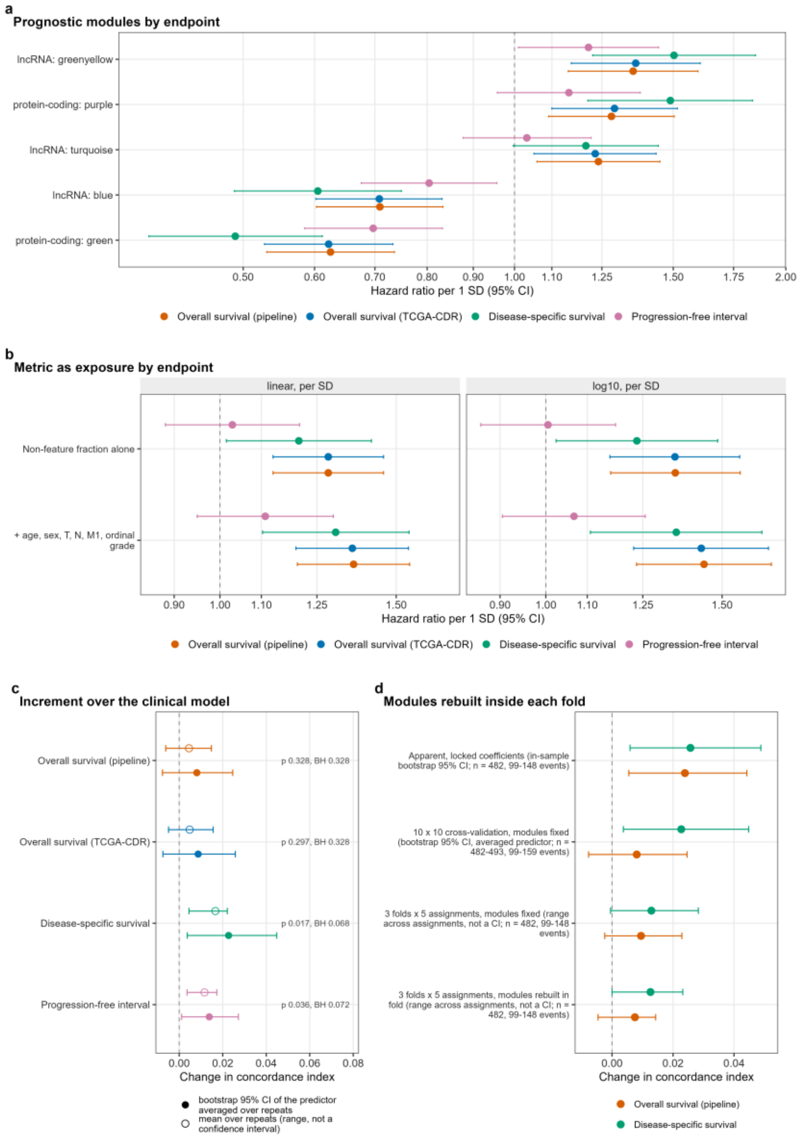


**Supplementary Fig. 13. Sensitivity of the results to the survival endpoint.** Events are deaths for overall survival, cancer-specific deaths for disease-specific survival, and progression or death for the progression-free interval. **a** The five prognostic modules under four endpoints in TCGA-KIRC (principal specification): overall survival from the pipeline and the Clinical Data Resource (502 to 504 patients, 163 to 165 deaths), disease-specific survival (491 to 493, 102 to 103) and the progression-free interval (500 to 502, 149 to 151). Sixteen of the 20 estimates reach a false discovery rate below 0.05. **b** The non-feature fraction as exposure under the same endpoints, on linear and log10 scales, with two models per scale (520 patients and 172 events for overall survival, 509 and 109 for disease-specific survival, 518 and 157 for the progression-free interval). The interval excludes 1 in all 12 overall and disease-specific estimates and in none of the 4 progression-free estimates. **c** Cross-validated increment in the concordance index from adding the 29 module eigengenes to the clinical model, by endpoint, in ten repeats of ten-fold cross-validation (493 patients and 159 events for overall survival, 482 and 99 for disease-specific survival, 491 and 146 for the progression-free interval). The interval excludes zero for disease-specific survival and the progression-free interval, and none of the four survives Benjamini-Hochberg adjustment over the endpoints (smallest adjusted p 0.068). The mean over the ten repeats is shown beside the bootstrap estimate, the increment of the repeat-averaged out-of-fold predictor, and is the smaller of the two for all four endpoints. Only the bootstrap estimate carries a confidence interval. **d** Fold-wise network rebuild for disease-specific and overall survival against matched fixed-module arms, with the ten-by-ten cross-validation rows for scale. Only the three-fold rows are matched arms, sharing patients, folds and inner splits and differing only in how modules are defined. Rebuilding lowers the increment by 0.0003 for disease-specific and by 0.0020 for overall survival. For disease-specific survival both three-fold rows (+0.0129 fixed, +0.0126 rebuilt) sit below the ten-by-ten value (+0.0227). Because the matched arms are nearly equal, this gap comes from the coarser cross-validation. Ranges span five fold assignments and are not confidence intervals. Source tables: `30_endpoint_modules.tsv`, `30_endpoint_exposure.tsv`, `30_endpoint_cv_increment.tsv`, `30_endpoint_foldwise.tsv`.

### Supplementary Tables
