## Supplementary Information for "Non-exonic read content reorganises lncRNA co-expression networks in renal and other cancers"

Supplementary Figures 1 to 13, Supplementary Tables 1 to 22 and the descriptions of Supplementary Data 1 to 5 follow the order of citation in the main article. Each value comes from the results file named in the note beneath its table. Superscript numbers refer to the main article's reference list, and Gene Ontology term names are quoted as published. A table cell reading n/a means not applicable. The non-feature read fraction is the percentage of reads that STAR assigns to no annotated feature, out of the unmapped, multimapping, non-feature, ambiguous and gene-assigned reads in its gene-counts summary. Hazard ratios are per standard deviation of the named quantity, with 95% confidence intervals. The clinical model comprises age, sex, T category, nodal status, M1 and ordinal grade. The augmented model adds the ESTIMATE stromal and immune scores and the three STAR metrics (non-feature fraction, multimapping fraction and log10 assigned reads). The principal specification for module models enters the module eigengene with the augmented covariates. Adjustment for the alignment metrics means residualisation of the expression matrix on the three STAR metrics before module definition. Benjamini-Hochberg control applies within the family stated in each note.

### Supplementary Figures

**Supplementary Fig. 1. The lncRNA axis under five normalisations.**

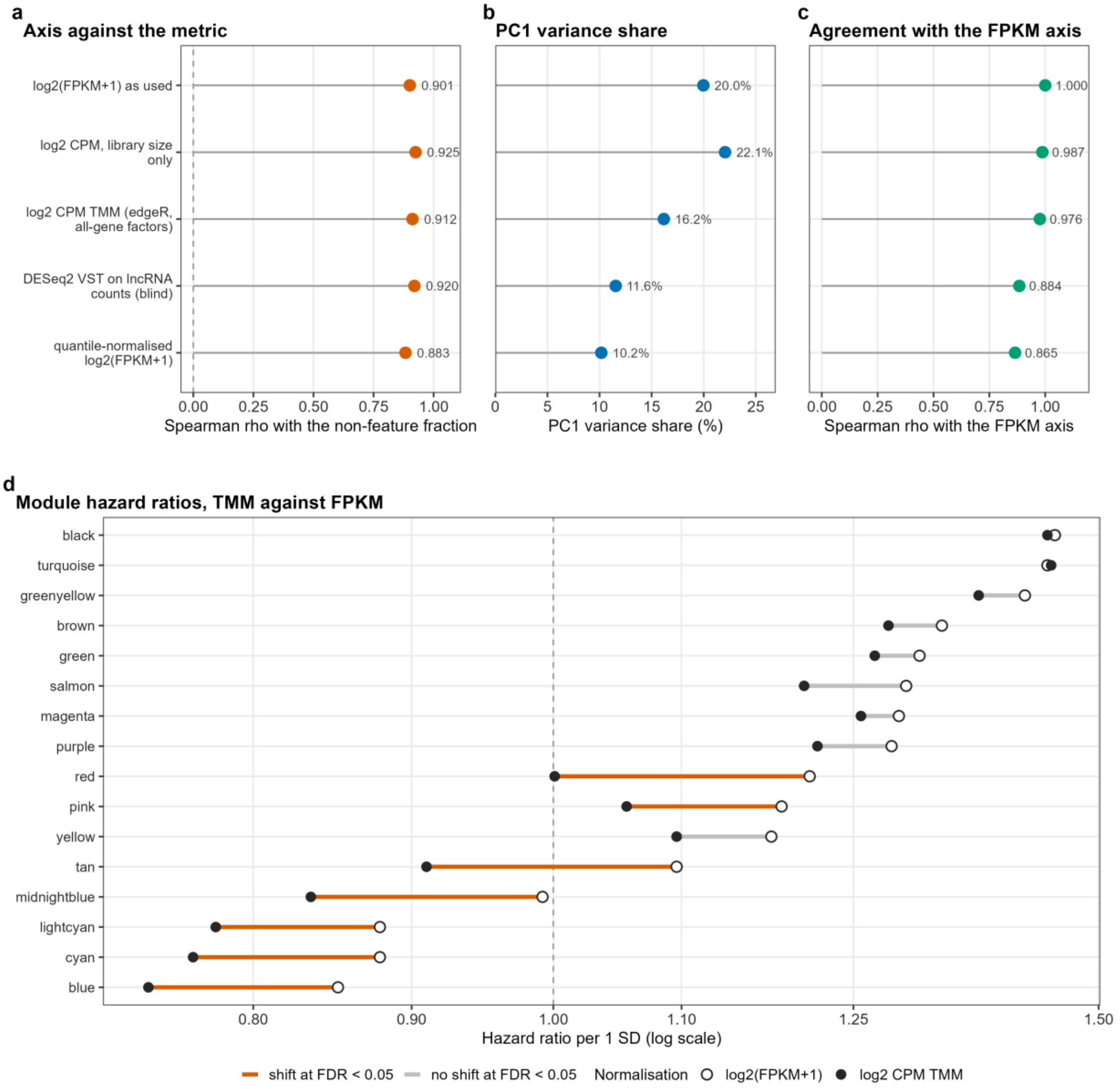


**a** Spearman correlation of the leading lncRNA principal component with the non-feature fraction under five normalisations of the same 3,442 genes and 511 libraries. The five arms are log2(FPKM + 1) as used throughout, log2 counts per million with library size only or with edgeR TMM factors, a blind DESeq2 variance-stabilising transform, and quantile-normalised log2(FPKM + 1). **b** PC1 variance share. Each PC1 is oriented against the log2(FPKM + 1) axis. **c** Spearman correlation of each arm's PC1 with the FPKM axis. Quantile normalisation removes the per-sample mean, leaving that arm's sign anchor indeterminate, so it is quoted as a magnitude. **d** Paired patient bootstrap difference in log hazard ratio between the TMM and FPKM scales for the 16 lncRNA modules (2,000 resamples, 504 patients, 163 deaths), with the same modules, patients and covariates on both scales. With Benjamini-Hochberg control within each arm, 7 of 16 differences reach a false discovery rate below 0.05 on the observed matrix and 0 of 16 on the residualised matrix. Before multiplicity control, 9 and 2 intervals exclude zero. Source tables: `29_normalisation_pc1.tsv`, `29_normalisation_module_contrast.tsv`.

**Supplementary Fig. 2. Matched normal-kidney libraries.**

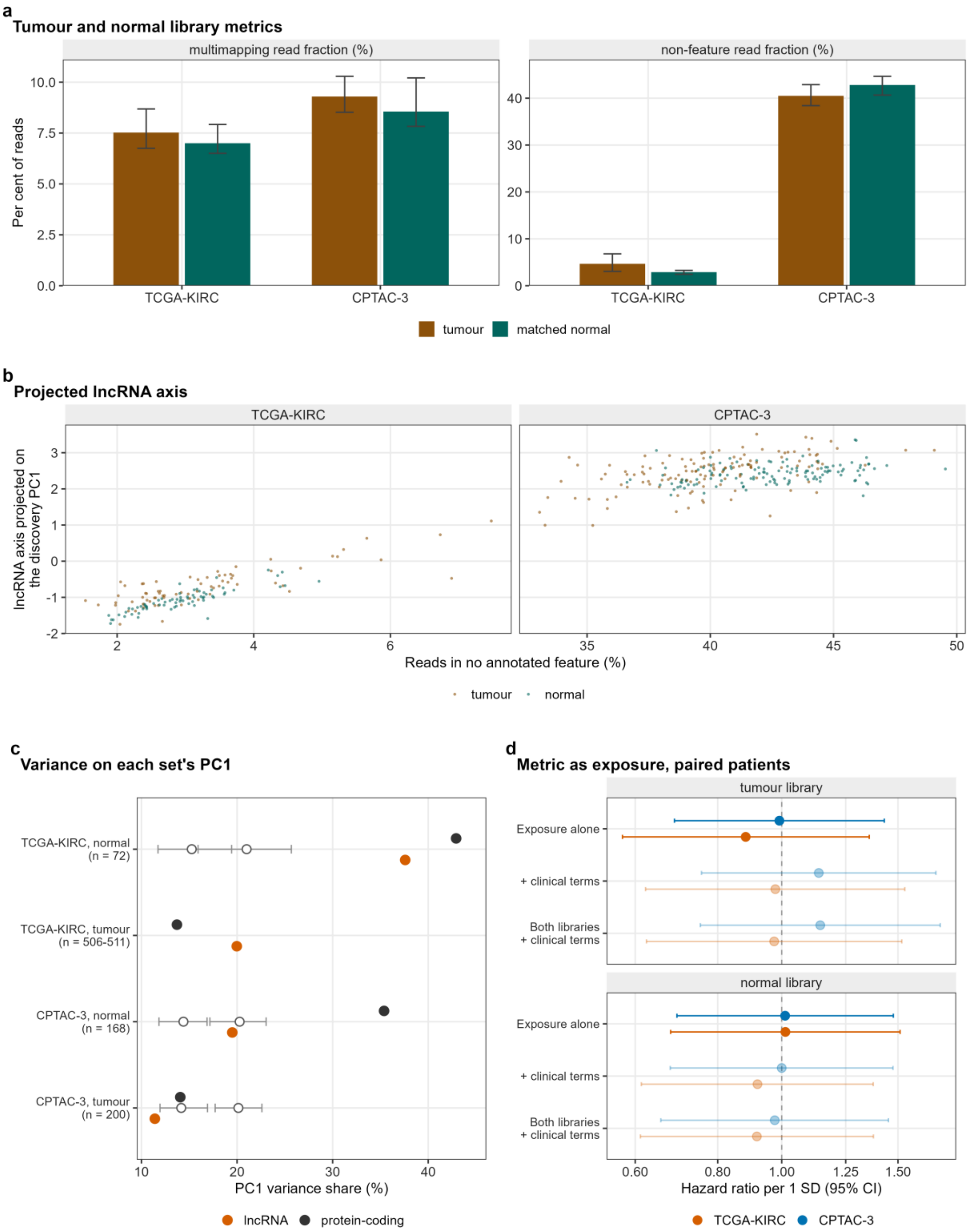


**a** Median and interquartile range of the multimapping and non-feature fractions in tumour and matched normal libraries of TCGA-KIRC (528 tumour, 72 normal) and CPTAC-3 (200 tumour,

168 normal). All four unpaired rank-sum tests reach a false discovery rate below 0.05, but the unpaired contrast mixes tissue with plate. In TCGA-KIRC the non-feature difference falls from 1.73 percentage points unpaired to 0.16 on the 68 plate-matched normal libraries (p 0.049), to 0.15 for the normals of the analysed tumours on shared plates (144 tumour and 67 normal libraries, p 0.057), and to 0.15 within patient (71 pairs, signed-rank p 0.10). **b** Projected lncRNA axis (the discovery lncRNA PC1 direction applied to each sample set) against the non-feature fraction in the paired libraries, faceted by cohort and coloured by tissue type: rho 0.77 in TCGA-KIRC tumours and 0.86 in their normals (71 pairs), and 0.54 and 0.35 in CPTAC-3 (129 pairs). **c** PC1 variance share of each sample set's own lncRNA and protein-coding matrix against a size-matched discovery baseline from 200 resampling draws (open grey points and bars, the median and range of the draws, available for the six non-discovery rows). The baseline range is not a confidence interval. Three of the eight observed shares lie above it. A share carries no sign, and the sign of three of the eight components cannot be fixed. **d** Forest plot of the non-feature fraction (linear scale, per standard deviation), measured on the tumour or the normal library, as exposure for overall survival in the paired patients under three models. Faded rows have an events-per-variable ratio below 5, as does every adjusted model in the panel (71 patients and 26 events in TCGA-KIRC, 129 and 28 in CPTAC-3). Source tables: `25_tumour_vs_normal_distribution.tsv`, `25_paired_scores_per_sample.tsv`, `25_normal_axis_projection.tsv`, `25_normal_metric_survival.tsv`.

**Supplementary Fig. 3. Network construction diagnostics.**

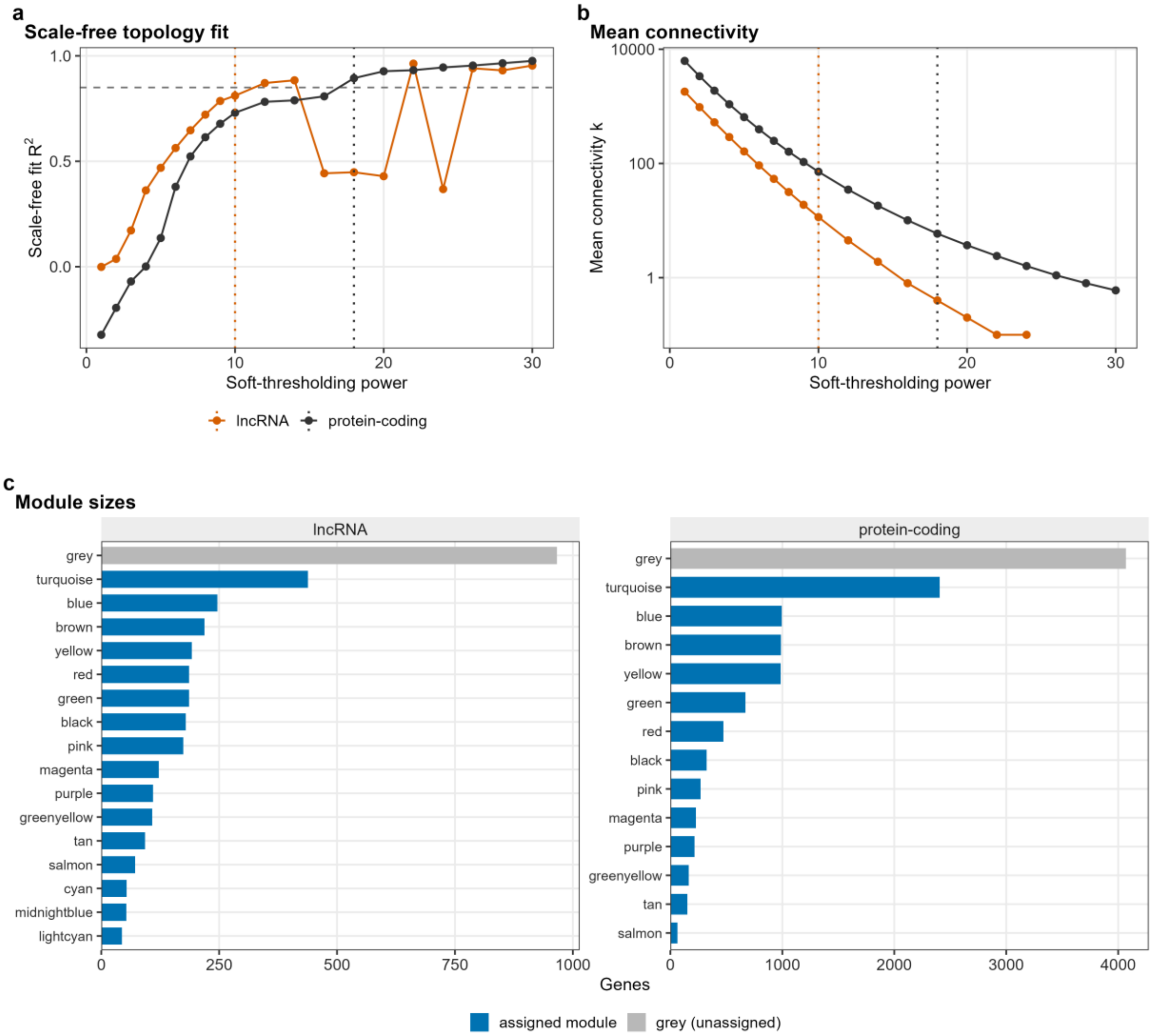


**a** Signed scale-free topology fit R-squared against soft-thresholding power for the protein-coding (12,000 genes, 506 samples) and lncRNA (3,442 genes, 511 samples) networks. The horizontal line marks the target of 0.85 and the vertical lines the chosen powers, 18 and 10, at which the signed fit is 0.894 and 0.811. The protein-coding power is the lowest reaching the target. The outcome-blind parameter sweep (Methods) fixed the lncRNA power at 10, where the automatic rule would have given 12. **b** Mean connectivity against power on a logarithmic axis, with the same chosen-power lines. **c** Module sizes, with unassigned (grey) genes shown separately. The protein-coding network resolves 13 non-grey modules, leaving 4,069 of 12,000 genes unassigned, and the lncRNA network 16, leaving 966 of 3,442. Source tables: `02_lncRNA_soft_threshold.tsv`, `02_mRNA_soft_threshold.tsv`, `02_network_summary.tsv`, `02_lncRNA_module_sizes.tsv`, `02_mRNA_module_sizes.tsv`.

**Supplementary Fig. 4. RNA integrity number and sequencing plate.**

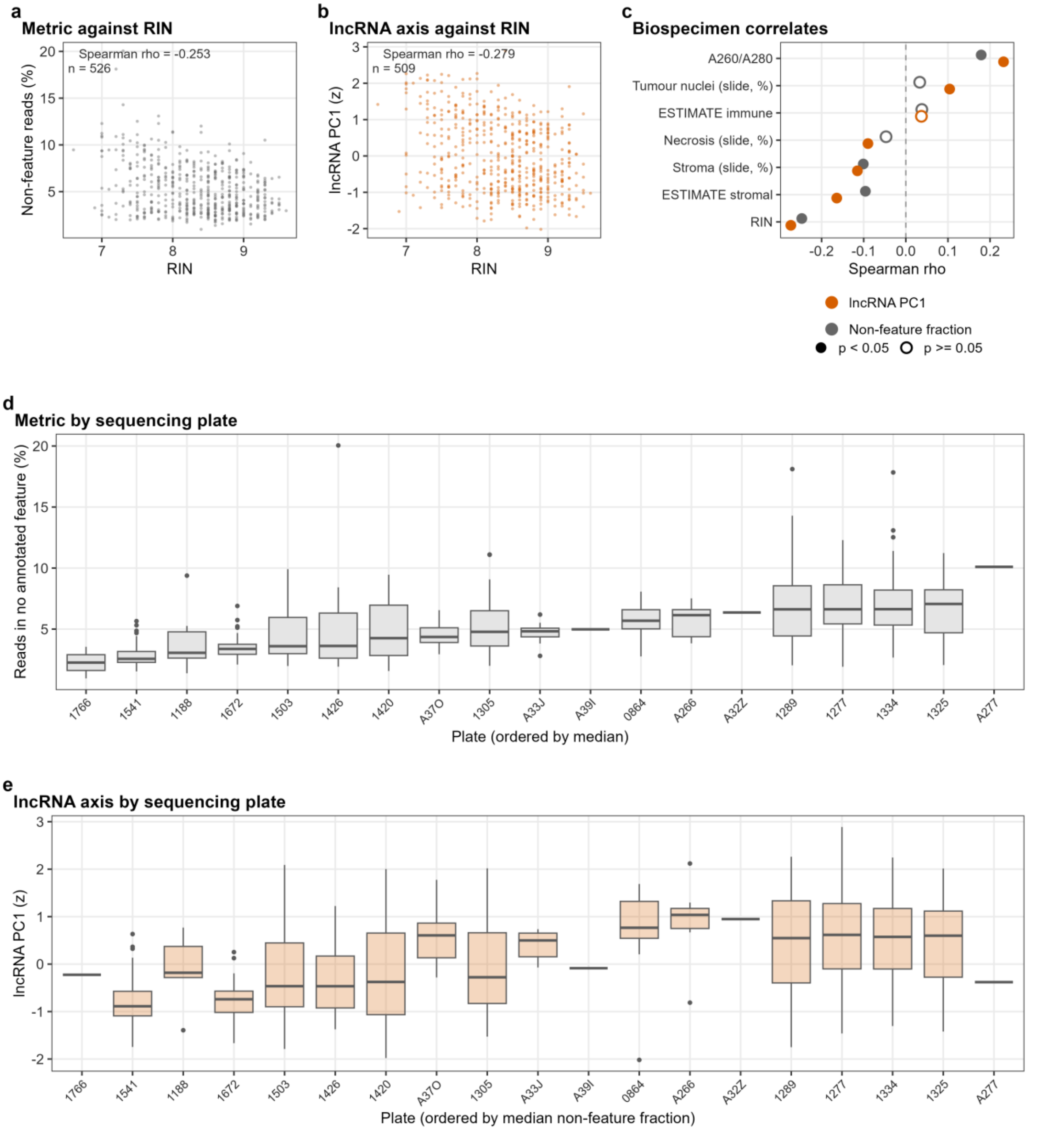


**a** Non-feature read fraction against RNA integrity number in TCGA-KIRC, with the integrity number from the Biospecimen Core Resource biospecimen record. One library with an RNA integrity number recorded as 0, 6.6 units below the next lowest, is treated as a coded missing value and excluded from panels **a** and **b**, so their sample size is one below the correlation table's. **b** Observed lncRNA PC1, the axis of Fig. 2a, against RNA integrity number, with the same exclusion. **c** Spearman correlation of the non-feature fraction and the lncRNA axis with seven biospecimen measures: RNA integrity number, A260/A280 ratio, slide percent necrosis, tumour nuclei and stromal cells, and the ESTIMATE stromal and immune scores (507 to 528 patients). Filled symbols mark the 10 of 14 tests with unadjusted p below 0.05. **d** Non-feature fraction by sequencing plate, ordered by median, for 528 libraries on 19 plates (1 to 59 per plate). **e** lncRNA axis by plate, in the same order, for the 511 libraries with an axis score. Biospecimen and batch metadata exist only for TCGA-KIRC. Source tables: `23_biospecimen_kirc.tsv`, `23_axis_vs_biospecimen.tsv`, `27_axis_scores_per_sample.tsv`.

**Supplementary Fig. 5. What the non-feature read fraction correlates with.**

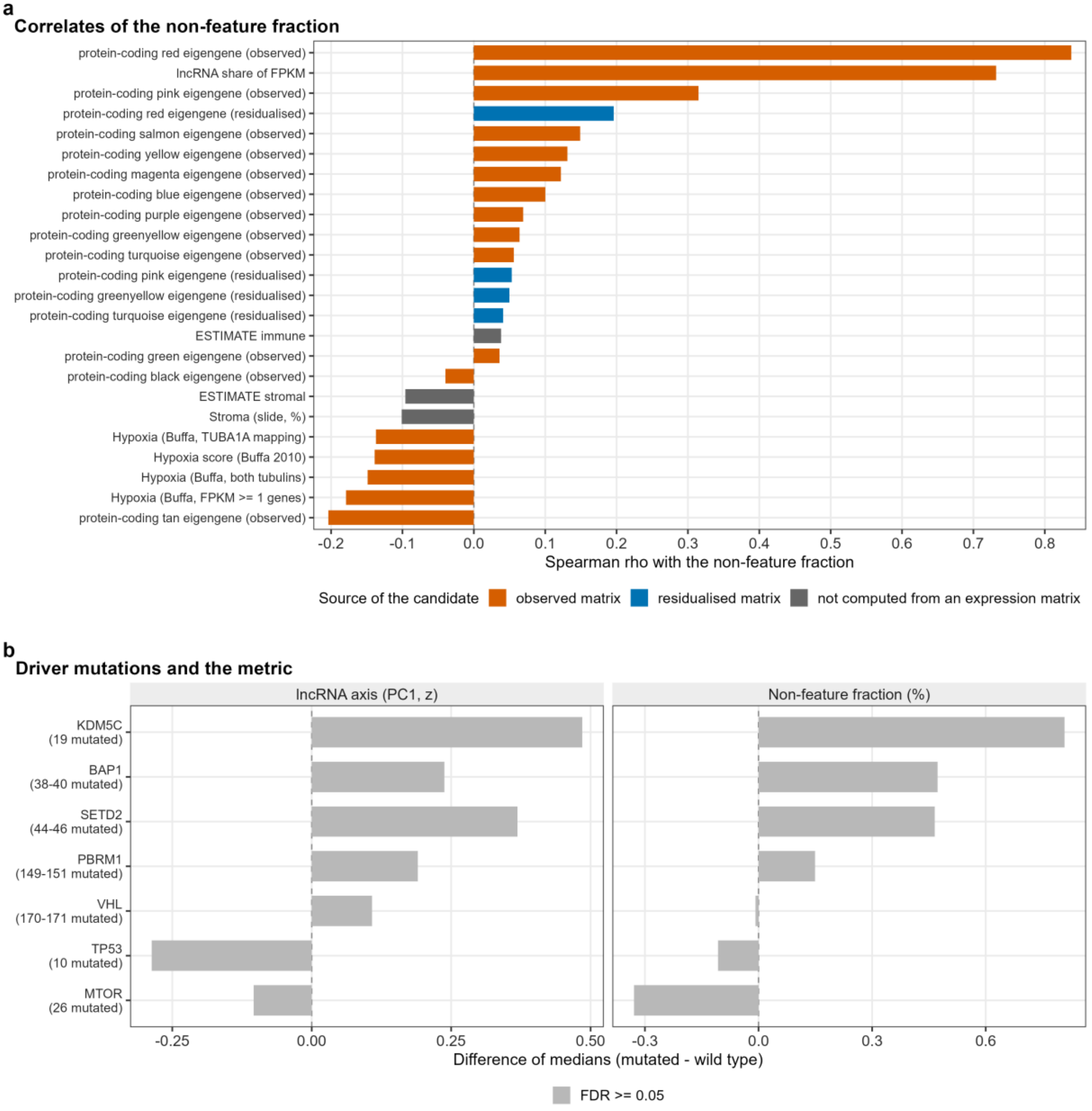


**a** The 24 strongest candidate correlates of the non-feature fraction in TCGA-KIRC (506 to 528 patients), coloured as eigengenes of the observed or the residualised expression matrix, or quantities computed outside it. Observed eigengenes cannot be read as biology, because the metric was measured on the same matrix. The annotated biological candidates are the hypoxia metagene, the proliferation score, the ESTIMATE stromal and immune scores and the slide tumour-nuclei percentage. **b** Difference of medians in the non-feature fraction and the lncRNA axis between mutated and wild-type patients for seven recurrent clear-cell drivers. None of the 14 tests (7 genes by 2 outcomes) reaches a 5% false discovery rate, on 358 to 368 patients per test and as few as 10 mutated patients. Testing is conditional on the 368 of 528 patients with an open-access masked mutation file, whose median non-feature fraction is lower than that of the others. Source tables: `27_metric_biology_correlations.tsv`, `28_noFeature_vs_mutations.tsv`.

**Supplementary Fig. 6. Single-cell ground truth for the correction.**

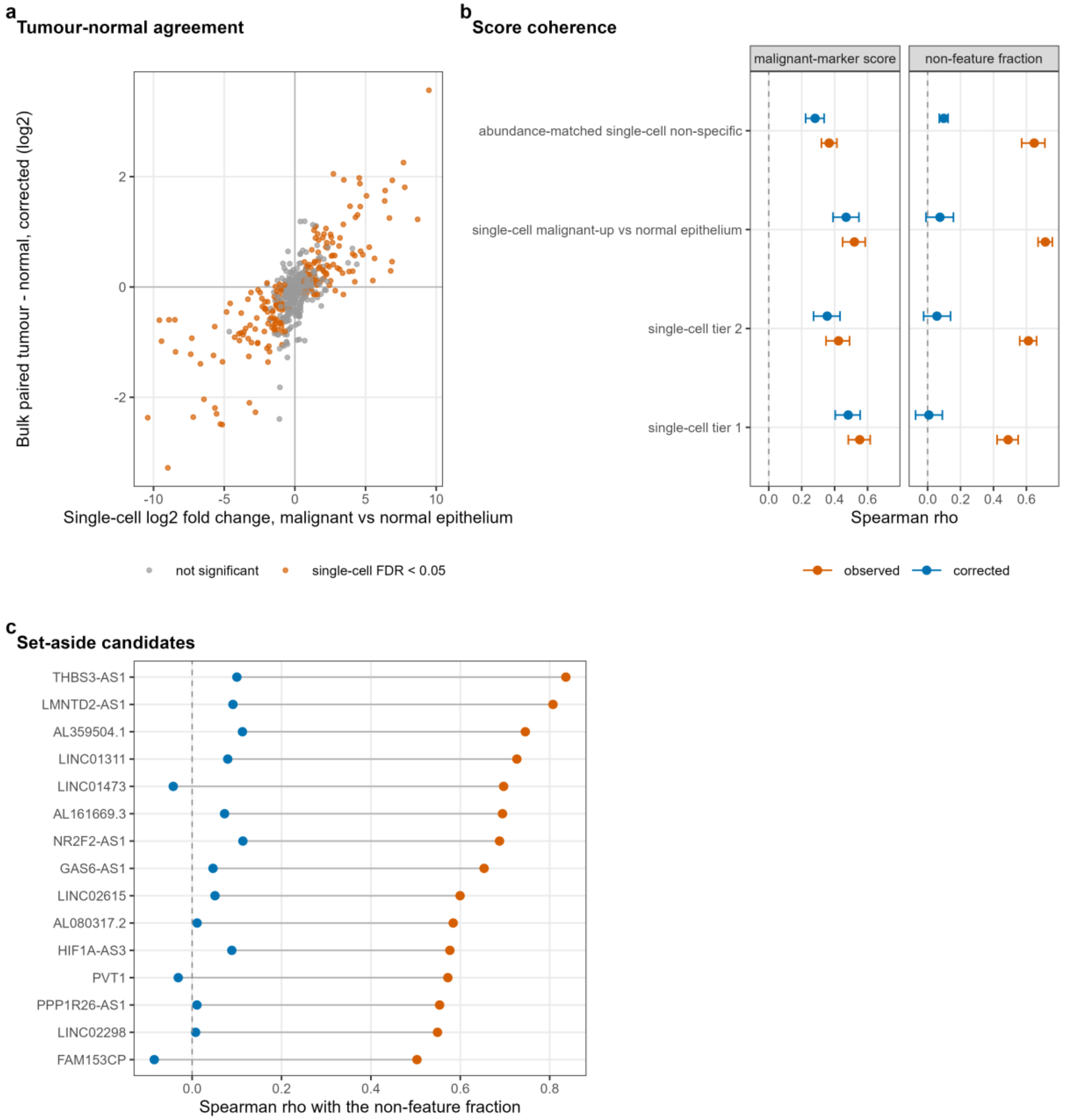


**a** Corrected bulk paired tumour-minus-normal difference against the single-cell fold change of malignant against normal epithelial cells, for 507 lncRNAs in 72 TCGA-KIRC patients. Orange, single-cell false discovery rate below 0.05. Agreement is Spearman rho 0.775 observed and 0.771 corrected, with sign agreement of 92.5% and 92.0% among single-cell-significant lncRNAs. **b** Spearman correlation of lncRNA scores in 532 tumours with the malignant-marker score and the non-feature fraction, observed (orange) and corrected (blue), with 95% bootstrap intervals. The marker score itself correlates with the metric at 0.22. The null row is 1,000 abundance-matched sets of single-cell non-specific lncRNAs. Correlation with the metric falls by construction, and the informative quantity is marker coherence relative to the null. **c** Correlation with the non-feature fraction, before and after correction, of the 15 single-cell candidates set aside in bulk as technically sensitive. Source tables: `42_sc_truth_concordance.tsv`, `42_sc_truth_score_coherence.tsv`, `42_sc_truth_technical_candidates.tsv`.

**Supplementary Fig. 7. Published lncRNA signatures against two random-signature nulls.**

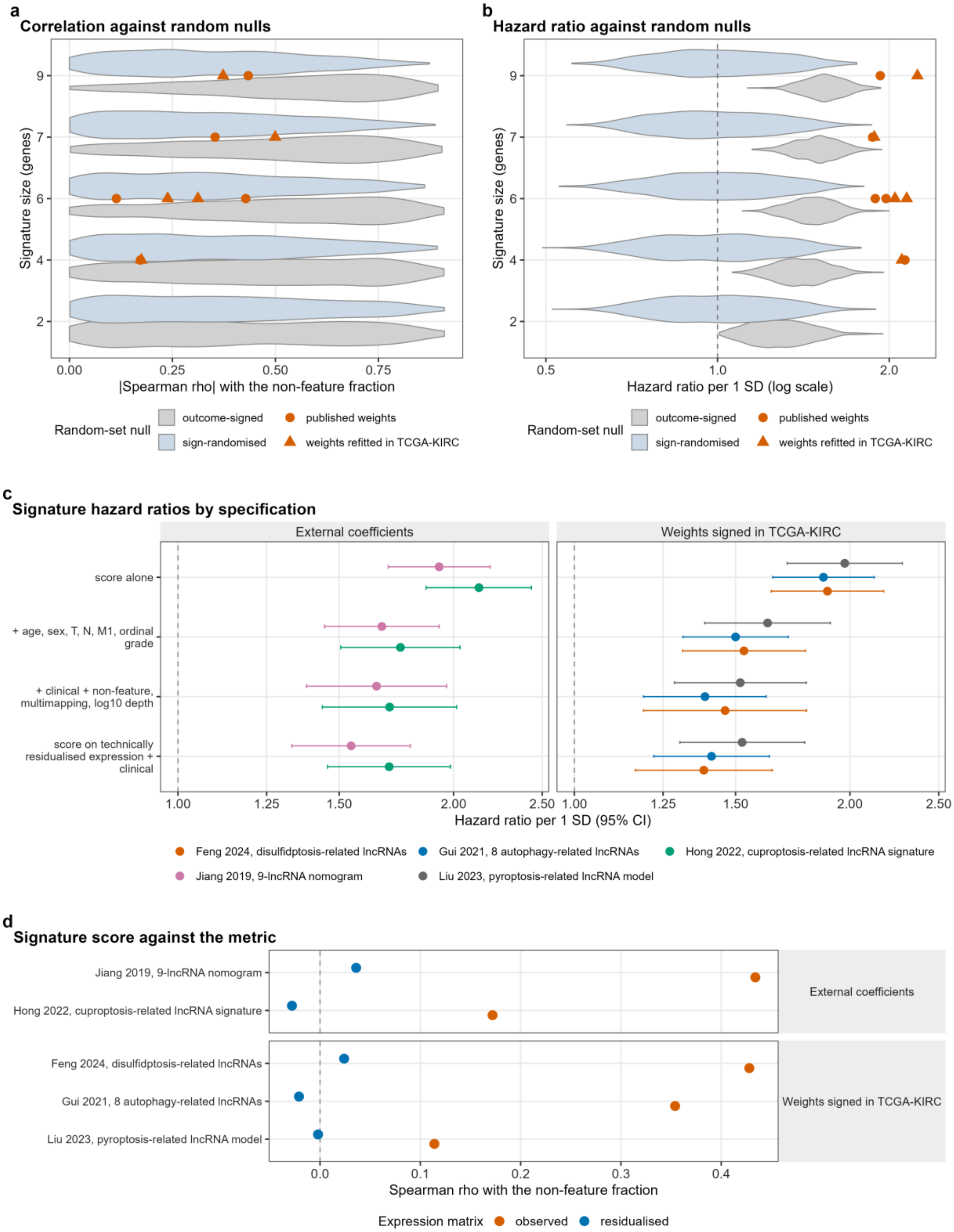


**a** Absolute Spearman correlation with the non-feature fraction (violins) for 1,000 random gene sets at each of five set sizes, under two weightings of the same draws: outcome-signed after

Venet and colleagues[11], and sign-randomised. The two nulls are the same 5,000 gene sets signed two ways. Points are the five primary-tier signatures, each with its published weights and with weights refitted in TCGA-KIRC. The Yu signature is held back for incomplete mapping. Against the sign-randomised null the signatures sit at percentiles 25.4 to 65.3 for the two with external coefficients and 19.1 to 64.9 for the three signed in sample, and against the outcome-signed null at 15.9 to 30.9 and 8.6 to 36.7. **b** Unadjusted hazard ratio per standard deviation (violins) for the same draws and weightings in 528 patients with 173 deaths, with the signatures overlaid. The sign-randomised null has a median hazard ratio of 1.00, whereas 95% of outcome-signed sets reach p below 0.05 by construction. The signatures sit at percentiles 99.8 to 100.0 (external coefficients) and 99.7 to 99.9 (signed in sample) of the outcome-signed null. **c** Hazard ratio per standard deviation (95% confidence interval) for the five primary-tier signatures under four specifications, grouped by weight provenance (520 to 528 patients, 172 to 173 deaths). All 20 intervals exclude 1. Adding the three STAR metrics changes the clinically adjusted log hazard ratio by -4.9% to -2.6% with external coefficients and -19.0% to -11.1% with weights signed in sample. Residualising the expression instead changes it by -15.0% to -5.1% and -23.6% to -13.2%. **d** Spearman correlation of each signature score with the non-feature fraction (528 patients), observed against residualised expression, by weight provenance: 0.172 to 0.434 observed with external coefficients and 0.114 to 0.428 signed in sample, and below 0.04 in absolute value for every signature after residualisation. Outcome-signing itself induces correlation with the metric in the null, so outcome-signed percentiles cannot show a signature to be less correlated with the metric than chance. Source tables:
`26_random_signature_null.tsv`, `26_published_signatures_models.tsv`,

26_published_signatures_correlations.tsv, 26_headline_ranges_by_provenance.tsv.

**Supplementary Fig. 8. Module preservation of the discovery networks in the three external cohorts.**

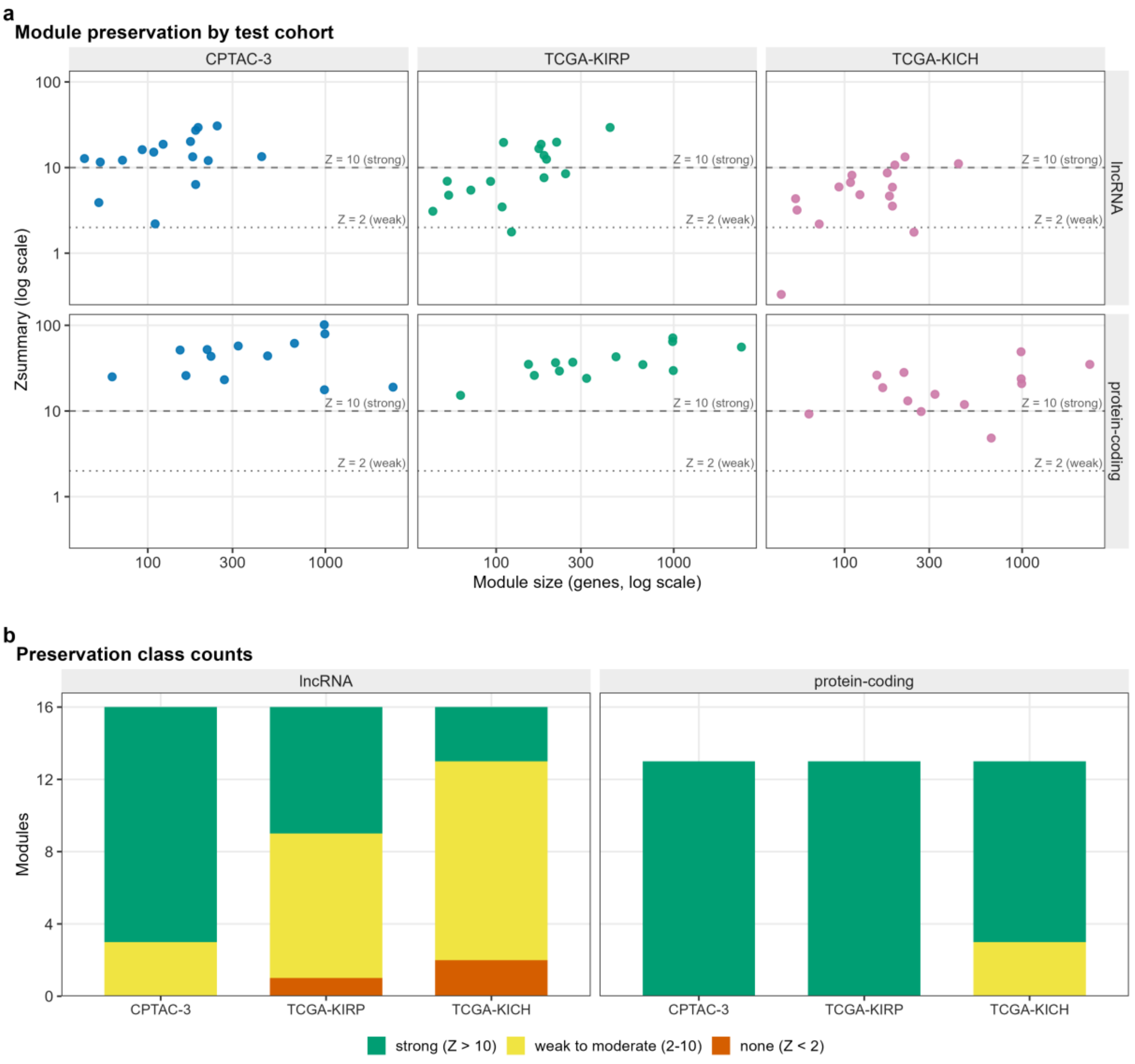


**a** WGCNA Zsummary against discovery module size by network and test cohort, for 29 modules and 87 module-cohort pairs, with lines at the conventional bands of Zsummary 2 and 10. Module size is the true discovery size, because the preservation function caps reported size at 1,000, which affects one module. The reference is the residualised TCGA-KIRC matrix used to define the modules, and each test network is residualised on its own cohort's STAR metrics. The lncRNA network used 100 permutations, the protein-coding network 30. **b** Stacked counts of modules strongly, weakly to moderately or not preserved, per cohort and network. Strongly preserved lncRNA modules number 13 of 16 in CPTAC-3, 7 of 16 in TCGA-KIRP and 3 of 16 in TCGA-KICH, and protein-coding modules 13 of 13, 13 of 13 and 10 of 13. The gold and grey rows of the underlying tables are WGCNA internal reference sets, not modules. Source tables: `24_module_preservation_lncRNA.tsv`, `24_module_preservation_mRNA.tsv`, `24_module_preservation_summary.tsv`, `02_lncRNA_module_sizes.tsv`, `02_mRNA_module_sizes.tsv`.

**Supplementary Fig. 9. Gene Ontology terms of correlated protein-coding partners and gene classes of the prognostic lncRNA modules.**

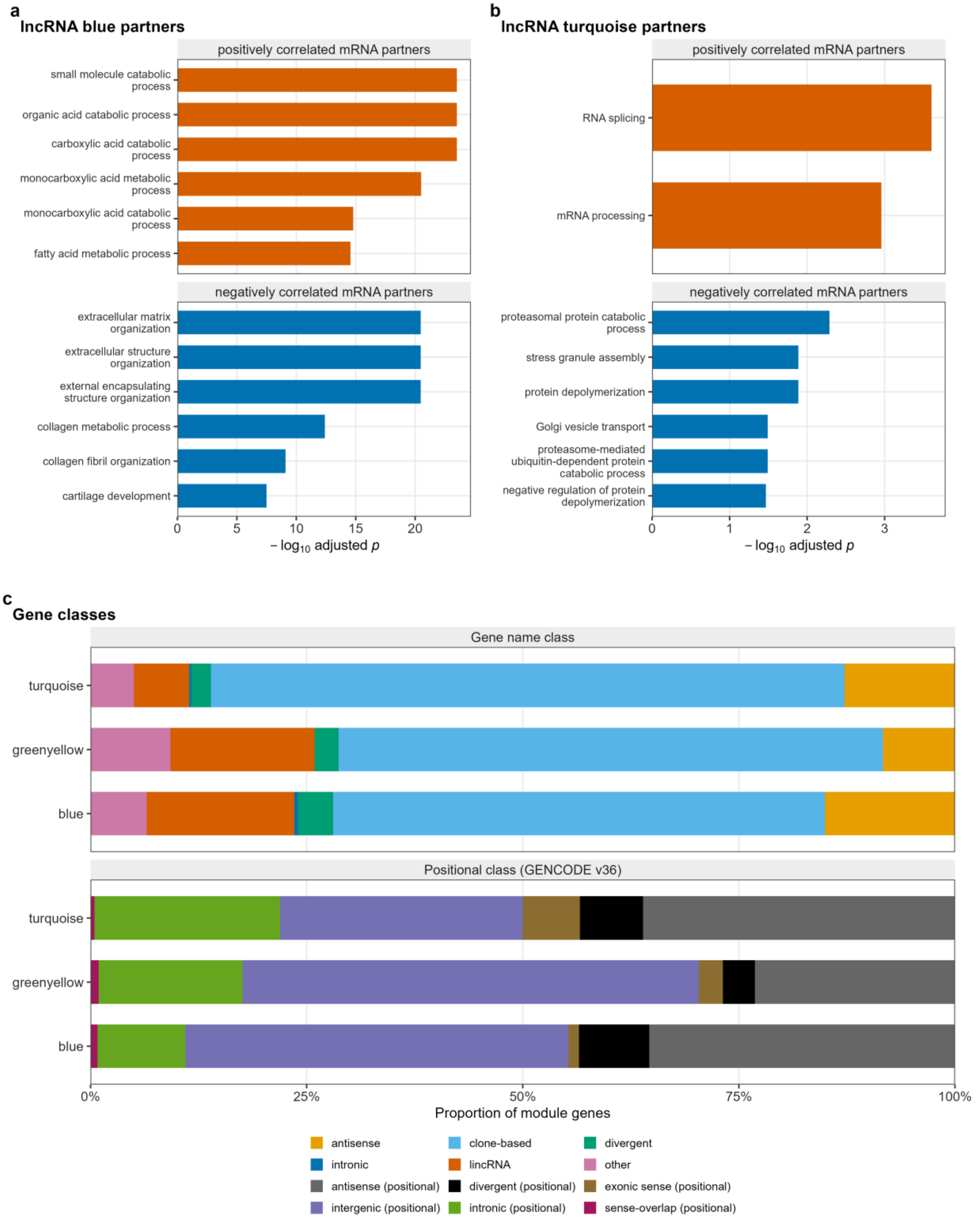


**a**, **b** Gene Ontology biological-process terms of the protein-coding partners of the lncRNA blue (**a**) and turquoise (**b**) modules, drawn as in Fig. 6c for greenyellow. Partners are split by the sign of their correlation with the module eigengene, the 300 most strongly correlated of each sign are tested separately against the 12,000 network protein-coding genes, and up to six terms with the smallest Benjamini-Hochberg adjusted p value are shown per sign. Turquoise has two enriched terms among its positive partners. Because partners are selected by correlation, the adjusted p values rank terms and are not calibrated tests. Under the principal specification the hazard ratio per standard deviation is 0.71 (0.60 to 0.83, false discovery rate 4.8e-04) for blue and 1.24 (1.06 to 1.45, 0.039) for turquoise. **c** Gene-class composition, as stacked proportions, of the three prognostic lncRNA modules, which hold 246 genes (blue), 108 (greenyellow) and 438 (turquoise), under two annotation schemes. The name-based classes are antisense, clone-based unnamed, divergent, intronic, named long intergenic and other. The positional classes, relative to GENCODE v36 protein-coding genes, are exonic sense, antisense, intronic, sense-overlapping, divergent and intergenic. Source tables: `10_lncRNA_guilt_by_association_GO_signed.tsv` (**a**, **b**), `10_lncRNA_module_composition.tsv` and `28_module_composition_by_class.tsv` (**c**).

#### Supplementary Fig. 10. Single-cell localisation of the 29 discovery modules.

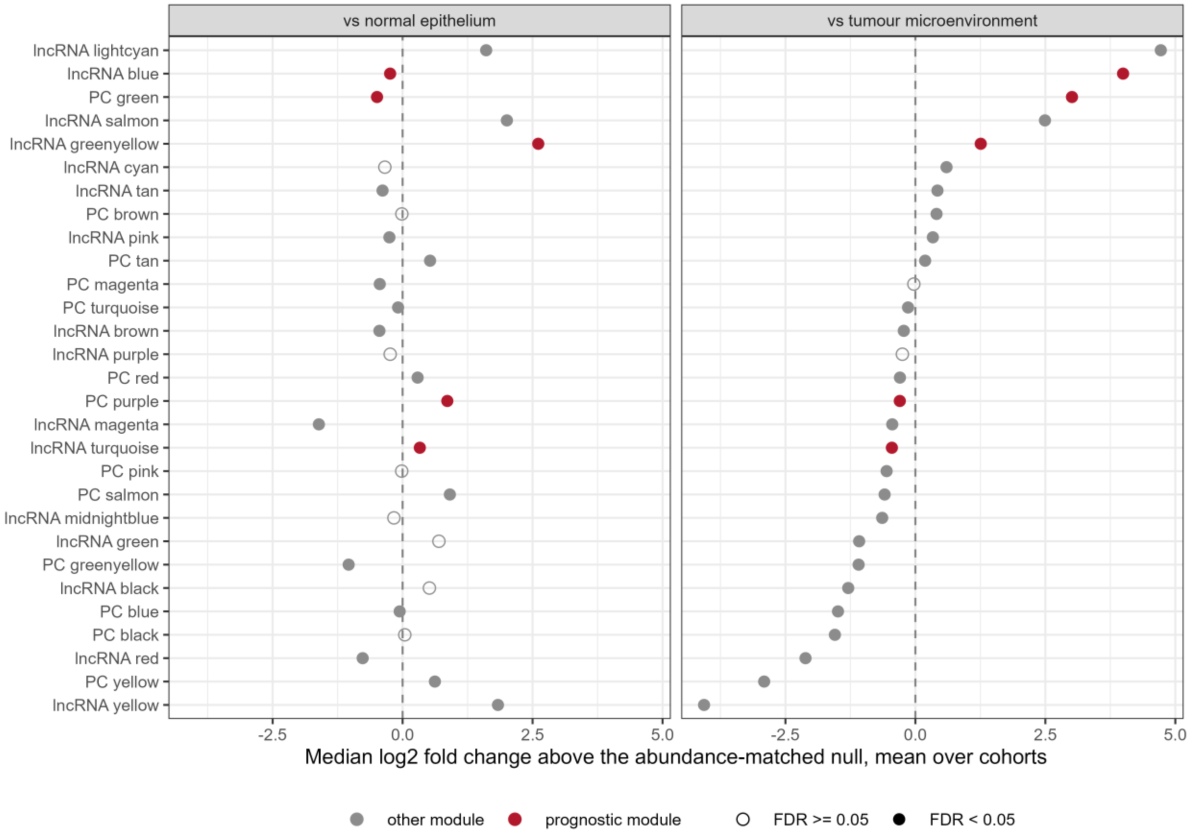


For each module, the median log2 fold change of its members minus the mean of 2,000 abundance-matched random gene sets, averaged over cohorts, for malignant cells against normal kidney epithelium (three cohorts, left) and against the rest of the tumour microenvironment (four cohorts, right). Positive values mean higher in malignant cells than a matched random set. Red points, the five prognostic modules. Filled symbols, a within-contrast false discovery rate below 0.05 by Stouffer combination of per-cohort z scores, which overstates precision because genes within a module are correlated. Per-cohort values are in Supplementary Table 19. Source table: `36_sc_module_localisation_combined.tsv`.

**Supplementary Fig. 11. Cross-validation of hub lncRNAs selected inside each fold.**

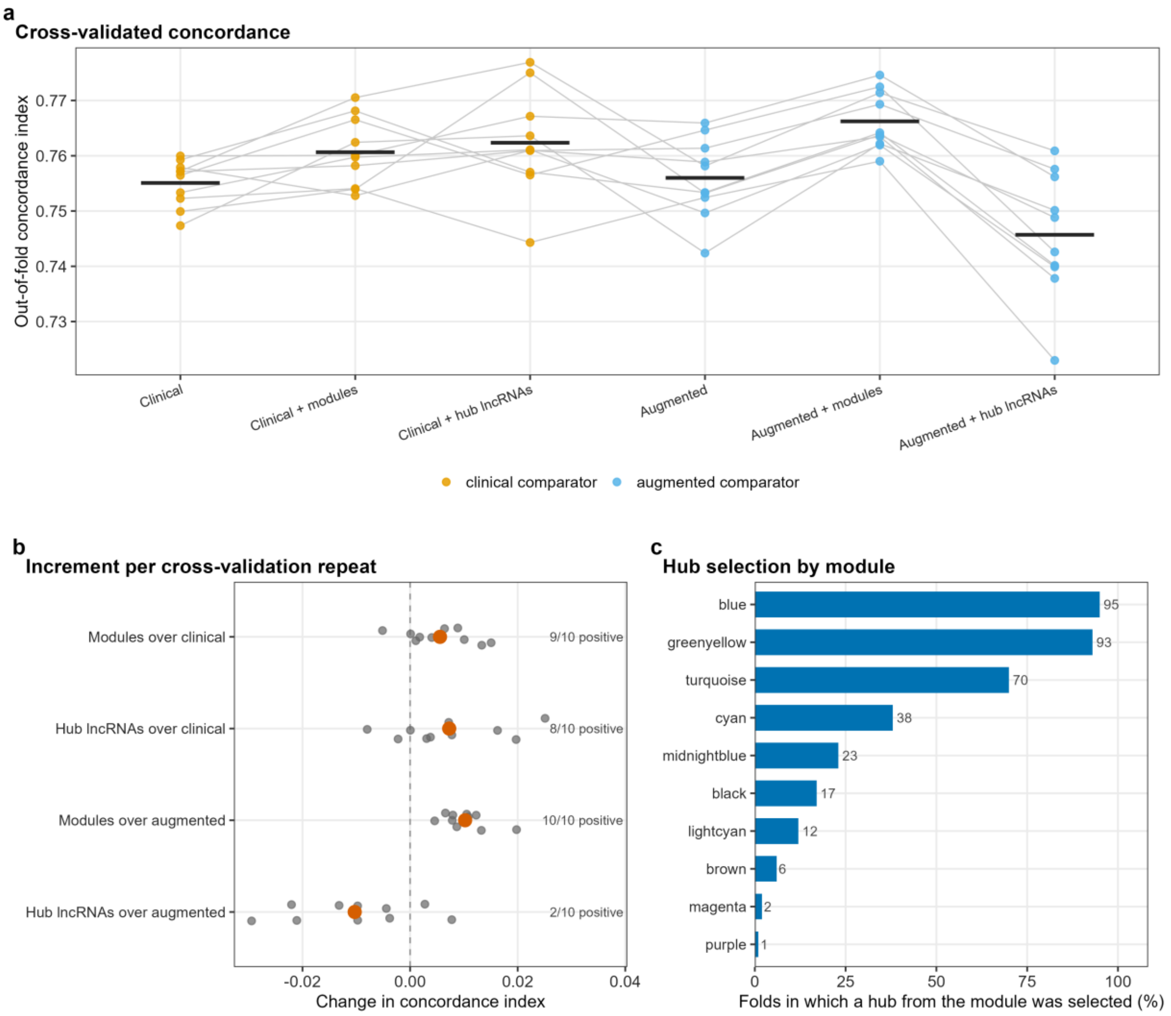


**a** Out-of-fold concordance index of six models over ten repeats of ten-fold cross-validation in 493 discovery patients with 159 events. The arms are the clinical and augmented models, each alone, with the 29 module eigengenes or with up to 200 hub lncRNAs added under an elastic-net penalty. Lines join the same repeat, and bars are the mean over repeats. **b** Per-repeat increment for four comparisons, with the number of repeats in which it is positive. Grey points are single repeats and red points the mean over the ten repeats. **c** Percentage of the 100 training folds (10 repeats by 10 folds, bar labels giving fold counts) in which a hub lncRNA from each module entered the penalised model, 95 of 100 for lncRNA blue, 93 for greenyellow and 70 for turquoise. Because the candidate pool is chosen inside each training fold, the panel shows the stability of feature selection, not a fixed signature. Source tables: `05_cv_cindex_per_repeat.tsv`, `05_hub_module_selection_frequency.tsv`.

**Supplementary Fig. 12. Sensitivity analyses of calibration and clinical utility.**

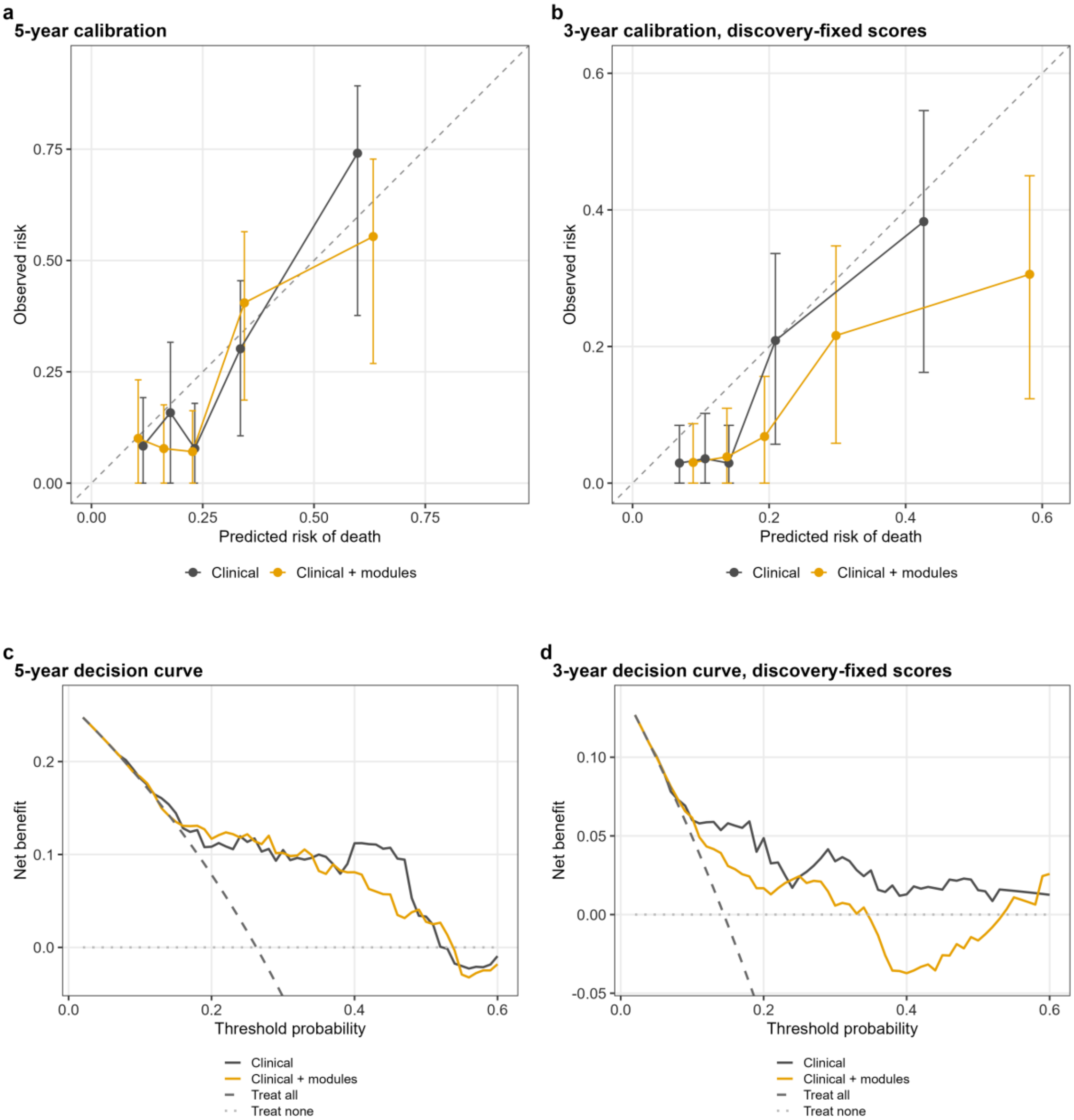


All panels are CPTAC-3, and error bars in **a** and **b** are Kaplan-Meier 95% confidence intervals. **a** Calibration at five years under cohort standardisation, by quintile of predicted risk, with the identity line. With twenty-three patients at risk at five years, this is the least stable estimate reported. The calibration slope is 0.97 (0.60 to 1.34) for the clinical model and 0.89 (0.58 to 1.20) with modules, and observed over expected risk 0.90 and 0.89. **b** Calibration at three years, the primary horizon, under discovery-fixed standardisation, with 95 patients at risk. Under discovery-fixed standardisation the combined model over-predicts risk about two-fold: the slope is 0.97 (0.61 to 1.34) for the clinical model and 0.89 (0.58 to 1.20) with modules, and observed over expected risk 0.76 and 0.55. **c** Decision curve at five years, cohort standardisation. The median change in net benefit over thresholds 0.10 to 0.40 is +0.003, positive at 19 of 31 thresholds. **d** Decision curve at three years, discovery-fixed standardisation, with a median change of -0.027, positive at 3 of 31 thresholds. In **c** and **d** the treat-all curve lies below the plotted range. All decision curves use the baseline hazard fixed in discovery, and none re-estimates it in CPTAC-3. Source tables: `12_calibration_bins.tsv`, `12_calibration_summary.tsv`, `12_validation_decision_curve.tsv`.

**Supplementary Fig. 13. Sensitivity of the results to the survival endpoint.**

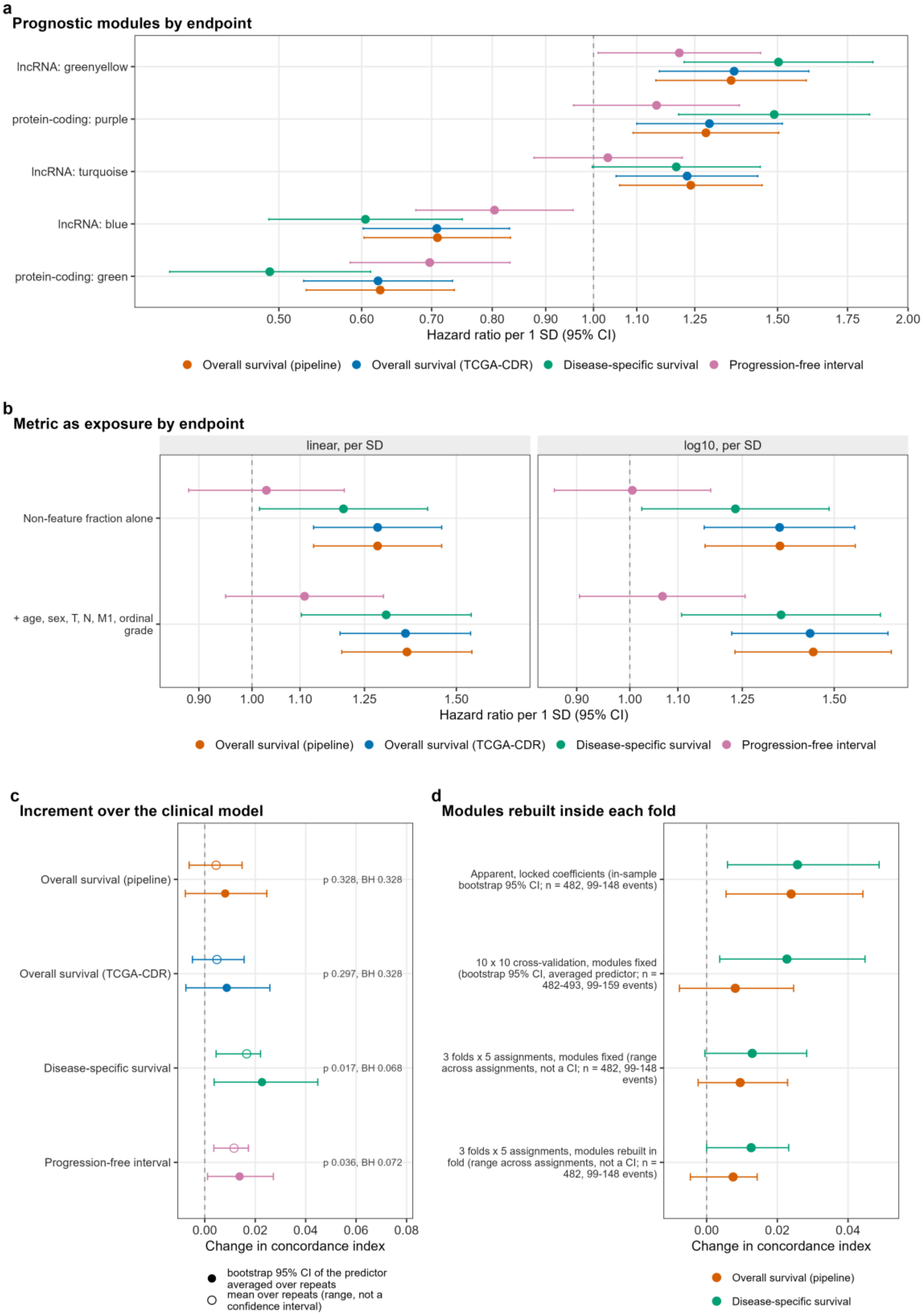

a
Prognostic modules by endpoint
lncRNA: greenyellow
protein-coding: purple
lncRNA: turquoise
lncRNA: blue
protein-coding: green
0.50
0.60
0.70
0.80
0.90
1.00
1.10
1.25
1.50
1.75
2.00
Hazard ratio per 1 SD (95% CI)
Overall survival (pipeline)
Overall survival (TCGA-CDR)
Disease-specific survival
Progression-free interval
b
Metric as exposure by endpoint
linear, per SD
log10, per SD
Non-feature fraction alone
+ age, sex, T, N, M1, ordinal grade
0.90
1.00
1.10
1.25
1.50
Hazard ratio per 1 SD (95% CI)
Overall survival (pipeline)
Overall survival (TCGA-CDR)
Disease-specific survival
Progression-free interval
c
Increment over the clinical model
Overall survival (pipeline)
p 0.328, BH 0.328
Overall survival (TCGA-CDR)
p 0.297, BH 0.328
Disease-specific survival
p 0.017, BH 0.068
Progression-free interval
p 0.036, BH 0.072
0.00
0.02
0.04
0.06
0.08
Change in concordance index
bootstrap 95% CI of the predictor averaged over repeats
mean over repeats (range, not a confidence interval)
d
Modules rebuilt inside each fold
Apparent, locked coefficients (in-sample bootstrap 95% CI; n = 482, 99-148 events)
10 x 10 cross-validation, modules fixed (bootstrap 95% CI, averaged predictor; n = 482-493, 99-159 events)
3 folds x 5 assignments, modules fixed (range across assignments, not a CI; n = 482, 99-148 events)
3 folds x 5 assignments, modules rebuilt in fold (range across assignments, not a CI; n = 482, 99-148 events)
0.00
0.02
0.04
Change in concordance index
Overall survival (pipeline)
Disease-specific survival

Events are deaths for overall survival, cancer-specific deaths for disease-specific survival, and progression or death for the progression-free interval. **a** The five prognostic modules under four endpoints in TCGA-KIRC (principal specification): overall survival from the pipeline and the Clinical Data Resource (502 to 504 patients, 163 to 165 deaths), disease-specific survival (491 to 493, 102 to 103) and the progression-free interval (500 to 502, 149 to 151). Sixteen of the 20 estimates reach a false discovery rate below 0.05. **b** The non-feature fraction as exposure under the same endpoints, on linear and log10 scales, with two models per scale (520 patients and 172 events for overall survival, 509 and 109 for disease-specific survival, 518 and 157 for the progression-free interval). The interval excludes 1 in all 12 overall and disease-specific estimates and in none of the 4 progression-free estimates. **c** Cross-validated increment in the concordance index from adding the 29 module eigengenes to the clinical model, by endpoint, in ten repeats of ten-fold cross-validation (493 patients and 159 events for overall survival, 482 and 99 for disease-specific survival, 491 and 146 for the progression-free interval). The interval excludes zero for disease-specific survival and the progression-free interval, and none of the four survives Benjamini-Hochberg adjustment over the endpoints (smallest adjusted p 0.068). The mean over the ten repeats is shown beside the bootstrap estimate, the increment of the repeat-averaged out-of-fold predictor, and is the smaller of the two for all four endpoints. Only the bootstrap estimate carries a confidence interval. **d** Fold-wise network rebuild for disease-specific and overall survival against matched fixed-module arms, with the ten-by-ten cross-validation rows for scale. Only the three-fold rows are matched arms, sharing patients, folds and inner splits and differing only in how modules are defined. Rebuilding lowers the increment by 0.0003 for disease-specific and by 0.0020 for overall survival. For disease-specific survival both three-fold rows (+0.0129 fixed, +0.0126 rebuilt) sit below the ten-by-ten value (+0.0227). Because the matched arms are nearly equal, this gap comes from the coarser cross-validation. Ranges span five fold assignments and are not confidence intervals. Source tables: `30_endpoint_modules.tsv`, `30_endpoint_exposure.tsv`, `30_endpoint_cv_increment.tsv`, `30_endpoint_foldwise.tsv`.

### Supplementary Tables

#### Supplementary Table 1. Driver mutations and the positional classes of the network lncRNAs

##### (a) Coverage of the discovery cohort by open-access masked mutation files

| Quantity | Value |
|---|---|
| Patients with a masked MAF | 368 of 528 (69.7%) |
| Primary-tumour MAF files used | 411 |
| Deaths among covered patients | 94 |
| Death rate, covered against uncovered | 25.5% against 49.4%, chi-square p 1.4e-07 |
| Overall survival by MAF availability | log-rank p 5.1e-05 |
| M1, covered against uncovered | 12.8% against 21.2%, p 0.019 |
| T3 or T4, covered against uncovered | 31.5% against 46.2%, p 0.0017 |
| Median non-feature fraction, covered against uncovered | 3.9% against 6.0%, Wilcoxon p 5.9e-07 |
| Coverage of the top decile of the metric | 52.8% |
| Coverage of the bottom decile of the metric | 86.8% |
| Median non-synonymous variants per covered patient | 45.5 (IQR 33 to 59) |

*Note.* Source: `28_mutation_coverage.tsv`. Mutation-file availability is associated with death, stage and the exposure itself. Any result in this subset is therefore conditional, and the absence of a genotype association is not an unconditional null.

##### (b) Driver frequencies and association with the metric and the axis

| Gene | Mutated of 368 (%) | Median metric, mutated against wild type (%) | p | FDR |
|---|---|---|---|---|
| VHL | 171 (46.5) | 3.9 against 3.9 | 0.68 | 0.69 |

| Gene | Mutated of 368 (%) | Median metric, mutated against wild type (%) | p | FDR |
|---|---|---|---|---|
| PBRM1 | 151 (41.0) | 4.1 against 3.9 | 0.39 | 0.69 |
| SETD2 | 46 (12.5) | 4.4 against 3.9 | 0.54 | 0.69 |
| BAP1 | 40 (10.9) | 4.4 against 3.9 | 0.29 | 0.69 |
| MTOR | 26 (7.1) | 3.7 against 4.1 | 0.45 | 0.69 |
| KDM5C | 19 (5.2) | 4.7 against 3.9 | 0.20 | 0.69 |
| TP53 | 10 (2.7) | 3.8 against 3.9 | 0.69 | 0.69 |
| Non-synonymous burden | n/a | Spearman rho -0.07 | 0.19 | 0.69 |

*Note.* Source: `28_mutation_frequency.tsv`, `28_noFeature_vs_mutations.tsv`.

Tested against the lncRNA axis on 358 patients, the same seven genes give raw p values of 0.065 to 0.88 (smallest false discovery rate 0.29). The largest difference of medians across the 16 tests, for KDM5C on 19 mutated patients, is 0.8 percentage points in the metric and 0.49 standard deviations in the axis, and neither is significant. Without an interval on the difference of medians, a tight null cannot be told from an uninformative one. The finding is no association at a 5% false discovery rate in the 368 patients with a masked MAF, which does not show that genotype fails to explain the metric.

**(c) The metric as exposure with genotype in the model**

| Model | HR | 95% CI | p | Events per variable |
|---|---|---|---|---|
| Metric + clinical model, covered subset | 1.31 | 1.10 to 1.57 | 0.0029 | 13.3 |
| + BAP1 | 1.29 | 1.07 to 1.55 | 0.0077 | 11.6 |
| + PBRM1 | 1.31 | 1.10 to 1.56 | 0.0029 | 11.6 |
| + SETD2 | 1.34 | 1.11 to 1.61 | 0.0019 | 11.6 |
| + BAP1 + PBRM1 + SETD2 | 1.32 | 1.09 to 1.60 | 0.0044 | 9.3 |
| + all seven driver genes | 1.33 | 1.09 to 1.62 | 0.0042 | 6.6 |
| + non-synonymous burden | 1.31 | 1.10 to 1.56 | 0.0026 | 11.6 |
| + all seven drivers + burden | 1.34 | 1.10 to 1.63 | 0.0032 | 6.2 |

*Note.* TCGA-KIRC, 361 patients and 93 events, overall survival, hazard ratio per standard deviation of the metric. Source: `28_mutation_survival_models.tsv`. Proportional-hazards p values across the eight models range from 0.036 to 0.058. The largest model has 6.2 events per variable, and its driver coefficients should not be interpreted alone. In the same file, TP53 has a clinically adjusted hazard ratio of 3.63 (1.53 to 8.61) on 10 mutated patients, and mutation burden 1.41 (1.10 to 1.80) per standard deviation of log1p(count). VHL violates proportional hazards in all three of its models.

**(d) Positional classes of the 3,442 analysed lncRNAs and their correlation with the metric**

| Positional class | Genes | Share of the set (%) | Median absolute rho with the metric | Fraction above 0.3 (%) | Mean observed PC1 loading | Median gene length (kb) |
|---|---|---|---|---|---|---|
| Intronic | 698 | 20.3 | 0.53 | 70.8 | 0.01512 | 1.01 |
| Exonic sense | 126 | 3.7 | 0.52 | 73.8 | 0.01566 | 11.60 |
| Sense-overlapping | 19 | 0.6 | 0.49 | 68.4 | 0.01179 | 28.00 |
| Antisense | 1,162 | 33.8 | 0.43 | 62.8 | 0.01261 | 8.16 |
| Intergenic | 1,122 | 32.6 | 0.25 | 43.5 | 0.01114 | 3.66 |
| Divergent | 315 | 9.2 | 0.25 | 43.2 | 0.01015 | 2.90 |
| All | 3,442 | 100.0 | 0.37 | 56.8 | 0.01252 | 3.55 |

*Note.* Classes are assigned per gene against GENCODE v36 protein-coding genes in a fixed priority order, so a lncRNA meeting several relations is counted once. Per-gene correlations use 511 libraries. Source: `28_lncRNA_positional_classes.tsv`, `28_quality_correlation_by_class.tsv`. The highest-signal subclass, intronic sense, has 317 genes, a median absolute rho of 0.59 and 78.2% above 0.3. No gene in any class falls below -0.3, so the correlation is positive wherever present. Positional class differs from Ensembl name class. Of the 1,122 positionally intergenic lncRNAs, 740 have clone-based names and only 264 are named lincRNAs.

**(e) Tests of the class effect**

| Quantity | Test | df | Statistic | p |
|---|---|---|---|---|
| Absolute rho with the metric, across the six classes | Kruskal-Wallis | 5 | 263.31 | 7.7e-55 |

| Quantity | Test | df | Statistic | p |
|---|---|---|---|---|
| Signed rho with the metric | Kruskal-Wallis | 5 | 265.65 | 2.4e-55 |
| Observed PC1 loading | Kruskal-Wallis | 5 | 109.97 | 4.2e-22 |
| Absolute rho, class entered after mean expression and log10 gene length | sequential ANOVA | 5 | F 54.80 | 6.2e-55 |
| PC1 loading, same adjustment | sequential ANOVA | 5 | F 18.29 | 5.8e-18 |
| PC1 loading against per-gene rho with the metric | Spearman | n/a | rho 0.78 | reported as 0 |
| PC1 score against the metric, sample level | Spearman | n/a | rho 0.90 | 1.4e-186 |

*Note.* 3,442 genes and 511 libraries. Source: `28_class_comparison_tests.tsv`. The class effect survives adjustment for expression level and gene length, so it is not a detectability artefact. It remains correlational, and residual confounding by other gene properties is not excluded.

**(f) Positional composition of the lncRNA modules**

| Module | Genes | Most enriched class (enrichment) | Goodness-of-fit p | FDR |
|---|---|---|---|---|
| green | 186 | intronic (2.0-fold) | 1.0e-06 | 4.3e-06 |
| pink | 174 | divergent (3.4-fold) | 1.0e-06 | 4.3e-06 |
| black | 179 | exonic sense (2.1-fold) | 1.0e-06 | 4.3e-06 |
| brown | 219 | divergent (2.6-fold) | 1.0e-06 | 4.3e-06 |
| cyan | 54 | divergent (3.4-fold) | 3.7e-04 | 7.9e-04 |
| tan | 93 | divergent (2.6-fold) | 2.6e-04 | 7.9e-04 |
| yellow | 192 | sense-overlapping (2.8-fold) | 3.3e-04 | 7.9e-04 |
| blue | 246 | intergenic (1.4-fold) | 3.6e-04 | 7.9e-04 |
| purple | 110 | intronic (1.5-fold) | 4.5e-04 | 8.5e-04 |
| lightcyan | 44 | intergenic (2.0-fold) | 0.0032 | 0.0046 |
| greenyellow | 108 | intergenic (1.6-fold) | 0.0032 | 0.0046 |
| turquoise | 438 | exonic sense (1.8-fold) | 0.0089 | 0.012 |
| magenta | 122 | divergent (2.0-fold) | 0.0095 | 0.012 |
| red | 186 | intergenic (1.3-fold) | 0.030 | 0.034 |
| salmon | 72 | sense-overlapping (2.5-fold) | 0.073 | 0.078 |
| midnightblue | 53 | sense-overlapping (3.4-fold) | 0.24 | 0.24 |

*Note.* Enrichment is relative to the class proportions of all 3,442 analysed lncRNAs. Source: `28_module_composition_by_class.tsv`. Over 16 modules and 6 classes on the 2,476 non-grey genes, the global test gives a chi-square of 558.24, with a simulated p at the floor of 1e-06 over one million replicates. Five module p values also sit on that floor and are upper bounds. Only midnightblue and salmon show no significant class deviation.

#### Supplementary Table 2. The lncRNA axis under five normalisations

##### (a) The leading lncRNA principal component under five normalisations

| Normalisation | PC1 variance share | rho with the non-feature fraction | p | rho with the cached FPKM axis | Absolute Pearson r with the per-sample mean | Per-gene median absolute rho |
|---|---|---|---|---|---|---|
| log2(FPKM + 1), as used | 0.1997 | 0.90 | 1.4e-186 | 1.00 | 0.988 | 0.37 |
| log2 counts per million, library size only | 0.2205 | 0.93 | 2.1e-215 | 0.99 | 0.985 | 0.37 |
| log2 counts per million, edgeR TMM | 0.1617 | 0.91 | 1.4e-199 | 0.98 | 0.956 | 0.25 |
| DESeq2 variance-stabilising transform, blind | 0.1155 | 0.92 | 4.7e-209 | 0.88 | 0.744 | 0.26 |
| Quantile-normalised log2(FPKM + 1) | 0.1017 | 0.88 in magnitude | 1.4e-169 | 0.87 | 0.084 | 0.28 |

*Note.* The same 3,442 genes and 511 libraries in every row. Source: `29_normalisation_pc1.tsv`. Raw counts re-read from the local files matched the STAR summary with a maximum absolute difference of 0 reads and 0 percentage points. Quantile normalisation removes the per-sample mean, leaving the quantile row's sign anchor indeterminate, so it is quoted as a magnitude. Per-sample median centring, whose sign is determinate, flips the sign and leaves the magnitude at 0.89.

##### (b) Size factors and compartment read shares against the metric

| Quantity | Median | rho with the non-feature fraction | p | rho with the FPKM axis |
|---|---|---|---|---|
| edgeR TMM normalisation factor | 0.9991 | 0.82 | 3.1e-129 | 0.81 |
| TMM effective library size | 5.815e7 | 0.38 | 1.7e-19 | 0.31 |
| DESeq2 median-of-ratios size factor | 1.045 | 0.13 | 0.0032 | 0.08 |
| DESeq2 size factor relative to library size | 1.030 | 0.46 | 3.4e-29 | 0.49 |
| Assigned reads | 5.854e7 | -0.06 | 0.15 | -0.13 |
| lncRNA read share, all 16,901 lncRNA genes | 0.01107 | 0.87 | 2.2e-166 | 0.90 |
| lncRNA read share, the 3,442 analysed lncRNAs | 0.00995 | 0.85 | 5.0e-151 | 0.90 |
| Protein-coding read share | 0.9671 | -0.32 | 3.8e-14 | -0.34 |

*Note.* TCGA-KIRC, 528 patients. Source: `29_size_factors_vs_quality.tsv`. The protein-coding read share and the FPKM-against-CPM per-sample scale are exact reciprocals, one measurement reported twice, so only one is given.

**(c) Module-level agreement between the FPKM and TMM scales**

| Quantity | Value |
|---|---|
| Modules significant at within-network FDR 0.05, observed arm, FPKM against TMM | 11 against 11 |
| The same, with the STAR metrics as covariates | 2 against 5 |
| The same, residualised arm | 7 against 5 |
| Lost, gained and robust on adjustment, FPKM against TMM | 8, 4, 3 against 6, 0, 5 |
| Median bootstrap difference in log HR, TMM minus FPKM, observed against residualised | -0.073 against -0.023 |
| Intervals excluding zero, observed against residualised | 9 of 16 against 2 of 16 |
| Surviving Benjamini-Hochberg within arm, observed against residualised | 7 of 16 against 0 of 16 |
| Modules whose FPKM and TMM intervals overlap | 16 of 16 |
| TMM point estimate inside the FPKM interval | 14 of 16 |
| Modules with the same significance call, observed, plus STAR, residualised | 8, 13 and 14 of 16 |
| Median eigengene correlation between scales, observed against residualised | 0.930 against 0.951 |
| Stability across 50 bootstrap seeds | 7 modules survive BH in 50 of 50 seeds |

The seven modules whose effect shifts detectably between scales on the observed matrix:

| Module | Bootstrap difference in log HR | 95% CI |
|---|---|---|
| red | -0.189 | -0.291 to -0.075 |
| tan | -0.186 | -0.282 to -0.097 |
| midnightblue | -0.173 | -0.276 to -0.085 |
| blue | -0.141 | -0.231 to -0.059 |
| cyan | -0.139 | -0.235 to -0.049 |
| lightcyan | -0.123 | -0.204 to -0.036 |
| pink | -0.115 | -0.198 to -0.048 |

*Note.* TCGA-KIRC, 504 patients and 163 events, 16 lncRNA modules, paired patient bootstrap over 2,000 resamples (128,064 Cox fits, 5 convergence warnings). Source: `29_normalisation_agreement.tsv`, `29_normalisation_module_summary.tsv`, `29_normalisation_module_contrast.tsv`, `29_bootstrap_seed_stability.tsv`. Two further modules, yellow (-0.070) and salmon (-0.075), have intervals excluding zero but fail Benjamini-Hochberg control (false discovery rates 0.076 and 0.085). Only the adjusted counts should be quoted. The bootstrap resamples patients only, holding module definitions, rotations, TMM factors and residualisation coefficients at their full-cohort values, so it compares two measurement scales, not two pipelines.

#### Supplementary Table 3. Matched normal-kidney libraries

##### (a) Libraries read and pairs formed

| Quantity | TCGA-KIRC | CPTAC-3 |
|---|---|---|
| Normal STAR-Counts files listed in the manifest and downloaded | 72 and 72 | 168 and 168 |
| Excluded | 0 | 0 |
| Normal libraries analysed | 72 | 168 |
| Tumour-normal pairs with survival | 71 | 129 |
| Deaths among the paired patients | 26 | 28 |
| Pairs on the same sequencing plate | 67 of 71 (94.4%) | not applicable |
| Normal libraries with a plate recorded | 72 | 0 |

*Note.* Source: `25_normal_consort.tsv`. All 168 CPTAC-3 normal libraries exceed the discovery non-feature cap of 30% as written, and, as for the CPTAC-3 tumours, none is excluded on that ground. No TCGA-KIRP or TCGA-KICH normal libraries are held locally. The 32 and 25 files at the GDC were not downloaded or analysed, so no subtype normal control exists (`25_subtype_normal_availability.tsv`).

**(b) The axis in normal kidney**

| Sample set | Matrix | n | Projected axis, rho with the metric (p) | Own PC1 variance share (size-matched discovery baseline) | Own PC1, rho with the metric (p) |
|---|---|---|---|---|---|
| TCGA-KIRC normal | lncRNA | 72 | 0.86 (8.9e-22) | 0.376 (0.210, 0.159 to 0.257) | 0.33 (0.0042) |
| TCGA-KIRC normal | protein-coding | 72 | 0.09 (0.48) | 0.429 (0.153, 0.117 to 0.194) | 0.30 in magnitude (0.010) |
| CPTAC-3 normal | lncRNA | 168 | 0.43 (4.1e-09) | 0.195 (0.203, 0.172 to 0.230) | 0.23 in magnitude (0.0030) |
| CPTAC-3 normal | protein-coding | 168 | 0.39 (2.2e-07) | 0.354 (0.144, 0.118 to 0.169) | 0.26 in magnitude (8.2e-04) |

*Note.* Source: `25_normal_axis_projection.tsv`. The projected axis is the discovery lncRNA or protein-coding PC1 direction applied to the sample set, and the own component is the set's own leading principal component. The two differ in the same samples, and lncRNA specificity holds for the projected axis but not for the own component. The size-matched baseline is the median of 200 resampling draws from the discovery matrix at the same sample size, with their minimum and maximum in brackets, and is not a confidence interval. Three of the eight own-component rows in this file carry a sign set by convention, not by the data, because the primary anchor is weak or the anchors disagree, and are quoted as magnitudes. Supplementary Table 4 gives the full six-sample-set comparison on the projected quantity.

**(c) Library metrics and axis scores, tumour against matched normal, TCGA-KIRC**

| Quantity | Tumour median | Normal median | Unpaired difference (p) | Plate-matched difference (p) | Within-patient difference (signed-rank p) |
|---|---|---|---|---|---|
| Non-feature fraction, % | 4.6 | 2.9 | 1.73 (1.6e-14) | 0.15 (0.057) | 0.15 (0.099) |
| Multimapping fraction, % | 7.5 | 7.0 | 0.48 (0.011) | -0.06 (0.18) | -0.14 (0.069) |
| Unmapped fraction, % | 3.3 | 2.7 | 0.53 (0.085) | 0.95 (6.0e-04) | 0.22 (0.0094) |
| Ambiguous fraction, % | 8.5 | 8.2 | 0.35 (7.0e-08) | 0.37 (5.5e-05) | 0.37 (0.0033) |
| log10 assigned reads | 7.767 | 7.804 | -0.037 (0.0015) | -0.030 (0.0050) | -0.020 (0.11) |
| Projected lncRNA axis | -0.117 | -1.094 | 0.977 (1.1e-20) | 0.397 (1.0e-08) | 0.372 (6.6e-06) |
| Projected protein-coding axis | 0.104 | 0.707 | -0.603 (9.1e-08) | -0.439 (1.4e-04) | -0.370 (3.7e-04) |
| Mean lncRNA expression | 0.718 | 0.617 | 0.102 (2.7e-10) | 0.013 (0.10) | 0.009 (0.20) |

*Note.* Source: `25_tumour_vs_normal_distribution.tsv`, rows for the normals of analysed-tumour patients. Unpaired comparisons use 528 tumour and 71 normal libraries (rank-sum test), plate-matched comparisons the 144 tumour and 67 normal libraries on plates both sides occupy, and within-patient comparisons the 71 pairs (signed-rank test). Medians of the four percentage metrics are given to one decimal and the three difference columns to two, because several plate-matched and within-patient differences fall below 0.1 percentage points. The file flags every TCGA-KIRC row as confounded with plate, since the tumours span 19 plates and the normals 5. For the non-feature fraction the unpaired difference is about eleven times the plate-matched one and must never be quoted alone. In CPTAC-3 the direction reverses, with 40.5% in 200 tumours against 42.5% in 129 paired normals, a within-patient difference of -2.22 points (signed-rank p 1.6e-10).

**(d) Whether the metric is shared between a patient's tumour and normal library: the primary test**

| Quantity | Role | Spearman rho, tumour against normal | p | FDR | Outcome |
|---|---|---|---|---|---|
| Non-feature fraction | primary | -0.02 | 0.88 | 0.88 | null |

| Quantity | Role | Spearman rho, tumour against normal | p | FDR | Outcome |
|---|---|---|---|---|---|
| Projected lncRNA axis | primary | -0.17 | 0.16 | 0.27 | null |
| Multimapping fraction | companion | -0.05 | 0.68 | 0.85 | null |
| log10 assigned reads | companion | 0.40 | 4.8e-04 | 0.0024 | shared |
| Projected protein-coding axis | intended negative control | 0.37 | 0.0013 | 0.0033 | shared |

*Note.* TCGA-KIRC, 71 pairs. Source: `25_paired_tumour_normal.tsv`. In CPTAC-3 (129 pairs) the values are 0.17 (FDR 0.15) for the metric, 0.11 (FDR 0.30) for the lncRNA axis and 0.24 (FDR 0.033) for the protein-coding axis. This test failed to discriminate and supports neither reading. It is also weak. Plate explains 7.8% of the variance in the metric within the paired libraries against 30.9% across all 600 analysed libraries, so the pairs carry little of the plate variation that drives the metric elsewhere. Nor is the intended negative control clean. In the discovery tumours the projected protein-coding axis tracks the multimapping fraction (rho -0.52), so its sharing within patient is evidence of a patient-level library component, not of tumour biology.

**(e) Variance in the metric and in the axis explained by plate and tissue type, by library set**

| Library set | Quantity | Factor | Libraries | Variance explained (%) | p |
|---|---|---|---|---|---|
| All analysed libraries | Non-feature fraction | plate | 600 | 30.9 | 1.1e-38 |
| Tumour libraries only | Non-feature fraction | plate | 528 | 26.6 | 6.5e-27 |
| Normal libraries only | Non-feature fraction | plate | 72 | 7.2 | 0.16 |
| Paired patients | Non-feature fraction | plate | 142 | 7.8 | 0.010 |
| Paired patients | Non-feature fraction | tissue type given patient | 142 | 6.1 partial | 0.037 |
| All analysed libraries | Projected lncRNA axis | plate | 600 | 31.2 | 3.4e-39 |
| Tumour libraries only | Projected lncRNA axis | plate | 528 | 25.6 | 1.4e-25 |

| Library set | Quantity | Factor | Libraries | Variance explained (%) | p |
|---|---|---|---|---|---|
| Normal libraries only | Projected lncRNA axis | plate | 72 | 3.9 | 0.43 |
| Paired patients | Projected lncRNA axis | plate | 142 | 0.8 | 0.79 |
| Paired patients | Projected lncRNA axis | tissue type given patient | 142 | 27.2 partial | 2.7e-06 |

*Note.* TCGA-KIRC. Source: `25_paired_variance_partition.tsv`. Sixty-eight of the 72 normal libraries sit on 3 of the 5 plates they span, against 19 plates for the 528 tumours, so the plate partition is a tumour-side result absent within the normals. CPTAC-3 lacks plate and site metadata and has no equivalent rows.

**(f) Prognostic value of the normal library's metric**

| Exposure library | Exposure | Model | HR | 95% CI | p |
|---|---|---|---|---|---|
| Normal | Non-feature fraction | alone | 1.01 | 0.68 to 1.51 | 0.95 |
| Normal | Non-feature fraction | + clinical model | 0.92 | 0.61 to 1.38 | 0.68 |
| Tumour | Non-feature fraction | alone | 0.88 | 0.57 to 1.36 | 0.57 |
| Tumour | Non-feature fraction | + clinical model | 0.98 | 0.62 to 1.54 | 0.92 |
| Normal | Projected lncRNA axis | alone | 1.00 | 0.67 to 1.47 | 0.98 |
| Normal | Projected lncRNA axis | + clinical model | 0.95 | 0.65 to 1.40 | 0.80 |
| Tumour | Projected lncRNA axis | alone | 0.83 | 0.54 to 1.28 | 0.40 |

*Note.* TCGA-KIRC, 71 paired patients and 26 events. Source: `25_normal_metric_survival.tsv`. Every adjusted model falls below the events-per-variable threshold of 5 (ratios 3.2 to 3.7), and the smallest false discovery rate in the 32-row table is 0.995 in TCGA-KIRC and 0.999 in CPTAC-3. The tumour metric is also null in this 71-patient subset, so the panel reflects low power and does not contradict the discovery result. Two of the 32 rows, both for the tumour axis alone, violate proportional hazards.

#### Supplementary Table 4. The projected lncRNA axis across all six sample sets

| Sample set | Tissue | Library protocol | n | Projected lncRNA axis, rho with the metric (p) | Projected protein-coding axis, rho with the metric (p) | Projected protein-coding axis, rho with the multimapping fraction |
|---|---|---|---|---|---|---|
| TCGA-KIRC | primary tumour | poly(A) | 528 | 0.90 (4.4e-194) | 0.06 (0.18) | -0.52 |
| TCGA-KIRP | primary tumour | poly(A) | 269 | 0.89 (2.6e-92) | 0.18 (0.0034) | -0.45 |
| TCGA-KIRC | matched normal | poly(A) | 72 | 0.86 (8.9e-22) | 0.09 (0.48) | 0.12 |
| TCGA-KICH | primary tumour | poly(A) | 65 | 0.59 (1.8e-07) | 0.29 (0.018) | -0.66 |
| CPTAC-3 | primary tumour | ribo-depleted total RNA | 200 | 0.55 (7.4e-17) | 0.29 (2.6e-05) | -0.05 |
| CPTAC-3 | matched normal | ribo-depleted total RNA | 168 | 0.43 (4.1e-09) | 0.39 (2.2e-07) | -0.51 |

Each cohort's own leading component, a different quantity in the same samples:

| Sample set | Matrix | n | Own PC1 variance share | Size-matched discovery baseline, median (minimum to maximum) | Own PC1, rho with the metric (p) |
|---|---|---|---|---|---|
| TCGA-KIRC tumour | lncRNA | 511 | 0.200 | not applicable | 0.90 (1.4e-186) |
| TCGA-KIRP tumour | lncRNA | 269 | 0.267 | 0.202 (0.181 to 0.219) | 0.83 (7.3e-69) |
| TCGA-KIRC normal | lncRNA | 72 | 0.376 | 0.210 (0.159 to 0.257) | 0.33 (0.0042) |
| TCGA-KICH tumour | lncRNA | 65 | 0.145 | 0.208 (0.165 to 0.269) | -0.13 (0.32) |
| CPTAC-3 tumour | lncRNA | 200 | 0.114 | 0.201 (0.177 to 0.226) | 0.20 (0.0041) |
| CPTAC-3 normal | lncRNA | 168 | 0.195 | 0.203 (0.172 to 0.230) | 0.23 in magnitude (0.0030) |

*Note.* The projected axis is the discovery lncRNA or protein-coding PC1 direction, defined by its gene loadings and the discovery gene means, applied to each sample set. The own component is the set's own leading principal component. Projected and own-component rows are different quantities and are not pooled. Source: `25_normal_axis_projection.tsv` and `35_axis_projection_subtypes.tsv`. Every projected row has an interpretable sign, inherited from the fixed discovery rotation. Of the own-component rows, three of the eight in the first file are not sign-interpretable and are quoted as magnitudes, and none of the four in the second is flagged.

On the projected quantity the axis is strong in every poly(A) sample set and materially weaker, though far from absent, under ribo-depleted total RNA. The separation is graded, not categorical, with the weakest poly(A) set, TCGA-KICH at 0.59, beside the strongest ribo-depleted set, CPTAC-3 tumour at 0.55. TCGA-KICH has 65 patients over a narrow exposure range, with a median non-feature fraction of 3.1% (interquartile range 2.6% to 3.5%) against 4.6% (3.1% to 6.8%) in TCGA-KIRC. Restricted range attenuates a correlation, so 0.59 is a lower bound. The projected value of 0.59 does not contradict this cohort's own component of -0.13. Its own lncRNA PC1 carries 14.5% of variance, below the size-matched discovery baseline of 20.8%, so no dominant leading component exists for the metric-linked direction to occupy. The direction is present without being first, which is why a fixed projected direction is used throughout.

The lncRNA specificity itself depends on protocol. Under poly(A) selection the projected lncRNA and protein-coding axes differ sharply, at 0.90 against 0.06 in the discovery tumours and 0.86 against 0.09 in the matched normals. Under ribo-depletion the difference nearly vanishes, at 0.43 against 0.39 in the CPTAC-3 normals. A protocol that retains unprocessed RNA raises intronic content for every gene, not only for lncRNAs overlapping protein-coding loci, so the effect is no longer lncRNA-specific.

#### Supplementary Table 5. The lncRNA network built with and without adjustment for the alignment metrics

| Comparison | Genes | Adjusted Rand index | Modules, unadjusted | Modules, adjusted |
|---|---|---|---|---|
| all genes, grey as a class | 3,442 | 0.062 | 3 | 16 |

| Comparison | Genes | Adjusted Rand index | Modules, unadjusted | Modules, adjusted |
|---|---|---|---|---|
| genes assigned in both networks | 1,722 | 0.047 | 3 | 16 |

Fate of each adjusted module in the unadjusted network (dominant unadjusted module: turquoise, 1,982 genes):

| Adjusted module | Genes | In the unadjusted dominant module (%) | In another unadjusted module (%) | Unassigned in the unadjusted network (%) |
|---|---|---|---|---|
| black | 179 | 100 | 0 | 0 |
| green | 186 | 99.5 | 0.5 | 0 |
| turquoise | 438 | 99.5 | 0 | 0.5 |
| salmon | 72 | 84.7 | 0 | 15.3 |
| pink | 174 | 66.7 | 0.6 | 32.8 |
| purple | 110 | 66.4 | 0 | 33.6 |
| brown | 219 | 61.6 | 0 | 38.4 |
| tan | 93 | 53.8 | 1.1 | 45.2 |
| magenta | 122 | 40.2 | 0.8 | 59 |
| blue | 246 | 36.6 | 0 | 63.4 |
| midnightblue | 53 | 34 | 1.9 | 64.2 |
| yellow | 192 | 32.8 | 49.5 | 17.7 |
| red | 186 | 25.3 | 47.3 | 27.4 |
| cyan | 54 | 18.5 | 1.9 | 79.6 |
| lightcyan | 44 | 13.6 | 0 | 86.4 |
| greenyellow | 108 | 13 | 0.9 | 86.1 |

Positional class of the lncRNAs in the unadjusted dominant module:

| Positional class | Network lncRNAs | In the dominant module | % | Fisher p | FDR |
|---|---|---|---|---|---|
| sense overlapping | 19 | 14 | 73.7 | 0.17 | 0.17 |
| intronic | 698 | 499 | 71.5 | 2.9e-17 | 8.7e-17 |
| exonic sense | 126 | 88 | 69.8 | 0.0044 | 0.0065 |
| antisense | 1162 | 732 | 63 | 4.3e-06 | 8.6e-06 |
| divergent | 315 | 159 | 50.5 | 0.0084 | 0.01 |
| intergenic | 1122 | 490 | 43.7 | 3e-30 | 1.8e-29 |

*Note.* Both networks used the same 511 discovery libraries and parameters, the unadjusted network on observed log2(FPKM + 1) and the adjusted network on expression residualised on the three STAR metrics. Mean intramodular connectivity over assigned genes is 70.5 unadjusted and 3.3 adjusted, each on its own matrix (`23_unadjusted_lncRNA_network_summary.tsv`). Adjusted and unadjusted module colours do not correspond. The false discovery rate is controlled across the six positional classes. The dominant module holds 57.6% of all network lncRNAs. Source tables: `37_network_partition_agreement.tsv`, `37_adjusted_module_fate.tsv`, `37_dominant_module_positional_class.tsv`.

#### Supplementary Table 6. The lncRNA axis, network reorganisation and prognosis of the metric across TCGA

*(a) Primary tumours, one row per project, with eligible projects (at least 100 retained tumours, TCGA-LAML excluded) entering the counts in the main text*

| Project | Eligible | Tumours | Median non-feature % | lncRNA PC1 rho | Protein-coding PC1 rho | Rule A | Intronic minus intergenic | Plate R-squared of metric | Spread (MAD of log metric) | Largest observed module | Largest adjusted module | ARI | Largest-module eigengene r | Rule C | Metric HR, plate-stratified (95% CI) | Deaths |
|---|---|---|---|---|---|---|---|---|---|---|---|---|---|---|---|---|
| GBM | yes | 256 | 9.1 | 0.95 | 0.87 | not met | 0.44 | 0.61 | 1.11 | 0.65 | 0.15 | 0.04 | 0.94 | met | 1.26 (1.00 to 1.59) | 203 |
| UVM | no | 80 | 1.8 | 0.95 | 0.83 | not met | 0.21 | n/a | n/a | n/a | n/a | n/a | n/a | n/a | n/a | n/a |
| KIRC | yes | 532 | 4.6 | 0.91 | 0.09 | met | 0.28 | 0.27 | 0.60 | 0.57 | 0.13 | 0.06 | 0.91 | met | 1.13 (0.96 to 1.33) | 174 |
| SKCM | yes | 103 | 2.8 | 0.89 | 0.65 | not met | 0.17 | 0.03 | 0.47 | 0.55 | 0.20 | 0.09 | 0.93 | met | 1.47 (0.98 to 2.20) | 27 |

| Project | Eligible | Tumours | Median non-feature % | lncRNA PC1 rho | Protein-coding PC1 rho | Rule A | Intronic minus intergenic | Plate R-squared of metric | Spread (MAD of log metric) | Largest observed module | Largest adjusted module | ARI | Largest-module eigengene r | Rule C | Metric HR, plate-stratified (95% CI) | Deaths |
|---|---|---|---|---|---|---|---|---|---|---|---|---|---|---|---|---|
| LUAD | yes | 517 | 4.4 | 0.88 | 0.20 | met | 0.22 | 0.39 | 0.39 | 0.44 | 0.16 | 0.21 | 0.91 | not met | 0.84 (0.68 to 1.04) | 176 |
| LUSC | yes | 500 | 4.1 | 0.86 | -0.19 | met | 0.22 | 0.40 | 0.48 | 0.25 | 0.18 | 0.36 | 0.72 | not met | 1.02 (0.87 to 1.21) | 205 |
| KIRP | yes | 290 | 4.1 | 0.83 | 0.71 | not met | 0.15 | 0.36 | 0.49 | 0.55 | 0.19 | 0.14 | 0.80 | met | 1.38 (1.00 to 1.89) | 39 |
| OV | yes | 426 | 5.2 | 0.82 | 0.52 | met | 0.24 | 0.26 | 0.41 | 0.29 | 0.15 | 0.28 | 0.94 | not met | 0.86 (0.74 to 1.01) | 261 |
| STAD | yes | 410 | 7.1 | 0.81 | 0.06 | met | 0.24 | 0.29 | 0.51 | 0.17 | 0.15 | 0.37 | 0.77 | not met | 1.11 (0.90 to 1.37) | 145 |
| READ | yes | 166 | 3.6 | 0.80 | 0.15 | met | 0.15 | 0.12 | 0.35 | 0.20 | 0.13 | 0.19 | 0.82 | not met | 0.73 (0.41 to 1.30) | 22 |
| PAAD | yes | 178 | 3.8 | 0.80 | 0.23 | met | 0.19 | 0.21 | 0.37 | 0.19 | 0.17 | 0.31 | 0.95 | not met | 1.02 (0.79 to 1.33) | 93 |
| COAD | yes | 446 | 3.4 | 0.79 | 0.07 | met | 0.17 | 0.17 | 0.34 | 0.18 | 0.16 | 0.26 | 0.81 | not met | 0.89 (0.70 to 1.14) | 91 |
| SARC | yes | 259 | 2.9 | 0.73 | 0.63 | not met | 0.06 | 0.13 | 0.39 | 0.24 | 0.15 | 0.30 | 0.74 | not met | 0.99 (0.80 to 1.23) | 98 |
| HNSC | yes | 520 | 2.5 | 0.72 | 0.48 | not met | 0.13 | 0.13 | 0.41 | 0.20 | 0.16 | 0.34 | 0.63 | not met | 0.82 (0.69 to 0.99) | 186 |
| LGG | yes | 516 | 4.5 | 0.68 | 0.05 | met | 0.11 | 0.26 | 0.29 | 0.18 | 0.10 | 0.32 | 0.73 | not met | 1.19 (0.94 to 1.51) | 117 |
| BLCA | yes | 406 | 3.5 | 0.68 | -0.54 | not met | 0.07 | 0.10 | 0.39 | 0.23 | 0.19 | 0.35 | 0.63 | not met | 0.90 (0.74 to 1.09) | 176 |
| BRCA | yes | 1094 | 3.9 | 0.61 | 0.44 | not met | 0.11 | 0.11 | 0.34 | 0.16 | 0.14 | 0.35 | 0.84 | not met | 0.78 (0.61 to 1.00) | 129 |
| UCS | no | 57 | 3.3 | 0.60 | -0.06 | met | 0.04 | n/a | n/a | n/a | n/a | n/a | n/a | n/a | n/a | n/a |

| Project | Eligible | Tumours | Median non-feature % | lncRNA PC1 rho | Protein-coding PC1 rho | Rule A | Intronic minus intergenic | Plate R-squared of metric | Spread (MAD of log metric) | Largest observed module | Largest adjusted module | ARI | Largest-module eigengene r | Rule C | Metric HR, plate-stratified (95% CI) | Deaths |
|---|---|---|---|---|---|---|---|---|---|---|---|---|---|---|---|---|
| DLBC | no | 48 | 3.1 | 0.58 | 0.66 | not met | 0.07 | 0.13 | n/a | n/a | n/a | n/a | n/a | n/a | n/a | n/a |
| THCA | yes | 505 | 3.3 | 0.57 | -0.28 | not met | 0.10 | 0.14 | 0.33 | 0.22 | 0.20 | 0.29 | 0.75 | not met | 1.19 (0.74 to 1.93) | 16 |
| ESCA | yes | 183 | 8.4 | 0.55 | 0.28 | not met | 0.25 | 0.32 | 0.45 | 0.40 | 0.28 | 0.31 | 0.86 | not met | 0.90 (0.68 to 1.19) | 64 |
| PCPG | yes | 179 | 3.3 | 0.52 | 0.42 | not met | 0.05 | 0.00 | 0.26 | 0.16 | 0.15 | 0.25 | 0.49 | not met | 1.76 (0.83 to 3.72) | 6 |
| KICH | no | 66 | 3.1 | 0.52 | -0.27 | not met | 0.05 | 0.02 | n/a | n/a | n/a | n/a | n/a | n/a | n/a | n/a |
| LAML | no | 63 | 27.7 | 0.50 | 0.33 | not met | 0.11 | 0.11 | n/a | n/a | n/a | n/a | n/a | n/a | n/a | n/a |
| CESC | yes | 304 | 3.5 | 0.49 | 0.22 | not met | 0.08 | 0.17 | 0.31 | 0.19 | 0.22 | 0.40 | 0.93 | not met | 0.90 (0.69 to 1.18) | 70 |
| UCEC | yes | 539 | 4.1 | 0.45 | 0.09 | not met | 0.07 | 0.16 | 0.39 | 0.20 | 0.11 | 0.52 | 0.50 | not met | 1.22 (1.00 to 1.49) | 91 |
| CHOL | no | 35 | 4.6 | 0.43 | 0.27 | not met | 0.05 | n/a | n/a | n/a | n/a | n/a | n/a | n/a | n/a | n/a |
| TGCT | yes | 150 | 4.8 | 0.42 | -0.39 | not met | 0.04 | 0.03 | 0.25 | 0.19 | 0.20 | 0.39 | -0.17 | not met | 0.68 (0.19 to 2.38) | 3 |
| ACC | no | 79 | 3.0 | 0.40 | 0.40 | not met | 0.06 | n/a | n/a | n/a | n/a | n/a | n/a | n/a | n/a | n/a |
| MESO | no | 87 | 3.2 | 0.36 | 0.09 | not met | 0.08 | 0.02 | n/a | n/a | n/a | n/a | n/a | n/a | n/a | n/a |
| PRAD | yes | 497 | 3.8 | 0.33 | 0.25 | not met | 0.09 | 0.29 | 0.31 | 0.19 | 0.18 | 0.26 | 0.32 | not met | 1.42 (0.68 to 2.97) | 10 |
| LIHC | yes | 371 | 2.3 | 0.26 | 0.39 | not met | 0.08 | 0.15 | 0.36 | 0.21 | 0.14 | 0.34 | 0.61 | not met | 1.00 (0.81 to 1.24) | 116 |
| THYM | yes | 120 | 3.8 | -0.42 | 0.57 | not met | 0.05 | 0.03 | 0.31 | 0.20 | 0.24 | 0.50 | 0.39 | not met | 0.75 (0.36 to 1.56) | 9 |

*(b) Solid-tissue normals with at least 20 retained libraries (descriptive)*

| Project | Normals | lncRNA PC1 rho | Protein-coding PC1 rho | Rule A thresholds met |
|---|---|---|---|---|
| LUAD | 59 | 0.95 | 0.66 | no |
| LUSC | 51 | 0.90 | 0.20 | yes |
| KICH | 25 | 0.84 | 0.86 | no |
| KIRP | 32 | 0.82 | 0.73 | no |
| STAD | 36 | 0.76 | -0.62 | no |
| BRCA | 113 | 0.71 | 0.49 | no |
| THCA | 59 | 0.61 | -0.03 | yes |
| UCEC | 35 | 0.51 | 0.45 | no |
| KIRC | 72 | 0.33 | 0.29 | no |
| PRAD | 52 | 0.26 | 0.12 | no |
| HNSC | 44 | 0.20 | 0.24 | no |
| COAD | 41 | 0.14 | -0.10 | no |
| LIHC | 50 | -0.16 | 0.37 | no |

*(c) Random-effects meta-analysis of the metric's hazard ratio for overall survival*

| Model | Cancer types | Pooled HR (95% CI) | p | I-squared (%) | Prediction interval | HR above 1 | p below 0.05 |
|---|---|---|---|---|---|---|---|
| primary (plate stratum) | 20 | 1.00 (0.93 to 1.08) | 0.98 | 50 | 0.79 to 1.26 | 10 | 4 |
| secondary (no plate stratum) | 20 | 1.01 (0.94 to 1.08) | 0.83 | 64 | 0.79 to 1.29 | 10 | 4 |

*(d) Post hoc dose-response: within-cohort spread of the metric against the rule quantities, Spearman across 25 cancer types*

| Spread measure | Outcome | rho | p | FDR |
|---|---|---|---|---|
| mad_log_metric | absolute rho of the lncRNA PC1 with the metric | 0.73 | 3.1e-05 | 0.00061 |
| mad_log_metric | lncRNA minus protein-coding absolute rho | 0.32 | 0.12 | 0.14 |
| mad_log_metric | largest observed module, share of genes | 0.69 | 0.00016 | 0.0016 |
| mad_log_metric | adjusted Rand index, observed vs adjusted | -0.33 | 0.1 | 0.13 |
| mad_log_metric | metric variance explained by plate | 0.53 | 0.0062 | 0.014 |
| cv_metric | absolute rho of the lncRNA PC1 with the metric | 0.64 | 0.00053 | 0.0025 |
| cv_metric | lncRNA minus protein-coding absolute rho | 0.39 | 0.057 | 0.081 |
| cv_metric | largest observed module, share of genes | 0.50 | 0.01 | 0.017 |

| Spread measure | Outcome | rho | p | FDR |
|---|---|---|---|---|
| cv_metric | adjusted Rand index, observed vs adjusted | -0.20 | 0.33 | 0.37 |
| cv_metric | metric variance explained by plate | 0.53 | 0.0062 | 0.014 |
| iqr_metric | absolute rho of the lncRNA PC1 with the metric | 0.64 | 0.00063 | 0.0025 |
| iqr_metric | lncRNA minus protein-coding absolute rho | 0.41 | 0.041 | 0.063 |
| iqr_metric | largest observed module, share of genes | 0.53 | 0.0069 | 0.014 |
| iqr_metric | adjusted Rand index, observed vs adjusted | -0.20 | 0.35 | 0.37 |
| iqr_metric | metric variance explained by plate | 0.65 | 0.00046 | 0.0025 |
| sd_metric | absolute rho of the lncRNA PC1 with the metric | 0.63 | 0.00081 | 0.0027 |
| sd_metric | lncRNA minus protein-coding absolute rho | 0.52 | 0.0076 | 0.014 |
| sd_metric | largest observed module, share of genes | 0.38 | 0.06 | 0.081 |
| sd_metric | adjusted Rand index, observed vs adjusted | -0.09 | 0.66 | 0.66 |
| sd_metric | metric variance explained by plate | 0.62 | 0.001 | 0.0029 |

*Note.* Decision rules were fixed in the pipeline configuration, committed and hashed (`38_decision_rules_lock.json`) before any non-renal project was analysed. We had analysed the renal projects during discovery, so we set the thresholds with those results known. Excluding TCGA-KIRC and TCGA-KIRP, Rule A is met in 8 of 23 projects and Rule C in 2 of 23. The rule letters correspond to the three measurements of the analysis plan. The second (B), the gene-level and positional-class comparison, had no decision threshold and is reported descriptively. Panel (d) was specified afterwards and is exploratory. Rule A requires the lncRNA PC1 to correlate with the non-feature fraction at an absolute rho of at least 0.5 and at least 0.3 above the protein-coding PC1. Rule C requires the largest observed lncRNA module to hold at least 40% of genes and its eigengene to correlate with the metric at an absolute r of at least 0.7, with an adjusted Rand index below 0.2. Overall survival from the TCGA Clinical Data Resource, censored at ten years, is adjusted for age, sex (omitted in single-sex cancers) and stage, and stratified by sequencing plate, pooling plates with fewer than 10 patients. Stage is AJCC pathologic where recorded for at least 70% of patients, otherwise clinical, otherwise omitted.

Projects with at least 20 deaths enter the meta-analysis. Each project uses its own expressed genes. Source tables: `38_pancancer_cohorts.tsv`, `39_pancancer_axis.tsv`, `39_pancancer_class_gradient.tsv`, `40_pancancer_network.tsv`, `41_pancancer_metric_survival.tsv`, `41_pancancer_metric_meta.tsv`, `43_pancancer_dose_response.tsv`, `43_pancancer_dose_response_projects.tsv`.

#### Supplementary Table 7. The lncRNA axis and the non-feature read fraction: extended models, batch structure and biospecimen correlates

##### (a) Nested Cox models for the leading lncRNA axis, hazard ratio per standard deviation of the axis, overall survival

| Model | HR | 95% CI | p | C | Proportional-hazards p |
|---|---|---|---|---|---|
| Axis alone | 1.31 | 1.12 to 1.53 | 5.3e-04 | 0.555 | 0.14 |
| + age, sex, T, N, M1, ordinal grade | 1.34 | 1.15 to 1.57 | 1.9e-04 | 0.773 | 0.11 |
| + ESTIMATE stromal and immune scores | 1.36 | 1.16 to 1.60 | 2.3e-04 | 0.773 | 0.10 |
| + non-feature fraction, multimapping fraction, log10 assigned reads | 1.08 | 0.81 to 1.43 | 0.61 | 0.777 | 0.093 |
| + RIN | 1.34 | 1.13 to 1.58 | 5.5e-04 | 0.773 | 0.089 |
| + RIN + slide necrosis | 1.39 | 1.17 to 1.64 | 1.5e-04 | 0.777 | 0.11 |
| + RIN + slide necrosis + the three STAR metrics | 1.12 | 0.84 to 1.50 | 0.44 | 0.779 | 0.099 |
| + RIN + slide necrosis + the three STAR metrics, stratified by sequencing plate | 0.90 | 0.62 to 1.29 | 0.55 | 0.760 | 0.57 |

*Note.* TCGA-KIRC, 503 patients and 163 events in every row, those with a recorded RNA integrity number and complete covariates. Source: `23_axis_nested_extended.tsv`. The association survives adding RIN and slide necrosis, falls only when the three STAR metrics enter, and disappears with stratification by sequencing plate.

**(b) The non-feature read fraction as the exposure, hazard ratio per standard deviation, overall survival**

| Model | Linear HR (95% CI) | p | log10 HR (95% CI) | p |
|---|---|---|---|---|
| Non-feature fraction alone | 1.28 (1.11 to 1.47) | 5.9e-04 | 1.32 (1.13 to 1.54) | 5.1e-04 |
| + age, sex, T, N, M1, ordinal grade | 1.36 (1.18 to 1.56) | 2.0e-05 | 1.41 (1.20 to 1.66) | 2.9e-05 |
| + ESTIMATE stromal and immune scores | 1.37 (1.18 to 1.58) | 2.0e-05 | 1.43 (1.21 to 1.69) | 2.4e-05 |
| + multimapping fraction, log10 assigned reads | 1.36 (1.17 to 1.58) | 4.5e-05 | 1.42 (1.20 to 1.68) | 4.5e-05 |
| + ESTIMATE, stratified by tissue source site | 1.44 (1.22 to 1.69) | 1.3e-05 | 1.48 (1.24 to 1.78) | 2.0e-05 |
| + RIN | 1.35 (1.17 to 1.56) | 6.0e-05 | 1.41 (1.20 to 1.67) | 4.6e-05 |
| + RIN + slide necrosis | 1.37 (1.19 to 1.59) | 2.5e-05 | 1.44 (1.22 to 1.71) | 1.9e-05 |
| + RIN + slide necrosis + multimapping, log10 assigned reads | 1.37 (1.18 to 1.59) | 4.7e-05 | 1.44 (1.21 to 1.70) | 3.0e-05 |
| + RIN + slide necrosis + multimapping, log10 assigned reads, stratified by plate | 1.23 (1.02 to 1.50) | 0.035 | 1.23 (1.00 to 1.51) | 0.049 |

*Note.* TCGA-KIRC, 503 patients and 163 events in every row, restricted to patients with a recorded RNA integrity number so that the RIN and plate rows are nested. Source: `23_noFeature_exposure_extended.tsv`. The headline figures in the main text use the wider complete-case set of 520 patients and 172 events (`07_noFeature_as_exposure.tsv`, reproduced in Supplementary Table 8c), so confidence limits differ slightly for the same named model. The exposure is right-skewed, so every model is reported on both scales. Unlike the axis in panel (a), the exposure survives stratification by sequencing plate.

**(c) Spearman correlations of the metric and of the axis with biospecimen measures**

| Measure | Non-feature fraction, rho (p, n) | lncRNA axis, rho (p, n) |
|---|---|---|
| RNA integrity number | -0.25 (9.4e-09, 527) | -0.27 (3.4e-10, 510) |
| A260/A280 ratio | 0.18 (3.6e-05, 528) | 0.23 (1.1e-07, 511) |
| Slide percent necrosis, mean over slides | -0.05 (0.28, 528) | -0.09 (0.042, 511) |
| Slide percent tumour nuclei, mean over slides | 0.03 (0.45, 528) | 0.10 (0.019, 511) |
| Slide percent stromal cells, mean over slides | -0.10 (0.021, 524) | -0.12 (0.0094, 507) |
| ESTIMATE stromal score | -0.10 (0.028, 528) | -0.16 (2.0e-04, 511) |
| ESTIMATE immune score | 0.04 (0.39, 528) | 0.04 (0.40, 511) |

*Note.* TCGA-KIRC only. No equivalent metadata exist for CPTAC-3, TCGA-KIRP or TCGA-KICH. Source: `23_axis_vs_biospecimen.tsv`. In the same file, the multimapping fraction correlates with RIN at rho -0.19 (p 1.3e-05), with slide necrosis at 0.12 (p 0.0078) and with the ESTIMATE stromal score at -0.21 (p 1.5e-06), and log10 assigned reads correlates with RIN at 0.12 (p 0.0051). RIN was recorded for 527 of 528 patients. One of those 527, recorded as 0, is a coded missing value, leaving 526 usable, although the regression of panel (e) uses all 527. Slide necrosis, with a median of 0 and an interquartile range of 0 to 0, is uninformative.

**(d) Variance in the metric and in the axis explained by batch variables**

| Quantity | Factor | Levels modelled | Variance explained (%) | ANOVA p | Kruskal-Wallis p | n |
|---|---|---|---|---|---|---|
| Non-feature fraction | Sequencing plate | 15 | 26.6 | 6.5e-27 | 4.0e-28 | 528 |
| Non-feature fraction | BCR batch | 15 | 24.6 | 4.2e-24 | 1.0e-25 | 528 |
| Non-feature fraction | Tissue source site | 11 | 11.1 | 2.3e-09 | 8.3e-11 | 528 |
| lncRNA axis | Sequencing plate | 15 | 26.2 | 2.1e-25 | 9.3e-22 | 511 |
| lncRNA axis | BCR batch | 15 | 26.0 | 4.3e-25 | 1.6e-21 | 511 |
| lncRNA axis | Tissue source site | 11 | 11.6 | 1.6e-09 | 3.1e-09 | 511 |

*Note.* One-way analysis of variance in TCGA-KIRC. Source: `23_batch_variance.tsv`. Nineteen sequencing plates and 15 BCR batches are recorded. Plate, batch and site levels with fewer than ten patients are pooled into one "other" stratum, giving 15, 15 and 11 modelled levels.

Across the 14 plates with at least ten patients the median non-feature fraction ranges from 2.6% to 7.1%.

**(e) Determinants of the non-feature read fraction**

| Term | df | Variance explained alone (%) | Partial variance explained (%) | p alone | p partial |
|---|---|---|---|---|---|
| RNA integrity number | 1 | 6.9 | 5.4 | 9.9e-10 | 1.5e-07 |
| Sequencing plate | 14 | 26.6 | 24.0 | 7.4e-27 | 9.4e-23 |
| Tissue source site | 10 | 11.1 | 7.8 | 2.3e-09 | 1.0e-05 |
| RIN + plate + site jointly | 25 | 36.2 | n/a | 4.1e-35 | n/a |

*Note.* TCGA-KIRC, 527 patients with a recorded RNA integrity number. Source: `23_noFeature_determinants.tsv`. Partial variance explained is conditional on the other two terms. Plate, batch and RIN proxy unrecorded extraction, library-preparation and run variables, so 36.2% is a lower bound on the technical share.

#### Supplementary Table 8. Disease-specific and progression-free endpoints

**(a) Endpoint coverage in the discovery cohort**

| Endpoint | Patients scoreable | Events after ten-year censoring | Unscoreable | Median follow-up (days) | Events per variable at 12 parameters |
|---|---|---|---|---|---|
| Overall survival, pipeline | 528 | 173 | 0 | 1,657 | 14.4 |
| Overall survival, Clinical Data Resource | 528 | 173 | 0 | 1,666 | 14.4 |
| Disease-specific survival | 517 | 109 | 11 | 1,495 | 9.1 |
| Progression-free interval | 526 | 158 | 2 | 1,471 | 13.2 |
| Disease-free interval | 114 | 14 | 414 | 1,525 | 1.2, not analysed |

*Note.* Source: `30_cdr_coverage_and_os_concordance.tsv`. The Clinical Data Resource overall-survival endpoint matches the pipeline endpoint on all 528 event indicators, with times correlated at Spearman 0.998. Six patients differ in time only, 5 from benign data-freeze differences and 1 flagged as not explained by a later freeze. All 11 patients unscorable for disease-specific survival are pipeline deaths, and both unscorable for the progression-free interval carry an event. Event-indicator dependence (phi) is 0.77 between overall and disease-specific survival, 0.52 between overall survival and the progression-free interval, and 0.79 between the latter two. The four endpoints are therefore not independent, and adjustment across them is crude in both directions. The Clinical Data Resource labels disease-specific survival approximate for this disease because cause of death is unverified. No patient is marked redacted.

**(b) The five prognostic modules under three endpoints, principal specification**

| Module | Overall survival | Disease-specific survival | Progression-free interval |
|---|---|---|---|
| Protein-coding green | 0.62 (0.53 to 0.74) | 0.49 (0.39 to 0.61) | 0.70 (0.59 to 0.83) |
| Protein-coding purple | 1.28 (1.09 to 1.50) | 1.49 (1.21 to 1.84) | 1.15 (0.96 to 1.38), p 0.14 |
| lncRNA blue | 0.71 (0.60 to 0.83) | 0.61 (0.49 to 0.75) | 0.80 (0.68 to 0.96) |
| lncRNA greenyellow | 1.35 (1.15 to 1.60) | 1.50 (1.22 to 1.85) | 1.21 (1.01 to 1.45), FDR 0.065 |
| lncRNA turquoise | 1.24 (1.06 to 1.45) | 1.20 (1.00 to 1.44), p 0.054 | 1.03 (0.88 to 1.22), p 0.70 |

*Note.* Hazard ratio per standard deviation of the module eigengene. Sample sizes and events for protein-coding modules are 502 and 165 (overall survival), 491 and 103 (disease-specific survival) and 500 and 151 (progression-free interval), and for lncRNA modules 504 and 163, 493 and 102, and 502 and 149. Source: `30_endpoint_modules.tsv`. The false discovery rate is controlled across the five modules within each endpoint, not across the 20 module-by-endpoint tests. Unmarked cells are below 0.05. Sixteen of the 20 tests reach a 5% false discovery rate. The Clinical Data Resource overall-survival column agrees with the pipeline column to within 0.01 throughout.

**(c) The non-feature read fraction as exposure, by endpoint**

| Endpoint | n / events | Alone | + clinical model | log10 scale, + clinical model |
|---|---|---|---|---|
| Overall survival, pipeline | 520 / 172 | 1.28 (1.13 to 1.46), p 1.2e-04 | 1.36 (1.20 to 1.55), p 3.0e-06 | 1.44 (1.23 to 1.68) |
| Overall survival, Clinical Data Resource | 520 / 172 | 1.28 (1.13 to 1.46) | 1.36 (1.19 to 1.54), p 4.0e-06 | 1.43 (1.22 to 1.67) |
| Disease-specific survival | 509 / 109 | 1.20 (1.02 to 1.42), p 0.032 | 1.31 (1.10 to 1.55), p 0.0020 | 1.35 (1.11 to 1.64) |
| Progression-free interval | 518 / 157 | 1.03 (0.88 to 1.20), p 0.71 | 1.11 (0.95 to 1.30), p 0.19 | 1.07 (0.91 to 1.26), p 0.44 |

**(d) The projected lncRNA axis, nested sequence, by endpoint**

| Endpoint | n / events | Alone | + clinical model | + ESTIMATE | + the three STAR metrics |
|---|---|---|---|---|---|
| Overall survival, pipeline | 504 / 163 | 1.31 (1.13 to 1.53) | 1.34 (1.15 to 1.57) | 1.36 (1.16 to 1.60) | 1.08 (0.81 to 1.43), p 0.61 |
| Overall survival, Clinical Data Resource | 504 / 163 | 1.31 | 1.33 | 1.35 | 1.07 (0.81 to 1.41), p 0.66 |
| Disease-specific survival | 493 / 102 | 1.20 (0.98 to 1.45), p 0.073 | 1.28 (1.04 to 1.56), p 0.017 | 1.30 (1.05 to 1.61) | 0.99 (0.70 to 1.39), p 0.94 |
| Progression-free interval | 502 / 149 | 0.97 (0.82 to 1.15), p 0.73 | 1.01 (0.85 to 1.19), p 0.92 | 1.09 (0.91 to 1.31) | 0.91 (0.67 to 1.24), p 0.57 |

*Note for panels (c) and (d).* Source: `30_endpoint_exposure.tsv`, `30_endpoint_axis_nested.tsv`. The metric and the axis are prognostic for overall and disease-specific survival but not for progression. The axis's variance-inflation factor reaches 4.5 once the three STAR metrics enter, so its collapse to 1.08 reflects collinearity as much as adjustment.

**(e) Cross-validated increment in the concordance index from adding the 29 module eigengenes to the clinical model**

| Endpoint | n | Events | C, clinical | C, clinical plus eigengenes | Mean over repeats | Bootstrap increment | 95% CI | p | BH over the four endpoints |
|---|---|---|---|---|---|---|---|---|---|
| Overall survival, pipeline | 493 | 159 | 0.7551 | 0.7596 | 0.0045 | 0.0081 | -0.0077 to 0.0246 | 0.33 | 0.33 |

| Endpoint | n | Events | C, clinical | C, clinical plus eigengenes | Mean over repeats | Bootstrap increment | 95% CI | p | BH over the four endpoints |
|---|---|---|---|---|---|---|---|---|---|
| Overall survival, Clinical Data Resource | 493 | 159 | 0.7540 | 0.7588 | 0.0048 | 0.0087 | -0.0075 to 0.0258 | 0.30 | 0.33 |
| Disease-specific survival | 482 | 99 | 0.8230 | 0.8396 | 0.0166 | 0.0227 | 0.0037 to 0.0448 | 0.017 | 0.068 |
| Progression-free interval | 491 | 146 | 0.8007 | 0.8123 | 0.0116 | 0.0138 | 0.0011 to 0.0272 | 0.036 | 0.072 |

*Note.* Ten repeats of ten-fold cross-validation, with module definitions fixed on the full cohort and the penalty tuned inside each training fold. Source: `30_endpoint_cv_increment.tsv`. The mean over repeats has no interval. The confidence interval and p value belong only to the bootstrap increment, the increment of the repeat-averaged out-of-fold predictor. Neither positive endpoint survives Benjamini-Hochberg adjustment over the four endpoints. Across 200 bootstrap seeds the lower bound ranges from 0.0010 to 0.0046 for disease-specific survival, excluding zero in 200 of 200, and from 0.0004 to 0.0025 for the progression-free interval, also 200 of 200, against 0 of 200 for overall survival.

**(f) The locked model of the main analysis re-scored against each endpoint, apparent and in sample**

| Endpoint | Comparator | n | Events | Modules retained | C comparator | C combined | Increment | 95% CI | p | BH over the eight rows |
|---|---|---|---|---|---|---|---|---|---|---|
| Overall survival, pipeline | clinical | 493 | 159 | 9 | 0.7662 | 0.7867 | 0.0205 | 0.0022 to 0.0400 | 0.026 | 0.052 |
| Overall survival, Clinical Data Resource | clinical | 493 | 159 | 9 | 0.7649 | 0.7858 | 0.0209 | 0.0024 to 0.0407 | 0.024 | 0.052 |
| Disease-specific survival | clinical | 482 | 99 | 9 | 0.8218 | 0.8475 | 0.0257 | 0.0059 to 0.0488 | 0.010 | 0.052 |

| Endpoint | Comparator | n | Events | Modules retained | C comparator | C combined | Increment | 95% CI | p | BH over the eight rows |
|---|---|---|---|---|---|---|---|---|---|---|
| Progression-free interval | clinical | 491 | 146 | 9 | 0.7924 | 0.8021 | 0.0098 | -0.0123 to 0.0319 | 0.40 | 0.45 |
| Overall survival, pipeline | augmented | 493 | 159 | 5 | 0.7750 | 0.7916 | 0.0165 | -0.0005 to 0.0347 | 0.056 | 0.075 |
| Overall survival, Clinical Data Resource | augmented | 493 | 159 | 5 | 0.7739 | 0.7910 | 0.0171 | -0.0003 to 0.0356 | 0.052 | 0.075 |
| Disease-specific survival | augmented | 482 | 99 | 5 | 0.8218 | 0.8458 | 0.0240 | 0.0051 to 0.0475 | 0.016 | 0.052 |
| Progression-free interval | augmented | 491 | 146 | 5 | 0.7850 | 0.7925 | 0.0074 | -0.0139 to 0.0293 | 0.52 | 0.52 |

*Note.* Source: `30_endpoint_locked_model.tsv`. Every row is apparent. The coefficients were fitted on the overall survival of exactly these patients, whose disease-specific deaths are a subset of the deaths used in fitting, as the file's own evaluation column states. On the same 482 patients the apparent overall-survival increment is 0.0239, essentially identical to the disease-specific figure, so the locked-model rows carry no information about differences between endpoints. The cross-validated rows of panel (e) do.

**(g) Rebuilding both networks inside every training fold, disease-specific survival**

| Analysis | Endpoint | n | Events | C clinical | C combined | Increment | SD | Range | Assignments |
|---|---|---|---|---|---|---|---|---|---|
| Ten by ten cross-validation, modules fixed | disease-specific | 482 | 99 | 0.8265 | 0.8493 | 0.0227 | 0.0050 | 0.0037 to 0.0448 (bootstrap) | 10 repeats |
| Three folds by five assignments, modules fixed | disease-specific | 482 | 99 | 0.8141 | 0.8271 | 0.0129 | 0.0124 | -0.0005 to 0.0283 | 5 |
| Three folds by five assignments, modules rebuilt inside each fold | disease-specific | 482 | 99 | 0.8141 | 0.8268 | 0.0126 | 0.0083 | 0.0000 to 0.0232 | 5 |

| Analysis | Endpoint | n | Events | C clinical | C combined | Increment | SD | Range | Assignments |
|---|---|---|---|---|---|---|---|---|---|
| Three folds by five assignments, modules fixed | overall survival | 482 | 148 | 0.7403 | 0.7498 | 0.0095 | 0.0096 | -0.0024 to 0.0229 | 5 |
| Three folds by five assignments, modules rebuilt inside each fold | overall survival | 482 | 148 | 0.7403 | 0.7478 | 0.0075 | 0.0075 | -0.0046 to 0.0143 | 5 |

*Note.* Source: `30_endpoint_foldwise.tsv`. The fixed-module three-fold arm shares folds and inner splits with the rebuilt arm, so the two differ only in module definition. Rebuilding both networks and all loadings inside every training fold costs 0.0003 for disease-specific and 0.0020 for overall survival, so the positive disease-specific increment is not an artefact of defining modules on the full cohort. The matched fixed-module arm shows that the drop from 0.0227 to about 0.013 reflects the coarser three-fold cross-validation (two-thirds of the data for training, no averaging over repeats). Rebuilt training-fold networks resolve 11 to 19 protein-coding and 11 to 17 lncRNA modules, giving 24 to 35 module eigengenes against the 29 fixed ones. The range across five fold assignments is not a confidence interval. CPTAC-3 lacks the disease-specific and progression-free endpoints, so neither positive result is externally replicated.

#### Supplementary Table 9. What the non-feature read fraction correlates with, and what attenuates it

##### (a) Candidate correlates of the metric and of the axis

| Candidate | Matrix | rho with the metric (FDR) | rho with the lncRNA axis (FDR) | Interpretable as biology |
|---|---|---|---|---|
| Protein-coding red eigengene | observed | 0.84 (3.5e-133) | 0.92 (1.5e-204) | no |
| Protein-coding red eigengene | residualised | 0.20 (1.2e-04) | 0.42 (6.8e-22) | yes |
| lncRNA share of total FPKM | observed | 0.73 (7.1e-89) | 0.74 (7.7e-89) | no |

| Candidate | Matrix | rho with the metric (FDR) | rho with the lncRNA axis (FDR) | Interpretable as biology |
|---|---|---|---|---|
| Protein-coding pink eigengene | observed | 0.32 (2.8e-12) | 0.22 (8.3e-06) | no |
| Protein-coding tan eigengene | observed | -0.20 (1.7e-05) | n/a | no |
| Hypoxia metagene, Buffa 2010[58] | observed | -0.14 (0.0056) | -0.12 (0.019) | yes |
| Proliferation score, 10 genes | observed | -0.01 (0.91) | -0.10 (0.031) | yes |
| ESTIMATE stromal score | n/a | -0.10 (0.056) | -0.16 (8.1e-04) | yes |
| ESTIMATE immune score | n/a | 0.04 (0.60) | 0.04 (0.40) | yes |
| ESTIMATE score | n/a | -0.02 (0.67) | -0.06 (0.22) | yes |
| Slide percent tumour nuclei | n/a | 0.03 (0.60) | 0.10 (0.031) | yes |
| Slide percent stromal cells | n/a | -0.10 (0.055) | -0.12 (0.019) | yes |

*Note.* TCGA-KIRC. Eigengene rows use 506 patients against the metric and 497 against the axis, other rows 528 or 511. Source: `27_metric_biology_correlations.tsv`. Benjamini-Hochberg families are declared per matrix. Eigengenes of the observed expression matrix cannot be read as biology, because the metric was measured on that matrix. All hypoxia sensitivity variants run negative: rho -0.18 (p 3.6e-05) for the metagene restricted to genes with a median FPKM of at least 1, and -0.14 and -0.15 for the two alternative tubulin mappings. A higher non-feature fraction thus goes with a lower hypoxia score, which is evidence against the hypoxia reading.

**(b) Which protein-coding genes carry the metric's signal**

| Module | Genes | Median per-gene rho with the metric | Fraction above 0.3 in absolute value | Eigengene rho, observed | Eigengene rho, residualised | Top GO term (adjusted p) |
|---|---|---|---|---|---|---|
| red | 473 | 0.68 | 94.5% | 0.84 | 0.20 | RNA splicing (4.5e-15) |
| pink | 270 | 0.22 | 37.0% | 0.32 | 0.05 | none |
| tan | 152 | -0.16 | 2.6% | -0.20 | -0.03 | none |

| Module | Genes | Median per-gene rho with the metric | Fraction above 0.3 in absolute value | Eigengene rho, observed | Eigengene rho, residualised | Top GO term (adjusted p) |
|---|---|---|---|---|---|---|
| salmon | 63 | 0.06 | 11.1% | 0.15 | 0.01 | none |
| yellow | 986 | 0.05 | 14.2% | 0.13 | 0.00 | none |
| magenta | 227 | 0.03 | 17.6% | 0.12 | 0.02 | none |
| blue | 994 | 0.04 | 12.1% | 0.10 | -0.01 | none |
| purple | 216 | 0.01 | 15.3% | 0.07 | 0.01 | chromosome segregation (1.5e-44) |
| greenyellow | 164 | 0.01 | 16.5% | 0.06 | 0.05 | none |
| turquoise | 2,405 | 0.01 | 11.7% | 0.06 | 0.04 | none |
| black | 323 | -0.04 | 1.9% | -0.04 | -0.03 | none |
| green | 670 | -0.01 | 7.5% | 0.04 | 0.02 | small molecule catabolic process (4.1e-38) |
| brown | 988 | -0.05 | 15.8% | -0.01 | -0.01 | none |

*Note.* Per-gene Spearman correlation with the metric on the observed matrix, 506 samples. Source: `27_module_gene_level_metric_correlation.tsv`, GO terms from `06_GO_enrichment_top10_per_module.tsv`. The only protein-coding module tracking the metric is the RNA-splicing and mRNA-processing module, as a shift towards unspliced and intronic reads predicts. This identifies the affected genes, not how much of the metric is biological.

**(c) Variance in the metric explained by candidate blocks, with negative controls**

| Block | Kind | Interpretable as biology | df | Variance explained (%) | Partial variance explained given the other block (%) |
|---|---|---|---|---|---|
| Annotated biology: hypoxia, proliferation, ESTIMATE stromal and immune, slide tumour nuclei | biological | yes | 5 | 6.1 | 5.0 |
| Technical: RIN, sequencing plate, tissue source site | technical | no, but interpretable as technical | 24 | 36.8 | 36.0 |

| Block | Kind | Interpretable as biology | df | Variance explained (%) | Partial variance explained given the other block (%) |
|---|---|---|---|---|---|
| Observed expression space, all terms | upper bound, not identifiable as biology | no | 18 | 77.9 | 71.5 |
| 13 observed protein-coding eigengenes alone | upper bound | no | 13 | 77.3 | 71.1 |
| Observed protein-coding red eigengene alone | upper bound | no | 1 | 64.7 | 60.4 |
| Negative control: 13 eigengenes of random gene sets of the same sizes, observed matrix | negative control | no | 13 | 61.0 | 54.3 |
| Negative control: 13 leading principal components of the observed matrix | negative control | no | 13 | 80.9 | 76.6 |
| 13 residualised protein-coding eigengenes | residualised expression space | no | 13 | 0.01 | 6.9 |
| Negative control: 13 random gene-set eigengenes, residualised matrix | negative control | no | 13 | 0.01 | 2.9 |
| Annotated biology plus observed eigengenes excluding red | upper bound | no | 17 | 40.6 | 41.4 |

*Note.* TCGA-KIRC, 501 complete cases, linear scale. Source: `27_noFeature_variance_partition.tsv`. Only the annotated-biology and technical blocks are interpretable shares. Values are given to one decimal, or to two for the two residualised rows, which would otherwise round to zero. Thirteen random gene sets with no biological content reach 61.0% and thirteen unsupervised components 80.9%, more than the block containing the real eigengenes. On the residualised matrix the same eigengenes explain 0.01%. These rows measure how much of the metric the observed expression matrix recovers, a property of the shared alignment and FPKM normalisation, not a biological cause. The random-gene-set row is one draw. Over 20 draws the mean is 60.8% (range 51.7% to 70.8%). The 77.9% and 77.3% rows are references for the negative controls, not estimates of any biological share.

Plate, batch and RIN only proxy unrecorded pre-analytical variables, so 36.8% is a lower bound on the technical share and 6.1% an upper bound on the annotated-biological share, as far as these data can measure.

**(d) What attenuates the hazard of the metric, and what does not**

| Model added to the base | Interpretable | HR | 95% CI | p | Standard-error inflation |
|---|---|---|---|---|---|
| Base: metric per SD + clinical model | n/a | 1.37 | 1.20 to 1.57 | 4.8e-06 | 1.00 |
| + hypoxia metagene | yes | 1.38 | 1.20 to 1.59 | 4.2e-06 | 1.02 |
| + proliferation score | yes | 1.37 | 1.19 to 1.56 | 5.8e-06 | 1.00 |
| + ESTIMATE stromal and immune | yes | 1.38 | 1.20 to 1.58 | 6.4e-06 | 1.03 |
| + slide percent tumour nuclei | yes | 1.37 | 1.20 to 1.57 | 6.0e-06 | 1.01 |
| + RIN | yes | 1.35 | 1.18 to 1.55 | 1.7e-05 | 1.02 |
| + sequencing plate stratum | yes | 1.24 | 1.04 to 1.48 | 0.018 | 1.32 |
| + residualised protein-coding red eigengene | yes | 1.40 | 1.21 to 1.63 | 5.9e-06 | 1.09 |
| + all 13 residualised eigengenes | yes | 1.39 | 1.19 to 1.62 | 2.3e-05 | 1.14 |
| + every interpretable candidate jointly, residualised | yes | 1.23 | 1.01 to 1.50 | 0.043 | 1.49 |
| + observed protein-coding red eigengene | no, collinearity diagnostic | 1.14 | 0.89 to 1.45 | 0.31 | 1.83 |
| + all 13 observed eigengenes | no, collinearity diagnostic | 1.13 | 0.85 to 1.49 | 0.40 | 2.07 |
| + lncRNA share of total FPKM | no, collinearity diagnostic | 1.27 | 1.04 to 1.54 | 0.018 | 1.45 |

*Note.* TCGA-KIRC, 501 patients and 165 events, overall survival, hazard ratio per standard deviation of the metric on the linear scale. Source: `27_metric_attenuation.tsv`. Rows marked as collinearity diagnostics add a covariate that is largely the exposure itself, measured through the expression matrix. For the lncRNA axis (492 patients, 159 events), the same sequence gives a base hazard ratio of 1.36 (1.17 to 1.59), falling to 1.13 (0.94 to 1.36, p 0.18) with stratification by plate and rising to 1.46 (1.22 to 1.75) with all 13 residualised eigengenes.

**(e) Collinearity diagnostics for the two rows that appear to attenuate**

| Candidate | Pearson r with the exposure | Variance inflation | Correlation of the orthogonal part with the exposure | Synthetic-proxy null, median HR (2.5 to 97.5 centile) | Contains the observed value |
|---|---|---|---|---|---|
| Observed protein-coding red eigengene | 0.80 | 2.89 | 0.99 | 1.37 (1.08 to 1.71) | yes, observed 1.14 |
| lncRNA share of total FPKM | 0.72 | 2.14 | n/a | 1.37 (1.15 to 1.62) | yes, observed 1.27 |

*Note.* TCGA-KIRC, 501 patients and 165 events, 300 synthetic draws. Source: `27_collinearity_diagnostics.tsv`. A synthetic covariate correlating 0.80 with the metric but carrying no survival information reproduces the apparent attenuation. The five residualised eigengenes, by comparison, correlate with the exposure between -0.003 and 0.003, with variance inflation of at most 1.02.

**(f) The metric against clinical, batch and biospecimen strata**

| Variable | Test | n | Summary | p |
|---|---|---|---|---|
| RIN tertile | Kruskal-Wallis | 527 | medians 6.0%, 4.3%, 4.1% | 4.7e-07 |
| Sequencing plate, 15 pooled levels | Kruskal-Wallis | 528 | medians 2.6% to 7.1% | 4.0e-28 |
| BCR batch, 14 levels | Kruskal-Wallis | 527 | medians 2.6% to 6.9% | 3.5e-26 |
| Sex | Kruskal-Wallis | 528 | 5.0% in women, 4.5% in men | 0.21 |
| Age tertile | Kruskal-Wallis | 528 | 4.7%, 4.6%, 4.3% | 0.74 |
| Stage group | Wilcoxon | 525 | 4.5% in stage I to II, 5.1% in III to IV | 0.35 |
| Grade group | Wilcoxon | 520 | 4.5% in G1 to G2, 4.7% in G3 to G4 | 0.97 |
| M1 | Wilcoxon | 528 | 4.6% M0, 4.8% M1 | 0.71 |
| T category | Spearman | 528 | rho 0.06 | 0.21 |
| Age | Spearman | 528 | rho -0.04 | 0.41 |
| Ordinal grade | Spearman | 520 | rho -0.03 | 0.57 |

*Note.* TCGA-KIRC. Source: `27_noFeature_reverse_check.tsv`, which reuses the clinical rows of `07_noFeature_vs_clinical.tsv` unchanged. The smallest p for any clinical variable is 0.21, whereas RIN, plate and batch each associate with the metric at p below 1e-06. The batch row drops the single-patient "other" stratum kept in Supplementary Table 7(d), so it has 14 levels on 527 patients, not 15 on 528.

#### Supplementary Table 10. Single-cell ground truth for the correction, TCGA-KIRC

*(a) Agreement of the bulk tumour-minus-normal difference with the single-cell malignant-versus-normal-epithelium fold change*

| Contrast | Correction | lncRNAs | Single-cell significant | rho observed | rho corrected | Change (95% interval) | Sign agreement observed | Sign agreement corrected |
|---|---|---|---|---|---|---|---|---|
| all pairs | tumour fitted | 507 | 213 | 0.775 | 0.771 | -0.004 (-0.008 to -0.001) | 92.5% | 92.0% |
| all pairs | joint fitted | 507 | 213 | 0.775 | 0.770 | -0.005 (-0.009 to -0.002) | 92.5% | 92.0% |
| shared-plate pairs | tumour fitted | 507 | 213 | 0.763 | 0.760 | -0.003 (-0.006 to 0.000) | 92.0% | 92.5% |
| shared-plate pairs | joint fitted | 507 | 213 | 0.763 | 0.760 | -0.003 (-0.006 to -0.000) | 92.0% | 92.5% |
| unpaired, all tumours vs all normals (added after review) | tumour fitted | 507 | 213 | 0.773 | 0.774 | 0.001 (-0.013 to 0.014) | 93.9% | 91.5% |
| unpaired, all tumours vs all normals (added after review) | joint fitted | 507 | 213 | 0.773 | 0.776 | 0.002 (-0.011 to 0.016) | 93.9% | 91.5% |

Tumour-minus-normal difference in the correction covariates, in pooled standard-deviation units. The unpaired contrast carries the between-library difference in alignment metrics that the correction can remove, the paired contrast very little of it.

| Covariate | Unpaired difference | Mean paired difference |
|---|---|---|
| pct_noFeature | 0.86 | 0.13 |
| pct_multimapping | 0.30 | -0.24 |
| log_depth | -0.37 | -0.21 |

*(b) lncRNA scores in 532 tumours against the malignant-marker score and the non-feature fraction*

| Gene set | Genes | Correction | rho with marker score (95% interval) | rho with non-feature fraction (95% interval) |
|---|---|---|---|---|
| single-cell tier 1 (tumour-specific) | 38 | observed | 0.55 (0.48 to 0.62) | 0.49 (0.43 to 0.55) |
| single-cell tier 1 (tumour-specific) | 38 | tumour fitted | 0.48 (0.41 to 0.55) | 0.01 (-0.07 to 0.09) |
| single-cell tier 1 (tumour-specific) | 38 | joint fitted | 0.48 (0.40 to 0.56) | 0.01 (-0.08 to 0.09) |
| single-cell tier 2 (malignant-compartment) | 77 | observed | 0.42 (0.35 to 0.49) | 0.61 (0.56 to 0.66) |
| single-cell tier 2 (malignant-compartment) | 77 | tumour fitted | 0.35 (0.27 to 0.43) | 0.06 (-0.02 to 0.14) |
| single-cell tier 2 (malignant-compartment) | 77 | joint fitted | 0.35 (0.27 to 0.43) | 0.05 (-0.03 to 0.13) |
| single-cell malignant-up vs normal epithelium (meta FDR < 0.05) | 106 | observed | 0.52 (0.45 to 0.59) | 0.71 (0.66 to 0.76) |
| single-cell malignant-up vs normal epithelium (meta FDR < 0.05) | 106 | tumour fitted | 0.47 (0.39 to 0.54) | 0.07 (-0.00 to 0.15) |
| single-cell malignant-up vs normal epithelium (meta FDR < 0.05) | 106 | joint fitted | 0.47 (0.39 to 0.54) | 0.07 (-0.01 to 0.15) |
| abundance-matched single-cell non-specific (null, 1000 draws) | 38 | observed | 0.37 (0.32 to 0.41) | 0.64 (0.57 to 0.71) |
| abundance-matched single-cell non-specific (null, 1000 draws) | 38 | tumour fitted | 0.28 (0.22 to 0.34) | 0.10 (0.07 to 0.12) |
| abundance-matched single-cell non-specific (null, 1000 draws) | 38 | joint fitted | 0.28 (0.22 to 0.33) | 0.08 (0.06 to 0.11) |

*(c) The 15 single-cell candidates set aside in bulk as technically sensitive*

| lncRNA | Tier | Single-cell log2FC vs normal epithelium | rho with metric, observed | rho with metric, corrected | rho with marker score, observed | rho with marker score, corrected | Bulk tumour minus normal, observed | Bulk tumour minus normal, corrected |
|---|---|---|---|---|---|---|---|---|
| THBS3-AS1 | tier2 | n/a | 0.84 | 0.10 | 0.27 | 0.14 | n/a | n/a |

| lncRNA | Tier | Single-cell log2FC vs normal epithelium | rho with metric, observed | rho with metric, corrected | rho with marker score, observed | rho with marker score, corrected | Bulk tumour minus normal, observed | Bulk tumour minus normal, corrected |
|---|---|---|---|---|---|---|---|---|
| LMNTD2-AS1 | tier2 | n/a | 0.81 | 0.09 | 0.35 | 0.26 | n/a | n/a |
| LINC01473 | tier2 | n/a | 0.70 | -0.04 | 0.25 | 0.14 | n/a | n/a |
| LINC01311 | tier2 | n/a | 0.73 | 0.08 | 0.32 | 0.22 | n/a | n/a |
| AL359504.1 | tier2 | n/a | 0.75 | 0.11 | 0.33 | 0.21 | n/a | n/a |
| AL161669.3 | tier2 | n/a | 0.69 | 0.07 | 0.40 | 0.34 | n/a | n/a |
| NR2F2-AS1 | tier1 | 1.42 | 0.69 | 0.11 | 0.40 | 0.34 | -0.10 | -0.14 |
| GAS6-AS1 | tier2 | n/a | 0.65 | 0.05 | 0.33 | 0.19 | n/a | n/a |
| PVT1 | tier1 | 5.64 | 0.57 | -0.03 | 0.37 | 0.26 | 0.54 | 0.52 |
| LINC02615 | tier1 | 3.01 | 0.60 | 0.05 | 0.22 | 0.09 | 0.37 | 0.34 |
| LINC02298 | tier2 | n/a | 0.55 | 0.01 | 0.50 | 0.43 | n/a | n/a |
| AL080317.2 | tier2 | n/a | 0.58 | 0.01 | 0.21 | 0.06 | n/a | n/a |
| PPP1R26-AS1 | tier2 | n/a | 0.55 | 0.01 | 0.24 | 0.12 | n/a | n/a |
| HIF1A-AS3 | tier1 | 6.38 | 0.58 | 0.09 | 0.62 | 0.59 | 1.89 | 1.75 |
| FAM153CP | tier1 | 5.52 | 0.50 | -0.08 | 0.33 | 0.17 | 0.16 | 0.12 |

*Note.* 532 tumours, 72 normals and 72 tumour-normal pairs (68 on a shared plate) from the pan-cancer build, which retains the 4 patients excluded from discovery for survival time. Of 4,088 lncRNAs expressed in either group, 507 have a single-cell estimate. The malignant-marker score (CA9, NDUFA4L2, ANGPTL4, VEGFA, EGLN3) itself correlates with the non-feature fraction at Spearman 0.22. The null row is 1,000 abundance-matched sets of single-cell non-specific lncRNAs (median, and 2.5th to 97.5th percentiles). Contrary to the pre-recorded expectation, these sets are not neutral. The single-cell candidates were locked before any bulk step. Source tables: `42_sc_truth_concordance.tsv`, `42_sc_truth_score_coherence.tsv`, `42_sc_truth_technical_candidates.tsv`, `42_sc_truth_gene_level.tsv`.

**Supplementary Table 11. Residualisation against RUV-III with pseudo-replicates, TCGA-KIRC**

*(a) The leading components and per-gene correlation with the non-feature fraction*

| Version | lncRNA PC1 rho with metric | Protein-coding PC1 rho with metric | Plate R-squared of lncRNA PC1 | Median per-gene abs rho, lncRNAs |
|---|---|---|---|---|
| observed | 0.91 | 0.08 | 0.26 | 0.38 |
| residualised | 0.06 | 0.03 | 0.06 | 0.04 |
| RUV-III (k = 3) | 0.05 | 0.01 | 0.05 | 0.05 |
| RUV-III (k = 1) | 0.10 | 0.19 | 0.04 | 0.10 |
| RUV-III (k = 5) | 0.02 | -0.01 | 0.05 | 0.05 |

*(b) The lncRNA network on the 3,442 discovery lncRNAs and 511 tumours, discovery parameters*

| Version | Modules | Largest module | Unassigned | ARI vs observed | ARI vs residualised | ARI vs RUV-III |
|---|---|---|---|---|---|---|
| observed | 3 | 57.6% | 36.9% | 1.00 | 0.07 | 0.07 |
| residualised | 14 | 13.5% | 29.5% | 0.07 | 1.00 | 0.23 |
| RUV-III (k = 3) | 10 | 14.1% | 45.1% | 0.07 | 0.23 | 1.00 |

*(c) Single-cell ground truth: coherence of the 38 tumour-specific lncRNAs with the malignant-marker score*

| Version | rho with marker score | Null median (500 abundance-matched sets) | Excess over null | rho with metric |
|---|---|---|---|---|
| observed | 0.55 | 0.36 | 0.20 | 0.49 |
| residualised | 0.48 | 0.26 | 0.22 | 0.01 |
| RUV-III (k = 3) | 0.28 | 0.17 | 0.11 | 0.03 |

*(d) The five prognostic modules, fixed discovery loadings, overall survival (Clinical Data Resource) adjusted for age, sex and stage*

| Version | Module | Patients | Deaths | HR per SD (95% CI) | p |
|---|---|---|---|---|---|
| observed | protein-coding green | 525 | 172 | 0.65 (0.56 to 0.75) | 7.2e-09 |
| observed | protein-coding purple | 525 | 172 | 1.45 (1.27 to 1.66) | 4.6e-08 |
| observed | lncRNA blue | 525 | 172 | 0.77 (0.66 to 0.90) | 0.0014 |
| observed | lncRNA greenyellow | 525 | 172 | 1.44 (1.26 to 1.65) | 1.5e-07 |
| observed | lncRNA turquoise | 525 | 172 | 1.42 (1.24 to 1.63) | 3.2e-07 |
| residualised | protein-coding green | 525 | 172 | 0.66 (0.57 to 0.76) | 2.7e-08 |

| Version | Module | Patients | Deaths | HR per SD (95% CI) | p |
|---|---|---|---|---|---|
| residualised | protein-coding purple | 525 | 172 | 1.44 (1.26 to 1.65) | 8.8e-08 |
| residualised | lncRNA blue | 525 | 172 | 0.70 (0.60 to 0.81) | 2.8e-06 |
| residualised | lncRNA greenyellow | 525 | 172 | 1.35 (1.17 to 1.56) | 3.2e-05 |
| residualised | lncRNA turquoise | 525 | 172 | 1.32 (1.13 to 1.53) | 0.00034 |
| RUV-III (k = 3) | protein-coding green | 525 | 172 | 0.64 (0.55 to 0.74) | 2.2e-09 |
| RUV-III (k = 3) | protein-coding purple | 525 | 172 | 1.48 (1.30 to 1.69) | 3.3e-09 |
| RUV-III (k = 3) | lncRNA blue | 525 | 172 | 0.64 (0.55 to 0.74) | 8.7e-09 |
| RUV-III (k = 3) | lncRNA greenyellow | 525 | 172 | 1.37 (1.19 to 1.58) | 1.8e-05 |
| RUV-III (k = 3) | lncRNA turquoise | 525 | 172 | 1.33 (1.13 to 1.56) | 0.00055 |

*Note.* Post hoc sensitivity analysis of 528 TCGA-KIRC tumours with a ccA/ccB class from the pan-cancer build. RUV-III (ruv package) removed k unwanted factors using 12 replicate sets of 66 pseudo-samples (plate, and the extremes of the non-feature fraction and of depth, within ccA/ccB by stage groups) and 1,000 negative-control genes. The single-cell null uses 500 draws, so its values differ slightly from Supplementary Table 10. Source tables: `44_prps_design.tsv`, `44_prps_axis.tsv`, `44_prps_network.tsv`, `44_prps_sc_truth.tsv`, `44_prps_module_hr.tsv`.

#### Supplementary Table 12. Module associations with and without adjustment for the alignment metrics, all 29 modules

##### (a) Clinical covariate set

| Network | Module | HR without adjustment (FDR) | HR with STAR metrics as covariates (FDR) | HR with adjustment for the alignment metrics (FDR) | Status |
|---|---|---|---|---|---|
| lncRNA | blue | 0.85 (0.096) | 0.73 (0.0091) | 0.73 (0.0015) | gained |
| lncRNA | greenyellow | 1.42 (8.6e-05) | 1.34 (0.0091) | 1.31 (0.0092) | robust |
| lncRNA | turquoise | 1.44 (1.3e-05) | 1.47 (0.056) | 1.26 (0.024) | robust |
| lncRNA | cyan | 0.88 (0.19) | 0.78 (0.089) | 0.78 (0.031) | gained |
| lncRNA | black | 1.45 (1.7e-05) | 1.49 (0.089) | 1.28 (0.042) | robust |

| Network | Module | HR without adjustment (FDR) | HR with STAR metrics as covariates (FDR) | HR with adjustment for the alignment metrics (FDR) | Status |
|---|---|---|---|---|---|
| lncRNA | lightcyan | 0.88 (0.18) | 0.82 (0.089) | 0.81 (0.042) | gained |
| lncRNA | midnightblue | 0.99 (0.93) | 0.81 (0.13) | 0.81 (0.043) | gained |
| lncRNA | brown | 1.34 (1.9e-04) | 1.22 (0.10) | 1.17 (0.079) | lost |
| lncRNA | tan | 1.10 (0.33) | 0.90 (0.45) | 0.87 (0.20) | not significant |
| lncRNA | magenta | 1.29 (0.0011) | 1.16 (0.19) | 1.14 (0.22) | lost |
| lncRNA | purple | 1.29 (0.0013) | 1.15 (0.26) | 1.12 (0.22) | lost |
| lncRNA | green | 1.31 (0.0013) | 1.07 (0.82) | 0.98 (0.90) | lost |
| lncRNA | pink | 1.19 (0.049) | 1.01 (0.94) | 0.99 (0.90) | lost |
| lncRNA | red | 1.21 (0.047) | 1.03 (0.85) | 1.01 (0.90) | lost |
| lncRNA | salmon | 1.30 (0.0031) | 1.03 (0.85) | 0.98 (0.90) | lost |
| lncRNA | yellow | 1.18 (0.047) | 0.97 (0.85) | 0.97 (0.90) | lost |
| protein-coding | green | 0.67 (1.5e-05) | 0.68 (8.5e-05) | 0.69 (3.8e-05) | robust |
| protein-coding | purple | 1.27 (0.012) | 1.22 (0.071) | 1.24 (0.036) | robust |
| protein-coding | turquoise | 0.80 (0.014) | 0.78 (0.071) | 0.81 (0.036) | robust |
| protein-coding | pink | 0.91 (0.36) | 0.80 (0.097) | 0.85 (0.10) | not significant |
| protein-coding | magenta | 0.86 (0.14) | 0.84 (0.17) | 0.87 (0.16) | not significant |
| protein-coding | red | 1.45 (2.5e-05) | 1.27 (0.18) | 1.17 (0.16) | lost |
| protein-coding | brown | 1.18 (0.11) | 1.15 (0.18) | 1.14 (0.18) | not significant |
| protein-coding | tan | 1.10 (0.36) | 1.15 (0.18) | 1.15 (0.18) | not significant |
| protein-coding | blue | 0.89 (0.35) | 0.86 (0.18) | 0.88 (0.22) | not significant |
| protein-coding | greenyellow | 0.90 (0.35) | 0.90 (0.36) | 0.92 (0.47) | not significant |
| protein-coding | yellow | 0.98 (0.77) | 0.92 (0.36) | 0.93 (0.47) | not significant |
| protein-coding | black | 1.04 (0.73) | 1.05 (0.58) | 1.06 (0.50) | not significant |
| protein-coding | salmon | 1.03 (0.77) | 0.97 (0.76) | 0.99 (0.93) | not significant |

604 **(b) Clinical covariate set with the ESTIMATE stromal and immune scores added**

| Network | Module | HR without adjustment (FDR) | HR with STAR metrics as covariates (FDR) | HR with adjustment for the alignment metrics (FDR) | Status |
|---|---|---|---|---|---|
| lncRNA | blue | 0.80 (0.018) | 0.68 (5.0e-04) | 0.67 (2.1e-05) | robust |
| lncRNA | greenyellow | 1.45 (3.3e-05) | 1.38 (0.0026) | 1.34 (0.0042) | robust |
| lncRNA | black | 1.47 (9.1e-06) | 1.56 (0.061) | 1.31 (0.023) | robust |
| lncRNA | cyan | 0.87 (0.17) | 0.76 (0.052) | 0.76 (0.023) | gained |

| Network | Module | HR without adjustment (FDR) | HR with STAR metrics as covariates (FDR) | HR with adjustment for the alignment metrics (FDR) | Status |
|---|---|---|---|---|---|
| lncRNA | lightcyan | 0.86 (0.11) | 0.81 (0.061) | 0.79 (0.023) | gained |
| lncRNA | turquoise | 1.50 (8.5e-06) | 1.55 (0.039) | 1.28 (0.023) | robust |
| lncRNA | tan | 1.06 (0.62) | 0.81 (0.11) | 0.78 (0.033) | gained |
| lncRNA | midnightblue | 1.00 (0.98) | 0.80 (0.11) | 0.81 (0.044) | gained |
| lncRNA | brown | 1.39 (2.5e-04) | 1.24 (0.11) | 1.16 (0.19) | lost |
| lncRNA | magenta | 1.29 (0.0034) | 1.13 (0.32) | 1.10 (0.44) | lost |
| lncRNA | purple | 1.30 (0.0020) | 1.15 (0.29) | 1.10 (0.44) | lost |
| lncRNA | red | 1.35 (0.0034) | 1.05 (0.77) | 1.12 (0.46) | lost |
| lncRNA | pink | 1.17 (0.11) | 0.96 (0.77) | 0.94 (0.54) | not significant |
| lncRNA | yellow | 1.68 (8.2e-05) | 1.44 (0.11) | 1.15 (0.54) | lost |
| lncRNA | salmon | 1.28 (0.0047) | 0.99 (0.93) | 0.96 (0.67) | lost |
| lncRNA | green | 1.33 (9.3e-04) | 1.09 (0.64) | 0.99 (0.93) | lost |
| protein-coding | green | 0.605 (2.0e-08) | 0.611 (2.0e-07) | 0.608 (1.4e-08) | robust |
| protein-coding | black | 1.50 (0.0054) | 1.33 (0.092) | 1.52 (0.0044) | robust |
| protein-coding | purple | 1.30 (0.0051) | 1.28 (0.017) | 1.27 (0.013) | robust |
| protein-coding | turquoise | 0.81 (0.024) | 0.78 (0.065) | 0.82 (0.052) | lost |
| protein-coding | pink | 0.93 (0.57) | 0.80 (0.092) | 0.86 (0.13) | not significant |
| protein-coding | magenta | 0.86 (0.21) | 0.82 (0.092) | 0.88 (0.22) | not significant |
| protein-coding | red | 1.48 (3.1e-05) | 1.29 (0.12) | 1.16 (0.22) | lost |
| protein-coding | tan | 1.08 (0.57) | 1.13 (0.24) | 1.14 (0.22) | not significant |
| protein-coding | brown | 1.16 (0.21) | 1.16 (0.24) | 1.11 (0.39) | not significant |
| protein-coding | blue | 0.93 (0.59) | 0.78 (0.12) | 0.91 (0.51) | not significant |
| protein-coding | yellow | 1.18 (0.59) | 1.09 (0.77) | 0.82 (0.51) | not significant |
| protein-coding | greenyellow | 0.94 (0.59) | 0.92 (0.56) | 0.97 (0.82) | not significant |
| protein-coding | salmon | 1.06 (0.59) | 0.97 (0.77) | 1.01 (0.92) | not significant |

*Note for panels (a) and (b).* TCGA-KIRC, 504 patients and 163 events for the lncRNA modules and 502 patients and 165 events for the protein-coding modules in every row. Source: `12_technical_adjustment_sensitivity.tsv`, with confidence intervals for every cell in `12_technical_adjustment_arms.tsv`. Membership is fixed at the definition made on the residualised matrix, and the three arms differ only in how the score is computed and what enters the model. Status is robust (significant in both arms), lost (without adjustment only), gained (with adjustment only) or never significant. Benjamini-Hochberg control is within one network, arm and covariate set, so over 13 or 16 tests, not 29. Under the clinical covariate set, 15 of 29 modules are significant without adjustment, 10 with adjustment for the alignment metrics and 3 with the STAR metrics as covariates. Nine are lost, 4 gained, 6 robust and 10 never significant. Under the clinical plus ESTIMATE set the counts are 16, 11 and 5, with 9 lost, 4 gained, 7 robust and 9 never significant. The protein-coding green row of panel (b) is given to three decimals because the three arms agree to two.

**(c) The same contrast with RIN and slide necrosis added to the clinical covariate set**

| Covariate set | Significant without adjustment | Significant with adjustment | Lost | Gained | Robust | Not significant |
|---|---|---|---|---|---|---|
| Clinical | 15 of 29 (11 lncRNA, 4 protein-coding) | 10 of 29 (7 lncRNA, 3 protein-coding) | 9 | 4 | 6 | 10 |
| Clinical + RIN + slide necrosis | 12 of 29 (9 lncRNA, 3 protein-coding) | 4 of 29 (3 lncRNA, 1 protein-coding) | 9 | 1 | 3 | 16 |

The five prognostic modules under the clinical, RIN and necrosis covariate set:

| Module | HR without adjustment (95% CI), FDR | HR with adjustment for the alignment metrics (95% CI), FDR | Status |
|---|---|---|---|
| lncRNA blue | 0.87 (0.73 to 1.04), 0.17 | 0.74 (0.63 to 0.87), 0.0035 | gained |
| lncRNA greenyellow | 1.41 (1.20 to 1.66), 1.8e-04 | 1.30 (1.10 to 1.54), 0.018 | robust |
| lncRNA turquoise | 1.47 (1.26 to 1.71), 1.2e-05 | 1.28 (1.08 to 1.51), 0.019 | robust |
| Protein-coding green | 0.69 (0.58 to 0.81), 4.9e-05 | 0.70 (0.60 to 0.83), 1.8e-04 | robust |
| Protein-coding purple | 1.26 (1.07 to 1.48), 0.021 | 1.24 (1.05 to 1.46), 0.068 | lost |

*Note.* TCGA-KIRC, 503 patients and 163 events for the lncRNA modules and 501 patients and 165 events for the protein-coding modules, one fewer in each network than panels (a) and (b) because one case lacks an RNA integrity number. Source: `23_technical_adjustment_with_rin.tsv`. Adding RIN and slide necrosis to the clinical block reduces the number of apparently prognostic modules before adjustment, and further after it. The direction of the contrast is unchanged.

#### Supplementary Table 13. Published TCGA-KIRC lncRNA signatures re-scored

##### (a) Provenance and mapping

| Signature | Weight provenance | Tier | Published genes | Scored | In the 3,442-gene production set |
|---|---|---|---|---|---|
| Jiang 2019[4], 9 lncRNA nomogram | published coefficients | primary | 9 | 9 | 7 |
| Gui 2021[5], 8 autophagy-related | unit weights signed in these same patients | primary | 8 | 7 | 6 |
| Yu 2021[6], m6A | footnote, incomplete mapping | footnote | 5 | 2 | 2 |
| Hong 2022[7], cuproptosis | published coefficients | primary | 4 | 4 | 3 |
| Liu 2023[8], pyroptosis | unit weights signed in these same patients | primary | 6 | 6 | 5 |
| Feng 2024[9], disulfidptosis | unit weights signed in these same patients | primary | 6 | 6 | 5 |

*Note.* Source: `26_signature_provenance_key.tsv`, `26_published_signatures_genes.tsv`. The six signatures publish 38 genes, of which 36 are in the current annotation and 34 are scored. The two unscored genes have a median FPKM of 0. Neither Ensembl host resolved two names. Hong 2022 and Feng 2024 share AC026401.3, so the six signatures are correlated arms, not independent tests. Gui 2021, Liu 2023 and Feng 2024 publish neither coefficients nor per-gene hazard directions. Their weights are therefore plus or minus one, signed by the univariate Cox coefficient in these same 528 patients, so their p values are invalid and their attenuation is inflated. Yu 2021 retained 2 of its 5 lncRNAs, lost the gene with the largest published coefficient and scores opposite to the published model. Ranges must never be quoted across two provenance classes.

##### (b) Correlation of each score with the alignment metrics

| Signature, published weights | rho with the non-feature fraction (p) | rho with the lncRNA axis | rho with the multimapping fraction | rho with the non-feature fraction after residualisation (p) |
|---|---|---|---|---|
| Jiang 2019 | 0.43 (1.1e-25) | 0.39 | 0.30 | 0.04 (0.41) |
| Gui 2021 | 0.35 (5.3e-17) | 0.32 | 0.20 | -0.02 (0.62) |
| Yu 2021 | -0.24 (1.8e-08) | -0.34 | 0.15 | 0.00 (0.96) |
| Hong 2022 | 0.17 (6.9e-05) | 0.18 | 0.33 | -0.03 (0.51) |
| Liu 2023 | 0.11 (0.0087) | -0.04 | 0.32 | 0.00 (0.96) |
| Feng 2024 | 0.43 (6.3e-25) | 0.53 | 0.32 | 0.02 (0.58) |

*Note.* TCGA-KIRC, 528 patients for the non-feature and multimapping fractions and 511 for the lncRNA axis. Source: `26_published_signatures_correlations.tsv`. With coefficients re-estimated in this cohort, correlations with the non-feature fraction range from 0.18 to 0.50. After residualising the expression matrix on the three STAR metrics, all twelve arms lie within 0.04 of zero (p at least 0.41), confirming that the adjustment works. Residualisation does not zero the correlations with the other two quantities. Against the lncRNA axis, the residualised published scores of Liu (-0.18), Feng (0.15) and Yu (-0.14) survive Benjamini-Hochberg control.

**(c) Hazard ratios per standard deviation, published weights**

| Signature | Score alone | + clinical model | + clinical model + the three STAR metrics | Residualised score + clinical model |
|---|---|---|---|---|
| Jiang 2019 | 1.93 (1.70 to 2.19) | 1.67 (1.45 to 1.93) | 1.65 (1.38 to 1.97) | 1.55 (1.33 to 1.79) |
| Gui 2021 | 1.87 (1.65 to 2.13) | 1.50 (1.31 to 1.71) | 1.39 (1.19 to 1.62) | 1.41 (1.22 to 1.63) |
| Yu 2021 | 1.63 (1.36 to 1.96) | 1.46 (1.21 to 1.77) | 1.51 (1.25 to 1.81) | 1.57 (1.31 to 1.86) |
| Hong 2022 | 2.13 (1.87 to 2.43) | 1.75 (1.51 to 2.03) | 1.70 (1.44 to 2.02) | 1.70 (1.46 to 1.99) |
| Liu 2023 | 1.97 (1.71 to 2.28) | 1.63 (1.39 to 1.90) | 1.52 (1.29 to 1.79) | 1.53 (1.30 to 1.79) |
| Feng 2024 | 1.89 (1.64 to 2.18) | 1.53 (1.31 to 1.79) | 1.46 (1.19 to 1.79) | 1.39 (1.17 to 1.65) |

**(d) Hazard ratios per standard deviation, coefficients re-estimated in TCGA-KIRC**

| Signature | + clinical model | + clinical model + the three STAR metrics | Residualised score + clinical model |
|---|---|---|---|
| Jiang 2019 | 1.88 (1.60 to 2.22) | 1.85 (1.53 to 2.24) | 1.70 (1.45 to 2.00) |
| Gui 2021 | 1.53 (1.35 to 1.74) | 1.46 (1.25 to 1.72) | 1.46 (1.28 to 1.67) |
| Yu 2021 | 1.63 (1.40 to 1.89) | 1.52 (1.28 to 1.80) | 1.57 (1.33 to 1.85) |
| Hong 2022 | 1.74 (1.50 to 2.01) | 1.68 (1.43 to 1.98) | 1.70 (1.46 to 1.98) |
| Liu 2023 | 1.60 (1.39 to 1.85) | 1.51 (1.27 to 1.80) | 1.52 (1.30 to 1.78) |
| Feng 2024 | 1.67 (1.40 to 1.98) | 1.59 (1.29 to 1.96) | 1.57 (1.30 to 1.89) |

*Note for panels (c) and (d).* Score-alone models use 528 patients and 173 events, adjusted models 520 patients and 172 events. Source: `26_published_signatures_models.tsv`. All 24 adjusted estimates remain significant (largest p 2.9e-04). The refit arm is fitted and evaluated on the same patients, so it is optimistic by construction.

**(e) Attenuation of the clinically adjusted log hazard ratio**

| Signature, published weights | Change on adding the three STAR metrics (%) | Change on residualising the expression (%) | Provenance |
|---|---|---|---|
| Jiang 2019 | -2.6 | -15.0 | published coefficients |
| Hong 2022 | -4.9 | -5.1 | published coefficients |
| Gui 2021 | -19.0 | -14.9 | signed in sample |
| Liu 2023 | -14.1 | -13.2 | signed in sample |
| Feng 2024 | -11.1 | -23.6 | signed in sample |
| Yu 2021 | 7.6 | 17.5 | footnote, 2 of 5 genes |

*Note.* Source: `26_published_signatures_models.tsv`, `26_headline_ranges_by_provenance.tsv`. For the only two signatures whose weights were not re-derived, the log hazard ratio moves by a few per cent and is not abolished. All six models were nonetheless fitted in TCGA-KIRC, so no arm is out of sample relative to the discovery cohort. With coefficients re-estimated in this cohort, the five refit arms move by -

12.6% to -2.8% on adding the STAR metrics and by -15.9% to -4.1% on residualising. Yu 2021 moves the other way and must not be pooled into any range.

**(f) Apparent increment in the concordance index over the reference model of the same specification**

| Signature, published weights | Over the clinical model | Over the clinical model plus the STAR metrics | Residualised score over the clinical model |
|---|---|---|---|
| Jiang 2019 | 0.0215 (0.0014 to 0.0445), p 0.041 | 0.0105 (-0.0071 to 0.0290), p 0.24 | 0.0138 (-0.0057 to 0.0350), p 0.17 |
| Gui 2021 | 0.0182 (-0.0007 to 0.0372), p 0.055 | 0.0117 (-0.0024 to 0.0260), p 0.11 | 0.0107 (-0.0061 to 0.0277), p 0.20 |
| Yu 2021 | 0.0086 (-0.0071 to 0.0245), p 0.30 | 0.0173 (0.0030 to 0.0330), p 0.022 | 0.0147 (-0.0032 to 0.0325), p 0.12 |
| Hong 2022 | 0.0236 (0.0030 to 0.0447), p 0.024 | 0.0169 (-0.0018 to 0.0364), p 0.078 | 0.0163 (-0.0034 to 0.0372), p 0.11 |
| Liu 2023 | 0.0181 (-0.0025 to 0.0397), p 0.074 | 0.0144 (-0.0017 to 0.0311), p 0.089 | 0.0140 (-0.0046 to 0.0328), p 0.14 |
| Feng 2024 | 0.0182 (0.0002 to 0.0359), p 0.046 | 0.0070 (-0.0061 to 0.0200), p 0.30 | 0.0107 (-0.0047 to 0.0253), p 0.18 |

*Note.* TCGA-KIRC, 520 patients and 172 events, paired patient bootstrap over 2,000 resamples. Source: `26_published_signatures_models.tsv`. All increments are apparent and in sample. Each score is evaluated on the patients whose model it entered, and four of the six signatures also derived their weights in this cohort. The modules of this paper have an apparent increment of 0.0205 and a cross-validated increment of 0.0092 (493 patients, 159 events). The relevant comparison is between apparent and cross-validated increments.

**(g) Position of each signature in two random-signature nulls**

| Signature, published weights | Set size | rho with the metric | Percentile of the absolute rho, outcome-signed null | Percentile of the absolute rho, sign-randomised null | Percentile of the univariate HR, outcome-signed null |
|---|---|---|---|---|---|
| Liu 2023 | 6 | 0.11 | 8.6 | 19.1 | 99.9 |
| Hong 2022 | 4 | 0.17 | 15.9 | 25.4 | 100.0 |
| Gui 2021 | 7 | 0.35 | 28.5 | 55.4 | 99.7 |
| Jiang 2019 | 9 | 0.43 | 30.9 | 65.3 | 99.8 |

| Signature, published weights | Set size | rho with the metric | Percentile of the absolute rho, outcome-signed null | Percentile of the absolute rho, sign-randomised null | Percentile of the univariate HR, outcome-signed null |
|---|---|---|---|---|---|
| Yu 2021 | 2 | -0.24 | 31.1 | 34.5 | 97.6 |
| Feng 2024 | 6 | 0.43 | 36.7 | 64.9 | 99.7 |

Median absolute correlation with the metric of the two nulls, by set size:

| Set size | Outcome-signed | Sign-randomised | Median paired difference | Wilcoxon p |
|---|---|---|---|---|
| 2 | 0.40 | 0.36 | 0.000 | 3.5e-06 |
| 4 | 0.48 | 0.34 | 0.056 | 5.9e-38 |
| 6 | 0.53 | 0.33 | 0.137 | 1.5e-63 |
| 7 | 0.54 | 0.32 | 0.144 | 2.3e-69 |
| 9 | 0.57 | 0.33 | 0.169 | 3.8e-86 |

*Note.* One set of 1,000 random gene sets per size, weighted two ways and paired on the same draws. Source: `26_random_signature_null.tsv`. The paired contrast shows directly that outcome-signing itself induces correlation with the metric in the null, so outcome-signed percentiles cannot show that a published signature is less correlated with the metric than chance. Against the sign-randomised null, three of the six published arms sit at or above the median. Percentiles carry Monte-Carlo error of about plus or minus 1.5 points. Neither null matches the magnitude of a published coefficient vector, because both weight every drawn gene plus or minus one.

#### Supplementary Table 14. Module hazard ratios under four codings of unassessed nodal and metastasis status

| Module | A: NX as N0, MX as M0, M1 reconciled from stage IV | B: NX and MX as their own categories | C: assessed N and M only | D: M1 as coded, no reconciliation |
|---|---|---|---|---|
| Protein-coding green | 0.62 (0.53 to 0.74), p 1.7e-08 | 0.62 (0.53 to 0.73), p 1.3e-08 | 0.55 (0.43 to 0.70), p 1.2e-06 | 0.63 (0.54 to 0.74), p 2.6e-08 |

| Module | A: NX as N0, MX as M0, M1 reconciled from stage IV | B: NX and MX as their own categories | C: assessed N and M only | D: M1 as coded, no reconciliation |
|---|---|---|---|---|
| Protein-coding purple | 1.28 (1.09 to 1.50), p 0.0025 | 1.30 (1.10 to 1.53), p 0.0018 | 1.33 (1.03 to 1.72), p 0.027 | 1.28 (1.09 to 1.50), p 0.0025 |
| lncRNA blue | 0.71 (0.60 to 0.83), p 3.0e-05 | 0.70 (0.60 to 0.83), p 2.3e-05 | 0.61 (0.48 to 0.78), p 6.3e-05 | 0.72 (0.61 to 0.84), p 5.9e-05 |
| lncRNA greenyellow | 1.35 (1.15 to 1.60), p 3.4e-04 | 1.36 (1.15 to 1.60), p 2.5e-04 | 1.45 (1.17 to 1.81), p 7.4e-04 | 1.37 (1.16 to 1.62), p 1.7e-04 |
| lncRNA turquoise | 1.24 (1.06 to 1.45), p 0.0074 | 1.26 (1.07 to 1.48), p 0.0054 | 1.48 (1.11 to 1.97), p 0.0082 | 1.27 (1.08 to 1.48), p 0.0039 |

*Note.* TCGA-KIRC, principal specification in every cell. Specification A is the primary analysis and reproduces the discovery column of Table 2 of the main text. Specifications A, B and D use 502 patients and 165 events for the protein-coding modules and 504 patients and 163 events for the lncRNA modules. Specification C uses 240 patients and 87 events for the protein-coding modules and 241 patients and 87 events for the lncRNA modules. Source: `03_nodal_metastasis_sensitivity.tsv`. All p values are nominal. Hazard ratios under specifications B and D lie within 0.020 and 0.026, respectively, of their specification A estimates. Specification C is a differently and more completely staged population, not a matched subset, so its larger effects do not show that the primary coding is conservative. Together, the four codings show that the primary coding did not create the associations.

#### Supplementary Table 15. Sex-disaggregated results

##### (a) Module scores associated with sex in TCGA-KIRC

| Module | Point-biserial r, male coded 1 | Wilcoxon p | BH over 29 modules | Higher in |
|---|---|---|---|---|
| lncRNA lightcyan | 0.23 | 4.3e-07 | 6.2e-06 | men |
| lncRNA greenyellow | 0.22 | 3.6e-07 | 6.2e-06 | men |
| Protein-coding green | -0.19 | 1.1e-05 | 1.0e-04 | women |
| Protein-coding purple | 0.10 | 0.0037 | 0.027 | men |
| lncRNA blue | -0.12 | 0.0052 | 0.028 | women |

| Module | Point-biserial r, male coded 1 | Wilcoxon p | BH over 29 modules | Higher in |
|---|---|---|---|---|
| Protein-coding tan | 0.12 | 0.0057 | 0.028 | men |
| Protein-coding black | 0.11 | 0.0071 | 0.029 | men |
| Protein-coding red | -0.11 | 0.012 | 0.044 | women |

*Note.* TCGA-KIRC, 511 patients for the lncRNA modules (178 women, 333 men) and 506 for the protein-coding modules (179 women, 327 men). Source: `33_sex_correlations.tsv`. Benjamini-Hochberg is applied to the Wilcoxon p values across the 29 module eigengenes within a cohort. Eight of 29 eigengenes differ by sex, including four of the five prognostic modules, all but lncRNA turquoise (r -0.03, BH 0.78). The counts are 3 of 29 in CPTAC-3 (1 prognostic), 10 of 29 in TCGA-KIRP (3 prognostic) and 1 of 29 in TCGA-KICH (none prognostic).

**(b) Alignment metrics by sex in TCGA-KIRC**

| Quantity | Median, women | Median, men | Hodges-Lehmann shift (95% CI) | Wilcoxon p | BH |
|---|---|---|---|---|---|
| Non-feature read fraction, % | 5.0 | 4.5 | -0.26 (-0.68 to 0.13) | 0.21 | 0.28 |
| Multimapping fraction, % | 7.2 | 7.7 | 0.37 (0.13 to 0.62) | 0.0027 | 0.011 |
| log10 assigned reads | 7.763 | 7.770 | 0.003 (-0.015 to 0.022) | 0.71 | 0.71 |
| lncRNA axis | -0.0018 | -0.0100 | -0.006 (-0.015 to 0.002) | 0.12 | 0.25 |

*Note.* 528 patients (185 women, 343 men) for the STAR metrics and 511 (178, 333) for the axis. Source: `33_noFeature_by_sex.tsv`. Of the alignment metrics, only the multimapping fraction differs by sex. The axis is in raw loading units in this file, so its effect size is not comparable with the per-standard-deviation axis used elsewhere.

**(c) Module by sex interaction, common-covariate model**

| Cohort and specification | Module | HR in women | HR in men | Interaction HR (95% CI) | Wald p | BH over 5 modules |
|---|---|---|---|---|---|---|
| TCGA-KIRC, principal | Protein-coding green | 0.64 | 0.62 | 0.96 (0.70 to 1.32) | 0.82 | 0.82 |

| Cohort and specification | Module | HR in women | HR in men | Interaction HR (95% CI) | Wald p | BH over 5 modules |
|---|---|---|---|---|---|---|
| TCGA-KIRC, principal | Protein-coding purple | 1.06 | 1.45 | 1.36 (1.03 to 1.80) | 0.030 | 0.15 |
| TCGA-KIRC, principal | lncRNA blue | 0.62 | 0.75 | 1.20 (0.87 to 1.65) | 0.26 | 0.35 |
| TCGA-KIRC, principal | lncRNA greenyellow | 1.58 | 1.29 | 0.81 (0.59 to 1.12) | 0.20 | 0.35 |
| TCGA-KIRC, principal | lncRNA turquoise | 1.06 | 1.28 | 1.21 (0.86 to 1.71) | 0.28 | 0.35 |
| CPTAC-3, six-parameter | lncRNA greenyellow | 2.33 | 1.23 | 0.53 (0.27 to 1.04) | 0.065 | 0.32 |
| CPTAC-3, six-parameter | Protein-coding purple | 1.85 | 1.26 | 0.68 (0.37 to 1.25) | 0.21 | 0.50 |
| TCGA-KIRP, six-parameter | lncRNA blue | 0.39 | 0.90 | 2.32 (1.12 to 4.82) | 0.024 | 0.067 |
| TCGA-KIRP, six-parameter | lncRNA turquoise | 3.00 | 0.97 | 0.32 (0.12 to 0.88) | 0.027 | 0.067 |
| TCGA-KIRP, six-parameter | lncRNA greenyellow | 1.72 | 1.32 | 0.77 (0.43 to 1.38) | 0.38 | 0.47 |

*Note.* The interaction model has one baseline hazard, covariate effects common to both sexes and a module-by-sex product term. Patients and events: TCGA-KIRC 502 and 165 for the protein-coding modules and 504 and 163 for the lncRNA modules, CPTAC-3 199 and 32, TCGA-KIRP 263 and 40. Source: `33_sex_interaction_models.tsv`. No interaction test survives Benjamini-Hochberg control in any cohort. The family is the five modules of one cohort-and-specification block, not all 20 tests. The smallest raw p over the 20 module interaction tests is 0.024. All five CPTAC-3 rows are below the events-per-variable threshold of 5, at 4.6. The metric exposures show no sex interaction. The non-feature fraction gives 1.41 in women and 1.34 in men (interaction hazard ratio 0.95, 0.73 to 1.24, p 0.71), and the axis 1.34 and 1.35 (1.01, 0.73 to 1.39, p 0.97).

**(d) Sex-specific fits, a different estimand**

| Cohort and specification | Module | HR in women (95% CI), n / events | HR in men (95% CI), n / events | Ratio, men over women (95% CI) | Wald p | BH over 5 modules |
|---|---|---|---|---|---|---|
| TCGA-KIRC, principal | lncRNA greenyellow | 1.97 (1.40 to 2.78), 174 / 57 | 1.20 (0.98 to 1.47), 330 / 106 | 0.61 (0.41 to 0.91) | 0.015 | 0.074 |
| TCGA-KIRC, principal | lncRNA blue | 0.59 (0.43 to 0.79), 174 / 57 | 0.76 (0.62 to 0.94), 330 / 106 | 1.30 (0.90 to 1.87) | 0.16 | 0.40 |
| TCGA-KIRC, principal | Protein-coding purple | 1.12 (0.85 to 1.47), 176 / 60 | 1.35 (1.09 to 1.68), 326 / 105 | 1.21 (0.85 to 1.71) | 0.29 | 0.48 |
| TCGA-KIRC, principal | lncRNA turquoise | 1.12 (0.80 to 1.57), 174 / 57 | 1.25 (1.04 to 1.50), 330 / 106 | 1.12 (0.76 to 1.64) | 0.58 | 0.72 |
| TCGA-KIRC, principal | Protein-coding green | 0.65 (0.49 to 0.85), 176 / 60 | 0.63 (0.49 to 0.80), 326 / 105 | 0.97 (0.68 to 1.39) | 0.87 | 0.87 |

*Note.* Separate baseline hazards and covariate effects in women and in men. Source: `33_sex_stratified_hr.tsv`, `33_sex_contrast_reconciliation.tsv`. Hazard ratios are per standard deviation of the pooled score. Per within-stratum standard deviation, greenyellow gives 1.76 (1.32 to 2.34) in women and 1.21 (0.98 to 1.49) in men, because the score varies less in women (standard deviations 0.83 and 1.04). The sex-specific fits and the interaction model of panel (c) estimate different quantities and disagree in 4 of 25 rows, so each hazard ratio must be quoted with its own model's p value. For lncRNA greenyellow the sex-specific fits give a difference at Wald p 0.015 and likelihood-ratio p 0.015, whereas the interaction model on the same patients gives 0.81 at p 0.20. Nothing survives multiplicity in either test. The five modules were chosen by overall survival in these same patients, so the family is post-selection and every false discovery rate is conditional on that selection. Fifteen of the 50 stratified models, all ten CPTAC-3 rows and the five TCGA-KIRP female rows, are below the events-per-variable threshold of 5.

**(e) X- and Y-encoded members of the lncRNA network**

| Set | X or Y encoded | Total | Odds ratio (95% CI) | Fisher p | BH over 17 tests |
|---|---|---|---|---|---|
| The analysed lncRNA network | 76 (2.2%) | 3,442 | 0.98 (0.75 to 1.27) | 0.95 | n/a |
| lncRNA turquoise | 9 (2.1%) | 438 | 0.91 (0.41 to 1.77) | 1.00 | 1.00 |
| lncRNA blue | 3 (1.2%) | 246 | 0.53 (0.11 to 1.59) | 0.38 | 0.81 |
| lncRNA greenyellow | 2 (1.9%) | 108 | 0.82 (0.10 to 3.06) | 1.00 | 1.00 |
| lncRNA black | 9 (5.0%) | 179 | 2.34 (1.04 to 4.59) | 0.020 | 0.20 |
| lncRNA red | 0 | 186 | 0 (0 to 0.87) | 0.024 | 0.20 |

*Note.* Chromosomes are from the GENCODE v36 annotation. The reference set is the GENCODE v36 lncRNA genes outside the tested set, of which 379 of 16,889 (2.2%) are X or Y encoded. Source: `33_xy_lncRNA_counts_by_module.tsv`. Sixty-eight of the 76 are on chrX and 8 on chrY. No module survives Benjamini-Hochberg control over the 17 tests. XIST is in the network but unassigned (module membership -0.26), as are 7 of the 8 chrY lncRNAs.

**(f) Sensitivity of the sex contrast**

| Analysis | lncRNA greenyellow, women | lncRNA greenyellow, men | p for the sex difference |
|---|---|---|---|
| Baseline | 1.97 | 1.20 | 0.015 |
| X- and Y-encoded members deleted, 2 of 108 genes | 2.00 | 1.20 | 0.012 |
| Recorded sex replaced by the marker call where the two disagree | 1.93 | 1.22 | 0.023 |
| The five discordant samples dropped | 1.96 | 1.21 | 0.018 |

*Note.* TCGA-KIRC, principal specification. Source: `33_sensitivity_analyses.tsv`, `33_sex_label_check.tsv`. Deleting the X- and Y-encoded members of each prognostic module (20, 6, 3, 2 and 9 genes) changes the score negligibly, with correlations of 0.9997 to 0.9999 against the full score. Recorded sex was checked against XIST and the mean of eight chrY lncRNAs in the 511 discovery samples of the lncRNA network: 448 concordant, 5 discordant and 58 ambiguous. The ambiguous calls are mostly recorded males with low chrY expression, consistent with loss of chromosome Y in renal tumour tissue, so the discordance rate is best quoted as 5 of the 453 samples the markers call. Marker cut-offs were fixed in advance. All sex-disaggregated analyses are descriptive.

#### Supplementary Table 16. Module estimates in chromophobe renal cell carcinoma

| Module | HR | 95% CI | Nominal p | Zsummary in TCGA-KICH |
|---|---|---|---|---|
| Protein-coding green | 0.88 | 0.20 to 3.90 | 0.87 | 4.83 |
| Protein-coding purple | 2.67 | 1.25 to 5.70 | 0.011 | 28.18 |
| lncRNA blue | 1.41 | 0.48 to 4.17 | 0.53 | 1.76 |
| lncRNA greenyellow | 5.50 | 1.06 to 28.61 | 0.043 | 6.70 |
| lncRNA turquoise | 1.77 | 0.60 to 5.19 | 0.30 | 11.10 |

*Note.* TCGA-KICH, 65 patients and 9 events. Hazard ratio per standard deviation of the projected module eigengene, from the shared six-parameter cross-cohort model: the eigengene plus age, sex, T category, M1 and the non-feature fraction. Grade is not modelled, being unrecorded for this cohort and inapplicable to chromophobe histology. Source: `11_subtype_module_effects.tsv`, preservation column from `24_module_preservation_lncRNA.tsv` and `24_module_preservation_mRNA.tsv`. Nine events against six parameters give 1.5 events per variable, below the adequacy threshold of 5, so these estimates are reported for completeness and not interpreted. The greenyellow interval spans a 27-fold range, and lncRNA blue is not preserved in this cohort. Undoing the reconciliation of stage IV to M1 changes nothing, because no TCGA-KICH patient was reconciled. With nine events, the twelve-covariate model shows complete separation, so the six-parameter set is used.

#### Supplementary Table 17. Module preservation of the discovery networks in the three external cohorts

**(a) Summary**

| Network | Test cohort | Modules | Strong | Weak to moderate | None | Median Zsummary | Minimum | Maximum | Test samples | Test genes | Permutations |
|---|---|---|---|---|---|---|---|---|---|---|---|
| lncRNA | CPTAC-3 | 16 | 13 | 3 | 0 | 13.39 | 2.20 | 30.65 | 200 | 3,442 | 100 |
| lncRNA | TCGA-KIRP | 16 | 7 | 8 | 1 | 8.04 | 1.77 | 29.39 | 269 | 3,442 | 100 |
| lncRNA | TCGA-KICH | 16 | 3 | 11 | 2 | 5.36 | 0.33 | 13.31 | 65 | 3,441 | 100 |
| Protein-coding | CPTAC-3 | 13 | 13 | 0 | 0 | 43.99 | 17.69 | 101.66 | 200 | 12,000 | 30 |
| Protein-coding | TCGA-KIRP | 13 | 13 | 0 | 0 | 35.08 | 15.24 | 71.33 | 269 | 12,000 | 30 |
| Protein-coding | TCGA-KICH | 13 | 10 | 3 | 0 | 18.72 | 4.83 | 49.11 | 65 | 12,000 | 30 |

**(b) lncRNA network, Zsummary per module**

| Module | Size | CPTAC-3 | TCGA-KIRP | TCGA-KICH |
|---|---|---|---|---|
| blue | 246 | 30.65 | 8.47 | 1.76 |
| greenyellow | 108 | 15.14 | 3.47 | 6.70 |
| turquoise | 438 | 13.42 | 29.39 | 11.10 |
| yellow | 192 | 29.39 | 12.51 | 10.71 |
| red | 186 | 27.21 | 7.61 | 5.90 |
| pink | 174 | 20.14 | 16.63 | 8.66 |
| magenta | 122 | 18.71 | 1.77 | 4.82 |
| tan | 93 | 16.18 | 6.90 | 5.94 |
| black | 179 | 13.36 | 18.72 | 4.65 |
| lightcyan | 44 | 12.74 | 3.09 | 0.33 |
| salmon | 72 | 12.19 | 5.45 | 2.19 |
| brown | 219 | 12.07 | 19.80 | 13.31 |
| cyan | 54 | 11.59 | 4.76 | 3.19 |
| green | 186 | 6.34 | 13.87 | 3.55 |
| midnightblue | 53 | 3.90 | 6.93 | 4.33 |
| purple | 110 | 2.20 | 19.64 | 8.14 |

**(c) Protein-coding network, Zsummary per module**

| Module | Size | CPTAC-3 | TCGA-KIRP | TCGA-KICH |
|---|---|---|---|---|
| green | 670 | 61.62 | 34.79 | 4.83 |
| purple | 216 | 52.20 | 36.70 | 28.18 |
| yellow | 986 | 101.66 | 64.67 | 49.11 |
| blue | 994 | 79.48 | 29.65 | 20.88 |
| black | 323 | 57.44 | 24.06 | 15.67 |
| tan | 152 | 51.48 | 35.08 | 26.22 |
| red | 473 | 43.99 | 42.96 | 11.90 |
| magenta | 227 | 43.50 | 29.28 | 13.13 |
| greenyellow | 164 | 25.93 | 26.05 | 18.72 |
| salmon | 63 | 25.06 | 15.24 | 9.22 |
| pink | 270 | 23.18 | 37.10 | 9.85 |
| turquoise | 2,405 | 18.99 | 55.77 | 35.00 |
| brown | 988 | 17.69 | 71.33 | 23.77 |

*Note.* WGCNA `modulePreservation`, with the residualised TCGA-KIRC matrix on which the modules were defined as reference and each test network residualised on its own cohort's STAR metrics. Bands: Zsummary above 10 strong, 2 to 10 weak to moderate, below 2 none. Source: `24_module_preservation_summary.tsv`, `24_module_preservation_lncRNA.tsv`, `24_module_preservation_mRNA.tsv`. The algorithm sub-sampled the protein-coding turquoise module to 1,000 genes. `Z.density` and `Z.connectivity` are not populated, and there is no per-module preservation p value. Median ranks (lower is better preserved) for lncRNA blue, greenyellow and turquoise and protein-coding green and purple are 5, 7, 13, 7 and 3 in CPTAC-3, 14, 17, 5, 12 and 3 in TCGA-KIRP, and 15, 7, 8, 14 and 2 in TCGA-KICH. Band edges are exact, so lncRNA blue at 1.76 in TCGA-KICH is not preserved, while salmon at 2.19 is weak to moderate.

#### Supplementary Table 18. The catabolic protein-coding module as a proximal-tubule programme

**(a) Marker set and module hubs**

| Quantity | Value |
|---|---|
| Proximal-tubule marker genes assembled | 21 |
| Of those, flagged core | 15 |
| Of those, named as proximal tubule by the Human Protein Atlas single-cell string | 18 |
| Of those, inside the protein-coding network | 14, all 14 members of the catabolic module |
| Leave-out markers, not members of that module | 7 |
| Markers exclusive to proximal tubule in the Human Protein Atlas | 7: LRP2, SLC34A1, SLC22A6, SLC13A3, SLC17A1, SLC7A13, MIOX |
| Top 20 module hubs whose Human Protein Atlas string names proximal tubule | 17 |
| Of those, exclusive to proximal tubule | 1, LRP2 |
| Top 20 hubs also in the 21-gene marker set | 3: PDZK1, LRP2, GLYAT |
| Top 20 hubs in ClearCode34 | 1: GIPC2, a ccA gene |

*Note.* Source: `34_pt_marker_set.tsv`, `34_green_hubs_pt_flag.tsv`. The Human Protein Atlas flag is permissive. Of the 17 flagged hubs, 2 name one other cell type and 14 appear in lists of 4 to 12 cell types. The flag has no background rate.

**(b) Correlations of the module and the marker score**

| Feature | Variable | Expression space | Pearson r | Spearman rho | n |
|---|---|---|---|---|---|
| Catabolic module eigengene | proximal-tubule marker score, 21 genes | residualised | 0.84 | 0.85 | 506 |
| Catabolic module eigengene | proximal-tubule marker score, 21 genes | observed | 0.86 | 0.88 | 506 |
| Catabolic module eigengene | leave-out marker score, 7 non-member genes | residualised | 0.29 | 0.42 | 506 |
| Catabolic module eigengene | ClearCode34 ccB minus ccA score | residualised | -0.74 | -0.70 | 506 |
| Catabolic module eigengene | slide percent tumour nuclei | residualised | 0.21 | 0.25 | 506 |
| Catabolic module eigengene | non-feature read fraction | observed | 0.02 | 0.04 | 506 |
| lncRNA blue | proximal-tubule marker score | residualised | 0.75 | 0.74 | 511 |
| lncRNA blue | non-feature read fraction | observed | 0.44 | 0.44 | 511 |
| lncRNA greenyellow | proximal-tubule marker score | residualised | -0.44 | -0.52 | 511 |

| Feature | Variable | Expression space | Pearson r | Spearman rho | n |
|---|---|---|---|---|---|
| lncRNA greenyellow | non-feature read fraction | observed | 0.22 | 0.25 | 511 |
| lncRNA turquoise | proximal-tubule marker score | residualised | 0.04 | 0.03 | 511 |
| lncRNA turquoise | non-feature read fraction | observed | 0.85 | 0.90 | 511 |
| lncRNA turquoise | non-feature read fraction | residualised | 0.00 | 0.20 | 511 |
| Proximal-tubule marker score | non-feature read fraction | observed | 0.08 | 0.05 | 528 |

*Note.* TCGA-KIRC. Source: `34_pt_score_correlations.tsv`. The catabolic module and the marker score are nearly the same measurement. The module does not track the alignment metrics, whereas the three prognostic lncRNA modules do. Residualisation removes most but not all of that dependence for lncRNA turquoise, whose residualised eigengene still correlates with the metric at Spearman 0.20 (p 5.3e-06), against 0.029 (p 0.69) in CPTAC-3. In CPTAC-3 the module correlates with the marker score at 0.87 on 199 patients.

**(c) Separability of the module and the marker score**

| Model, clinical frame | Terms | HR (95% CI), p | n / events | Variance inflation |
|---|---|---|---|---|
| Marker score, 21 genes | pt_full | 0.69 (0.58 to 0.81), p 6.8e-06 | 520 / 172 | 1.07 |
| Leave-out marker score, 7 genes | pt_leaveout | 0.90 (0.75 to 1.09), p 0.29 | 520 / 172 | 1.05 |
| Catabolic module alone | green | 0.69 (0.59 to 0.81), p 2.9e-06 | 502 / 165 | 1.13 |
| Marker score on the module's analysis set | pt_full | 0.70 (0.59 to 0.82), p 2.6e-05 | 502 / 165 | 1.10 |
| Module and marker score together | green / pt_full | 0.74 (0.55 to 1.00), p 0.046 / 0.92 (0.67 to 1.25), p 0.58 | 502 / 165 | 4.05 / 3.98 |
| Module and lncRNA blue together | blue / green | 1.18 (0.79 to 1.77), p 0.41 / 0.61 (0.40 to 0.91), p 0.015 | 493 / 159 | 7.53 / 7.59 |
| Module and lncRNA turquoise together | turquoise / green | 1.34 (1.14 to 1.59), p 5.5e-04 / 0.68 (0.58 to 0.80), p 2.0e-06 | 493 / 159 | 1.15 / 1.17 |
| Module and lncRNA greenyellow together | greenyellow / green | 1.14 (0.94 to 1.40), p 0.19 / 0.76 (0.63 to 0.91), p 0.0035 | 493 / 159 | 1.83 / 1.58 |

Wald contrasts and nested likelihood-ratio tests:

| Test | Estimate (SE) or statistic | p | Pairwise r |
|---|---|---|---|
| Contrast, module minus marker score, clinical frame | -0.2105 (0.2949) | 0.48 | 0.84 |
| Contrast, module minus marker score, principal frame | -0.5617 (0.3385) | 0.097 | 0.84 |
| Contrast, lncRNA blue minus module, clinical frame | 0.6722 (0.4053) | 0.097 | 0.91 |
| Adding the module on top of the marker score, clinical frame | 3.801 | 0.051 | n/a |
| Adding the marker score on top of the module, clinical frame | 0.319 | 0.57 | n/a |
| Adding the module on top of the marker score, principal frame | 7.637 | 0.0057 | n/a |
| Adding the marker score on top of the module, principal frame | 0.078 | 0.78 | n/a |
| Adding lncRNA turquoise on top of the module | 11.509 | 6.9e-04 | n/a |
| Adding lncRNA greenyellow on top of the module | 1.700 | 0.19 | n/a |
| Adding lncRNA blue on top of the module | 0.683 | 0.41 | n/a |

*Note.* TCGA-KIRC, overall survival, hazard ratio per standard deviation. Source: `34_pt_adjusted_models.tsv`. The module eigengene and the marker score are statistically indistinguishable as predictors. They correlate at 0.84 with variance inflation of about 4, and the Wald contrast gives p 0.48, so the split of p 0.046 against p 0.58 in the joint model is not evidence of absorption. Whether the module adds to the marker score depends on the specification (nested-test p 0.051 under the clinical frame, p 0.0057 under the principal frame) and remains unresolved. The module and lncRNA blue are worse still, correlating at 0.91 with variance inflation of 7.5 (493 patients, 159 events), so no clean attribution is possible there either. Every CPTAC-3 proximal-tubule model is below the events-per-variable threshold of 5 and is descriptive.

**(d) ccA and ccB classification**

| Quantity | TCGA-KIRC | CPTAC-3 |
|---|---|---|
| Patients classified | 528 | 200 |
| ccB, n (%) | 219 (41.5) | 81 (40.5) |

| Quantity | TCGA-KIRC | CPTAC-3 |
|---|---|---|
| Agreement of the two published classifier genes | 98.5% | 99.0% |
| Mean catabolic eigengene, ccA against ccB | 0.492 against -0.753, p 1.5e-44 | 0.534 against -0.782, p 3.0e-20 |
| ccB, unadjusted HR | 2.33 (1.72 to 3.15), p 4.2e-08, 528 / 173 | 1.80 (0.89 to 3.61), p 0.10, 199 / 32 |
| ccB + clinical model | 1.62 (1.16 to 2.27), p 0.0052, 520 / 172 | 1.51 (0.74 to 3.11), p 0.26 |
| ccB + clinical model + the catabolic module | 1.03 (0.67 to 1.58), p 0.89, with the module at 0.70 (0.58 to 0.84) | 0.96 (0.36 to 2.57), p 0.94 |

*Note.* Source: `34_ccAB_class_summary.tsv`, `34_pt_adjusted_models.tsv`.

ClearCode34 is used as a template axis, not as the published nearest-centroid classifier. Of its 34 published genes, 31 are in the protein-coding network and 12 in the catabolic module, all 12 of them ccA genes, so the contrast between the class and the module is partly definitional.

**(e) Stromal and immune content of the catabolic module**

| Module | Genes | ESTIMATE stromal genes in the module | Expected | p | ESTIMATE immune genes | Expected | p |
|---|---|---|---|---|---|---|---|
| Catabolic (green) | 670 | 4 (0.6%) | 6.36 | 0.89 | 0 | 7.48 | 1.00 |
| black | 323 | 28 (8.7%) | 3.07 | 1.5e-19 | 0 | 3.61 | 1.00 |
| yellow | 986 | 40 (4.1%) | 9.37 | 4.6e-16 | 111 (11.3%) | 11.01 | 6.9e-99 |
| blue | 994 | 28 (2.8%) | 9.44 | 1.2e-07 | 11 (1.1%) | 11.10 | 0.56 |
| turquoise | 2,405 | 2 (0.1%) | 22.85 | 1.00 | 4 (0.2%) | 26.86 | 1.00 |

*Note.* Hypergeometric test against a universe of the 12,000 network genes, which include 114 of the 141 ESTIMATE stromal genes and 134 of the 141 immune genes. Source: `34_green_estimate_overlap.tsv`. The catabolic module carries fewer stromal genes than chance and no immune genes. The stromal and immune programmes sit in other modules.

**(f) Normal-kidney admixture in the catabolic module**

| Cohort | Feature | Expression space | Normal mean in tumour SD units (AUC) | Wilcoxon p | Normals above the tumour maximum | Confounded by the alignment metrics | Direction survives adjustment |
|---|---|---|---|---|---|---|---|
| TCGA-KIRC | Catabolic module | observed | 0.358 (0.587) | 0.017 | 0 of 72 | no | yes |
| TCGA-KIRC | Catabolic module | residualised | 0.313 (0.576) | 0.036 | 0 of 72 | no | yes |
| TCGA-KIRC | Proximal-tubule markers in the network | observed | 0.744 (0.663) | 7.8e-06 | 6 of 72 | no | yes |
| TCGA-KIRC | lncRNA blue | observed | -0.398 (0.359) | 1.1e-04 | 0 of 72 | yes | no |
| TCGA-KIRC | lncRNA blue | residualised | -0.053 (0.467) | 0.37 | 0 of 72 | no | no |
| TCGA-KIRC | lncRNA greenyellow | observed | -0.970 (0.159) | 7.8e-21 | 0 of 72 | yes | yes |
| TCGA-KIRC | lncRNA greenyellow | residualised | -0.758 (0.259) | 3.2e-11 | 0 of 72 | no | yes |
| TCGA-KIRC | lncRNA turquoise | observed | -0.896 (0.230) | 1.1e-13 | 0 of 72 | yes | no |
| TCGA-KIRC | lncRNA turquoise | residualised | -0.202 (0.441) | 0.11 | 0 of 72 | no | no |
| CPTAC-3 | Catabolic module | observed | 1.524 (0.902) | 2.5e-40 | 6 of 168 | no | yes |
| CPTAC-3 | Proximal-tubule markers in the network | observed | 2.109 (0.950) | 5.3e-50 | 119 of 168 | no | yes |
| CPTAC-3 | lncRNA blue | observed | 0.666 (0.740) | 2.4e-15 | 0 of 168 | yes | yes |
| CPTAC-3 | lncRNA greenyellow | observed | -1.069 (0.124) | 1.5e-35 | 0 of 168 | no | yes |
| CPTAC-3 | lncRNA turquoise | observed | 0.294 (0.615) | 1.5e-04 | 0 of 168 | yes | no |

*Note.* Scores are standardised to the tumour distribution, so tabulated values are mean normal-library scores in tumour standard deviations. Source: `34_normal_admixture.tsv`. In TCGA-KIRC, an entirely normal library sits 0.358 tumour standard deviations above the mean tumour on the catabolic module, on a tumour range of 5.07 standard deviations. No normal library exceeds the tumour maximum, so no mixing fraction can reach the top of the tumour distribution. The same cannot be asserted for CPTAC-3, where the protocol differs and 6 of 168 normals exceed the tumour maximum on the module eigengene and 119 of 168 on the marker score. The like-for-like marker-score figure in TCGA-KIRC is 6 of 72. Five rows, all on the observed matrix, are flagged as confounded by the alignment metrics. After residualisation the apparent opposite direction of lncRNA blue and turquoise in normal kidney disappears and only greenyellow survives. The file also gives the normal fraction needed for a one-standard-deviation shift. That column assumes the score is linear in the normal fraction, which it is not, and is flagged as anti-conservative, so the argument rests on the observed ceiling.

#### Supplementary Table 19. Single-cell localisation of the five prognostic modules

| Contrast | Network | Module | Cohort | Members tested | Median log2FC | Null mean | z | Empirical p | Members up (%) | Members down (%) |
|---|---|---|---|---|---|---|---|---|---|---|
| vs normal epithelium | lncRNA | blue | GSE159115 | 69 | -0.11 | -0.08 | -0.2 | 0.68 | 6 | 3 |
| vs normal epithelium | lncRNA | blue | GSE222703 | 96 | -0.32 | 0.23 | -2.8 | 0.0015 | 24 | 22 |
| vs normal epithelium | lncRNA | blue | Li2022 | 55 | 0.04 | 0.17 | -0.6 | 0.57 | 16 | 7 |
| vs normal epithelium | lncRNA | greenyellow | GSE159115 | 18 | 1.04 | -0.08 | 3.5 | 0.0025 | 11 | 0 |
| vs normal epithelium | lncRNA | greenyellow | GSE222703 | 14 | 4.38 | 0.40 | 7.2 | 0.0005 | 64 | 0 |
| vs normal epithelium | lncRNA | greenyellow | Li2022 | 14 | 2.86 | 0.14 | 6.4 | 0.0005 | 50 | 0 |
| vs normal epithelium | lncRNA | turquoise | GSE159115 | 60 | 0.24 | -0.10 | 2.1 | 0.049 | 7 | 0 |
| vs normal epithelium | lncRNA | turquoise | GSE222703 | 44 | 0.82 | 0.34 | 1.6 | 0.12 | 34 | 2 |
| vs normal epithelium | lncRNA | turquoise | Li2022 | 33 | 0.25 | 0.08 | 0.6 | 0.61 | 18 | 0 |
| vs normal epithelium | protein-coding | green | GSE159115 | 640 | -0.09 | -0.00 | -2.4 | 0.021 | 3 | 1 |
| vs normal epithelium | protein-coding | green | GSE222703 | 633 | -0.98 | -0.11 | -20.4 | 0.0005 | 9 | 42 |

| Contrast | Network | Module | Cohort | Members tested | Median log2FC | Null mean | z | Empirical p | Members up (%) | Members down (%) |
|---|---|---|---|---|---|---|---|---|---|---|
| vs normal epithelium | protein-coding | green | Li2022 | 627 | -0.27 | 0.24 | -12.6 | 0.0005 | 9 | 10 |
| vs normal epithelium | protein-coding | purple | GSE159115 | 143 | 0.88 | -0.02 | 9.5 | 0.0005 | 10 | 0 |
| vs normal epithelium | protein-coding | purple | GSE222703 | 102 | 0.69 | 0.07 | 5.4 | 0.0005 | 26 | 3 |
| vs normal epithelium | protein-coding | purple | Li2022 | 131 | 1.26 | 0.21 | 10.5 | 0.0005 | 24 | 1 |
| vs microenvironment | lncRNA | blue | GSE159115 | 118 | 3.60 | 0.42 | 22.4 | 0.0005 | 84 | 0 |
| vs microenvironment | lncRNA | blue | GSE207493 | 161 | 3.57 | 0.45 | 20.5 | 0.0005 | 89 | 1 |
| vs microenvironment | lncRNA | blue | GSE222703 | 89 | 5.04 | 0.32 | 15.6 | 0.0005 | 81 | 1 |
| vs microenvironment | lncRNA | blue | Li2022 | 85 | 5.33 | 0.38 | 16.3 | 0.0005 | 78 | 9 |
| vs microenvironment | lncRNA | greenyellow | GSE159115 | 49 | 1.73 | 0.44 | 5.6 | 0.0015 | 55 | 4 |
| vs microenvironment | lncRNA | greenyellow | GSE207493 | 69 | 1.51 | 0.45 | 4.4 | 0.0005 | 65 | 16 |
| vs microenvironment | lncRNA | greenyellow | GSE222703 | 21 | 1.56 | 0.50 | 1.4 | 0.16 | 43 | 10 |
| vs microenvironment | lncRNA | greenyellow | Li2022 | 28 | 1.89 | 0.27 | 3.3 | 0.011 | 54 | 21 |
| vs microenvironment | lncRNA | turquoise | GSE159115 | 164 | 0.32 | 0.41 | -0.7 | 0.49 | 22 | 5 |
| vs microenvironment | lncRNA | turquoise | GSE207493 | 243 | -0.04 | 0.43 | -3.7 | 0.0025 | 33 | 31 |
| vs microenvironment | lncRNA | turquoise | GSE222703 | 83 | -0.40 | 0.29 | -2.5 | 0.0075 | 8 | 22 |
| vs microenvironment | lncRNA | turquoise | Li2022 | 91 | -0.38 | 0.19 | -2.1 | 0.018 | 14 | 29 |
| vs microenvironment | protein-coding | green | GSE159115 | 661 | 2.29 | -0.02 | 55.2 | 0.0005 | 79 | 1 |
| vs microenvironment | protein-coding | green | GSE207493 | 668 | 2.63 | -0.01 | 53.4 | 0.0005 | 93 | 2 |
| vs microenvironment | protein-coding | green | GSE222703 | 601 | 3.21 | -0.14 | 46.7 | 0.0005 | 81 | 0 |
| vs microenvironment | protein-coding | green | Li2022 | 646 | 3.59 | -0.15 | 50.9 | 0.0005 | 85 | 3 |

| Contrast | Network | Module | Cohort | Members tested | Median log2FC | Null mean | z | Empirical p | Members up (%) | Members down (%) |
|---|---|---|---|---|---|---|---|---|---|---|
| vs microenvironment | protein-coding | purple | GSE159115 | 216 | -0.33 | -0.00 | -3.8 | 0.0015 | 6 | 25 |
| vs microenvironment | protein-coding | purple | GSE207493 | 216 | -0.31 | -0.01 | -3.0 | 0.0025 | 23 | 49 |
| vs microenvironment | protein-coding | purple | GSE222703 | 154 | -0.31 | -0.08 | -1.4 | 0.15 | 15 | 21 |
| vs microenvironment | protein-coding | purple | Li2022 | 211 | -0.52 | -0.17 | -2.6 | 0.011 | 13 | 31 |

Combined across cohorts, all 29 modules:

| Contrast | Network | Module | Prognostic | Cohorts | Mean median log2FC | Stouffer z | FDR | Cohorts above null |
|---|---|---|---|---|---|---|---|---|
| vs normal epithelium | lncRNA | greenyellow | yes | 3 | 2.76 | 9.9 | 1.3e-22 | 3 |
| vs normal epithelium | lncRNA | turquoise | yes | 3 | 0.44 | 2.4 | 0.022 | 3 |
| vs normal epithelium | lncRNA | blue | yes | 3 | -0.13 | -2.1 | 0.0498 | 0 |
| vs normal epithelium | protein-coding | purple | yes | 3 | 0.94 | 14.7 | 5.2e-48 | 3 |
| vs normal epithelium | protein-coding | green | yes | 3 | -0.45 | -20.4 | 3.7e-91 | 0 |
| vs normal epithelium | lncRNA | salmon | no | 3 | 2.20 | 6.3 | 5.8e-10 | 3 |
| vs normal epithelium | lncRNA | lightcyan | no | 3 | 1.70 | 5.4 | 1.1e-07 | 3 |
| vs normal epithelium | lncRNA | yellow | no | 3 | 2.00 | 5.3 | 2.1e-07 | 3 |
| vs normal epithelium | lncRNA | black | no | 3 | 0.58 | 2.0 | 0.053 | 2 |
| vs normal epithelium | lncRNA | green | no | 3 | 0.96 | 1.8 | 0.084 | 3 |
| vs normal epithelium | lncRNA | midnightblue | no | 3 | -0.08 | -0.5 | 0.65 | 1 |
| vs normal epithelium | lncRNA | purple | no | 3 | 0.00 | -0.7 | 0.57 | 1 |
| vs normal epithelium | lncRNA | cyan | no | 3 | -0.31 | -1.2 | 0.26 | 1 |

| Contrast | Network | Module | Prognostic | Cohorts | Mean median log2FC | Stouffer z | FDR | Cohorts above null |
|---|---|---|---|---|---|---|---|---|
| vs normal epithelium | lncRNA | pink | no | 3 | -0.14 | -2.3 | 0.035 | 0 |
| vs normal epithelium | lncRNA | red | no | 3 | -0.70 | -2.6 | 0.014 | 0 |
| vs normal epithelium | lncRNA | tan | no | 3 | -0.29 | -2.6 | 0.014 | 0 |
| vs normal epithelium | lncRNA | brown | no | 3 | -0.36 | -3.6 | 0.00072 | 0 |
| vs normal epithelium | lncRNA | magenta | no | 3 | -1.53 | -7.5 | 2.5e-13 | 0 |
| vs normal epithelium | protein-coding | yellow | no | 3 | 0.70 | 22.3 | 3e-108 | 3 |
| vs normal epithelium | protein-coding | tan | no | 3 | 0.58 | 10.8 | 3.4e-26 | 3 |
| vs normal epithelium | protein-coding | salmon | no | 3 | 0.99 | 10.1 | 2e-23 | 3 |
| vs normal epithelium | protein-coding | red | no | 3 | 0.38 | 8.7 | 1.2e-17 | 3 |
| vs normal epithelium | protein-coding | brown | no | 3 | 0.04 | -0.4 | 0.71 | 1 |
| vs normal epithelium | protein-coding | pink | no | 3 | 0.06 | -0.6 | 0.61 | 1 |
| vs normal epithelium | protein-coding | black | no | 3 | 0.15 | -1.1 | 0.32 | 1 |
| vs normal epithelium | protein-coding | blue | no | 3 | 0.03 | -3.0 | 0.0044 | 1 |
| vs normal epithelium | protein-coding | turquoise | no | 3 | -0.02 | -7.4 | 5e-13 | 0 |
| vs normal epithelium | protein-coding | magenta | no | 3 | -0.37 | -10.4 | 1.6e-24 | 0 |
| vs normal epithelium | protein-coding | greenyellow | no | 3 | -0.95 | -20.1 | 9.5e-89 | 0 |
| vs microenvironment | lncRNA | blue | yes | 4 | 4.39 | 37.4 | 4.7e-305 | 4 |
| vs microenvironment | lncRNA | greenyellow | yes | 4 | 1.67 | 7.3 | 5.1e-13 | 4 |
| vs microenvironment | lncRNA | turquoise | yes | 4 | -0.12 | -4.5 | 1e-05 | 0 |
| vs microenvironment | protein-coding | green | yes | 4 | 2.93 | 103.1 | 0 | 4 |

| Contrast | Network | Module | Prognostic | Cohorts | Mean median log2FC | Stouffer z | FDR | Cohorts above null |
|---|---|---|---|---|---|---|---|---|
| vs microenvironment | protein-coding | purple | yes | 4 | -0.37 | -5.4 | 9.1e-08 | 0 |
| vs microenvironment | lncRNA | lightcyan | no | 4 | 5.18 | 16.7 | 1.8e-62 | 4 |
| vs microenvironment | lncRNA | salmon | no | 4 | 2.95 | 8.5 | 2.8e-17 | 4 |
| vs microenvironment | lncRNA | pink | no | 4 | 0.66 | 3.1 | 0.0028 | 4 |
| vs microenvironment | lncRNA | tan | no | 4 | 0.76 | 2.8 | 0.0068 | 4 |
| vs microenvironment | lncRNA | cyan | no | 4 | 0.97 | 2.3 | 0.027 | 4 |
| vs microenvironment | lncRNA | purple | no | 4 | 0.05 | -1.1 | 0.26 | 1 |
| vs microenvironment | lncRNA | brown | no | 4 | 0.12 | -2.2 | 0.034 | 1 |
| vs microenvironment | lncRNA | magenta | no | 4 | -0.08 | -2.2 | 0.031 | 0 |
| vs microenvironment | lncRNA | midnightblue | no | 4 | -0.26 | -2.3 | 0.027 | 0 |
| vs microenvironment | lncRNA | green | no | 4 | -0.70 | -4.5 | 9.4e-06 | 0 |
| vs microenvironment | lncRNA | black | no | 4 | -0.94 | -7.9 | 4.4e-15 | 0 |
| vs microenvironment | lncRNA | red | no | 4 | -1.58 | -18.2 | 1.2e-73 | 0 |
| vs microenvironment | lncRNA | yellow | no | 4 | -3.74 | -35.7 | 4.2e-278 | 0 |
| vs microenvironment | protein-coding | brown | no | 4 | 0.38 | 17.6 | 1e-68 | 4 |
| vs microenvironment | protein-coding | tan | no | 4 | 0.36 | 4.2 | 3.8e-05 | 3 |
| vs microenvironment | protein-coding | magenta | no | 4 | -0.14 | -0.5 | 0.63 | 2 |
| vs microenvironment | protein-coding | salmon | no | 4 | -0.56 | -5.3 | 2.1e-07 | 1 |
| vs microenvironment | protein-coding | red | no | 4 | -0.42 | -7.3 | 4.1e-13 | 1 |
| vs microenvironment | protein-coding | turquoise | no | 4 | -0.27 | -10.0 | 2.7e-23 | 0 |

| Contrast | Network | Module | Prognostic | Cohorts | Mean median log2FC | Stouffer z | FDR | Cohorts above null |
|---|---|---|---|---|---|---|---|---|
| vs microenvironment | protein-coding | pink | no | 4 | -0.69 | -11.6 | 8e-31 | 0 |
| vs microenvironment | protein-coding | greenyellow | no | 4 | -1.24 | -19.2 | 2.2e-81 | 0 |
| vs microenvironment | protein-coding | black | no | 4 | -1.59 | -37.7 | 0 | 0 |
| vs microenvironment | protein-coding | blue | no | 4 | -1.56 | -67.7 | 0 | 0 |
| vs microenvironment | protein-coding | yellow | no | 4 | -2.98 | -127.4 | 0 | 0 |

*Note.* Cohorts: GSE159115 (discovery cohort of the single-cell pipeline), Li 2022, GSE222703 and GSE207493 (microenvironment contrast only, no normal kidney). Contrasts are donor-level pseudobulk edgeR quasi-likelihood fits from the outcome-blind single-cell pipeline. The null for each module is 2,000 random same-size sets from the same gene class of the network universe, among genes tested in that cohort, matched on deciles of log10 mean counts per million. The empirical p is two-sided with a floor of 1/2,000. Members up or down are the percentage of tested members with an upstream false discovery rate below 0.05 over all genes. The combined false discovery rate is controlled across the 29 modules within each contrast. Combined z scores overstate precision because genes within a module are correlated. Member coverage in at least one cohort is in `36_sc_module_gene_coverage.tsv`. Against normal epithelium it is 25% for greenyellow, 44% for blue, 18% for turquoise, 97% for green and 74% for purple. Source tables: `36_sc_module_localisation_per_dataset.tsv`, `36_sc_module_localisation_combined.tsv`, `36_sc_module_lncRNA_pooled_meta_summary.tsv`.

#### Supplementary Table 20. The locked prediction model

**(a) Coefficients**

| Term | Clinical comparator, coefficient (HR) | Clinical combined, coefficient (HR) | Augmented comparator, coefficient (HR) | Augmented combined, coefficient (HR) |
|---|---|---|---|---|
| Age, standardised | 0.4011 (1.49) | 0.4095 (1.51) | 0.3936 (1.48) | 0.3927 (1.48) |
| Male | -0.0171 (0.98) | -0.1383 (0.87) | 0.0361 (1.04) | -0.0926 (0.91) |
| T category, 1 to 4 | 0.1418 (1.15) | 0.1203 (1.13) | 0.1274 (1.14) | 0.1009 (1.11) |
| Node positive | 0.8099 (2.25) | 0.7823 (2.19) | 0.6734 (1.96) | 0.6604 (1.94) |
| M1 | 1.0394 (2.83) | 1.0747 (2.93) | 1.0343 (2.81) | 1.0890 (2.97) |
| Ordinal grade, 1 to 4 | 0.3966 (1.49) | 0.2886 (1.33) | 0.4979 (1.65) | 0.3532 (1.42) |
| ESTIMATE stromal, standardised | n/a | n/a | -0.0148 (0.99) | -0.0976 (0.91) |
| ESTIMATE immune, standardised | n/a | n/a | -0.0785 (0.92) | -0.1072 (0.90) |
| Non-feature fraction, standardised | n/a | n/a | 0.2930 (1.34) | 0.3141 (1.37) |
| Multimapping fraction, standardised | n/a | n/a | 0.0661 (1.07) | 0.0977 (1.10) |
| log10 assigned reads, standardised | n/a | n/a | -0.0179 (0.98) | -0.0026 (1.00) |
| Protein-coding black eigengene | n/a | -0.0589 (0.94) | n/a | n/a |
| Protein-coding blue eigengene | n/a | -0.0096 (0.99) | n/a | n/a |
| Protein-coding green eigengene | n/a | -0.2774 (0.76) | n/a | -0.3285 (0.72) |
| Protein-coding pink eigengene | n/a | -0.0558 (0.95) | n/a | -0.1263 (0.88) |
| Protein-coding yellow eigengene | n/a | -0.1255 (0.88) | n/a | n/a |
| lncRNA black eigengene | n/a | 0.0398 (1.04) | n/a | 0.0729 (1.08) |
| lncRNA blue eigengene | n/a | -0.1171 (0.89) | n/a | -0.0486 (0.95) |
| lncRNA midnightblue eigengene | n/a | -0.0202 (0.98) | n/a | n/a |
| lncRNA turquoise eigengene | n/a | 0.1982 (1.22) | n/a | 0.1106 (1.12) |

*Note.* Fitted on the 493 TCGA-KIRC patients of the locked prediction set, with 159 events.

Source: `22_locked_model_coefficients.tsv`. Comparator terms are unpenalised. The 29 module eigengenes are offered to an elastic net (mixing parameter 0.5) with the penalty at the minimum of ten-fold cross-validated partial likelihood. The chosen penalty is 0.0254 for the clinical arm (9 module scores retained) and 0.0220 for the augmented arm (5 retained). Coefficients are per standard deviation of the standardised covariate or module score. Neither arm retains lncRNA greenyellow, the lncRNA module with the strongest external estimate. A separate full-cohort reporting fit in `05_final_model_coefficients.tsv` retains the same nine module scores at a slightly different penalty, with every coefficient different. Validation, calibration, Brier scores, decision curves and the endpoint analyses all use the model tabulated here.

**(b) Module score definitions and retention**

| Module score | Genes | Variance explained by PC1 | Score scale | Retained, clinical arm | Retained, augmented arm |
|---|---|---|---|---|---|
| protein-coding black | 323 | 0.4347 | 11.849 | yes | no |
| protein-coding blue | 994 | 0.3936 | 19.779 | yes | no |
| protein-coding brown | 988 | 0.3815 | 19.414 | no | no |
| protein-coding green | 670 | 0.3624 | 15.582 | yes | yes |
| protein-coding greenyellow | 164 | 0.4506 | 8.597 | no | no |
| protein-coding magenta | 227 | 0.4646 | 10.270 | no | no |
| protein-coding pink | 270 | 0.4521 | 11.048 | yes | yes |
| protein-coding purple | 216 | 0.4467 | 9.823 | no | no |
| protein-coding red | 473 | 0.3849 | 13.494 | no | no |
| protein-coding salmon | 63 | 0.4601 | 5.384 | no | no |
| protein-coding tan | 152 | 0.4813 | 8.553 | no | no |
| protein-coding turquoise | 2,405 | 0.3649 | 29.622 | no | no |
| protein-coding yellow | 986 | 0.4009 | 19.881 | yes | no |
| lncRNA black | 179 | 0.3408 | 7.810 | yes | yes |
| lncRNA blue | 246 | 0.2765 | 8.247 | yes | yes |
| lncRNA brown | 219 | 0.2654 | 7.624 | no | no |
| lncRNA cyan | 54 | 0.2912 | 3.966 | no | no |
| lncRNA green | 186 | 0.2614 | 6.972 | no | no |

| Module score | Genes | Variance explained by PC1 | Score scale | Retained, clinical arm | Retained, augmented arm |
|---|---|---|---|---|---|
| lncRNA greenyellow | 108 | 0.2505 | 5.201 | no | no |
| lncRNA lightcyan | 44 | 0.3138 | 3.716 | no | no |
| lncRNA magenta | 122 | 0.3070 | 6.120 | no | no |
| lncRNA midnightblue | 53 | 0.2893 | 3.916 | yes | no |
| lncRNA pink | 174 | 0.3057 | 7.294 | no | no |
| lncRNA purple | 110 | 0.3255 | 5.983 | no | no |
| lncRNA red | 186 | 0.2994 | 7.462 | no | no |
| lncRNA salmon | 72 | 0.2992 | 4.642 | no | no |
| lncRNA tan | 93 | 0.2983 | 5.267 | no | no |
| lncRNA turquoise | 438 | 0.2655 | 10.784 | yes | yes |
| lncRNA yellow | 192 | 0.3065 | 7.671 | no | no |

*Note.* Source: `22_module_score_constants.tsv`. A module eigengene is the first principal component of the module's gene-standardised residualised expression, sign-aligned to mean module expression and divided by the score scale to give unit variance in discovery. The score centre is numerically zero in every row (of order 1e-16). Thirteen protein-coding and 16 lncRNA modules give 29 scores. The per-gene loadings, gene means and gene standard deviations that define each score are in Supplementary Data 2.

**(c) Standardisation constants of the comparator covariates, discovery cohort**

| Covariate | Source | Mean | SD |
|---|---|---|---|
| Age, years | clinical record | 61.1754 | 12.0387 |
| ESTIMATE stromal score | ESTIMATE | 842.4403 | 476.0470 |
| ESTIMATE immune score | ESTIMATE | 1683.9339 | 640.6478 |
| Non-feature read fraction, % | STAR summary | 5.1418 | 2.5660 |
| Multimapping fraction, % | STAR summary | 7.8333 | 1.7663 |
| log10 assigned reads | STAR summary | 7.7592 | 0.1078 |

*Note.* Source: `22_locked_model_standardisation.tsv`. These constants define the discovery-fixed arm of the eigengene standardisation. They are not applied to the CPTAC-3 comparator covariates, which are z-scored within CPTAC-3 in both arms. Because of the protocol difference, the CPTAC-3 non-feature fraction lies far outside the discovery range.

**(d) Discovery baseline survival**

| Comparator | Model | Mean discovery linear predictor | S0 at 1, 3, 5 years | S at the mean linear predictor, 1, 3, 5 years | Apparent C |
|---|---|---|---|---|---|
| clinical | comparator only | 1.4891 | 0.9838 / 0.9551 / 0.9240 | 0.9300 / 0.8156 / 0.7043 | 0.766 |
| clinical | combined | 1.0812 | 0.9777 / 0.9367 / 0.8907 | 0.9356 / 0.8247 / 0.7108 | 0.787 |
| augmented | comparator only | 1.7604 | 0.9885 / 0.9675 / 0.9436 | 0.9352 / 0.8250 / 0.7134 | 0.775 |
| augmented | combined | 1.2453 | 0.9825 / 0.9491 / 0.9102 | 0.9405 / 0.8342 / 0.7211 | 0.792 |

*Note.* Source: `22_locked_model_baseline.tsv`, 493 patients and 159 events. Absolute risk at time t is 1 minus exp(-H0(t) times exp(LP - mean LP)), where H0 is the Breslow baseline cumulative hazard estimated in discovery and LP is the locked linear predictor. No clinical test is proposed. The model is reported in full for evaluation, not for use. Its primary arm standardises the eigengenes within the scored cohort, so it cannot score a single patient. The discovery-fixed arm can score a single patient, at the cost of the two-fold over-prediction in panel (b) of Supplementary Table 21.

#### Supplementary Table 21. External validation: discrimination, standardisation, calibration, Brier scores and clinical utility

**(a) Concordance index of the locked model**

| Cohort | Comparator | Model | Standardisation | n | Events | C (95% CI) |
|---|---|---|---|---|---|---|
| TCGA-KIRC (apparent) | clinical | comparator only | cohort | 493 | 159 | 0.766 (0.729 to 0.804) |
| TCGA-KIRC (apparent) | clinical | + module eigengenes | cohort | 493 | 159 | 0.787 (0.752 to 0.822) |

| Cohort | Comparator | Model | Standardisation | n | Events | C (95% CI) |
|---|---|---|---|---|---|---|
| CPTAC-3 | clinical | comparator only | cohort | 199 | 32 | 0.750 (0.657 to 0.843) |
| CPTAC-3 | clinical | + module eigengenes | cohort | 199 | 32 | 0.759 (0.671 to 0.847) |
| CPTAC-3 | clinical | comparator only | discovery-fixed | 199 | 32 | 0.750 (0.657 to 0.843) |
| CPTAC-3 | clinical | + module eigengenes | discovery-fixed | 199 | 32 | 0.754 (0.663 to 0.844) |
| TCGA-KIRC (apparent) | augmented | comparator only | cohort | 493 | 159 | 0.775 (0.738 to 0.812) |
| TCGA-KIRC (apparent) | augmented | + module eigengenes | cohort | 493 | 159 | 0.792 (0.757 to 0.826) |
| CPTAC-3 | augmented | comparator only | cohort | 199 | 32 | 0.730 (0.642 to 0.818) |
| CPTAC-3 | augmented | + module eigengenes | cohort | 199 | 32 | 0.745 (0.650 to 0.839) |

*Note.* Source: `09_validation_cindex.tsv`. The two TCGA-KIRC rows per comparator are apparent, scoring the locked model on the patients it was fitted to. Cohort standardisation, the primary arm, standardises the module eigengenes within the scored cohort. Discovery-fixed standardisation applies the discovery gene-level and score-level constants. In both arms the clinical, ESTIMATE and STAR covariates are standardised within CPTAC-3, and CPTAC-3 expression is residualised on CPTAC-3's own STAR metrics, so the discovery-fixed arm fixes only the eigengene constants.

**(b) Calibration**

| Cohort | Comparator | Standardisation | Model | Calibration slope (95% CI) |
|---|---|---|---|---|
| TCGA-KIRC (apparent) | clinical | cohort | comparator | 1.00 (0.83 to 1.17) |
| TCGA-KIRC (apparent) | clinical | cohort | comparator + modules | 1.05 (0.89 to 1.22) |
| CPTAC-3 | clinical | cohort | comparator | 0.97 (0.60 to 1.34) |
| CPTAC-3 | clinical | cohort | comparator + modules | 0.89 (0.58 to 1.20) |
| CPTAC-3 | clinical | discovery-fixed | comparator | 0.97 (0.61 to 1.34) |

| Cohort | Comparator | Standardisation | Model | Calibration slope (95% CI) |
|---|---|---|---|---|
| CPTAC-3 | clinical | discovery-fixed | comparator + modules | 0.89 (0.58 to 1.20) |
| TCGA-KIRC (apparent) | augmented | cohort | comparator | 1.00 (0.84 to 1.16) |
| TCGA-KIRC (apparent) | augmented | cohort | comparator + modules | 1.04 (0.88 to 1.19) |
| CPTAC-3 | augmented | cohort | comparator | 0.90 (0.51 to 1.28) |
| CPTAC-3 | augmented | cohort | comparator + modules | 0.84 (0.51 to 1.16) |

Observed against expected risk at each horizon:

| Cohort | Comparator | Standardisation | Model | 1 year, expected (O/E) | 3 years, expected (O/E) | 5 years, expected (O/E) |
|---|---|---|---|---|---|---|
| TCGA-KIRC | clinical | cohort | comparator | 0.094 (1.00) | 0.225 (1.03) | 0.335 (1.08) |
| TCGA-KIRC | clinical | cohort | + modules | 0.095 (0.99) | 0.226 (1.02) | 0.337 (1.08) |
| CPTAC-3 | clinical | cohort | comparator | 0.079 (0.71) | 0.192 (0.75) | 0.292 (0.90) |
| CPTAC-3 | clinical | cohort | + modules | 0.083 (0.68) | 0.194 (0.74) | 0.295 (0.89) |
| CPTAC-3 | clinical | discovery-fixed | comparator | 0.079 (0.71) | 0.191 (0.76) | 0.290 (0.91) |
| CPTAC-3 | clinical | discovery-fixed | + modules | 0.115 (0.49) | 0.260 (0.55) | 0.384 (0.68) |
| CPTAC-3 | augmented | cohort | comparator | 0.073 (0.77) | 0.182 (0.79) | 0.283 (0.93) |
| CPTAC-3 | augmented | cohort | + modules | 0.075 (0.75) | 0.184 (0.79) | 0.284 (0.93) |

*Note.* Source: `12_calibration_summary.tsv`. The calibration slope, the coefficient of the linear predictor in a Cox model of the outcome on that predictor, does not depend on the horizon, so one slope is reported per arm. Observed risk is the cohort-wide Kaplan-Meier estimate at the horizon: 0.094, 0.231 and 0.363 in TCGA-KIRC and 0.056, 0.144 and 0.263 in CPTAC-3. Patients at risk at 1, 3 and 5 years number 408, 275 and 141 in TCGA-KIRC and 144, 95 and 23 in CPTAC-3. The discovery TCGA-KIRC rows are apparent, and the comparator slope of exactly 1.00 there is an artefact of scoring a model on its training data. Both models over-predict risk in CPTAC-3, a lower-risk cohort, and under discovery-fixed standardisation the combined model over-predicts roughly two-fold.

**(c) Inverse-probability-of-censoring-weighted Brier scores, lower is better**

| Cohort | Comparator | Standardisation | Years | Comparator | Comparator + modules | Kaplan-Meier null |
|---|---|---|---|---|---|---|
| TCGA-KIRC | clinical | cohort | 1 | 0.0715 | 0.0675 | 0.0850 |
| TCGA-KIRC | clinical | cohort | 3 | 0.1443 | 0.1386 | 0.1779 |
| TCGA-KIRC | clinical | cohort | 5 | 0.1935 | 0.1795 | 0.2319 |
| CPTAC-3 | clinical | cohort | 1 | 0.0578 | 0.0607 | 0.0536 |
| CPTAC-3 | clinical | cohort | 3 | 0.0995 | 0.0979 | 0.1260 |
| CPTAC-3 | clinical | cohort | 5 | 0.1392 | 0.1455 | 0.1950 |
| CPTAC-3 | clinical | discovery-fixed | 1 | 0.0576 | 0.0671 | 0.0564 |
| CPTAC-3 | clinical | discovery-fixed | 3 | 0.0994 | 0.1134 | 0.1370 |
| CPTAC-3 | clinical | discovery-fixed | 5 | 0.1394 | 0.1625 | 0.2086 |
| TCGA-KIRC | augmented | cohort | 1 | 0.0711 | 0.0669 | 0.0850 |
| TCGA-KIRC | augmented | cohort | 3 | 0.1421 | 0.1385 | 0.1779 |
| TCGA-KIRC | augmented | cohort | 5 | 0.1826 | 0.1697 | 0.2321 |
| CPTAC-3 | augmented | cohort | 1 | 0.0564 | 0.0574 | 0.0533 |
| CPTAC-3 | augmented | cohort | 3 | 0.1073 | 0.1041 | 0.1251 |
| CPTAC-3 | augmented | cohort | 5 | 0.1384 | 0.1417 | 0.1944 |

*Note.* TCGA-KIRC, 493 patients and 159 events. CPTAC-3, 199 patients and 32 events. Source: `12_brier_scores.tsv`. The null model assigns every patient the cohort Kaplan-Meier risk at the horizon. The TCGA-KIRC rows are apparent. In CPTAC-3 the combined model is worse than the comparator at 1 and 5 years under cohort standardisation and at every horizon under discovery-fixed standardisation. At 1 year both models are worse than the null.

**(d) Decision curves in CPTAC-3, baseline hazard fixed in discovery**

| Comparator | Standardisation | Years | Threshold window | Thresholds | Median net-benefit gain | Positive thresholds | Bootstrap 95% CI | p |
|---|---|---|---|---|---|---|---|---|
| clinical | cohort | 3 | 0.10 to 0.40 | 31 | -0.0037 | 9 | -0.0223 to 0.0061 | 0.42 |

| Comparator | Standardisation | Years | Threshold window | Thresholds | Median net-benefit gain | Positive thresholds | Bootstrap 95% CI | p |
|---|---|---|---|---|---|---|---|---|
| clinical | cohort | 3 | 0.05 to 0.30 | 26 | -0.0005 | 11 | -0.0162 to 0.0061 | 0.87 |
| clinical | cohort | 5 | 0.10 to 0.40 | 31 | 0.0026 | 19 | -0.0183 to 0.0219 | 0.85 |
| clinical | cohort | 5 | 0.05 to 0.30 | 26 | 0.0016 | 16 | -0.0075 to 0.0133 | 0.74 |
| clinical | discovery-fixed | 3 | 0.10 to 0.40 | 31 | -0.0268 | 3 | -0.0421 to -0.0004 | 0.048 |
| clinical | discovery-fixed | 3 | 0.05 to 0.30 | 26 | -0.0147 | 5 | -0.0322 to 0.0010 | 0.10 |
| clinical | discovery-fixed | 5 | 0.10 to 0.40 | 31 | -0.0200 | 2 | -0.0570 to 0.0104 | 0.22 |
| clinical | discovery-fixed | 5 | 0.05 to 0.30 | 26 | -0.0071 | 2 | -0.0286 to 0.0024 | 0.26 |
| augmented | cohort | 3 | 0.10 to 0.40 | 31 | 0.0043 | 19 | -0.0083 to 0.0147 | 0.50 |
| augmented | cohort | 3 | 0.05 to 0.30 | 26 | -0.0007 | 12 | -0.0125 to 0.0087 | 0.80 |
| augmented | cohort | 5 | 0.10 to 0.40 | 31 | -0.0026 | 14 | -0.0217 to 0.0174 | 0.84 |
| augmented | cohort | 5 | 0.05 to 0.30 | 26 | -0.0009 | 11 | -0.0127 to 0.0112 | 0.82 |

*Note.* CPTAC-3, 199 patients and 32 events. Source: `12_validation_decision_curve_summary.tsv`. The median gain is the median, across the threshold window, of the net-benefit difference between the combined model and the comparator, with a paired patient bootstrap interval over 500 resamples. Every decision curve uses the discovery baseline hazard, and no arm re-estimates it in CPTAC-3. The only interval excluding zero does so in the harmful direction.

**(e) Drop-one-module sensitivity of the external increment, CPTAC-3**

| Comparator and standardisation | Model | C | Increment | 95% CI | p |
|---|---|---|---|---|---|
| clinical, cohort | all retained modules, 9 | 0.759 | 0.0087 | -0.0331 to 0.0478 | 0.68 |
| clinical, cohort | minus protein-coding black | 0.757 | 0.0068 | -0.0357 to 0.0466 | 0.77 |
| clinical, cohort | minus protein-coding blue | 0.758 | 0.0081 | -0.0320 to 0.0465 | 0.71 |

| Comparator and standardisation | Model | C | Increment | 95% CI | p |
|---|---|---|---|---|---|
| clinical, cohort | minus protein-coding green | 0.749 | -0.0013 | -0.0339 to 0.0320 | 0.96 |
| clinical, cohort | minus protein-coding pink | 0.755 | 0.0050 | -0.0368 to 0.0438 | 0.84 |
| clinical, cohort | minus protein-coding yellow | 0.746 | -0.0042 | -0.0480 to 0.0370 | 0.82 |
| clinical, cohort | minus lncRNA black | 0.758 | 0.0084 | -0.0304 to 0.0456 | 0.66 |
| clinical, cohort | minus lncRNA blue | 0.761 | 0.0113 | -0.0258 to 0.0471 | 0.56 |
| clinical, cohort | minus lncRNA midnightblue | 0.758 | 0.0081 | -0.0336 to 0.0464 | 0.70 |
| clinical, cohort | minus lncRNA turquoise | 0.754 | 0.0037 | -0.0361 to 0.0447 | 0.87 |
| clinical, cohort | minus all lncRNA modules | 0.760 | 0.0105 | -0.0192 to 0.0409 | 0.49 |
| clinical, discovery-fixed | all retained modules | 0.754 | 0.0039 | -0.0383 to 0.0433 | 0.88 |
| clinical, discovery-fixed | minus lncRNA blue | 0.762 | 0.0121 | -0.0255 to 0.0478 | 0.53 |
| clinical, discovery-fixed | minus lncRNA turquoise | 0.751 | 0.0010 | -0.0411 to 0.0447 | 0.98 |
| clinical, discovery-fixed | minus all lncRNA modules | 0.762 | 0.0115 | -0.0176 to 0.0419 | 0.43 |
| augmented, cohort | all retained modules, 5 | 0.745 | 0.0147 | -0.0192 to 0.0478 | 0.37 |
| augmented, cohort | minus protein-coding green | 0.728 | -0.0021 | -0.0256 to 0.0208 | 0.87 |
| augmented, cohort | minus protein-coding pink | 0.741 | 0.0105 | -0.0269 to 0.0463 | 0.58 |
| augmented, cohort | minus lncRNA black | 0.750 | 0.0197 | -0.0130 to 0.0518 | 0.23 |
| augmented, cohort | minus lncRNA blue | 0.745 | 0.0152 | -0.0167 to 0.0459 | 0.33 |
| augmented, cohort | minus lncRNA turquoise | 0.744 | 0.0142 | -0.0242 to 0.0503 | 0.44 |
| augmented, cohort | minus all lncRNA modules | 0.743 | 0.0126 | -0.0227 to 0.0461 | 0.46 |

*Note.* CPTAC-3, 199 patients and 32 events, paired patient bootstrap over 2,000 resamples. Source: `20_bootstrap_delta_cindex.tsv`, which also holds the remaining drop-one rows under discovery-fixed standardisation. The comparator concordance index is fixed at 0.750 for the clinical arm and 0.730 for the augmented arm. No single module carries the external increment. Removing every lncRNA module leaves the increment as large or larger, so the small external increment is protein-coding. Every interval spans zero.

**(f) Agreement between the two standardisations, per module eigengene, in CPTAC-3**

| Module score | Pearson r | Spearman rho | Mean under discovery-fixed standardisation | SD under discovery-fixed standardisation |
|---|---|---|---|---|
| protein-coding black | 0.9999 | 0.9998 | -0.06 | 0.95 |
| protein-coding blue | 0.9998 | 0.9996 | -0.14 | 0.88 |

| Module score | Pearson r | Spearman rho | Mean under discovery-fixed standardisation | SD under discovery-fixed standardisation |
|---|---|---|---|---|
| protein-coding brown | 0.9997 | 0.9992 | -1.47 | 0.45 |
| protein-coding green | 0.9999 | 0.9998 | 0.23 | 0.98 |
| protein-coding greenyellow | 0.9996 | 0.9992 | 2.27 | 0.78 |
| protein-coding magenta | 0.9997 | 0.9993 | 1.57 | 0.88 |
| protein-coding pink | 0.9979 | 0.9968 | 3.09 | 0.66 |
| protein-coding purple | 0.9999 | 0.9995 | 0.32 | 0.94 |
| protein-coding red | 0.9991 | 0.9981 | 1.01 | 0.80 |
| protein-coding salmon | 0.9999 | 0.9995 | -0.66 | 0.94 |
| protein-coding tan | 0.9997 | 0.9995 | -1.07 | 1.07 |
| protein-coding turquoise | 0.9993 | 0.9983 | 2.87 | 0.45 |
| protein-coding yellow | 0.9999 | 0.9998 | 0.66 | 0.89 |
| lncRNA black | 0.9974 | 0.9959 | 5.35 | 0.90 |
| lncRNA blue | 0.9988 | 0.9975 | 2.37 | 1.24 |
| lncRNA brown | 0.9977 | 0.9957 | 1.95 | 0.72 |
| lncRNA cyan | 0.9961 | 0.9954 | 3.85 | 1.19 |
| lncRNA green | 0.9972 | 0.9958 | 11.65 | 0.90 |
| lncRNA greenyellow | 0.9973 | 0.9964 | 1.43 | 1.12 |
| lncRNA lightcyan | 0.9860 | 0.9828 | 2.32 | 1.45 |
| lncRNA magenta | 0.9973 | 0.9929 | 2.41 | 0.98 |
| lncRNA midnightblue | 0.9864 | 0.9734 | 6.63 | 0.61 |
| lncRNA pink | 0.9978 | 0.9969 | 1.68 | 0.89 |
| lncRNA purple | 0.9933 | 0.9918 | 1.35 | 0.49 |
| lncRNA red | 0.9986 | 0.9986 | 2.59 | 1.30 |
| lncRNA salmon | 0.9930 | 0.9933 | 3.92 | 1.29 |
| lncRNA tan | 0.9968 | 0.9950 | 2.79 | 1.06 |
| lncRNA turquoise | 0.9978 | 0.9971 | 4.64 | 0.83 |
| lncRNA yellow | 0.9989 | 0.9985 | 3.00 | 1.26 |

*Note.* CPTAC-3, 199 patients. Source: `20_score_spread_by_standardisation.tsv`. The two standardisations rank patients almost identically (Pearson correlation at least 0.986 for every module), but the discovery constants displace the CPTAC-3 means by up to 11.7 standard deviations. This offset is why discovery-fixed standardisation barely changes the concordance index in panel (a) yet destroys calibration in panel (b).

**(g) External association estimates in CPTAC-3 under the two standardisations**

| Module | HR, cohort standardisation (95% CI) | FDR | HR, discovery-fixed standardisation (95% CI) | FDR |
|---|---|---|---|---|
| Protein-coding green | 0.73 (0.51 to 1.04) | 0.14 | 0.73 (0.52 to 1.04) | 0.14 |
| Protein-coding purple | 1.52 (1.08 to 2.14) | 0.042 | 1.52 (1.08 to 2.14) | 0.042 |
| lncRNA blue | 0.78 (0.55 to 1.11) | 0.21 | 0.78 (0.54 to 1.11) | 0.20 |
| lncRNA greenyellow | 1.59 (1.12 to 2.25) | 0.042 | 1.59 (1.12 to 2.25) | 0.042 |
| lncRNA turquoise | 1.11 (0.81 to 1.53) | 0.51 | 1.12 (0.81 to 1.54) | 0.49 |

*Note.* CPTAC-3, 199 patients and 32 events, twelve-parameter principal specification, module membership and eigengene loadings fixed in discovery. Source: `20_replication_unitmatched.tsv`. Benjamini-Hochberg control is across the five modules within each arm. The two standardisations differ by at most 0.006 in the hazard ratio, for lncRNA turquoise. This panel repeats the association analysis of Table 2b of the main text, whereas panels (a) to (f) evaluate the fixed prediction rule.

#### Supplementary Table 22. Specimen handling time in CPTAC-3

**(a) Spearman correlation of handling time with the alignment metrics, the projected lncRNA axis and tumour severity**

| Sample set | Handling time | Variable | Expected direction | n | Spearman rho (95% CI) | p | FDR |
|---|---|---|---|---|---|---|---|
| CPTAC-3 tumours | excision to freezing | Non-feature fraction | positive | 175 | -0.02 (-0.16 to 0.12) | 0.79 | 0.79 |
| CPTAC-3 tumours | excision to freezing | Projected lncRNA axis | positive | 175 | -0.09 (-0.23 to 0.06) | 0.25 | 0.43 |
| CPTAC-3 tumours | excision to freezing | Multimapping fraction | n/a | 175 | -0.04 (-0.18 to 0.10) | 0.62 | 0.72 |
| CPTAC-3 tumours | excision to freezing | log10 assigned reads | n/a | 175 | -0.04 (-0.19 to 0.10) | 0.55 | 0.72 |

| Sample set | Handling time | Variable | Expected direction | n | Spearman rho (95% CI) | p | FDR |
|---|---|---|---|---|---|---|---|
| CPTAC-3 tumours | excision to freezing | T category | positive | 174 | 0.14 (-0.02 to 0.28) | 0.06 | 0.14 |
| CPTAC-3 tumours | excision to freezing | Ordinal stage | positive | 175 | 0.15 (0.00 to 0.29) | 0.04 | 0.14 |
| CPTAC-3 tumours | excision to freezing | Ordinal grade | n/a | 175 | 0.32 (0.18 to 0.44) | 1.9e-05 | 1.3e-04 |
| CPTAC-3 tumours | clamping to freezing | Non-feature fraction | positive | 35 | 0.16 (-0.20 to 0.50) | 0.36 | 0.82 |
| CPTAC-3 tumours | clamping to freezing | Projected lncRNA axis | positive | 35 | -0.26 (-0.56 to 0.07) | 0.13 | 0.82 |
| CPTAC-3 tumours | clamping to freezing | Multimapping fraction | n/a | 35 | -0.08 (-0.43 to 0.28) | 0.66 | 0.82 |
| CPTAC-3 tumours | clamping to freezing | log10 assigned reads | n/a | 35 | 0.08 (-0.27 to 0.38) | 0.67 | 0.82 |
| CPTAC-3 tumours | clamping to freezing | T category | positive | 35 | -0.07 (-0.39 to 0.26) | 0.70 | 0.82 |
| CPTAC-3 tumours | clamping to freezing | Ordinal stage | positive | 35 | -0.11 (-0.42 to 0.23) | 0.54 | 0.82 |
| CPTAC-3 tumours | clamping to freezing | Ordinal grade | n/a | 35 | 0.00 (-0.28 to 0.30) | 0.99 | 0.99 |
| CPTAC-3 normal tissue | excision to freezing | Non-feature fraction | positive | 168 | -0.10 (-0.25 to 0.06) | 0.21 | 0.41 |
| CPTAC-3 normal tissue | excision to freezing | Projected lncRNA axis | positive | 168 | 0.01 (-0.14 to 0.16) | 0.93 | 0.93 |
| CPTAC-3 normal tissue | excision to freezing | Multimapping fraction | n/a | 168 | 0.07 (-0.09 to 0.21) | 0.38 | 0.51 |
| CPTAC-3 normal tissue | excision to freezing | log10 assigned reads | n/a | 168 | 0.24 (0.09 to 0.38) | 0.002 | 0.008 |

**(b) Handling time and overall survival, CPTAC-3 tumours**

| Model | n | Events | HR per SD (95% CI) | p | Events per variable |
|---|---|---|---|---|---|
| excision-to-freezing time alone | 175 | 23 | 1.45 (0.96 to 2.19) | 0.08 | 23.0 |
| + age, sex, T category, M1, ordinal grade | 174 | 23 | 1.18 (0.76 to 1.83) | 0.45 | 3.8 |

*Note.* Handling times are the Genomic Data Commons sample fields for the time from excision and from clamping to freezing, in minutes, taken from the sample whose library was sequenced. Excision to freezing is recorded for 175 of the 200 validation tumours (median 16, range 3 to 30, the CPTAC protocol cap) and all 168 normal-tissue libraries, and clamping to freezing for 35 tumours. Intervals are from 2,000 bootstrap resamples, and Benjamini-Hochberg control is within each sample set and handling-time variable. Median excision-to-freezing time is 12, 14, 17 and 21 minutes for grades 1 to 4. The 25 tumours without a recorded time account for 9 of the 32 deaths, so the survival estimates in panel (b) rest on 23 events. Source tables: `45_cptac_handling_time_correlations.tsv`, `45_cptac_handling_time_survival.tsv`, `45_cptac_handling_time_per_sample.tsv`.

### Supplementary Data

Five Supplementary Data items, supplied as tab-separated files, accompany this Supplementary Information.

#### Supplementary Data 1. Case list and file manifest

`SupplementaryData1_case_list.tsv`, 1,549 rows and 25 columns, one row per STAR-Counts expression file in the GDC manifest of each of the four projects. The 25 columns are, in groups:

- `cohort` and `role`, the project and its part in the study.
- `patient`, `sample_barcode` and `sample_type`, the submitter identifiers and the GDC sample type.
- `file_id`, `file_name`, `gdc_md5sum`, `gdc_file_size`, `gdc_data_release`, `gdc_metadata_fetched`, `gdc_state`, `gdc_workflow_version`, `gdc_created_datetime` and `gdc_updated_datetime`, the GDC record of the file.

- `local_file_present`, `local_file_size`, `local_file_date`, `local_md5sum` and `local_md5_matches_gdc`, the local copy and its verification.
- `assigned_reads`, the STAR library size used throughout the pipeline.
- `retained_in_analysed_cohort` and `exclusion_reason`, whether the file supplied the expression profile of one patient in an analysed cohort and, if not, why not.
- `in_adjusted_models` and `in_prediction_set`, the two downstream analysis sets.

`SupplementaryData1_column_definitions.tsv` is a 25-row data dictionary giving the type, definition and applicable cohorts of every column.

All metadata were read from GDC Data Release 46.0 of 10 August 2026. Local copies exist for 1,491 of the 1,549 files, and all 1,491 md5 checksums match the GDC record. The other 58 files, the subtype normal libraries and one additional new primary, were never downloaded.

| Cohort | Role | Files listed | Patients retained | In adjusted models | In the prediction set |
|---|---|---|---|---|---|
| TCGA-KIRC | discovery | 614 | 528 | 520 | 493 |
| CPTAC-3 | validation | 521 | 200 | 199 | 199 |
| TCGA-KIRP | subtype | 323 | 269 | 263 | not applicable |
| TCGA-KICH | subtype | 91 | 65 | 65 | not applicable |
| All cohorts | n/a | 1,549 | 1,062 | 1,047 | 692 |

Exclusions by reason, TCGA-KIRC: non-primary sample 73, failed library 5, duplicate aliquot 4, non-positive or missing survival time 4. CPTAC-3: non-primary sample 168, duplicate aliquot 92 files from 40 patients, non-clear-cell histology 38, non-positive or missing survival time 23. TCGA-KIRP: non-primary sample 33, non-positive or missing survival time 3, missing age or sex 18. TCGA-KICH: non-primary sample 25, non-positive or missing survival time 1. The all-cohorts patient total of 1,150 in the source table is de-duplicated, because one patient can appear in more than one row. The file count and the retained-patient count are simple sums. Source: `31_supplementary_data1_summary.tsv`.

#### Supplementary Data 2. Module gene lists, module membership and eigengene loadings

Three files. 02_lncRNA_module_genes.tsv (3,442 rows) and 02_mRNA_module_genes.tsv (12,000 rows) give one row per analysed gene. Their columns are gene_id (versioned GENCODE v36 Ensembl identifier), gene_name, module (WGCNA colour label, grey for genes assigned to no module) and kME (correlation of the gene with its own module eigengene).

22_module_loadings.tsv gives one row per gene per module, 10,407 rows over the 29 non-grey modules. Its columns are module (mRNA_ME<colour> or lnc_ME<colour>), gene_id, gene_name, loading (element of the first principal component of the module's gene-standardised residualised discovery expression), gene_center (discovery mean of the gene on the log2(FPKM + 1) scale) and gene_scale (its discovery standard deviation). To score a new sample, each gene is standardised with its gene_center and gene_scale, the inner product is taken with loading, and the result is divided by the score_scale in Supplementary Table 20b. The three files fully specify every module eigengene.

#### Supplementary Data 3. Survival results for all 29 modules

03_all_modules_principal.tsv, 29 rows, one per non-grey module of either network under the principal specification. The columns are:

- biotype, module, n_genes, n and events.
- HR, lo and hi, the hazard ratio per standard deviation with its 95% confidence limits, and p.

- `fdr_within`, the Benjamini-Hochberg false discovery rate within the module's own network of 13 or 16 tests, and `fdr_joint`, the same across all 29 modules of both networks.
- `ph_p_ME` and `ph_p_global`, the proportional-hazards p values for the eigengene term and for the global test.

Five modules reach a 5% false discovery rate under either family. This file is also the source of Table 2 of the main text.

##### Supplementary Data 4. Gene Ontology enrichment of the protein-coding modules

`06_GO_enrichment_all_modules.tsv`, 472 rows, one per enriched biological-process term per module. The columns are `module`, `ID`, `Description`, `GeneRatio`, `BgRatio`, `pvalue`, `p.adjust`, `Count` and `geneID`, the slash-separated list of module genes annotated to the term. `clusterProfiler::enrichGO` was run for every module significant under the continuity specification, against a universe of the 12,000 protein-coding genes in the network, with Benjamini-Hochberg adjustment and cut-offs of 0.05 for p and 0.10 for q. Three modules return enriched terms: the catabolic module (229 terms), the proliferation module (232) and the splicing module (11).

##### Supplementary Data 5. Pan-cancer file manifest

`38_pancancer_file_manifest.tsv`, 11,505 rows, one per STAR-Counts file of the 33 TCGA projects at Data Release 46.0, with the columns `file_id`, `project`, `file_name`, `md5`, `sample_type`, `group` and `reason` (retained, failed library, duplicate library of a retained patient, or not a primary tumour or solid-tissue normal). Every retained file was verified against its md5 checksum.
